# d-Lysergic acid diethylamide has major potential as a cognitive enhancer

**DOI:** 10.1101/866814

**Authors:** Felipe Augusto Cini, Isis Ornelas, Encarni Marcos, Livia Goto-Silva, Juliana Nascimento, Sergio Ruschi, José Salerno, Karina Karmirian, Marcelo Costa, Eduardo Sequerra, Dráulio de Araújo, Luis Fernando Tófoli, César Rennó-Costa, Daniel Martins-de-Souza, Amanda Feilding, Stevens Rehen, Sidarta Ribeiro

**Affiliations:** Brain Institute, Federal University of Rio Grande do Norte (UFRN), Natal, Brazil; D’Or Institute, Rio de Janeiro, Brazil; Instituto de Neurociencias de Alicante, Consejo Superior de Investigaciones Científicas-Universidad Miguel Hernández de Elche, San Juan de Alicante, Spain; Institute of Biomedical Sciences, Federal University of Rio de Janeiro (UFRJ), Rio de Janeiro, Brazil; University of Campinas (UNICAMP), Campinas, Brazil; Digital Metropolis Institute, Federal University of Rio Grande do Norte (UFRN), Natal, Brazil; The Beckley Foundation, Oxford, UK

## Abstract

Psychedelic agonists of serotonin receptors induce neural plasticity and synaptogenesis, but their potential to enhance learning remains uncharted. Here we show that a single dose of d-LSD, a potent serotonergic agonist, increased novel object preference in young and adult rats several days after treatment. d-LSD alone did not increase preference in old animals, but could rescue it to young levels when followed by a 6-day exposure to enriched environment (EE). Mass spectrometry-based proteomics in human brain organoids treated with d-LSD showed upregulation of proteins from the presynaptic active zone. A computational model of synaptic connectivity in the hippocampus and prefrontal cortex suggests that d-LSD enhances novelty preference by combining local synaptic changes in mnemonic and executive regions, with alterations of long-range synapses. Better pattern separation within EE explained its synergy with d-LSD in rescuing novelty preference in old animals. These results advance the use of d-LSD in cognitive enhancement.

## Introduction

Normal aging is associated with a decline in cognitive abilities, such as learning capacity, processing speed, working memory, and executive functions (1, 2). Cognitive aging is thought to reflect decreased synaptogenesis in elderly individuals (3). In the 1960s and 1970s, psychedelic substances that are agonists of serotonin receptors were extensively used in psychotherapy because of their potential to cause long-lasting psychological changes (4–7). Decades of prohibitive regulation prevented research on the cognitive benefits of serotonergic agonists, but in the past years psychedelic research is undergoing a major renaissance (8–11), particularly in psychiatric conditions. Agonists of serotonin receptors have been shown to have foremost utility for the treatment of depression (12, 13) and terminal anxiety (14–16). The use of serotonergic psychedelics is associated with decreased psychological suffering and lower suicide rates (17). These substances have also been shown to induce neural plasticity and synaptogenesis both *in vitro* and *in vivo* (18–20), and are therefore likely to enhance learning capacity. Indeed, psychedelics have been shown to enhance memory consolidation when administered after fear learning or novel object exploration (21, 22). In the former, a single dose immediately after learning enhanced memory consolidation in adult mice, while treatment immediately before learning facilitated the extinction of fear memory (21). In the latter, chronic treatment of adult rats with d-LSD for 11 days improved learning in animals deprived of olfaction by bulbectomy, but no effects were observed in sham controls (22). The lesion increases hippocampal 5-HT2A levels, and d-LSD down-regulated these to normal levels, with no effects in sham animals. The main interpretation proposed for these findings was that the post-learning activation of 5-HT2A receptors improves memory consolidation, and promotes positive mood changes. Neither study investigated the effects of d-LSD treatment several days before the learning task. Thus, we hypothesized that d-LSD treatment before a given learning task would increase synaptogenesis, creating a rich synaptic landscape that should favor the encoding of new memories.

## Results & Discussion

To test this hypothesis we first explored how a pre-treatment with d-LSD at various concentrations and time intervals (Figure 1A) would affect subsequent novel object preference (Figure 1B) in young, adult and old rats (**Suppl. Table S1**). The reference dose chosen for treatment (0.13 mg/kg) was determined by the current literature as required for the activation of 5-HT2A receptors, as indicated by the occurrence of wet dog shakes (23). For comparison, we also tested half and triple doses (0.065 mg/kg and 0.39 mg/kg, respectively). The choice of serotonergic agonist – lysergic acid diethylamide (d-LSD) – was determined by its extremely low toxicity and effective dose, which makes d-LSD stand out among all other similar molecules (24).

**Figure 1:**
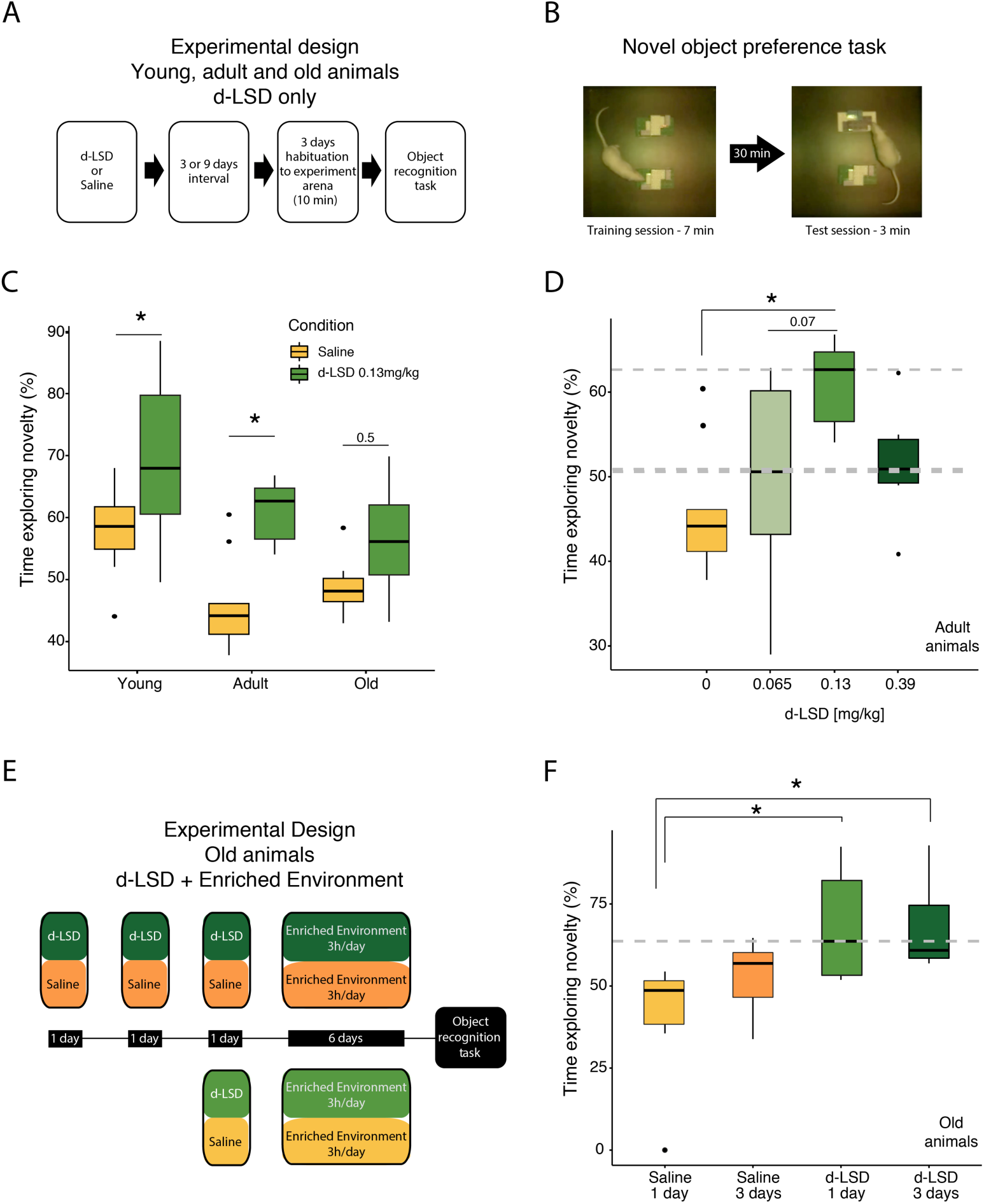
Treatment with d-LSD enhanced novel object preference in adults rats, and old rats showed similar effects when d-LSD was combined with enriched environment exposure. **(A)** Rats of different ages were treated with d-LSD or saline and tested for novel object preference 6 days later. **(B)** Novel object preference was assessed during test sessions with one new object and one familiar object. **(C)** In young and adult rats, but not old animals, a single d-LSD dose significantly increased the preference for novel objects 6 days later. **(D)** In adult rats, the biggest effect was observed for the intermediary dose of 0.13 mg/kg (medians marked by grey dashed lines). **(E)** Experimental design combining d-LSD with EE in old animals. **(F)** Cognitive rescue was obtained in old rats treated with d-LSD treatment and then exposed to EE, as shown by the increased time exploring novelty in this group. # indicates a significant difference from the performance of saline controls unexposed to EE (median marked by grey dashed line).

Using the reference dose, we found that d-LSD treatment significantly increased novel object preference in adults (p = 0.008), from 44.16 ± 7.49% in saline controls to 62.65 ± 5.07% in d-LSD treated animals (median ± sd). In young animals the effect was also detected (p=0.05), with an increase from 58.57 ± 9.06% in saline controls to 67.95 %± 13.13% in d-LSD treated animals (median ± sd; Figure 1C; **Suppl. Tables S2,S3**). When different d-LSD doses were compared in adult animals, a maximum of novel object preference was found for the intermediate dose of 0.13 mg/kg (Figure 1D). Daily doses of 0.13 mg/kg for 3 consecutive days were not as effective as the single dose; nor did a single application of a triple dose (0.39 mg/kg) produce more gain with a longer interval (**Suppl. Figure S1A**). Importantly, old animals did not display differences in novel object exploration after a single d-LSD dose (Figure 1C). Moreover, the efficiency of d-LSD in increasing explorative behavior correlated negatively with both weight and age (**Suppl. Figures S1B,C; Suppl. Table S4**).

Since exposure to an enriched environment (EE) enhances learning & memory in aging rodents (25, 26), we hypothesized that d-LSD treatment followed by several days of EE exposure could promote a cognitive rescue of the old animals. To test this idea we treated old rats with 0.13 mg/kg of d-LSD for 1 or 3 days, followed by 3h of EE exposure for 6 days (Figure 1E). Both combinations of d-LSD treatment with EE led to a significant increase from baseline levels (i.e. saline controls without EE) (Figure 1F). Treatment with d-LSD for 1 day followed by EE also led to significantly more novel object exploration than saline followed by EE; a non-significant trend in the same direction was detected for 3 days of d-LSD treatment followed by EE (Figure 1F). The different age groups did not show significant differences in the total time exploring objects (**Suppl. Figure S1D**). Overall the data indicate that a single dose of d-LSD increases cognition whenever plasticity is present, but is insufficient to rescue cognitive losses caused by ageing. However, it also indicates that once brain plasticity is by other means stimulated in old rats, d-LSD can boost the effect.

To investigate the molecular mechanisms underlying the cognitive benefits of d-LSD treatment, we tested whether d-LSD is able to alter expression of genes associated with brain plasticity *in vitro*. While this has been demonstrated in rats both *in vitro* and *in vivo* (20), similar effects remain to be shown in human neurons. One of the greatest caveats for studying cellular and molecular effects of any psychedelic compound is the relative difficulty of obtaining brain cells from human donors. To overcome this limitation, we used human induced pluripotent stem cells (iPSCs) to generate brain organoids that recapitulate some aspects of patterning, organization, and connectivity observed in the embryonic human brain (27).

Brain organoids generated from human iPSCs were grown in agitation and used for the experiments at day 45, when they already express most of the transcription factors and a protein profile consistent with differentiated brain regions (28). Organoids were treated for 24 h with 10 nM d-LSD and processed by mass spectrometry-based proteomics in order to systematically approach the lysergic effects in human neural cells. Liquid chromatography–mass spectrometry (LC-MS) based proteomics of control and d-LSD identified 3,448 proteins in 3 biological replicates (**Suppl. Table S5**). We used a String network to analyze a total of 234 proteins with expression significantly modified by d-LSD treatment (p<0.05). This tool allows searching for experimental and correlational data connecting proteins in an interaction network (Figure 2A). From this network we identified enriched categories of cell components, such as the term ‘neuron part’ (red circles, 14% of hits, *q* = 0.01), and the term ‘synapse’ (blue circles, 9 % of hits, *q* = 0.02).

**Figure 2:**
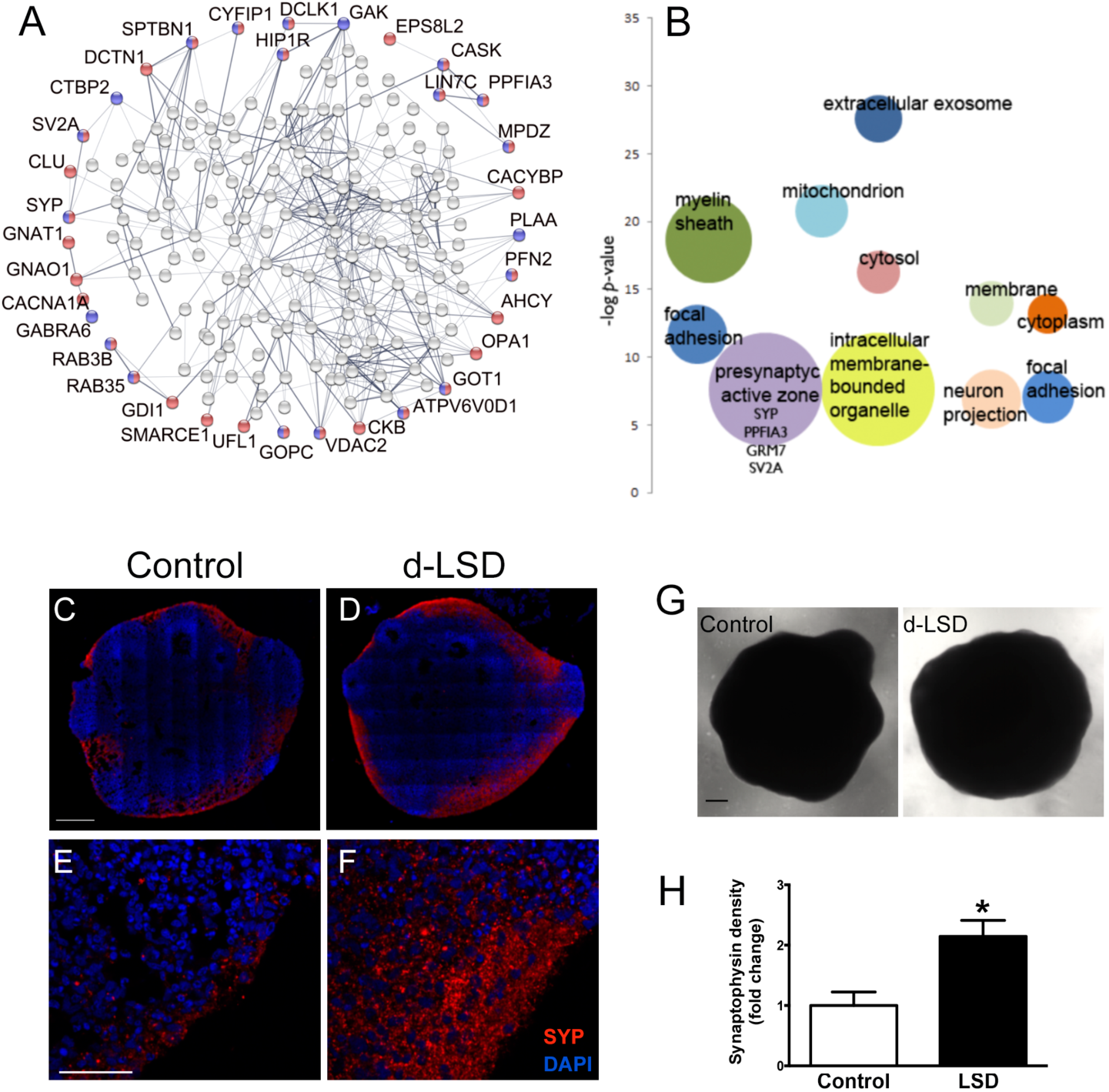
d-LSD regulated synaptic protein expression in human brain organoids. Brain organoids derived from human iPSCs were grown for 45 days and treated with 10 nM d-LSD for proteomic analysis. **(A)** String network analysis showing neuron part (red, q=0.01) and synapse (blue q=0.02) enriched proteins in the dataset. **(B)** Cell compartment gene ontology categories enriched in proteomic hits from the dataset (p<0.05). Presynaptic active zone proteins are listed. –log p-value of enriched categories and fold increase enrichment (bubble size) are plotted. **(C-F)** Representative images of SYP (red) and DAPI (blue, nuclear) staining of control and d-LSD-treated organoids. The top panels (C,D) show a section of the whole organoid and the bottom panels (E,F) represent magnified pictures. **(G)** Phase contrast images of control and d-LSD-treated organoids displaying similar morphology and appearance. **(H)** Quantification of SYP intensity expressed as fold change of control. Plot represents the mean ± SEM, *p<0.05. Organoids were collected from 4 independent experiments and at least 4 organoids per condition. Scale bar = 250 μm in C,D, G and 50 μm in E, F.

In addition, gene ontology (GO) enrichment analysis, using the DAVID bioinformatics tool shows other ‘cell compartment’ terms enriched (Figure 2B). See highlighted the enrichment for the term ‘presynaptic active zone’ (*p*= 0,005; *q*<0.2). The proteins synaptophysin (SYP = 3.63 ± 0.92), PTPRF interacting protein alpha 3 (PPFIA3 = 1.28 ± 0.38), glutamate metabotropic receptor 7 (GRM7 = −2.4 ± 0.55), and synaptic vesicle glycoprotein 2A (SV2A = 1.17 ± 0.29) are some of the proteins present in this enriched GO category (values expressed as normalized fold change ± SD). Typical neuronal markers like class III β-tubulin, MAP2 and Tbr1 did not show a significant difference in their expression (**Suppl. Table S4**), suggesting that the increase in synaptic proteins was not due to an increase in the number of neurons.

SYP is a major integral membrane protein that binds to synaptobrevin and is ubiquitously expressed in synapses throughout the mammalian brain (29). Synapse formation is one of the main neuroplastic changes and correlates with cognition. Interestingly, SYP stands out as a highly upregulated protein within the categories of the GO analysis. In order to confirm whether d-LSD was affecting synaptic proteins, and SYP in particular, we performed an immunofluorescence staining for SYP in brain organoids kept in the presence or absence of d-LSD for 4 days. SYP was mainly expressed in the part of the organoid that resembles the structural organization of the cortical plate (27), the preferential location of the neurons (Figure 2C-F). d-LSD induced a more than 2-fold increase in the levels of SYP in the cortical plate-like region of the human brain organoids (Figure 2H). These data suggest that d-LSD stimulates young neurons to increase the expression of SYP, thus becoming more prone to form additional new synapses.

To understand the association of SYP up-regulation with cognitive enhancement, we manipulated the proprieties of synaptic connectivity in a composite model of the hippocampus (30) and the prefrontal cortex (31, 32) used to emulate the novel object preference task (Figure 3A). Altering the balance of pattern separation and completion in the hippocampus (Figure 3B), the internal connectivity of prefrontal cortex (Figure 3C), and the strength of coupling between these areas (Figure 3D) impacted the estimated preference for novelty independently and allowed for an interpretation of the differences in the behavioral results of adult and old rats (Figure 3E). The use of long-range connectivity as a proxy for age (33, 34) provided an explanation for age differences in baseline performance and cognitive enhancement (Figure 3D): Young rats start at a high baseline level, but still with room for improvement; adult and old rats start at neutral performance level, but with the connectivity levels for adult rats sitting next to an inflection and for old rats at a plateau. A small increase in connectivity within the local circuits of the prefrontal cortex further impedes the cognitive enhancement in old rats (Figure 3C). Superior pattern separation in an enriched environment (35) does not affect baseline performance level for old rats but amplifies the gain added for extra connectivity (Figure 3B).

**Figure 3:**
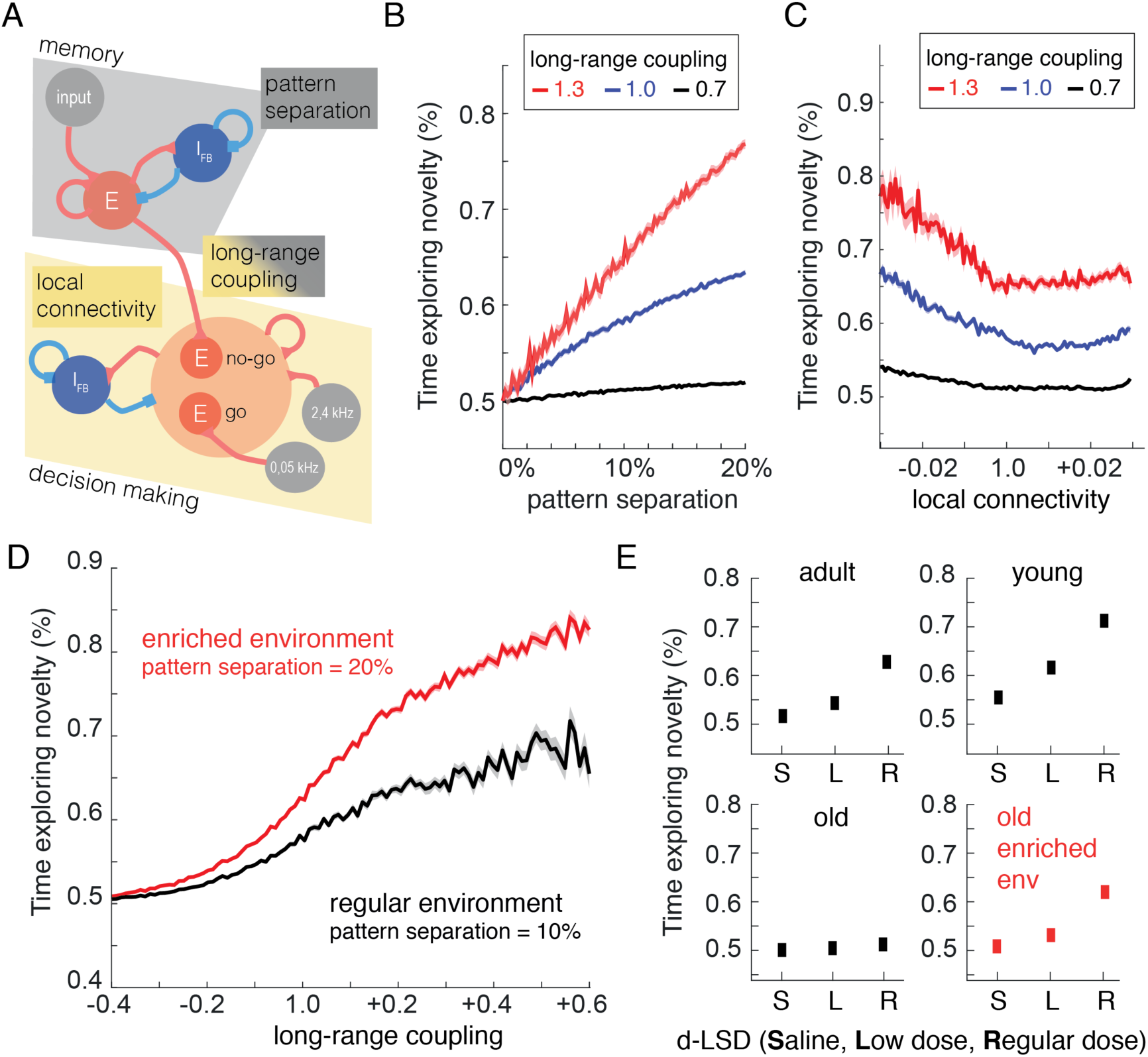
Composite mechanism for d-LSD associated behavioral alterations. (**A**) Cortico-hippocampal model. The mnemonic circuit outputs the level of novelty of an input pattern. Familiar patterns serve as a reference in training cycles before testing. The novelty signal feeds the downstream decision circuit through a magnified long-range synapse. The decision circuit promotes a convergent competitive process between two neuronal pools, one excited by a constant signal that biases the network towards exploration and the other by the novelty signal. The pools are, respectively, associated with the decision to explore or not. The simulations include multiple trials with novel or familiar inputs, allowing an estimation of the time exploring novelty. Time exploring novelty reported as a function of (**B**) hippocampal pattern separation; (**C**) local cortical connectivity; and (**D**) long-range connectivity between hippocampus and cortex. (**E**) The behavioral results were replicated by considering long-range connectivity as a proxy for age (baseline of +0.2 for young, −0.1 for adult and −0.4 for old) and SYP level (increase of +0.1 for low dose and +0.3 for regular dose); and increase of pattern separation as a surrogate of environmental enrichment (10% for normal and 20% for enriched).

The results indicate that d-LSD can substantially increase synaptic plasticity in human neurons *in vitro*, and novelty preference in behaving rats. Altogether, they support the use of d-LSD to enhance learning, chart the molecular pathways underlying this effect, and show for the first time that d-LSD modulates synaptic proteins in human neurons. Of note, the computational modeling of synaptic connectivity in mnemonic and executive brain regions demonstrated that superior pattern separation in the EE is sufficient to explain its synergy with d-LSD in the cognitive rescue of old animals.

The results also suggest that the potent anti-depressant effects of serotonergic psychedelics (12–16, 36–38) may stem in part from a boost in synaptogenesis that increases learning capacity and drives curiosity outwards. The same mechanisms make this class of drugs one of the greatest promises among cognitive enhancers (39, 40), side by side with Δ-9-tetrahydrocannabinol, which also involves SYP (41), and likely overlap with the mechanisms of sleep-induced synaptogenesis (42). Future studies must determine how to optimize the action of serotonergic psychedelics so as to rescue the learning deficits brought by natural or pathological aging (1–3).

## Support

This project was supported by the Beckley Foundation; Fundação de Amparo à Pesquisa do Estado do Rio de Janeiro, Coordenação de Aperfeiçoamento de Pessoal de Nível Superior, Conselho Nacional de Desenvolvimento Científico e Tecnológico grants 308775/2015-5 and 408145/2016-1, Sao Paulo Research Foundation grants 2013/07699-0 (Neuromathematics), 2014/10068-4, 2017/25588-1 and 2019/00098-7, intramural grants from D’Or Institute and Federal University of Rio Grande do Norte, and a Juan de la Cierva-Incorporación Scholarship (IJCI-2016-27864 from the Spanish Ministry of Science, Innovation and Universities).

## Acknowledgments

We thank the Institute of Chemistry of UFRN for NMR analysis, F. G. Menezes and R. S. Fernandes for analytical characterization support, A. C. Souza for behavioral advice, M. Medeiros and J. Pontes for animal care, D. Costa and I. S. Pereira for documentation support, and A.E.A. Oliveira, G. Santana and K. Rocha for miscellaneous support.

## Author contributions

SR, SR, AF, IO, FAC, DM-S, CR-C, LFT and DA contributed to the design of the work; FAC, IO, LG-S, JN, SR, JS, KK, MC, ES, SR, SR, AF, DM-S, CR-C, LFT and DA contributed to the acquisition, analysis, or interpretation of data; EM and CR-C contributed to the creation of new software used in the work; SR, SR, AF, IO, FAC, DM-S, ES, CR-C, LFT and DA have drafted the work or substantively revised it.

The authors declare no competing interests.

## Material & Methods

### Research permits

ANVISA AEP no. 003/2018, ANVISA AEP no. 019/2019 and CEUA UFRN # 011.2015.

### Drug

The study used high-purity d-LSD dissolved in ultrapure water (18.2 MΩ.cm). The solution was manipulated at room temperature and light exposure was avoided. The analytical characterization is described on **Suppl. Information**.

### Animals

Wistar rats (n=96) on different age groups comprising young (2 months), adult (9 months) and old (12-18 months) animals received 1 or 3 daily intra-peritoneal injections of saline or d-LSD doses of 0.065mg/kg, 0.13mg/kg and 0.39mg/kg (**Suppl. Table S1**). Before the injection the animals were housed with other individuals, after the application of saline or d-LSD the animals were housed individually. For ten hours following the injection, animals were video-recorded and stayed alone during the acute effect of the drug/saline. After that, each rat was housed in cages with 4 more individuals.

### Novel object preference task

The square arena where the novel object preference task took place (44 × 44 × 41 cm) had two 3D printed bases fixed on the floor 11 cm apart, for the firm placement of the objects, which were assembled with the same number and types of LEGO**™** pieces. On the 3 days preceding the task, animals were habituated for 10 min to the task arena. Two identical objects were used for the training session, with 12 edges and 12 vertices each. For the test session, a new object, with 14 edges and 14 vertices, replaced one of the previous objects. Novel and familiar objects had identical height and base area dimensions. At the onset of each session, the animals were always placed facing the same wall of the arena, with their backs towards the objects. The test session was performed 30 min after the end of the training session. Novelty preference was calculated as B/(A+B)*100, where B is the time exploring the new object, and A the time exploring the familiar object. New object preference was analyzed using two-way (Age and Dose) ANOVA, followed by Tukey HSD (post-hoc analysis suited to compare group with different sample sizes).

### Enriched Environment (EE)

Old animals were placed in an EE for 3h every day, during 6 days. On the last 3 days, they were also habituated for 10 min to the arena used in the novel object preference task. The EE was made of card boxes with 8 different rooms, in which it was possible to find PVC tubes, toilet paper cardboard tubes, wood-made objects and hidden fruit loops to stimulate exploratory behavior. Animals explored the environment freely in groups of 3-5 animals. The effect of d-LSD followed by EE was analyzed using one-way ANOVA, followed by Tukey HSD. The statistical analyses were performed using open source programming language R (Version 3.6.0). The Mann-Whitney U test was used to compare the performance of old animals exposed to EE to that of old animals treated with saline and unexposed to EE.

### Brain organoid preparation

GM23279A, an induced pluripotent cell (iPSC) line from the NIGMS Human Genetic Cell Repository obtained and certified by the Coriell cell repository was used in this study. iPSCs were maintained with mTeSR1 medium and upon confluence passaged manually or with EDTA. iPSC cultures with no differentiated cells were used to prepare brain organoids following a previously described protocol (28). Shortly, iPSCs were detached using Accutase for 5 min at 37°C, detached cells were then added to phosphate-buffered saline (PBS; LGC Biotechnology, USA) containing 10μM ROCK inhibitor (ROCKi, Y27632; Merck Millipore, USA) to a final concentration of 10 μM and dissociated to single cells. Cells were centrifuged at 300 rpm for 5 min and pellet was resuspended in hESC medium (20% knockout serum replacement; Life Technologies), 3% ESC-quality fetal bovine serum (Thermo Fisher Scientific, USA), 1% GlutaMAX (Life Technologies, Canada), 1% minimum essential medium non-essential amino acids (MEM-NEAAs; Life Technologies), 0.7% 2-mercapto-ethanol, and 1% penicillin-streptomycin (P/S; Life Technologies). 9,000 cells were plated per well of an ultra-low attachment 96 well plate in hESC medium containing 50 µM ROCKi and 4 ng/ml b-FGF. After plating the plate was spinned at 300 rpm for 1 min. On days 3 and 5 medium was changed. At this stage, embryoid bodies (EBs) were formed and grew larger. On day 7 EBs were transferred to ultra-low attachment 24 well plates and media was changed to neuroinduction media [1% N2 supplement (Gibco), 1% GlutaMAX (Life Technologies), 1% MEM-NEAAs, 1% P/S, and 1 μg/ml heparin in DMEM/F12 (Life Technologies)]. The neuroinduction media was changed at day 9 and at day 11 the tissue was submitted to 1 h Matrigel bath. After that, media was changed to differentiation media minus vitamin A (50% neurobasal medium, 0.5% N2, 1% B27 supplement without vitamin A, 1:100 2-mercapto-ethanol, 0.5% MEM-NEAA, 1% GlutaMAX, and 1:100 P/S in DMEM/F12) and on day 13 the media was changed. On day 15 organoids were transferred to agitation in 6 well plates at 90 rpm, media was changed to differentiation media plus vitamin A (50% neurobasal medium, 0.5% N2, 1% B27 supplement with vitamin A, 1:100 2-mercapto-ethanol, 0.5% MEM-NEAA, 1% GlutaMAX, and 1:100 P/S in DMEM/F12), and replaced every 4 days until day 45.

### Liquid-chromatography mass spectrometry (LC-MS)

45-day old human brain organoids (four to five per condition) were treated for 24 hours with 10 nM d-LSD in water, or only medium (control). The organoids were collected from three independent experimental batches. Organoids were lysed in buffer containing 7 M Urea, 2 M thiourea, 1% CHAPS, 70 mM DTT, and Complete Protease Inhibitor Cocktail (Roche). Supernatant protein extracts (50 µg) were digested *in gel* with trypsin (1:100, w/w) overnight. Peptides were injected into a reverse-phase liquid chromatographer [Acquity UPLC M-Class System (Waters Corporation, USA)], coupled to a Synapt G2-Si mass spectrometer (Waters Corporation, USA). Data was acquired with Data-Independent Acquisitions (DIA), with ion mobility separation (high-definition data-independent mass spectrometry; HDMS^E^) (43). Peptides were loaded in first-dimension chromatography onto an M-Class BEH C18 Column (130 Å, 5 µm, 300 µm X 50 mm, Waters Corporation, Milford, MA). Three discontinuous fractionation steps were performed (13%, 18%, and 50% acetonitrile). Peptide loads were directed, after each step, to second-dimension chromatography on a nanoACQUITY UPLC HSS T3 Column (1.8 µm, 75 µm X 150 mm; Waters Corporation, USA). Peptides were then eluted using a 7 to 40% (v/v) acetonitrile gradient for 54 min at a flow rate of 0.4 µL/min directly into a Synapt G2-Si. MS/MS analyses were performed by nano-electrospray ionization in positive ion mode nanoESI (+) and a NanoLock Spray (Waters Corporation, Manchester, UK) ionization source. The lock mass channel was sampled every 30 sec. [Glu1]-Fibrinopeptide B human (Glu-Fib) solution was used to calibrate the mass spectrometer with an MS/MS spectrum reference, using the NanoLock Spray source. Samples were all run in technical duplicates of biological triplicates.

### Database search and quantification

All HDMS^E^ raw files were processed for identification and quantification using Progenesis^®^ QI for proteomics version 4.0, including software package with Apex3D, peptide 3D, and ion accounting informatics (Waters). Search algorithms and cross-matched with the Uniprot human proteome database, version 2018/09 (reviewed and unreviewed), with the default parameters for ion accounting and quantitation. To assess the false-positive identification rate, reversed database queries were appended to the original database. Protein and peptide level false discovery rate (FDR) were set at 1%. Digestion by trypsin allowed a limit of one missed cleavage, and methionine oxidation was considered a variable modification and carbamidomethylation (C), a fixed modification. Identifications that did not satisfy these criteria were rejected. The software starts with LC-MS data loading, then performs alignment and peak detection, which creates a list of interesting peptide ions (peptides) that are explored within Peptide Ion Stats by multivariate statistical methods; the final step is protein identity. Relative quantitation of proteins used the Hi-N (3) method of comparison of peptides (44).

### Data analysis

Further statistical analyses of the list with identified proteins and normalized intensity data was performed using the Perseus software (45, 46). Fold change of the intensities assigned to each protein was calculated within each experimental batch. Fold change values were normalized using the vsn package in RStudio (47, 48). Statistical analyses were performed with one sample t-test (*p*<0.05) against 0 value (no change). Significant hits were subjected to gene enrichment using String (49) and DAVID (50) online bioinformatics tools.

### Immunostaining

Brain organoids were treated with 10 nM d-LSD for 4 days, and fixed overnight in 4% paraformaldehyde (Sigma-Aldrich, USA). Organoids were then incubated in 30% sucrose for dehydration, frozen in optimal cutting temperature compound on dry ice, and stored at −80°C. The organoids were sectioned (20 μm thickness) with a cryostat (Leica Biosystems, Germany). For immunofluorescence slides were thawed for 30 min, washed in PBS and permeabilized in 0.3% Triton X-100 in PBS for 15 min. Sections were blocked in a solution containing 3% BSA in PSB for 1 h. Primary antibody anti-synaptophysin (1:200, Millipore, #MAB368) was diluted in blocking solution and incubated at 4°C overnight. After primary incubation, slides were washed in PBS 3 times for 5 min. Sections were then incubated in AlexaFluor secondary antibody goat anti-mouse (1:400, Invitrogen, #A-11003) for 1 hour at room temperature and washed 3 times for 5 min in PBS. For nuclear staining, sections were incubated with DAPI for 5 min. Slides were washed again 3 times for 5 min in PBS and then cover-slipped with Aqua-Poly/Mount (Polysciences, Inc). Images were acquired using a Leica TCS SP8 confocal microscope. For quantification, images were taken from 4-6 different fields of the borders of each brain organoid. Within an experiment 4-5 organoids per condition (control and d-LSD) were evaluated. Organoids were collected from 4 independent experiments. Image J software was used for the analysis. For each section the area of the tissue was delineated and integrated density used as parameter to estimate SYP expression. Statistical testing was performed using two-tailed t-test with GraphPad Prism 6 software. Statistical significance was defined as p < 0.05.

### Computational model

The architecture of the neural network included two modules: a mnemonic circuit, inspired by the hippocampus to compute the level of novelty of an input pattern, and a decision circuit, modeled with the prefrontal cortex as a reference to decide to explore or not to explore. The mnemonic circuit projects the spiking activity of 800 neurons to the decision circuit. Independent Poisson processes determine the activity of the neurons. On familiar trials, the basal firing rate is 50 Hz. Novelty is encoded as a reduction of the basal firing rate with a standard value of 45 Hz (−10%). In some experiments, the firing rate for novelty trials was changed to emulate variations in the level of pattern separation. In emulation of the enriched environments, we reduced the firing rate to 40Hz (−20%). The projection of the mnemonic circuit to the decision circuit is subject to a modulation factor (standard value of 1.0), analog to the strength of long-range synapses connecting the two areas. The strength of this connection can be modified. The decision circuit comprises 2000 leaky integrate-and-fire neurons (80% excitatory and 20% inhibitory) and represents pools of neurons in the prefrontal cortex that implement decision-making mechanisms. All parameters are identical to those presented in a previous work (31) except for the weight of the connections, which are changed during simulations. The network has all-to-all connectivity via three types of receptors: AMPA, NMDA, and GABA_A_. All neurons in the network receive an external input via excitatory connections (AMPA mediated) that simulates the background activity with a mean rate of ν_ext_ = 2.4kHz, following an independent Poisson process for each neuron. Of the excitatory neurons, 240 (15%) form a non-exploratory pool that receives input from the mnemonic circuit. Other 240 excitatory neurons form an exploratory pool that receives an additional input of constant firing rate calculated as the average hippocampal input during training with familiar patterns. Neurons sharing same external input have recurrent connections of w_+_= 1.7 and connect to other excitatory neurons with w_−_= 1 - f(w+-1)/(1-f), with f = 0.15 (31). Inhibitory neurons have recurrent connections and connect to excitatory neurons with unitary strength. The two pools compete with each other through shared recurrent inhibitory connections mediated by the interneurons. All connections within the decision circuit could also be altered to allow the manipulation of decision dynamics. In the model, a decision occurred when the difference in mean activity between the exploratory and non-exploratory pool was above 12 Hz (32). To estimate the time exploring novelty, we computed the proportion of novel trials in which a decision was made towards exploration compared to all cases, i.e., novel and familiar pattern simulations. We simulated 100 blocks of 100 trials per condition (connectivity variation and novel/familiar pattern presented).

