## Supplemental Information for "d-Lysergic acid diethylamide has major potential as a cognitive enhancer"

### Supplementary Information

Analytical Characterization of d-LSD: An Agilent 1260 (Ontario, Canada) and Shimadzu (Kyoto, Japan) high-performance liquid chromatography systems coupled with UV-Vis detector, Fluorescence Detector (Ontario, Canada) and a micrOTOF II (Bruker Daltonics, Billerica, MA, USA) with an electrospray ion (ESI) source were used for the characterization analysis. Purity chromatographic scans for UV-Vis and FLD were achieved using a Poroshell 120 EC-C18 column (50 mm × 3.0 mm, 2.7 μm, Agilent Technologies, Ontario, Canada) and a Kromasil C-18 column (250 mm × 4.6 mm, 5.0 μm, Kromasil, Bohus, Sweden) was employed for High-resolution electrospray ionization mass spectrometry (HRESIMS) analysis. The mobile phase consisted of 0.1% formic acid in ultrapure water and methanol HPLC-grade. The exploratory gradient was performed to elution. For UV-Vis/FLD spectra acquisition, the flow rate of the mobile phase was 0.5 mL/min for 7 minutes, and the temperature of 30 °C. The injection volume was 1 μL. The components of the d-LSD solution were monitored based on peaks at 310 nm to UV-Vis. For FLD detector was set the wavelength 325 and 425 nm for excitation and emission respectively. For HRESIMS, the gradual elution lasted 60 minutes. The injection volume was 20 μL. The analysis parameters were the capillary 4.5 kV, ESI in positive mode, final plate offset 500 V, 40 psi nebulizer, dry gas ($N_2$) with a flow rate of 8 mL/min and a temperature of 300 °C. $^1$H NMR spectra were recorded on Bruker Avance DPX 300 MHz. Chemical shifts are reported relative to chloroform (δ = 7.26). IR spectra were measured on a Shimadzu IRAffinity-1 FTIR Spectrophotometer (Shimadzu, Japan) and a horizontal ATR sampling accessory. All spectra were recorded within a range of 4000-700 $cm^{-1}$ with a 4 $cm^{-1}$ resolution. Each spectrum was calculated as the average of 32 scans and subjected to background subtraction.

d-LSD: 99.3% and 97.8% of the total area on HPLC analysis for UV-Vis and FLD detectors respectively. $^1$H NMR (300 MHz, $CDCl_3$) δ 7.97 (s, 1H), 7.22-7,17 (m, 3H), 6.93 (s, 1H), 6.36 (s, 1H), 4.09-4.01 (m, 1H), 3.63-3.54 (m, 1H), 3.52-3.40 (m, 5H), 3.29-3.21 (m, 1H), 3.13-3.03 (m, 1H), 2.92-2.81 (m, 1H), 2.74 (s, 3H), 1.25 (t, 3H), 1.17 (t, 3H). IR ($\nu_{max}/cm^{-1}$): 3262, 2972, 2932, 1622, 1447, 1342,

1215, 1107, 1069, 972, 907, 849, 779, 748. HRESIMS (m/z): calcd. for $C_{20}H_{26}N_3O$ $[M+H]^+$ 324.2076; found: 324.2063.
