## Supplemental Figure S1 for "d-Lysergic acid diethylamide has major potential as a cognitive enhancer"

### Supplementary Figure

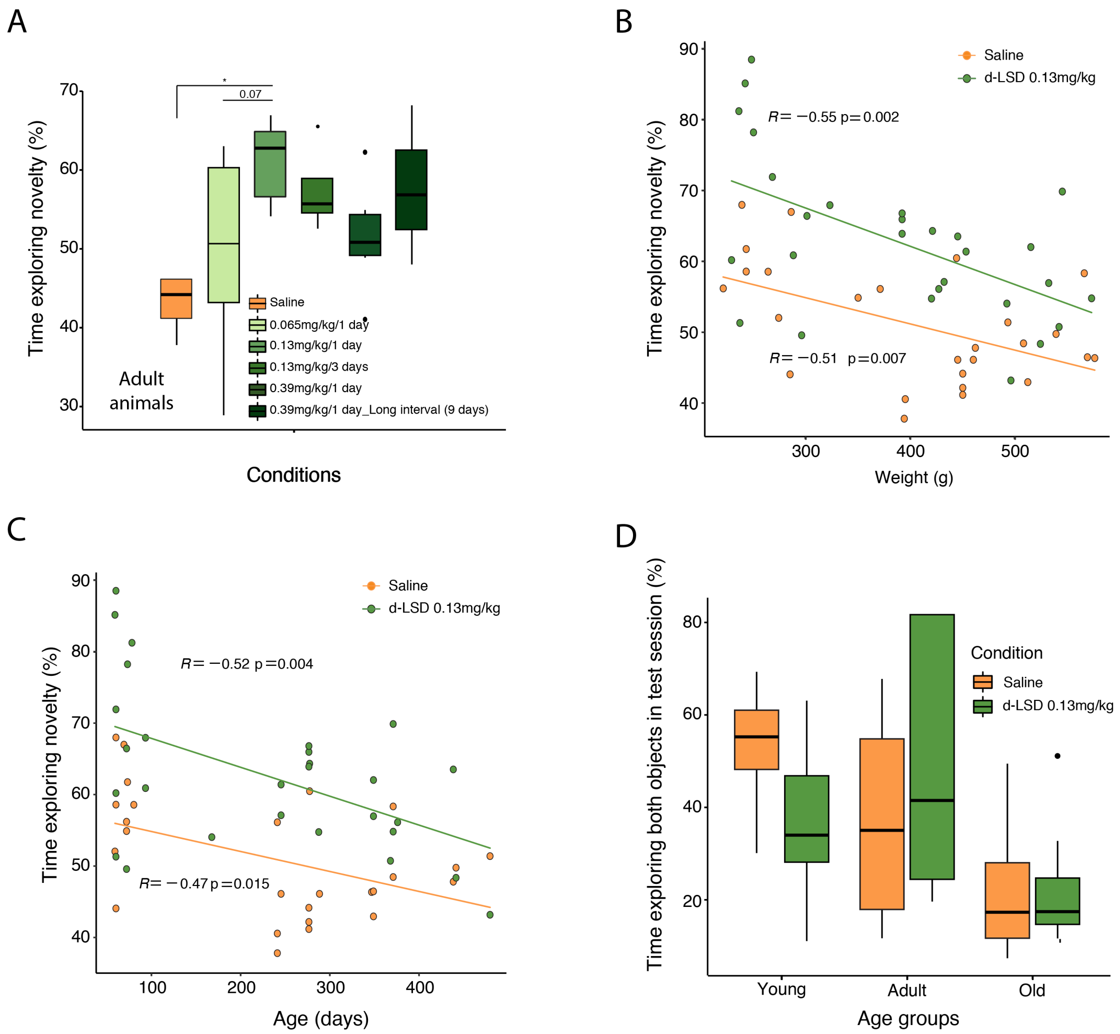


**Supplementary Figure S1: Novel object exploration was enhanced after d-LSD treatment. (A)** In adult rats, maximal cognitive enhancement was observed 6 days after a single dose of d-LSD 0.13 mg/kg. **(B)** Time exploring novelty was negatively correlated with weight; d-LSD treatment increased novelty preference across the entire weight range. **(C)** Time exploring novelty was negatively correlated with age; d-LSD treatment increased novelty preference across the entire age range . (D) Total time exploring objects did not differ across age groups nor between d-LSD and saline treated animals.
