## Supplemental Table S1 for "d-Lysergic acid diethylamide has major potential as a cognitive enhancer"

**Supplementary Table S1**

| Group | N | Mean age (months) | SD age | Median age (months) | Mean weight (g) | SD weight | Median weight (g) | Sex |
|---|---|---|---|---|---|---|---|---|
| d-LSD 0.065 mg/kg_1 day_Adult | 5 | 9,38 | 1,24 | 9,93 | 389,50 | 56,81 | 372,00 | M |
| d-LSD 0.13 mg/kg_1 day_Adult | 8 | 8,51 | 1,30 | 9,17 | 424,25 | 35,10 | 420,50 | M |
| d-LSD 0.13 mg/kg_1 day_Old | 9 | 13,05 | 1,56 | 12,30 | 511,00 | 47,73 | 524,00 | M |
| d-LSD 0.13 mg/kg_1 day_Young | 11 | 2,36 | 0,43 | 2,40 | 265,27 | 31,81 | 250,00 | M |
| d-LSD 0.13 mg/kg_3 days_Adult | 4 | 8,17 | 0,71 | 8,30 | 371,67 | 11,59 | 370,00 | M |
| d-LSD 0.39 mg/kg_1 day_Adult | 6 | 8,86 | 1,16 | 9,53 | 430,25 | 33,00 | 444,50 | M |
| d-LSD 0.39 mg/kg_1 day_Adult_long interval | 3 | 9,09 | 0,04 | 9,07 | 430,00 | 46,89 | 423,00 | M |
| Saline_1 day_Adult | 8 | 9,07 | 0,48 | 9,17 | 449,83 | 5,67 | 450,00 | M |
| Saline_1 day_Old | 8 | 13,04 | 1,72 | 12,30 | 528,13 | 41,03 | 525,50 | M |
| Saline_1 day_Young | 10 | 2,24 | 0,26 | 2,30 | 267,22 | 38,21 | 264,00 | M |
| Saline_3 days_Adult | 4 | 8,17 | 0,71 | 8,30 | 371,00 | 23,07 | 363,00 | M |
| **Enriched Environment** | | | | | | | | |
| d-LSD 0.13 mg/kg_1 day_Old | 6 | 16,12 | 1,34 | 15,68 | 266,00 | 27,27 | 267,00 | F |
| d-LSD 0.13 mg/kg_3 days_Old | 5 | 16,38 | 1,45 | 16,10 | 269,83 | 17,63 | 262,50 | F |
| Saline_1 day_Old | 5 | 16,00 | 1,28 | 15,40 | 271,14 | 29,17 | 274,00 | F |
| Saline_3 days_Old | 4 | 16,75 | 1,57 | 16,87 | 265,14 | 19,36 | 272,00 | F |
