## Supplemental Table S2 for "d-Lysergic acid diethylamide has major potential as a cognitive enhancer"

**Supplementary Table S2**

**One Way Anova comparing old animals exposed to enriched environment on the amount of novel object exploration (%) (d-LSD 0.13mg/kg 1 day and Saline)**

| | Df | Sum Sq | Mean Sq | F Value | Pr(>F) |
|---|---|---|---|---|---|
| Dose | 4 | 3353 | 838,4 | 3,505 | 0,0242 |
| Residuals | 21 | 5023 | 239,2 | | |

---
Signif. codes: 0 '***' 0.001 '**' 0.01 '*' 0.05 '.' 0.1 ' ' 1

Tukey multiple comparisons of means
95% family-wise confidence level

Fit: aov(formula = B_ratio ~ Dose, data = Old with )

| Condition | diff | lwr | upr | p |
|---|---|---|---|---|
| d-LSD 0.13 mg/kg 3days - d-LSD 0.13 mg/kg 1 day | 0,058474 | -29,0803 | 29,19721 | 1 |
| Saline - d-LSD 0.13mg/kg 1 day | -28,8161 | -56,7143 | -0,9179 | 0,040734 |
| Saline 1day/noEE - d-LSD 0.13mg/kg 1 day | -17,5259 | -46,6647 | 11,61279 | 0,404056 |
| Saline 3days - d-LSD 0.13mg/kg 1 day | -16,2548 | -45,3935 | 12,88397 | 0,477128 |
| Saline - d-LSD 0.13 mg/kg 3days | -28,8746 | -56,7728 | -0,97638 | 0,040201 |
| Saline 1day/noEE - d-LSD 0.13mg/kg 3days | -17,5844 | -46,7232 | 11,55432 | 0,400829 |
| Saline 3days - d-LSD0.13 mg/kg 3days | -16,3132 | -45,452 | 12,82549 | 0,473659 |
| Saline 1day/noEE - Saline | 11,29018 | -16,608 | 39,18839 | 0,748235 |
| Saline 3days - Saline | 12,56135 | -15,3369 | 40,45957 | 0,669599 |
| Saline 3days - Saline 1day/noEE | 1,271173 | -27,8676 | 30,40991 | 0,99993 |

**Two Way Anova -d-LSD 0.13mg/kg 1 day and Saline for novelty preference (Condition and Age)**

| | Df | Sum Sq | MeanSq | F value | Pr(>F) |
|---|---|---|---|---|---|
| Condition | 1 | 1827 | 1827,4 | 25,238 | 7,44E-06 |
| age | 2 | 1426 | 712,8 | 9,845 | 0,000261 |
| Condition:age | 2 | 127 | 63,3 | 0,874 | 0,423883 |
| Residuals | 48 | 3476 | 72,4 | | |

---
Signif. codes: 0 '***' 0.001 '**' 0.01 '*' 0.05 '.' 0.1 ' ' 1

**Post Hoc Tukey HSD - Condition and Age**

Tukey multiple comparisons of means
95% family-wise confidence level

Fit: aov(formula = Novelty Preference ~ Condition + age + age:Condition, data =d-LSD 0.13mg/kg 1 day and Saline)

| | diff | lwr | upr | p |
|---|---|---|---|---|
| d-LSD 0.13mg/kg 1 day:Adults-Saline:Adults | 14,96368 | 2,692176 | 27,23517 | 0,008758 |
| Saline:Old-Saline:Adults | 2,866154 | -9,40534 | 15,13765 | 0,981863 |
| d-LSD 0.13mg/kg 1 day:Old-Saline:Adults | 10,10441 | -1,8007 | 22,00951 | 0,13891 |
| Saline:Young-Saline:Adults | 11,81908 | -0,08602 | 23,72418 | 0,052697 |
| d-LSD 0.13mg/kg 1 day:Young-Saline:Adults | 23,14447 | 11,7934 | 34,49554 | 0,000003 |
| Saline:Old-d-LSD 0.13mg/kg 1 day:Adults | -12,0975 | -24,7248 | 0,529747 | 0,067496 |
| d-LSD 0.13mg/kg 1 day:Old-d-LSD 0.13mg/kg 1 day:Adults | -4,85927 | -17,1308 | 7,412229 | 0,846317 |
| Saline:Young-d-LSD 0.13mg/kg 1 day:Adults | -3,14459 | -15,4161 | 9,126905 | 0,972762 |
| d-LSD 0.13mg/kg 1 day:Young-d-LSD 0.13mg/kg 1 day:Adults | 8,180797 | -3,55398 | 19,91558 | 0,320474 |
| d-LSD 0.13mg/kg 1 day:Old-Saline:Old | 7,238251 | -5,03325 | 19,50975 | 0,50638 |
| Saline:Young-Saline:Old | 8,952927 | -3,31857 | 21,22443 | 0,272796 |
| d-LSD 0.13mg/kg 1 day:Young-Saline:Old | 20,27832 | 8,543541 | 32,0131 | 7,32E-05 |
| Saline:Young-d-LSD 0.13mg/kg 1 day:Old | 1,714676 | -10,1904 | 13,61978 | 0,99807 |
| d-LSD 0.13mg/kg 1 day:Young-d-LSD 0.13mg/kg 1 day:Old | 13,04007 | 1,688997 | 24,39114 | 0,015786 |
| d-LSD 0.13mg/kg 1 day:Young-Saline:Young | 11,32539 | -0,02568 | 22,67646 | 0,050831 |

**Tukey HSD only for factor Age**

Tukey multiple comparisons of means
95% family-wise confidence level

Fit: anova(formula = NoveltyPref ~ Condition + age + age:Condition, data = d-LSD)

| Age | diff | lwr | upr | p |
|---|---|---|---|---|
| Old-Adults | -1,02842 | -8,08712 | 6,030294 | 0,933947 |
| Young-Adults | 10,08176 | 3,292913 | 16,87061 | 0,002189 |
| Young-Old | 11,11018 | 4,321328 | 17,89903 | 0,000717 |

**Tukey HSD only for factor Condition**

Tukey multiple comparisons of means
95% family-wise confidence level

Fit: aov(formula = NoveltyPref ~ Condition + age + age:Condition, data = d-LSD)

$Condition

| | diff | lwr | upr | p |
|---|---|---|---|---|
| d-LSD 0.13mg/kg 1 day - Saline | 11,64254 | 6,982866 | 16,30221 | 7,40E-06 |

**One way ANOVA for Adults with different doses (.065; .13; .39) inverted U for Novelty Preference**

| | Df | Sum Sq | Mean Sq | F-value | P value |
|---|---|---|---|---|---|
| Condition | 5 | 1178 | 235,68 | 3,485 | 0,0138 |
| Residuals | 29 | 1962 | 67,64 | | |

---
Signif. codes: 0 '***' 0.001 '**' 0.01 '*' 0.05 '.' 0.1 ' ' 1

**Post Hoc Tukey HSD**

Tukey multiple comparisons of means
95% family-wise confidence level

Fit: aov(formula = NoveltyPref ~ Condition, data = new)

| Condition | diff | lwr | upr | p |
|---|---|---|---|---|
| d-LSD 0.065mg/kg 1 day - Saline | 3,130393 | -10,8537 | 17,11445 | 0,982535 |
| d-LSD 0.13mg/kg 1 day - Saline | 14,96368 | 2,781254 | 27,1461 | 0,009382 |
| d-LSD 0.13mg/kg 3 days - Saline | 12,13666 | -4,57749 | 28,8508 | 0,262373 |
| d-LSD 0.39mg/kg - Saline | 5,55251 | -7,0822 | 18,18722 | 0,760957 |
| d-LSD 0.039mg/kg Long Wait-Saline | 11,52256 | -5,19159 | 28,2367 | 0,314397 |
| d-LSD 0.13mg/kg 1 day - d-LSD 0.065 mg/kg 1 day | 11,83328 | -2,45951 | 26,12608 | 0,0741 |
| d-LSD 0.13mg/kg 3 days - d-LSD 0.065 mg/kg 1 day | 9,006265 | -9,30316 | 27,31569 | 0,667242 |
| d-LSD 0.39mg/kg - d-LSD 0.065 mg/kg 1 day | 2,422117 | -12,2581 | 17,10232 | 0,995663 |
| d-LSD 0.039mg/kg LongWai t- d-LSD 0.065 mg/kg 1 day | 8,392165 | -9,91726 | 26,70159 | 0,728257 |
| d-LSD 0.13mg/kg 3 days - d-LSD 0.13mg/kg 1 day | -2,82702 | -19,8003 | 14,14628 | 0,995467 |
| d-LSD 0.39mg/kg - d-LSD 0.13mg/kg 1 day | -9,41117 | -22,3868 | 3,564421 | 0,263485 |
| d-LSD 0.039mg/kg LongWait-d-LSD 0.13mg/kg 1 day | -3,44112 | -20,4144 | 13,53218 | 0,988793 |
| d-LSD 0.39mg/kg - d-LSD 0.13mg/kg 3 days | -6,58415 | -23,8849 | 10,71664 | 0,851651 |
| d-LSD 0.039mg/kg LongWait-3 days d-LSD | -0,6141 | -21,0847 | 19,85646 | 0,999999 |
| d-LSD 0.039mg/kg LongWait- d-LSD 0.39mg/kg | 5,970048 | -11,3307 | 23,27083 | 0,895986 |
