## Supplemental Table S3 for "d-Lysergic acid diethylamide has major potential as a cognitive enhancer"

**Supplementary Table S3**

**d-LSD 0.13mg/kg/1day vs saline**

Time exploring novelty (s)

| Condition | age | mean | sd | median | SE |
|---|---|---|---|---|---|
| Saline | Adults | 16,48778 | 8,261676 | 21,19 | 2,753892 |
| Saline | Old | 11,30375 | 8,329244 | 8,185 | 2,944833 |
| Saline | Young | 31,25111 | 9,025564 | 33,21 | 3,008521 |
| d-LSD 0.13mg/kg 1 day | Adults | 31,42375 | 19,74251 | 25,19 | 6,980031 |
| d-LSD 0.13mg/kg 1 day | Old | 12,39556 | 6,602593 | 11,13 | 2,200864 |
| d-LSD 0.13mg/kg 1 day | Young | 24,36273 | 8,27041 | 24,45 | 2,493622 |

Novelty preference (%)

| Condition | age | mean | sd | median | SE |
|---|---|---|---|---|---|
| Saline | Adults | 46,07834 | 7,497176 | 44,1667 | 2,499059 |
| Saline | Old | 48,9445 | 4,5535 | 48,1339 | 1,609905 |
| Saline | Young | 57,89742 | 7,398955 | 58,571 | 2,466318 |
| d-LSD 0.13mg/kg 1 day | Adults | 61,04202 | 5,074933 | 62,65574 | 1,79426 |
| d-LSD 0.13mg/kg 1 day | Old | 56,18275 | 8,197743 | 56,12285 | 2,732581 |
| d-LSD 0.13mg/kg 1 day | Young | 69,22281 | 13,133466 | 67,95017 | 3,959889 |

Time exploring both objects in teste session

| Condition | age | mean | sd | median | SE |
|---|---|---|---|---|---|
| Saline | Adults | 37,17111 | 21,16396 | 35,04 | 7,054654 |
| Saline | Old | 22,22625 | 15,08686 | 17,335 | 5,33401 |
| Saline | Young | 53,50111 | 11,90236 | 55,26 | 3,967454 |
| d-LSD 0.13mg/kg 1 day | Adults | 49,5575 | 28,25082 | 41,51 | 9,988172 |
| d-LSD 0.13mg/kg 1 day | Old | 22,57556 | 12,44237 | 17,45 | 4,147456 |
| d-LSD 0.13mg/kg 1 day | Young | 36,72545 | 14,46603 | 33,99 | 4,361673 |

**OLD exposed to environment enrichment**

Time exploring novelty (s)

| Condition | mean | sd | median | SE |
|---|---|---|---|---|
| d-LSD 0.13mg/kg 1 day | 12,562 | 9,316656 | 8,96 | 4,166535 |
| d-LSD 0.13mg/kg 3 days | 17,702 | 6,455313 | 18,85 | 2,886904 |
| Saline 1 day | 8,07 | 6,80697 | 8,01 | 2,778934 |
| Saline 3 days | 10,862 | 6,149668 | 12,56 | 2,750215 |

Novelty preference (%)

| Condition | mean | sd | median | SE |
|---|---|---|---|---|
| d-LSD 0.13mg/kg 1 day | 68,68111 | 17,96789 | 63,62775 | 8,035485 |
| d-LSD 0.13mg/kg 3 days | 68,73959 | 15,1837 | 60,86535 | 6,790357 |
| Saline 1 day | 39,86499 | 20,62403 | 48,65869 | 8,419724 |
| Saline 3 days | 52,42634 | 12,35442 | 56,8979 | 5,525062 |

Time exploring both objects in teste session

| Condition | mean | sd | median | SE |
|---|---|---|---|---|
| d-LSD 0.13mg/kg 1 day | 18,74 | 14,06595 | 16,82 | 6,290486 |
| d-LSD 0.13mg/kg 3 days | 27,868 | 14,01314 | 30,97 | 6,266869 |
| Saline 1 day | 16,19667 | 12,28561 | 16,095 | 5,015581 |
| Saline 3 days | 21,274 | 10,3215 | 25,66 | 4,615916 |

**Adults inverted U curve**

Time exploring novelty (s)

| Condition | age | mean | sd | median | se |
|---|---|---|---|---|---|
| Saline | Adults | 16,77778 | 8,539644 | 21,19 | 2,846548 |
| d-LSD 0.065mg/kg 1 day | Adults | 10,478 | 3,837222 | 8,96 | 1,716058 |
| d-LSD 0.13mg/kg 1 day | Adults | 31,41875 | 19,806795 | 25,19 | 7,00276 |
| d-LSD 0.13mg/kg 3 days | Adults | 15,63333 | 4,501826 | 14,81 | 2,59913 |
| d-LSD 0.39mg/kg 1 day | Adults | 10,53714 | 6,493088 | 7,99 | 2,454157 |
| d-LSD 0.39mg/kg Long Wait | Adults | 14,81 | 3,715172 | 15,61 | 2,144955 |

Novelty preference (%)

| Condition | age | mean | sd | median | se |
|---|---|---|---|---|---|
| Saline | Adults | 46,07834 | 7,497176 | 44,1667 | 2,499059 |
| d-LSD 0.065mg/kg 1 day | Adults | 49,20874 | 13,716101 | 50,64267 | 6,134027 |
| d-LSD 0.13mg/kg 1 day | Adults | 61,04202 | 5,074933 | 62,65574 | 1,79426 |
| d-LSD 0.13mg/kg 3 days | Adults | 58,215 | 7,733215 | 55,133 | 4,464773 |
| d-LSD 0.39mg/kg 1 day | Adults | 51,63085 | 6,54211 | 50,91912 | 2,472685 |
| d-LSD 0.39mg/kg Long Wait | Adults | 57,6009 | 10,064812 | 56,76364 | 5,810922 |

Time exploring both objects in test session (s)

| Condition | age | mean | sd | median | se |
|---|---|---|---|---|---|
| Saline | Adults | 37,73111 | 21,748238 | 35,04 | 7,249413 |
| d-LSD 0.065mg/kg 1 day | Adults | 22,03 | 6,961368 | 19,45 | 3,113219 |
| d-LSD 0.13mg/kg 1 day | Adults | 49,73 | 28,519877 | 41,51 | 10,083299 |
| d-LSD 0.13mg/kg 3 days | Adults | 27,39 | 10,094459 | 22,1 | 5,828039 |
| d-LSD 0.39mg/kg 1 day | Adults | 19,85857 | 9,879473 | 17,07 | 3,73409 |
| d-LSD 0.39mg/kg Long Wait | Adults | 26,98333 | 10,924167 | 27,5 | 6,307071 |
