## Supplemental Table S4 for "d-Lysergic acid diethylamide has major potential as a cognitive enhancer"

**Supplementary Table S4**

**Pearson Correlations for Novelty preference x Age(days) for group d-LSD 0.13mg/kg 1 day**

Pearson's product-moment correlation

data: d-LSD 0.13mg/kg 1 day - Age and Novelty Preference
t = -3.1281, df = 26, p-value = 0.004303
alternative hypothesis: true correlation is not equal to 0
95 percent confidence interval:
-0.7497273 -0.1861505
sample estimates:
cor
r = -0.522912

**Pearsons Correlation for Novelty preference x Age (days) for group Saline**

Pearson's product-moment correlation

data: Saline Age and Novelty Preference
t = -2.5605, df = 18, p-value = 0.01966
alternative hypothesis: true correlation is not equal to 0
95 percent confidence interval:
-0.78071471 -0.09617787
sample estimates:
cor
r = -0.5167072

**Pearsons Correlation for Novelty preference x Weight (g) for group d-LSD 0.13 mg/kg 1 day**

Pearson's product-moment correlation

data: d-LSD 0.13mg/kg 1 day - Weight and Novelty Preference
t = -3.3976, df = 26 p-value = 0.002198
alternative hypothesis: true correlation is not equal to 0
95 percent confidence interval:
-0.7685825 -0.2287496
sample estimates:
cor
r =-0.5545044

**Pearsons Correlation for Novelty preference x Weight (g) for group Saline**

Pearson's product-moment correlation

data: Saline Weight and Novelty Preference
t = -2.9304, df = 24, p-value = 0.007318
alternative hypothesis: true correlation is not equal to 0
95 percent confidence interval:
-0.7512960 -0.1572468
sample estimates:
cor
r = -0.5133318
