## Supplemental Table S5 for "d-Lysergic acid diethylamide has major potential as a cognitive enhancer"

### Supplementary Table S5

| LSD/control (1) | LSD/control (2) | LSD/control (3) | Average (1,2,3) | SD (1, 2, 3) | T-test p-value | Acession | GeneName |
|---|---|---|---|---|---|---|---|
| 4,58897 | 4,75132 | 4,74052 | 4,74052 | 0,0907759 | 0,0001247 | A0A0C4DGI | TTC7B |
| -1,79896 | -1,89769 | -1,81356 | -1,81356 | 0,0532895 | 0,0002804 | Q96JE7;E9F | SEC16B |
| -1,95527 | -1,88314 | -1,83594 | -1,88314 | 0,0600975 | 0,0003363 | P04844;Q5. | RPN2 |
| -3,38723 | -3,39908 | -3,185 | -3,38723 | 0,1203243 | 0,0004366 | Q13733;E9I | ATP1A4 |
| 1,27254 | 1,36582 | 1,36288 | 1,36288 | 0,0530269 | 0,0005265 | P56545;Q5! | CTBP2 |
| 0,492568 | 0,479468 | 0,451039 | 0,479468 | 0,0212308 | 0,000667 | Q14108;A0. | SCARB2 |
| -2,76453 | -2,97101 | -2,68469 | -2,76453 | 0,1477541 | 0,0009225 | P78332;E9F | RBM6 |
| 1,44083 | 1,56911 | 1,42232 | 1,44083 | 0,0799434 | 0,0009746 | P23786;A0/ | CPT2 |
| 3,78059 | 3,63528 | 3,36151 | 3,63528 | 0,2127961 | 0,0011675 | F8VXU5;Q9 | VPS29 |
| -0,731137 | -0,645236 | -0,702494 | -0,702494 | 0,0437376 | 0,0013253 | P62269;J3J! | RPS18 |
| -1,23097 | -1,38413 | -1,37217 | -1,37217 | 0,0851846 | 0,0013666 | P25789;H0\ | PSMA4 |
| 2,73721 | 2,39606 | 2,54608 | 2,54608 | 0,1709873 | 0,001484 | P48739;A0/ | PITPNB |
| -0,476552 | -0,502663 | -0,544383 | -0,502663 | 0,0342135 | 0,0015093 | P49321;Q5 | NASP |
| 4,58606 | 4,89108 | 4,24869 | 4,58606 | 0,3213307 | 0,0016401 | A0A1B0GU! | CACNA1A |
| 0,267693 | 0,270184 | 0,303532 | 0,270184 | 0,0200114 | 0,0016926 | P13674 | P4HA1 |
| 13,7309 | 13,8574 | 12,1515 | 13,7309 | 0,9504912 | 0,0017117 | P23284 | PPIB |
| 1,85197 | 1,63871 | 1,63268 | 1,63871 | 0,1249028 | 0,0017783 | H3BSG2;O7 | CDH16 |
| 1,31277 | 1,38221 | 1,52188 | 1,38221 | 0,1065024 | 0,0019081 | Q6ZVL6;H0 | KIAA1549L |
| 1,4561 | 1,71386 | 1,5845 | 1,5845 | 0,1288803 | 0,0021973 | Q03938 | ZNF90 |
| 2,26642 | 1,86447 | 2,04412 | 2,04412 | 0,2013518 | 0,0031746 | Q9NRR5 | UBQLN4 |
| 3,20161 | 3,87379 | 3,38803 | 3,38803 | 0,3470209 | 0,0032835 | Q8IV35;H7( | WDR49 |
| 1,73409 | 1,63524 | 1,41486 | 1,63524 | 0,163425 | 0,0034824 | Q9BUJ2;A0. | HNRNPUL1 |
| -1,74981 | -1,43805 | -1,74695 | -1,74695 | 0,1791748 | 0,0039316 | O00178;F5I | GTPBP1 |
| -2,85515 | -3,181 | -3,56039 | -3,181 | 0,3529586 | 0,0040337 | P10412 | HIST1H1E |
| 1,9412 | 1,54334 | 1,79788 | 1,79788 | 0,2015043 | 0,0043369 | Q96H55;A0 | MYO19 |
| 0,44954 | 0,471671 | 0,556533 | 0,471671 | 0,0564784 | 0,0043535 | Q12888;A6 | TP53BP1 |
| 1,96643 | 2,2667 | 2,49074 | 2,2667 | 0,263077 | 0,0045611 | C9K025;P1 | RPL35A |
| 0,629188 | 0,634555 | 0,772806 | 0,634555 | 0,0814128 | 0,0047601 | Q15555;K7I | MAPRE2 |
| -2,74922 | -3,48049 | -3,00867 | -3,00867 | 0,370739 | 0,0047966 | Q12965;H0 | MYO1E |
| -2,3843 | -2,4161 | -2,94386 | -2,4161 | 0,3142847 | 0,0049046 | Q8IWV7;AC | UBR1 |
| -1,70095 | -1,45817 | -1,33652 | -1,45817 | 0,1855398 | 0,005071 | P40227 | CCT6A |
| 3,07051 | 3,82933 | 3,85641 | 3,82933 | 0,4461278 | 0,0051212 | O00411;K7I | POLRMT |
| 1,77047 | 1,37063 | 1,61311 | 1,61311 | 0,2014244 | 0,005342 | Q5T8A7 | PPP1R26 |
| -2,34865 | -1,96105 | -1,85095 | -1,96105 | 0,2614259 | 0,0053587 | O75970;F5I | MPDZ |
| 0,642107 | 0,642386 | 0,79607 | 0,642386 | 0,0888101 | 0,0054218 | P24534;C9J | EEF1B2 |
| 1,38948 | 1,07822 | 1,34183 | 1,34183 | 0,1676522 | 0,0057604 | Q8TF40 | FNIP1 |
| -3,07619 | -3,06462 | -3,83856 | -3,07619 | 0,4435322 | 0,0058739 | Q9UKY7 | CDV3 |
| -0,724875 | -0,564272 | -0,72467 | -0,72467 | 0,0926651 | 0,0062921 | O75947;F5I | ATP5PD |
| -1,86162 | -2,12797 | -1,60479 | -1,86162 | 0,2616044 | 0,0064964 | O60313;C9. | OPA1 |
| -2,52915 | -3,25046 | -2,6049 | -2,6049 | 0,396395 | 0,0066388 | O14976 | GAK |
| -1,6658 | -2,06332 | -1,59428 | -1,6658 | 0,2526974 | 0,0066921 | Q14566 | MCM6 |
| -0,663285 | -0,621068 | -0,810945 | -0,663285 | 0,0996987 | 0,0067237 | Q6UXN9 | WDR82 |
| -0,616695 | -0,745627 | -0,564409 | -0,616695 | 0,0932713 | 0,0069571 | O15075;Q5 | DCLK1 |
| 0,936373 | 1,20546 | 1,23291 | 1,20546 | 0,1638574 | 0,0069983 | P51826;C9J | AFF3 |
| -1,43005 | -1,18541 | -1,59614 | -1,43005 | 0,2066131 | 0,0071428 | P35232;C9J | PHB |
| 2,65831 | 3,27134 | 3,57518 | 3,27134 | 0,467043 | 0,0071658 | Q9GZV4;C9 | EIF5A2 |
| 1,42382 | 1,91939 | 1,72094 | 1,72094 | 0,2494168 | 0,0071987 | E9PK80;E9F | SIRT3 |
| -2,23455 | -3,02294 | -2,68351 | -2,68351 | 0,395461 | 0,0073582 | C9JRR5;F8V | BBS9 |
| 0,431036 | 0,531527 | 0,590255 | 0,531527 | 0,0805172 | 0,0079697 | O75820 | ZNF189 |

| | | | | | | | |
|---|---|---|---|---|---|---|---|
| 1,00611 | 1,35976 | 1,1186 | 1,1186 | 0,1806841 | 0,0079698 | Q68CZ2 | TNS3 |
| 2,41639 | 1,82832 | 2,4433 | 2,41639 | 0,3475512 | 0,0080043 | F2Z2M7 | GALNT10 |
| -1,44454 | -1,31558 | -1,0543 | -1,31558 | 0,1988237 | 0,0080524 | P08195;F5( | SLC3A2 |
| -2,28186 | -2,93136 | -2,22818 | -2,28186 | 0,3914064 | 0,008198 | Q13310;B1. | PABPC4 |
| -3,04776 | -2,93862 | -3,89506 | -3,04776 | 0,5235466 | 0,0083167 | Q8IWZ3;E9 | ANKHD1 |
| 2,02631 | 1,79824 | 1,45159 | 1,79824 | 0,2893917 | 0,0089049 | Q9BQ70;H3 | TCF25 |
| 4,03654 | 3,26085 | 2,93978 | 3,26085 | 0,5638651 | 0,0089792 | Q15257;A6 | PTPA |
| -1,32966 | -0,97718 | -1,31931 | -1,31931 | 0,2005834 | 0,009055 | H0Y3K2;Q6 | UBR3 |
| -1,31819 | -1,04741 | -1,47564 | -1,31819 | 0,2166 | 0,0094045 | P09471;A0/ | GNAO1 |
| 3,24981 | 3,15511 | 4,26453 | 3,24981 | 0,6150119 | 0,0098213 | Q9BZL4 | PPP1R12C |
| -2,48811 | -3,44164 | -3,39656 | -3,39656 | 0,5379797 | 0,0098353 | H3BPZ1;H3 | HACD3 |
| -1,6244 | -1,23619 | -1,19857 | -1,23619 | 0,2357447 | 0,009968 | Q6IAA8;F5( | LAMTOR1 |
| 2,76766 | 2,18247 | 1,93849 | 2,18247 | 0,4261253 | 0,0112855 | P48637;A0/ | GSS |
| -1,48144 | -1,06361 | -1,51754 | -1,48144 | 0,2523019 | 0,0113736 | P26196;Q8I | DDX6 |
| 0,98486 | 1,00415 | 0,700455 | 0,98486 | 0,1700436 | 0,0117811 | P53602;H3I | MVD |
| -1,50315 | -2,18721 | -1,74631 | -1,74631 | 0,3467606 | 0,0119853 | P43246;V9I | MSH2 |
| 3,20637 | 2,80272 | 2,16076 | 2,80272 | 0,5273118 | 0,012268 | Q9H3S7 | PTPN23 |
| -1,17099 | -1,53932 | -1,07529 | -1,17099 | 0,2449998 | 0,0123337 | Q12860;H0 | CNTN1 |
| -0,669935 | -0,867245 | -1,00243 | -0,867245 | 0,167212 | 0,0127573 | P38606;C9J | ATP6V1A |
| -1,47309 | -1,20942 | -0,981385 | -1,20942 | 0,2460676 | 0,0132628 | P51957;E7E | NEK4 |
| 3,33884 | 2,44999 | 3,68947 | 3,33884 | 0,6389192 | 0,0133592 | F5GYQ1;P6 | ATP6V0D1 |
| -1,17007 | -0,938665 | -1,4163 | -1,17007 | 0,2388558 | 0,0134958 | Q96AE4;E9 | FUBP1 |
| -0,755762 | -1,15207 | -0,983006 | -0,983006 | 0,1988645 | 0,0139015 | K7EK33;K7E | DAZAP1 |
| 1,39984 | 2,0029 | 2,11337 | 2,0029 | 0,3840595 | 0,014233 | Q6ZS17;A0. | RIPOR1 |
| 3,35283 | 2,23508 | 2,57824 | 2,57824 | 0,5725838 | 0,0144304 | Q6YN16 | HSDL2 |
| -2,04336 | -3,14218 | -2,73281 | -2,73281 | 0,5553273 | 0,0144367 | A0A0B4J1T. | EPHA6 |
| -2,03663 | -1,5397 | -2,37501 | -2,03663 | 0,4201554 | 0,0146252 | A6NFX8;A6 | NUDT5 |
| 2,209 | 2,48846 | 1,61076 | 2,209 | 0,4483946 | 0,0148212 | Q9H4G0;A( | EPB41L1 |
| 2,77027 | 1,77945 | 2,17175 | 2,17175 | 0,4989739 | 0,016134 | Q9HB71;B2 | CACYBP |
| -0,457472 | -0,30223 | -0,464533 | -0,457472 | 0,0917353 | 0,0164308 | P35080;C9J | PFN2 |
| 1,18929 | 0,994406 | 1,54745 | 1,18929 | 0,2805103 | 0,0165372 | Q96N67;A0 | DOCK7 |
| 1,75693 | 1,64836 | 1,1159 | 1,64836 | 0,3430793 | 0,016839 | J3QRH7 | MARCH10 |
| -0,968557 | -1,5252 | -1,18461 | -1,18461 | 0,2806338 | 0,0170174 | Q01581;D6 | HMGCS1 |
| -2,12753 | -2,96623 | -1,95578 | -2,12753 | 0,540667 | 0,017193 | Q13724;C9. | MOGS |
| -3,08954 | -2,34 | -3,77306 | -3,08954 | 0,7167834 | 0,0177178 | A2RRP1;H0 | NBAS |
| -1,34345 | -1,45994 | -2,05029 | -1,45994 | 0,3789692 | 0,017802 | A2A3F3;A2. | TRPM3 |
| -2,3506 | -2,22543 | -3,3577 | -2,3506 | 0,620746 | 0,0178741 | Q92928 | RAB1C |
| -2,44194 | -1,66554 | -2,68322 | -2,44194 | 0,5317715 | 0,0179044 | P20337 | RAB3B |
| 3,27125 | 3,60998 | 2,23888 | 3,27125 | 0,7141943 | 0,0179046 | Q9NXV6 | CDKN2AIP |
| -1,19953 | -1,29056 | -0,80419 | -1,19953 | 0,2585653 | 0,0179847 | Q15738;C9. | NSDHL |
| -1,42399 | -1,97509 | -1,27661 | -1,42399 | 0,3681726 | 0,0180973 | Q6VY07;B4 | PACS1 |
| 0,990019 | 1,50827 | 1,57736 | 1,50827 | 0,321021 | 0,0181079 | Q08378;A0. | GOLGA3 |
| -1,4855 | -1,04139 | -1,69509 | -1,4855 | 0,3337877 | 0,0182397 | P31150;G5I | GDI1 |
| -1,97806 | -1,33771 | -2,16716 | -1,97806 | 0,4347019 | 0,0183397 | P23526 | AHCY |
| -0,714268 | -1,1684 | -1,01053 | -1,01053 | 0,2305537 | 0,0185229 | P13796;Q5 | LCP1 |
| -1,09502 | -0,708234 | -0,776307 | -0,776307 | 0,2064846 | 0,018685 | O94874 | UFL1 |
| 2,64114 | 1,92216 | 3,15851 | 2,64114 | 0,6209086 | 0,0188504 | Q9BRA2;I3I | TXNDC17 |
| 2,00661 | 1,40418 | 2,32234 | 2,00661 | 0,4664806 | 0,0192884 | F5H5K1;J3K | LRRC37B |
| -1,66262 | -2,75003 | -2,4041 | -2,4041 | 0,5555659 | 0,0193501 | Q14831 | GRM7 |
| -3,25638 | -2,35651 | -3,90957 | -3,25638 | 0,7797883 | 0,0195302 | Q15067;I3L | ACOX1 |
| 2,54436 | 1,85456 | 1,59336 | 1,85456 | 0,4913333 | 0,0195789 | J3KNV4;G3' | ITGA7 |
| 1,1811 | 1,82425 | 1,93004 | 1,82425 | 0,405328 | 0,0196401 | D6RBT3;O7 | NDUFS6 |

| | | | | | | | |
|---|---|---|---|---|---|---|---|
| 0,446858 | 0,715497 | 0,712841 | 0,712841 | 0,1543378 | 0,019723 | Q9H0L4 | CSTF2T |
| 1,93368 | 2,36999 | 3,14437 | 2,36999 | 0,6131614 | 0,0197325 | Q5T6L9;K7E | ERMARD |
| 1,12138 | 1,15765 | 0,709752 | 1,12138 | 0,2487856 | 0,0201601 | P38646;D6I | HSPA9 |
| 4,47054 | 2,95622 | 2,97465 | 2,97465 | 0,8690216 | 0,0203054 | D6REX8 | G3BP2 |
| -1,43302 | -1,93109 | -2,40457 | -1,93109 | 0,4858269 | 0,0206223 | Q14203;E7I | DCTN1 |
| 3,54858 | 2,7562 | 4,59016 | 3,54858 | 0,9197975 | 0,0207203 | F2Z3E1;P08 | SYP |
| -1,84156 | -3,09293 | -2,84933 | -2,84933 | 0,663434 | 0,0211064 | Q9Y295;H0 | DRG1 |
| 3,13383 | 2,43433 | 1,85588 | 2,43433 | 0,6399298 | 0,021571 | Q16568 | CARTPT |
| 2,05425 | 1,7563 | 2,8702 | 2,05425 | 0,5766746 | 0,0216303 | P13591;A0A | NCAM1 |
| 0,816608 | 1,33319 | 0,949245 | 0,949245 | 0,2682857 | 0,0217526 | F8VXY3;P0( | OAS1 |
| 0,926506 | 1,14161 | 1,54959 | 1,14161 | 0,3164783 | 0,0221969 | Q5T670;Q7 | MCM10 |
| 0,878509 | 1,37537 | 0,910903 | 0,910903 | 0,2779838 | 0,0223723 | M0R2J8;B6 | DCDC1 |
| 4,91539 | 2,86226 | 3,85337 | 3,85337 | 1,0267691 | 0,0225902 | E7EVR1;Q9 | ZNF302 |
| 0,411911 | 0,34051 | 0,573776 | 0,411911 | 0,1195209 | 0,0235106 | A0A0X1KG7 | COBLL1 |
| 0,639066 | 0,961374 | 1,13471 | 0,961374 | 0,2515256 | 0,0244437 | A0A087WV | TEX2 |
| -3,09825 | -1,7805 | -2,3159 | -2,3159 | 0,6627204 | 0,0245221 | Q495B1 | ANKDD1A |
| -2,06607 | -3,20249 | -3,66645 | -3,20249 | 0,8234 | 0,0245431 | F8W7R3;Q9 | FANCI |
| -2,54788 | -2,45254 | -1,47052 | -2,45254 | 0,5964 | 0,0245491 | A0A075B7G | ZNF595 |
| -1,70697 | -0,962461 | -1,36829 | -1,36829 | 0,3727589 | 0,0246277 | Q09428;A0 | ABCC8 |
| -1,88897 | -3,32002 | -3,09411 | -3,09411 | 0,7693397 | 0,0248019 | B1AHB1;P3 | MCM5 |
| -1,7552 | -3,13477 | -2,5686 | -2,5686 | 0,6934673 | 0,0249662 | Q04917 | YWHAH |
| -1,17422 | -1,3839 | -0,774082 | -1,17422 | 0,3098263 | 0,0249681 | P27361;E9F | MAPK3 |
| 1,24291 | 1,46379 | 2,10563 | 1,46379 | 0,4481503 | 0,0250438 | E7EU09;P1 | FGFR1 |
| -0,294557 | -0,528065 | -0,456846 | -0,456846 | 0,1196772 | 0,0252574 | P57737;A0A | CORO7 |
| 0,599342 | 0,61423 | 0,959026 | 0,61423 | 0,2035021 | 0,0253253 | C9J4R3;F8V | PASK |
| -0,175886 | -0,318537 | -0,260883 | -0,260883 | 0,0717609 | 0,0260275 | H7BXZ6;Q8 | RHOT1 |
| -0,711718 | -0,441357 | -0,449604 | -0,449604 | 0,1537676 | 0,0265222 | P45880;A0A | VDAC2 |
| -2,26628 | -1,35949 | -1,47881 | -1,47881 | 0,492716 | 0,0268309 | Q7L576;A0A | CYFIP1 |
| -0,658845 | -0,442702 | -0,386246 | -0,442702 | 0,1438838 | 0,0269299 | E7EV01;O1 | CAPN5 |
| -1,19048 | -0,690862 | -0,807685 | -0,807685 | 0,261342 | 0,0271868 | P20618 | PSMB1 |
| 1,51071 | 0,950088 | 0,928653 | 0,950088 | 0,3300371 | 0,0272854 | A0A2R8Y6C | CLPB |
| 0,833101 | 0,754484 | 1,27271 | 0,833101 | 0,2792833 | 0,0274304 | P0CG38 | POTEI |
| 0,656107 | 1,16958 | 1,17122 | 1,16958 | 0,2969283 | 0,0282093 | Q7L0J3 | SV2A |
| 1,25918 | 0,738809 | 0,82126 | 0,82126 | 0,2796897 | 0,0282796 | Q9Y383;A0 | LUC7L2 |
| -1,36711 | -0,812635 | -0,874497 | -0,874497 | 0,3038467 | 0,0284309 | Q9BWD1 | ACAT2 |
| 7,19241 | 6,31037 | 3,85755 | 6,31037 | 1,7279859 | 0,0284597 | A0A087WX | ABRAXAS1 |
| -0,393216 | -0,330449 | -0,209812 | -0,330449 | 0,0932112 | 0,0286336 | P49915 | GMPS |
| 1,7933 | 2,35895 | 3,26362 | 2,35895 | 0,7416455 | 0,0287183 | Q68DQ2 | CRYBG3 |
| 1,65732 | 0,886069 | 1,31726 | 1,31726 | 0,3865218 | 0,0287794 | O75145;R4 | PPFIA3 |
| -1,71863 | -2,40364 | -3,18504 | -2,40364 | 0,7337328 | 0,0289403 | O15054;I3L | KDM6B |
| 3,6449 | 1,92903 | 3,08396 | 3,08396 | 0,8749026 | 0,0292953 | A0A087WY | SEZ6L2 |
| -1,14063 | -0,954015 | -1,69639 | -1,14063 | 0,3861811 | 0,0297485 | Q15008;C9 | PSMD6 |
| 5,40367 | 4,40651 | 2,84325 | 4,40651 | 1,2905981 | 0,0298206 | O95985 | TOP3B |
| -2,33535 | -2,20758 | -1,25179 | -2,20758 | 0,5921658 | 0,0299291 | B7Z645;H0Y | SYNCRIP |
| -1,55011 | -1,8994 | -0,995061 | -1,55011 | 0,4560541 | 0,0301643 | Q00610;A0 | CLTC |
| 0,882577 | 1,33325 | 0,75064 | 0,882577 | 0,3054908 | 0,0303737 | Q86U42;B4 | PABPN1 |
| 1,06725 | 1,468 | 0,786365 | 1,06725 | 0,3425695 | 0,0304588 | A6NEC2 | NPEPPSL1 |
| -0,777222 | -1,02871 | -1,4441 | -1,02871 | 0,3367792 | 0,0307359 | Q2TAC2 | CCDC57 |
| -2,06585 | -3,58503 | -3,96931 | -3,58503 | 1,0065402 | 0,0313069 | Q70CQ4;A0 | USP31 |
| -1,96803 | -2,44685 | -3,60286 | -2,44685 | 0,8404658 | 0,0314199 | O60281;J3K | ZNF292 |
| -1,74901 | -3,39832 | -2,94205 | -2,94205 | 0,8516406 | 0,0316796 | O95831;E9I | AIFM1 |
| -0,865932 | -0,65998 | -0,449652 | -0,65998 | 0,2081438 | 0,0317254 | P60174;U3I | TPI1 |

| | | | | | | | |
|---|---|---|---|---|---|---|---|
| 1,83533 | 0,985402 | 1,27621 | 1,27621 | 0,4319649 | 0,0317694 | Q05519;Q5 | SRSF11 |
| -1,45923 | -0,821549 | -0,938242 | -0,938242 | 0,3395298 | 0,0317927 | P12277;H0Y | CKB |
| 0,814384 | 1,19569 | 1,57018 | 1,19569 | 0,3779031 | 0,0318364 | A0A087WT | C11orf54 |
| 3,27811 | 5,98764 | 4,00992 | 4,00992 | 1,4016939 | 0,0318543 | Q6NV74;C9 | KIAA1211L |
| -1,32298 | -0,734829 | -1,40747 | -1,32298 | 0,3664028 | 0,0319418 | P17174 | GOT1 |
| -3,07325 | -1,97907 | -3,87719 | -3,07325 | 0,9527512 | 0,0324971 | Q16445 | GABRA6 |
| 0,964398 | 1,83947 | 1,31565 | 1,31565 | 0,4403628 | 0,032613 | P17036;C9J | ZNF3 |
| -1,20149 | -1,0579 | -0,612073 | -1,0579 | 0,3073522 | 0,0326945 | Q96Q15;J3I | SMG1 |
| -2,04363 | -1,63241 | -1,03785 | -1,63241 | 0,5056674 | 0,0328312 | Q5JX54 | CPNE1 |
| 0,63146 | 0,727058 | 0,368445 | 0,63146 | 0,1857055 | 0,0329833 | O14936;A0. | CASK |
| -1,40321 | -0,866647 | -1,71929 | -1,40321 | 0,4310465 | 0,0332885 | P40925;B9A | MDH1 |
| 1,88934 | 2,29254 | 1,15356 | 1,88934 | 0,577526 | 0,0333989 | Q14839;A0. | CHD4 |
| 0,527329 | 0,409799 | 0,77266 | 0,527329 | 0,1851435 | 0,0334231 | Q9UDT6;H7 | CLIP2 |
| 1,29528 | 0,69185 | 1,33329 | 1,29528 | 0,3598652 | 0,0334786 | O75146 | HIP1R |
| -2,25724 | -1,78476 | -3,35408 | -2,25724 | 0,8050943 | 0,0337582 | O00399 | DCTN6 |
| -1,51942 | -1,34538 | -0,764314 | -1,34538 | 0,3954139 | 0,0338179 | P53007 | SLC25A1 |
| -0,867116 | -1,56091 | -0,98132 | -0,98132 | 0,372003 | 0,0339103 | Q9NRD9;H( | DUOX1 |
| -0,858806 | -0,897966 | -1,49139 | -0,897966 | 0,3544592 | 0,0339181 | P22314;Q5. | UBA1 |
| -2,00503 | -1,03022 | -1,44634 | -1,44634 | 0,4891395 | 0,0339285 | P10909;H0Y | CLU |
| -1,90543 | -3,15228 | -3,81324 | -3,15228 | 0,9687829 | 0,0339671 | P05198 | EIF2S1 |
| -0,751213 | -1,09643 | -0,574583 | -0,751213 | 0,2654233 | 0,0341858 | Q5SSJ5;B0( | HP1BP3 |
| 0,962122 | 0,611289 | 1,225 | 0,962122 | 0,3079042 | 0,0344523 | A0A2R8Y7K | LOXHD1 |
| 3,80726 | 2,24674 | 2,21137 | 2,24674 | 0,9113487 | 0,0345913 | H3BN72;H3 | COX4I1 |
| -0,959419 | -0,883937 | -0,480876 | -0,883937 | 0,2572804 | 0,0348497 | O00505 | KPNA3 |
| -1,10064 | -0,586348 | -0,721362 | -0,721362 | 0,2666386 | 0,0348614 | P32119;A6I | PRDX2 |
| 2,48561 | 4,58193 | 2,90006 | 2,90006 | 1,1101811 | 0,0352593 | Q86W11;A( | ZSCAN30 |
| 1,57968 | 0,774932 | 1,27548 | 1,27548 | 0,4063466 | 0,0355958 | P50851;H0Y | LRBA |
| 2,42763 | 1,36526 | 1,4545 | 1,4545 | 0,5892899 | 0,0358145 | Q9NUP9;G3 | LIN7C |
| -1,93066 | -1,34043 | -2,68461 | -1,93066 | 0,6737497 | 0,0363147 | A0A0D9SFF | HADH |
| 0,805558 | 1,66227 | 1,42552 | 1,42552 | 0,4424099 | 0,0366222 | O60303 | KIAA0556 |
| -0,227083 | -0,386674 | -0,469171 | -0,386674 | 0,1230729 | 0,0366321 | P34897;H0Y | SHMT2 |
| -1,85444 | -2,42691 | -3,61539 | -2,42691 | 0,8982531 | 0,0366938 | Q9UEG4 | ZNF629 |
| -0,866024 | -1,79319 | -1,4915 | -1,4915 | 0,4729119 | 0,0368069 | Q96ER9;C9. | CCDC51 |
| -1,02493 | -0,593151 | -1,23208 | -1,02493 | 0,3259792 | 0,0370743 | P07195;A8I | LDHB |
| -0,433245 | -0,863546 | -0,598098 | -0,598098 | 0,2171014 | 0,0371969 | Q5JZY3;J3K | EPHA10 |
| 1,18366 | 2,46465 | 1,96618 | 1,96618 | 0,6457225 | 0,0374661 | O00743 | PPP6C |
| -1,15493 | -2,28276 | -1,51454 | -1,51454 | 0,5761195 | 0,0382852 | O15417;H9 | TNRC18 |
| -2,45522 | -1,35593 | -1,43894 | -1,43894 | 0,6121212 | 0,0384453 | Q14697;F5I | GANAB |
| -1,26147 | -0,980513 | -1,92766 | -1,26147 | 0,4864554 | 0,0384905 | Q02878;F8Y | RPL6 |
| -0,305657 | -0,569222 | -0,642074 | -0,569222 | 0,1769889 | 0,0384957 | O60502;H7 | OGA |
| -2,41101 | -1,49757 | -3,14615 | -2,41101 | 0,8258954 | 0,0387423 | F5H1Y4;Q9 | GOPC |
| 0,401224 | 0,342175 | 0,652887 | 0,401224 | 0,1650066 | 0,0394347 | P12004 | PCNA |
| -2,6158 | -1,227 | -2,33631 | -2,33631 | 0,7345576 | 0,0398767 | Q9H6S3 | EPS8L2 |
| -1,22575 | -1,85978 | -0,924433 | -1,22575 | 0,4774341 | 0,0399931 | Q9NP80 | PNPLA8 |
| -0,772845 | -0,751539 | -0,371313 | -0,751539 | 0,2259254 | 0,0400667 | Q9ULL1 | PLEKHG1 |
| 1,48833 | 2,10447 | 1,00893 | 1,48833 | 0,5491904 | 0,0401716 | A0A0A0MR | FADS1 |
| -0,385719 | -0,798827 | -0,554699 | -0,554699 | 0,20769 | 0,040216 | O14828 | SCAMP3 |
| 1,19577 | 0,897822 | 0,56054 | 0,897822 | 0,3178179 | 0,0404262 | Q08AF3;B4 | SLFN5 |
| -0,832196 | -1,47534 | -1,81422 | -1,47534 | 0,4988061 | 0,0412371 | Q5THR3 | EFCAB6 |
| 0,76881 | 0,414972 | 0,905611 | 0,76881 | 0,2531938 | 0,0413415 | A0A0G2JIR: | EHMT2 |
| 1,44678 | 3,1196 | 2,86368 | 2,86368 | 0,9010575 | 0,0414 | A0A0B4J1Y. | TTC21A |
| -0,503985 | -1,02047 | -1,07599 | -1,02047 | 0,3154439 | 0,0414203 | O95747;C9. | OXSR1 |

| | | | | | | | |
|---|---|---|---|---|---|---|---|
| 3,10994 | 3,91716 | 1,79474 | 3,10994 | 1,0712938 | 0,0415056 | Q08AE8;J3I | SPIRE1 |
| 3,28709 | 1,5012 | 2,55453 | 2,55453 | 0,8977334 | 0,0420352 | Q63HM2;B! | PCNX4 |
| 10,4593 | 10,3341 | 4,94249 | 10,3341 | 3,1496118 | 0,0421141 | Q6P179 | ERAP2 |
| -0,95554 | -1,90574 | -2,0845 | -1,90574 | 0,6068204 | 0,0423158 | O15327;E7I | INPP4B |
| 2,01747 | 2,30912 | 3,89987 | 2,30912 | 1,0131613 | 0,0426165 | Q14584 | ZNF266 |
| -0,631208 | -1,13485 | -0,616753 | -0,631208 | 0,2950392 | 0,0430465 | Q96PY6;H0 | NEK1 |
| -1,05788 | -0,601431 | -0,556023 | -0,601431 | 0,2775692 | 0,0440106 | Q9NXD2 | MTMR10 |
| -2,12357 | -1,67271 | -0,945612 | -1,67271 | 0,5943528 | 0,0440407 | A0A0A0MS | ABCA8 |
| 1,43204 | 2,72096 | 3,23392 | 2,72096 | 0,928369 | 0,0442624 | P11488 | GNAT1 |
| -1,89275 | -1,24673 | -2,71736 | -1,89275 | 0,7371201 | 0,0443798 | Q9UMD9;H | COL17A1 |
| 2,92781 | 2,57822 | 1,29967 | 2,57822 | 0,857102 | 0,0444346 | P12532;F8V | CKMT1A |
| 2,74883 | 1,34614 | 1,69833 | 1,69833 | 0,7297405 | 0,0444502 | A7E2Y1;A0/ | MYH7B |
| 2,93792 | 3,70704 | 1,6395 | 2,93792 | 1,0450009 | 0,0445673 | Q7Z3T8 | ZFYVE16 |
| 0,992791 | 1,11523 | 0,495165 | 0,992791 | 0,3284061 | 0,0445773 | Q9UBU7;B4 | DBF4 |
| 2,36023 | 4,34769 | 5,35289 | 4,34769 | 1,5229597 | 0,0446553 | Q96EA4;D6 | SPDL1 |
| 0,556596 | 0,803295 | 1,19961 | 0,803295 | 0,3243951 | 0,044965 | Q9Y263;E5I | PLAA |
| -0,485109 | -0,896678 | -1,11098 | -0,896678 | 0,3180746 | 0,0455345 | A0A0A0MR | TNIP1 |
| -0,749119 | -1,38212 | -0,739244 | -0,749119 | 0,3683471 | 0,0460165 | Q01082;A0. | SPTBN1 |
| 1,73852 | 1,03942 | 2,36677 | 1,73852 | 0,6639901 | 0,0465128 | Q2M2I5 | KRT24 |
| 4,1504 | 1,87324 | 2,7827 | 2,7827 | 1,1462387 | 0,0472521 | H0Y9Y7 | TACC2 |
| 1,39354 | 0,595409 | 1,1083 | 1,1083 | 0,4044404 | 0,0475358 | Q13151 | HNRNPA0 |
| 2,18497 | 1,70091 | 0,932347 | 1,70091 | 0,6316734 | 0,0478888 | A8MQ14 | ZNF850 |
| -0,990301 | -0,531795 | -1,25096 | -0,990301 | 0,36409 | 0,0480209 | Q969G3;B4 | SMARCE1 |
| 0,64713 | 0,560197 | 0,274207 | 0,560197 | 0,195115 | 0,0482947 | P55809;E9F | OXCT1 |
| -0,998642 | -0,4255 | -0,906597 | -0,906597 | 0,3077928 | 0,0485402 | P50990;H7( | CCT8 |
| 0,992957 | 0,426276 | 0,934836 | 0,934836 | 0,3117528 | 0,0487953 | Q15637;C9. | SF1 |
| -1,35966 | -0,848545 | -0,627008 | -0,848545 | 0,3757429 | 0,0488605 | Q9P0M6;Q! | H2AFY2 |
| 0,193047 | 0,4386 | 0,441473 | 0,4386 | 0,1426067 | 0,0491092 | P62263;A0/ | RPS14 |
| -1,19283 | -1,09125 | -0,502964 | -1,09125 | 0,37245 | 0,0496221 | Q9Y4A5;F2. | TRRAP |
| 3,30041 | 1,41211 | 2,32426 | 2,32426 | 0,9443307 | 0,0500103 | Q9P2R7;A0 | SUCLA2 |
| 0,872253 | 1,78014 | 2,10409 | 1,78014 | 0,6385693 | 0,0500467 | Q14315 | FLNC |
| -0,725486 | -1,53189 | -1,74825 | -1,53189 | 0,539003 | 0,0502604 | Q96EK9 | KTI12 |
| -1,79813 | -2,32629 | -3,8843 | -2,32629 | 1,0846239 | 0,0508625 | Q6P0N0;G5 | MIS18BP1 |
| -3,09273 | -1,43828 | -1,82464 | -1,82464 | 0,865499 | 0,0513827 | O15397;F5( | IPO8 |
| -1,06833 | -2,37901 | -1,46151 | -1,46151 | 0,6725919 | 0,0519686 | K7EJV4 | MYCBPAP |
| 0,352439 | 0,466364 | 0,775054 | 0,466364 | 0,2186595 | 0,0520899 | Q96JN2;C9I | CCDC136 |
| 1,42121 | 0,572773 | 1,13438 | 1,13438 | 0,4315706 | 0,0526275 | A0A0A6YYH | C15orf38-AP3S |
| -1,16132 | -1,27472 | -2,3607 | -1,27472 | 0,6621587 | 0,0526908 | O95347;Q5 | SMC2 |
| -0,575801 | -1,27994 | -1,42043 | -1,27994 | 0,4525755 | 0,0527595 | Q8N111 | CEND1 |
| 0,709965 | 0,692918 | 0,295314 | 0,692918 | 0,2346327 | 0,0527767 | Q92896;H3 | GLG1 |
| 0,730554 | 1,05018 | 1,66889 | 1,05018 | 0,477046 | 0,0528633 | O95487 | SEC24B |
| 0,750992 | 0,42654 | 1,04321 | 0,750992 | 0,3084754 | 0,0532993 | H0Y8W2;H( | RACK1 |
| 3,44864 | 3,49266 | 6,78601 | 3,49266 | 1,9142505 | 0,0536829 | G3XAE0;Q5 | ZNF766 |
| 3,55159 | 1,41578 | 2,73617 | 2,73617 | 1,0778083 | 0,0540112 | O95235 | KIF20A |
| 2,55048 | 2,26184 | 1,0144 | 2,26184 | 0,8163906 | 0,0541538 | Q9Y2I8;A0/ | WDR37 |
| -1,49639 | -3,50686 | -2,2328 | -2,2328 | 1,0171462 | 0,0544785 | Q9UL15;G3 | BAG5 |
| -0,345016 | -0,665463 | -0,332109 | -0,345016 | 0,1888464 | 0,0545436 | P07237;H7I | P4HB |
| -1,22718 | -3,11913 | -2,73142 | -2,73142 | 0,9993775 | 0,0549302 | O96019;H7 | ACTL6A |
| 0,36268 | 0,922954 | 0,71057 | 0,71057 | 0,2828549 | 0,0552858 | Q92785;J3K | DPF2 |
| 2,85041 | 1,49626 | 3,79974 | 2,85041 | 1,1576535 | 0,05558 | Q8NDZ4;C9 | C3orf58 |
| -0,880863 | -0,401437 | -1,03131 | -0,880863 | 0,3289437 | 0,0556314 | A0A1B0GTI | FTO |
| 1,1998 | 2,45826 | 3,08362 | 2,45826 | 0,9594768 | 0,0557337 | H3BLV9;Q9 | SRPK1 |

| | | | | | | | |
|---|---|---|---|---|---|---|---|
| 1,74354 | 1,35712 | 3,02476 | 1,74354 | 0,8729135 | 0,0558681 | F2Z2U2;Q9 | ERGIC1 |
| 1,58493 | 2,72485 | 3,98973 | 2,72485 | 1,202941 | 0,05763 | Q7L1W4;E9 | LRRC8D |
| -1,28401 | -3,29267 | -2,34285 | -2,34285 | 1,004823 | 0,0578294 | F6WFR7;Q9 | NTM |
| 3,29197 | 1,75837 | 1,53302 | 1,75837 | 0,9571326 | 0,0579541 | Q14161;F8 | GIT2 |
| 1,01842 | 0,510183 | 0,502648 | 0,510183 | 0,2956299 | 0,0580673 | P60763;J3K | RAC3 |
| -1,91925 | -1,34242 | -0,749097 | -1,34242 | 0,5850959 | 0,0583154 | P52209;K7E | PGD |
| 0,682271 | 1,1773 | 1,73584 | 1,1773 | 0,5271034 | 0,0588447 | Q8TE82;D6 | SH3TC1 |
| 4,18628 | 2,23152 | 1,92915 | 2,23152 | 1,2252313 | 0,0589794 | J3KNK3;Q1 | NKPD1 |
| 5,51116 | 2,89156 | 2,56797 | 2,89156 | 1,6139693 | 0,0592207 | Q13111;K7 | CHAF1A |
| 0,293452 | 0,774889 | 0,561263 | 0,561263 | 0,2412262 | 0,0598917 | P23368;A0 | ME2 |
| -0,393244 | -0,269397 | -0,650749 | -0,393244 | 0,1945406 | 0,0599605 | P19367;B1 | HK1 |
| -1,13887 | -2,04464 | -0,895756 | -1,13887 | 0,6054555 | 0,0601833 | Q9BZV1;K7 | UBXN6 |
| 0,857887 | 0,319028 | 0,647036 | 0,647036 | 0,2715439 | 0,0605196 | Q3LXA3;H0 | TKFC |
| 1,7665 | 1,40751 | 0,645343 | 1,40751 | 0,5725332 | 0,0612815 | Q9HCJ0 | TNRC6C |
| -1,81189 | -0,677901 | -1,29567 | -1,29567 | 0,5677518 | 0,0613404 | J3KT51;Q9 | JPT1 |
| -1,57308 | -3,29017 | -1,62779 | -1,62779 | 0,9759524 | 0,061617 | H3BT97 | MMP15 |
| 1,2838 | 0,671844 | 1,80135 | 1,2838 | 0,5654102 | 0,0617226 | P98171;E7E | ARHGAP4 |
| 1,27219 | 0,993457 | 2,29838 | 1,27219 | 0,6872144 | 0,0617796 | Q15858;A0 | SCN9A |
| 1,1767 | 2,99476 | 1,90112 | 1,90112 | 0,9152572 | 0,0618903 | P40818 | USP8 |
| -2,01944 | -0,738557 | -1,52483 | -1,52483 | 0,6459522 | 0,0619679 | Q9NSD9 | FARSB |
| 1,57208 | 2,93275 | 1,29427 | 1,57208 | 0,8768521 | 0,0622525 | P42357 | HAL |
| -2,49651 | -1,02291 | -1,46787 | -1,46787 | 0,7558204 | 0,0625108 | J3KNX4;Q2 | MUM1 |
| 0,62825 | 1,46188 | 1,753 | 1,46188 | 0,583774 | 0,0627737 | H7BZK9;Q9 | C7orf26 |
| -0,519487 | -1,4501 | -1,19106 | -1,19106 | 0,4803042 | 0,0628211 | Q9H0R4;K7 | HDHD2 |
| -0,922589 | -2,36757 | -2,53375 | -2,36757 | 0,8861363 | 0,0629647 | B5BUE6;J3K | DDX5 |
| 1,97237 | 0,72931 | 1,37006 | 1,37006 | 0,6216291 | 0,0633516 | Q9H9E3;J3I | COG4 |
| 1,66757 | 0,822778 | 2,27309 | 1,66757 | 0,7284382 | 0,0635424 | A0A087X16 | PHF19 |
| 0,575807 | 1,6189 | 1,27798 | 1,27798 | 0,5318703 | 0,0637196 | Q8N283;F6 | ANKRD35 |
| 0,756464 | 2,01068 | 1,32685 | 1,32685 | 0,6279625 | 0,0638921 | A0A0D9SEV | PNKP |
| -1,03829 | -1,38348 | -2,48144 | -1,38348 | 0,7535863 | 0,0641233 | P17612;Q1 | PRKACA |
| -0,694313 | -1,64079 | -1,98034 | -1,64079 | 0,6664556 | 0,0646854 | Q9NYD6 | HOXC10 |
| 2,46128 | 3,5828 | 1,30655 | 2,46128 | 1,1381654 | 0,0649924 | E9PQR7;Q9 | VPS28 |
| -1,13005 | -1,36601 | -2,61107 | -1,36601 | 0,7957462 | 0,0657322 | H3BS70;P4 | ECI1 |
| 1,1065 | 1,24292 | 2,47183 | 1,24292 | 0,7519925 | 0,0658564 | B7WPL9;F8 | PPIP5K1 |
| -1,55227 | -2,40469 | -0,893145 | -1,55227 | 0,7578296 | 0,0660667 | Q8WUF5;K | PPP1R13L |
| 0,659342 | 1,60917 | 1,91239 | 1,60917 | 0,6537386 | 0,0661527 | P12036 | NEFH |
| -0,819383 | -0,660424 | -0,281841 | -0,660424 | 0,2761474 | 0,0664527 | P10809;E7E | HSPD1 |
| -2,31114 | -1,12291 | -3,19578 | -2,31114 | 1,0401337 | 0,0665535 | Q9BV86;S4 | NTMT1 |
| 1,34132 | 3,55249 | 2,2062 | 2,2062 | 1,114285 | 0,0665949 | Q5JRD6 | PDSS2 |
| 3,38008 | 1,18105 | 3,22764 | 3,22764 | 1,2279727 | 0,0671458 | O95104 | SCAF4 |
| 1,47429 | 0,729159 | 0,659434 | 0,729159 | 0,4516769 | 0,0672311 | O43314;A0 | PPIP5K2 |
| -0,463114 | -0,229566 | -0,206705 | -0,229566 | 0,1418995 | 0,0672339 | Q9BXM9;A | FSD1L |
| -1,94188 | -0,661527 | -1,56564 | -1,56564 | 0,6580629 | 0,0672885 | Q15293 | RCN1 |
| -1,25201 | -3,50227 | -3,537 | -3,50227 | 1,3093291 | 0,0673435 | Q9HBT8;A8 | ZNF286A |
| 1,23876 | 2,34771 | 0,989138 | 1,23876 | 0,7231642 | 0,0674446 | Q8IXT1;E9P | DDIAS |
| 2,11973 | 0,870147 | 1,14218 | 1,14218 | 0,6571483 | 0,0682063 | Q96JE9 | MAP6 |
| 1,40906 | 1,49557 | 0,51359 | 1,40906 | 0,5436965 | 0,0682231 | P07355;H0 | ANXA2 |
| 2,68907 | 1,44716 | 1,10421 | 1,44716 | 0,8338401 | 0,068268 | O60299 | LZTS3 |
| 1,85958 | 2,04294 | 0,693799 | 1,85958 | 0,7317613 | 0,068337 | Q86WA8;H | LONP2 |
| 3,18058 | 4,53212 | 1,57642 | 3,18058 | 1,4796482 | 0,0684004 | P41970 | ELK3 |
| 0,790653 | 2,28497 | 1,61307 | 1,61307 | 0,7484209 | 0,0686591 | P34932;A0 | HSPA4 |
| 1,14314 | 0,441188 | 1,31474 | 1,14314 | 0,4628312 | 0,0686789 | Q96T51;J3K | RUFY1 |

| | | | | | | | |
|---|---|---|---|---|---|---|---|
| 0,926093 | 0,461752 | 1,32967 | 0,926093 | 0,4343134 | 0,0688116 | E9PCX8;C9J | TNS3 |
| 0,573661 | 0,397119 | 1,01556 | 0,573661 | 0,3185674 | 0,0692447 | O43809;H3 | NUDT21 |
| 0,440851 | 0,346271 | 0,838025 | 0,440851 | 0,2609325 | 0,0693852 | Q13045;J3K | FLII |
| 2,58288 | 2,75232 | 0,930256 | 2,58288 | 1,0066275 | 0,0694657 | Q7Z7G8 | VPS13B |
| 2,33694 | 0,890945 | 1,36208 | 1,36208 | 0,7374756 | 0,0694723 | O00423;F8Y | EML1 |
| -1,15558 | -0,779524 | -2,01866 | -1,15558 | 0,6353192 | 0,0694844 | O00232 | PSMD12 |
| 1,78416 | 0,978508 | 2,75106 | 1,78416 | 0,8874975 | 0,0696975 | Q05513;E9I | PRKCZ |
| 0,410113 | 1,23005 | 1,11393 | 1,11393 | 0,4436851 | 0,0698054 | Q8IU85 | CAMK1D |
| -2,40428 | -0,805662 | -1,77243 | -1,77243 | 0,805135 | 0,0701923 | P55209;F5H | NAP1L1 |
| 3,41874 | 1,93827 | 1,31772 | 1,93827 | 1,0794411 | 0,0702889 | Q9NSC5;M( | HOMER3 |
| 0,364785 | 0,242363 | 0,12663 | 0,242363 | 0,1190932 | 0,0707423 | Q03252 | LMNB2 |
| 0,825882 | 2,23308 | 2,51436 | 2,23308 | 0,9046438 | 0,0707542 | A0A1B0GV( | RAP1GAP2 |
| -0,905384 | -1,58757 | -0,602691 | -0,905384 | 0,5044779 | 0,0712607 | P48047;H7( | ATP5PO |
| -2,60247 | -0,9729 | -2,99568 | -2,60247 | 1,0725166 | 0,0714598 | A0A1W2PP | STRADA |
| -1,14274 | -0,610799 | -0,453611 | -0,610799 | 0,3611482 | 0,0717795 | Q8N653;H7 | LZTR1 |
| -1,19279 | -1,27281 | -0,417756 | -1,19279 | 0,4722637 | 0,0719082 | P49411;H3I | TUFM |
| 1,18044 | 0,622725 | 0,472052 | 0,622725 | 0,373176 | 0,0720875 | Q8IZQ1;D6I | WDFY3 |
| 0,855321 | 2,48378 | 2,59671 | 2,48378 | 0,9744287 | 0,0722001 | Q5T3Q7;Q9 | HEATR1 |
| -2,48198 | -1,21579 | -1,0542 | -1,21579 | 0,7818679 | 0,0724943 | Q9Y5G2 | PCDHGB2 |
| -1,75305 | -1,40499 | -0,561788 | -1,40499 | 0,6125412 | 0,0726001 | P46782;M0 | RPS5 |
| -3,04989 | -1,17494 | -1,67028 | -1,67028 | 0,9716072 | 0,0727154 | P11137;A8I | MAP2 |
| 1,58071 | 2,3417 | 0,786585 | 1,58071 | 0,7776163 | 0,0729672 | A0A0G2JN( | HNRNPCL2 |
| 0,699995 | 1,21359 | 0,447786 | 0,699995 | 0,3902659 | 0,0730744 | Q6ZVF9 | GPRIN3 |
| -2,38028 | -0,886494 | -1,34569 | -1,34569 | 0,7651399 | 0,0735608 | Q86VP6;A0 | CAND1 |
| 0,634142 | 2,00961 | 1,63008 | 1,63008 | 0,7103811 | 0,0738228 | Q12929;A0 | EPS8 |
| -0,674521 | -1,44849 | -0,632497 | -0,674521 | 0,4594632 | 0,0742413 | Q13439;H0 | GOLGA4 |
| -1,61522 | -1,10784 | -0,529916 | -1,10784 | 0,543034 | 0,0743923 | Q13838;F6Y | DDX39B |
| 0,498637 | 0,729983 | 1,32519 | 0,729983 | 0,4264157 | 0,0744227 | P14866;M0 | HNRNPL |
| -0,94896 | -2,49648 | -3,03956 | -2,49648 | 1,0847705 | 0,0746622 | A0A087WY | ZMYND8 |
| -0,635319 | -2,01207 | -1,85342 | -1,85342 | 0,7532577 | 0,0747309 | P21333;Q6( | FLNA |
| 8,55022 | 5,01958 | 3,05147 | 5,01958 | 2,7861302 | 0,0749412 | Q12770;C9 | SCAP |
| 0,4439 | 0,638392 | 0,206645 | 0,4439 | 0,2162262 | 0,0750456 | O15550;A0 | KDM6A |
| 1,00123 | 2,66478 | 3,23643 | 2,66478 | 1,1611992 | 0,0754238 | Q9ULD4;E9 | BRPF3 |
| -0,269281 | -0,851398 | -0,812239 | -0,812239 | 0,3253708 | 0,0755046 | Q15154;H0 | PCM1 |
| -0,661605 | -0,250075 | -0,811855 | -0,661605 | 0,2908404 | 0,0758363 | Q15365 | PCBP1 |
| 0,597556 | 0,645146 | 1,39218 | 0,645146 | 0,445674 | 0,0761545 | Q5JY65;Q9I | CRNKL1 |
| 0,808489 | 2,26972 | 1,30894 | 1,30894 | 0,7426019 | 0,0762532 | P08603 | CFH |
| -1,42321 | -0,439097 | -1,07591 | -1,07591 | 0,4991036 | 0,0767326 | P05023;Q5 | ATP1A1 |
| -2,78523 | -3,29484 | -1,00184 | -2,78523 | 1,2040232 | 0,0768514 | P04114;A8I | APOB |
| 0,558792 | 0,838743 | 1,52718 | 0,838743 | 0,4983462 | 0,0771538 | A0A087WV | CACNB2 |
| -2,71139 | -1,46597 | -1,00277 | -1,46597 | 0,8836485 | 0,0773096 | Q9NTK5;J3I | OLA1 |
| 1,05912 | 2,55481 | 1,20156 | 1,20156 | 0,8254961 | 0,0779865 | O75376;A0 | NCOR1 |
| -1,14335 | -0,343512 | -0,999479 | -0,999479 | 0,4263669 | 0,0780335 | Q06830;A0 | PRDX1 |
| -2,44812 | -0,93268 | -1,26107 | -1,26107 | 0,7972334 | 0,0782471 | P35609;F6T | ACTN2 |
| 1,3592 | 2,80692 | 1,12515 | 1,3592 | 0,910954 | 0,0785808 | Q96KP4;J3( | CNDP2 |
| -1,84584 | -0,617937 | -1,11015 | -1,11015 | 0,6179616 | 0,0791847 | Q8TDY2 | RB1CC1 |
| -2,41637 | -3,00559 | -0,890316 | -2,41637 | 1,0916658 | 0,0792133 | E9PDF1;Q8 | SH3TC2 |
| 3,46948 | 1,81 | 1,29209 | 1,81 | 1,1374776 | 0,0793324 | Q8NBU5 | ATAD1 |
| -0,745399 | -1,9955 | -1,03576 | -1,03576 | 0,6542378 | 0,0794474 | P12814;H9I | ACTN1 |
| 2,88165 | 1,73367 | 0,96111 | 1,73367 | 0,9663661 | 0,0794981 | Q5T7W7;Q | TSTD2 |
| 0,842746 | 2,53844 | 2,85726 | 2,53844 | 1,0828428 | 0,079726 | O43854 | EDIL3 |
| 2,61725 | 2,60167 | 0,798419 | 2,60167 | 1,045634 | 0,0798839 | E7ER45;O4 | MGAM |

| | | | | | | | |
|---|---|---|---|---|---|---|---|
| 1,10939 | 2,18314 | 0,826252 | 1,10939 | 0,7158039 | 0,0799002 | Q5T4S7;X6I | UBR4 |
| 0,744727 | 0,417377 | 1,25052 | 0,744727 | 0,4197443 | 0,0800537 | Q9H992 | MARCH7 |
| -1,70391 | -0,656229 | -0,841618 | -0,841618 | 0,5590993 | 0,0805773 | Q9HB07;F8 | C12orf10 |
| -2,21796 | -3,19649 | -0,970406 | -2,21796 | 1,115748 | 0,0806803 | P13639 | EEF2 |
| 3,13637 | 3,74528 | 1,08189 | 3,13637 | 1,3955468 | 0,0810824 | Q9ULE3;F8\ | DENND2A |
| 1,54997 | 1,04797 | 0,473042 | 1,04797 | 0,5388754 | 0,0812705 | J9JIE5;Q14 | NFE2L1 |
| 0,771399 | 1,42237 | 2,36356 | 1,42237 | 0,8004768 | 0,0814117 | A6NK75;Q9 | ZNF98 |
| 0,398599 | 0,645151 | 0,206495 | 0,398599 | 0,2198905 | 0,0816013 | P35612;C9J | ADD2 |
| -0,74988 | -0,745592 | -1,73207 | -0,74988 | 0,5683095 | 0,0817677 | Q9NYU2;H7 | UGGT1 |
| -0,574388 | -2,01486 | -1,63643 | -1,63643 | 0,7467822 | 0,0822947 | P05141 | SLC25A5 |
| -0,129414 | -0,432021 | -0,440782 | -0,432021 | 0,1772934 | 0,0824394 | A0A0G2JRV | SMARCB1 |
| 1,99122 | 4,724 | 2,07107 | 2,07107 | 1,5552331 | 0,0825255 | P62854 | RPS26 |
| -2,73355 | -0,79185 | -2,60202 | -2,60202 | 1,0850664 | 0,0825888 | P49327;A0/ | FASN |
| 1,39228 | 0,99038 | 2,71883 | 1,39228 | 0,904507 | 0,0827685 | Q5JRA6;A0 | MIA3 |
| 1,12452 | 2,39111 | 0,937836 | 1,12452 | 0,790686 | 0,0829671 | Q96DA2 | RAB39B |
| -0,997405 | -3,03178 | -3,53291 | -3,03178 | 1,3427955 | 0,0829875 | A0A2R8Y5N | TSC1 |
| -0,886627 | -0,257443 | -0,635893 | -0,635893 | 0,316745 | 0,0833007 | Q9BZ29;A0 | DOCK9 |
| -2,03651 | -0,910183 | -0,834345 | -0,910183 | 0,6732464 | 0,0833901 | Q9UMX0;H | UBQLN1 |
| -0,0767237 | -0,251912 | -0,269149 | -0,251912 | 0,1064703 | 0,0834305 | Q86TI0;H0Y | TBC1D1 |
| -2,21888 | -0,842122 | -1,07452 | -1,07452 | 0,7370019 | 0,0835163 | P43304;F5C | GPD2 |
| -0,876592 | -1,1751 | -2,36756 | -1,1751 | 0,7888867 | 0,0837629 | Q13574;E9I | DGKZ |
| -1,3786 | -0,40046 | -0,974254 | -0,974254 | 0,4915101 | 0,0837665 | A0A0J9YX8 | GOLGA8Q |
| -1,95903 | -1,2988 | -3,70431 | -1,95903 | 1,242872 | 0,0837691 | Q8WX93;H | PALLD |
| -0,515338 | -1,84327 | -1,62497 | -1,62497 | 0,7120793 | 0,0839623 | Q6ZS30 | NBEAL1 |
| 2,0246 | 0,806362 | 0,923034 | 0,923034 | 0,6722058 | 0,0842183 | O76041;Q5 | NEBL |
| -1,7158 | -1,12192 | -0,513104 | -1,12192 | 0,6013635 | 0,0845512 | P20700;E9F | LMNB1 |
| 1,00135 | 0,362603 | 0,503676 | 0,503676 | 0,335554 | 0,0847172 | P29083;C9I | GTF2E1 |
| -2,09046 | -0,693359 | -1,16883 | -1,16883 | 0,7103245 | 0,0847506 | P55036;Q5 | PSMD4 |
| 2,57187 | 0,763601 | 1,6827 | 1,6827 | 0,9041758 | 0,0851408 | Q9UBP0;AC | SPAST |
| 0,788776 | 2,67995 | 1,76767 | 1,76767 | 0,9457825 | 0,0855029 | Q9Y653;H3 | ADGRG1 |
| 3,5739 | 1,02426 | 3,61607 | 3,5739 | 1,4843585 | 0,0855763 | Q9BXU1;C9 | STK31 |
| -3,25891 | -2,78154 | -0,890063 | -2,78154 | 1,2527971 | 0,0856255 | Q8WYQ5 | DGCR8 |
| -0,469648 | -0,555368 | -0,151385 | -0,469648 | 0,2128542 | 0,0857678 | Q8WXI2;A0 | CNKSR2 |
| 0,248005 | 0,633432 | 0,288023 | 0,288023 | 0,2119209 | 0,0859968 | E7EUN2;A0 | AGAP1 |
| 0,668563 | 0,391964 | 0,210643 | 0,391964 | 0,2306061 | 0,0861624 | Q99829;B0 | CPNE1 |
| 1,35824 | 1,11529 | 2,9217 | 1,35824 | 0,9803527 | 0,0864077 | Q06124;A0 | PTPN11 |
| -0,933933 | -2,64142 | -3,42878 | -2,64142 | 1,2753894 | 0,0867268 | Q96RL7;H0 | VPS13A |
| 0,694877 | 0,554163 | 0,187498 | 0,554163 | 0,2619405 | 0,0869354 | B1AK87;B1 | CAPZB |
| 0,681043 | 2,2105 | 2,53206 | 2,2105 | 0,989015 | 0,0869454 | Q9NR48;F8 | ASH1L |
| -1,38446 | -1,65479 | -0,443444 | -1,38446 | 0,6358654 | 0,0871327 | P49821;G3 | NDUFV1 |
| -0,776171 | -2,90626 | -2,377 | -2,377 | 1,1090573 | 0,087509 | Q8N5M1;C | ATPAF2 |
| -2,33341 | -1,00683 | -0,940649 | -1,00683 | 0,7857033 | 0,0879331 | P07205 | PGK2 |
| 1,64801 | 0,552154 | 2,09035 | 1,64801 | 0,7918977 | 0,0887965 | O94888;C9 | UBXN7 |
| -2,39754 | -0,629931 | -2,14655 | -2,14655 | 0,9563447 | 0,0890197 | Q14240;E7I | EIF4A2 |
| 4,54901 | 1,7877 | 6,52238 | 4,54901 | 2,3782422 | 0,0891125 | F5GX75;P0 | VTN |
| -1,47965 | -1,37066 | -0,390621 | -1,37066 | 0,5997693 | 0,0892083 | Q9NRX4 | PHPT1 |
| -0,38735 | -1,4077 | -1,44228 | -1,4077 | 0,5993312 | 0,0892674 | Q9UNM6;A | PSMD13 |
| -0,637108 | -0,610798 | -1,50706 | -0,637108 | 0,5100317 | 0,0892675 | Q15424;K7I | SAFB |
| -0,143743 | -0,0375237 | -0,125481 | -0,125481 | 0,0567928 | 0,0892787 | Q9BRX8 | PRXL2A |
| 0,658739 | 1,04689 | 1,98905 | 1,04689 | 0,6841118 | 0,0892924 | Q9H3H9 | TCEAL2 |
| -2,59144 | -0,675913 | -2,14806 | -2,14806 | 1,0027501 | 0,0892964 | P60842;J3K | EIF4A1 |
| -1,34405 | -0,350462 | -1,11629 | -1,11629 | 0,52051 | 0,0893098 | F5GWN5;O | PIK3C2B |

| | | | | | | | |
|---|---|---|---|---|---|---|---|
| 1,37065 | 3,48815 | 1,51798 | 1,51798 | 1,1823058 | 0,0894999 | A0A096LNH | PCDHB6 |
| 0,287258 | 1,10371 | 0,984422 | 0,984422 | 0,4409953 | 0,0897041 | P49840;A8I | GSK3A |
| 1,17158 | 2,43563 | 0,868518 | 1,17158 | 0,8312149 | 0,0897587 | O75367;B4 | H2AFY |
| 0,434338 | 0,405481 | 1,01674 | 0,434338 | 0,3448822 | 0,0897988 | Q9NWT6;E! | HIF1AN |
| -0,634215 | -1,23714 | -2,10483 | -1,23714 | 0,7392691 | 0,0899332 | Q86YS3;K7I | RAB11FIP4 |
| 1,47384 | 2,07926 | 0,558669 | 1,47384 | 0,7655356 | 0,09015 | Q9Y5B6 | PAXBP1 |
| -0,789894 | -3,07074 | -2,66881 | -2,66881 | 1,2175201 | 0,0903895 | A0A024RC\ | MSH5-SAPCD1 |
| 5,68375 | 7,50591 | 1,96133 | 5,68375 | 2,826041 | 0,090439 | P19784;H3I | CSNK2A2 |
| 1,22037 | 0,540801 | 1,91195 | 1,22037 | 0,6855833 | 0,0905432 | Q14966 | ZNF638 |
| -2,15106 | -1,0948 | -0,718208 | -1,0948 | 0,7428068 | 0,0911639 | P39656;A0/ | DDOST |
| -0,215072 | -0,414983 | -0,716699 | -0,414983 | 0,2525294 | 0,0912692 | Q9P2J5;A0/ | LARS |
| -1,13447 | -1,12815 | -2,77777 | -1,13447 | 0,9505894 | 0,0921862 | P02787;C9J | TF |
| 0,926228 | 0,238942 | 0,902769 | 0,902769 | 0,3902091 | 0,0922713 | Q7Z7A4;W! | PXK |
| 1,15306 | 0,30174 | 0,816289 | 0,816289 | 0,4287426 | 0,0923457 | Q9UL36;J9J | ZNF236 |
| 2,11045 | 1,12842 | 3,75545 | 2,11045 | 1,3273843 | 0,0931914 | G3V1V8;P1 | MYL2 |
| 3,10513 | 0,974787 | 1,64986 | 1,64986 | 1,0887222 | 0,093388 | F8VWY2;Q! | MGAT4C |
| 1,69139 | 0,769004 | 2,74762 | 1,69139 | 0,9900622 | 0,0934667 | P35499 | SCN4A |
| -1,12349 | -2,67479 | -1,04039 | -1,12349 | 0,9205705 | 0,0935925 | Q9UI12;G3' | ATP6V1H |
| 1,41195 | 0,346965 | 1,19957 | 1,19957 | 0,5636538 | 0,0938213 | G3V321 | ATL1 |
| -0,420146 | -1,70566 | -1,53748 | -1,53748 | 0,698721 | 0,0940045 | P15153;B1/ | RAC2 |
| 4,97338 | 1,6144 | 2,51958 | 2,51958 | 1,7379697 | 0,0940851 | Q9Y5H1 | PCDHGA2 |
| 1,72947 | 0,422225 | 1,43121 | 1,43121 | 0,6850668 | 0,0944031 | A0A087WV | NCAM1 |
| -1,05076 | -1,04362 | -2,60565 | -1,05076 | 0,8997844 | 0,0946052 | Q14289;C9. | PTK2B |
| 2,24932 | 2,15489 | 5,48804 | 2,24932 | 1,8977229 | 0,0949445 | Q14525 | KRT33B |
| -1,87064 | -1,2279 | -0,492889 | -1,2279 | 0,6893903 | 0,0950427 | P08238 | HSP90AB1 |
| -1,24338 | -1,02214 | -2,80951 | -1,24338 | 0,9743717 | 0,0950759 | Q9NUU7;I3 | DDX19A |
| -0,297732 | -1,05345 | -0,622961 | -0,622961 | 0,3790788 | 0,0951001 | P01111 | NRAS |
| -1,85485 | -0,495747 | -2,0436 | -1,85485 | 0,8444561 | 0,0952311 | A0A0A0MR | WDR47 |
| 1,89715 | 3,95243 | 1,32365 | 1,89715 | 1,3822423 | 0,0956749 | P46100;A0/ | ATRX |
| -1,14621 | -0,888847 | -0,27781 | -0,888847 | 0,4460419 | 0,0958099 | P14314;K7E | PRKCSH |
| 0,894318 | 3,75362 | 3,11377 | 3,11377 | 1,50061 | 0,0962229 | Q9P2G3 | KLHL14 |
| -0,752285 | -1,17726 | -2,35108 | -1,17726 | 0,8281106 | 0,0963258 | Q07955;J3k | SRSF1 |
| -0,332657 | -1,03151 | -0,510553 | -0,510553 | 0,3631893 | 0,0965617 | P23258;Q9I | TUBG1 |
| -1,17441 | -1,76046 | -0,44793 | -1,17441 | 0,6575159 | 0,0971106 | P35998;A0/ | PSMC2 |
| -2,07145 | -0,725703 | -0,930373 | -0,930373 | 0,7251415 | 0,0972534 | P61026 | RAB10 |
| -0,32762 | -1,12659 | -1,39574 | -1,12659 | 0,555529 | 0,0975877 | F5H658 | DHX8 |
| -0,228036 | -0,612986 | -0,909766 | -0,612986 | 0,341814 | 0,0978529 | P61981 | YWHAG |
| 3,02457 | 0,729684 | 3,03813 | 3,02457 | 1,3288848 | 0,0982049 | Q15370 | ELOB |
| 0,4451 | 0,509476 | 0,11778 | 0,4451 | 0,210043 | 0,0984008 | Q9UHD1;E! | CHORDC1 |
| -0,467836 | -1,84865 | -1,19503 | -1,19503 | 0,6907336 | 0,0991202 | Q96SI9 | STRBP |
| -1,0989 | -0,266677 | -1,14153 | -1,0989 | 0,4932512 | 0,099151 | P46781;A0/ | RPS9 |
| -0,591237 | -0,168989 | -0,324705 | -0,324705 | 0,2135338 | 0,0992172 | E5RGN3;E5 | ATOX1 |
| 1,43805 | 3,45433 | 1,27407 | 1,43805 | 1,2142081 | 0,0992931 | P47813;O1· | EIF1AX |
| -1,45409 | -0,632933 | -2,40374 | -1,45409 | 0,8861801 | 0,0996633 | Q9UBE0;M( | SAE1 |
| -0,847774 | -2,22321 | -3,40212 | -2,22321 | 1,2784324 | 0,0998057 | Q16795 | NDUFA9 |
| -0,61882 | -1,41432 | -2,35391 | -1,41432 | 0,8685416 | 0,100221 | Q969E4;Q6 | TCEAL3 |
| 0,353714 | 0,528147 | 0,128906 | 0,353714 | 0,2001495 | 0,100254 | A0A087WZ | NAPB |
| -0,575799 | -1,07818 | -0,311657 | -0,575799 | 0,3893831 | 0,100322 | P01112 | HRAS |
| -1,48895 | -0,382825 | -0,919286 | -0,919286 | 0,5531455 | 0,100399 | Q92870;G5 | APBB2 |
| -2,69331 | -0,742819 | -1,51535 | -1,51535 | 0,9822431 | 0,100562 | P51610;A6I | HCFC1 |
| 1,75366 | 2,23219 | 5,08315 | 2,23219 | 1,8001142 | 0,100664 | Q9NYQ7 | CELSR3 |
| 0,099925 | 0,432162 | 0,429448 | 0,429448 | 0,1910385 | 0,100829 | A0A2U3TZ\ | PLCH1 |

| | | | | | | | |
|---|---|---|---|---|---|---|---|
| -2,42292 | -1,80523 | -0,55642 | -1,80523 | 0,9508671 | 0,100877 | F8W8I6;P3: | TIA1 |
| 0,250829 | 0,855581 | 1,10809 | 0,855581 | 0,4405266 | 0,101043 | E5RG59 | ZNF395 |
| -1,15752 | -1,8551 | -3,76357 | -1,8551 | 1,3490967 | 0,101186 | Q9BZK7;A0 | TBL1XR1 |
| 1,20865 | 0,724199 | 0,314328 | 0,724199 | 0,447679 | 0,101295 | Q8IY63 | AMOTL1 |
| 3,19077 | 0,753624 | 3,33938 | 3,19077 | 1,4518896 | 0,101393 | F8VU39;J3K | BAZ2A |
| 0,301679 | 1,08234 | 1,35513 | 1,08234 | 0,5467487 | 0,101631 | Q9C0D5;A0 | TANC1 |
| 0,356924 | 1,2844 | 0,705031 | 0,705031 | 0,4685187 | 0,101695 | Q93050;B7. | ATP6V0A1 |
| 0,149852 | 0,679613 | 0,592794 | 0,592794 | 0,2841308 | 0,101777 | A0A075B7C | APBB1 |
| -0,683795 | -2,6519 | -3,11636 | -2,6519 | 1,2914161 | 0,102109 | Q9NQG5 | RPRD1B |
| 0,750453 | 1,006 | 0,226847 | 0,750453 | 0,3971874 | 0,102205 | A0A2R8YEK | CASK |
| 2,55133 | 0,596462 | 2,68126 | 2,55133 | 1,1679593 | 0,102294 | P46939 | UTRN |
| 0,530139 | 2,20571 | 2,42012 | 2,20571 | 1,0348541 | 0,102591 | O75175;B7. | CNOT3 |
| 4,11657 | 1,00939 | 2,60751 | 2,60751 | 1,5538027 | 0,102773 | Q9Y536;A0. | PPIAL4A |
| 2,15994 | 0,640658 | 2,87543 | 2,15994 | 1,1412237 | 0,102896 | E2QRB3;J3K | PYCR1 |
| -0,456916 | -1,18614 | -0,453488 | -0,456916 | 0,4220107 | 0,103095 | A0A0G2JQN | AP2A2 |
| 1,06482 | 3,13414 | 1,3593 | 1,3593 | 1,1194389 | 0,103193 | A0A0A0MR | ANKRD2 |
| -0,317295 | -0,820124 | -1,29839 | -0,820124 | 0,4905987 | 0,103201 | Q16204 | CCDC6 |
| -0,917024 | -0,592788 | -1,91779 | -0,917024 | 0,6906871 | 0,103286 | A0A0A0MS | PRKACB |
| -2,41006 | -0,789958 | -1,07834 | -1,07834 | 0,8642311 | 0,103715 | B2R6F3;P84 | SFRS3 |
| -0,350158 | -0,0917176 | -0,199631 | -0,199631 | 0,1298044 | 0,104015 | P54577;A0/ | YARS |
| -0,296089 | -1,0405 | -0,539137 | -0,539137 | 0,3796017 | 0,104043 | P00568;Q5 | AK1 |
| -1,37598 | -1,85751 | -0,406602 | -1,37598 | 0,7389969 | 0,104603 | O76054 | SEC14L2 |
| -0,478492 | -0,188095 | -0,175679 | -0,188095 | 0,1713575 | 0,104981 | P50213;H0 | IDH3A |
| -0,733894 | -0,230587 | -1,03788 | -0,733894 | 0,4077269 | 0,105134 | Q16658;H7 | FSCN1 |
| 0,535133 | 0,82069 | 1,74244 | 0,82069 | 0,6309723 | 0,10516 | P46060 | RANGAP1 |
| 1,3236 | 1,49752 | 0,312422 | 1,3236 | 0,6399461 | 0,105661 | A0A0A0MS | OTOGL |
| -1,26226 | -0,579588 | -2,2448 | -1,26226 | 0,8370939 | 0,106197 | A0A1C7CYX | DPYSL2 |
| -0,341585 | -0,930796 | -1,46079 | -0,930796 | 0,5598635 | 0,106201 | Q13315;E9I | ATM |
| -0,513606 | -2,46633 | -2,33522 | -2,33522 | 1,0915279 | 0,106658 | F8VYC4;P0I | RGPD1 |
| -1,34932 | -0,91488 | -2,95712 | -1,34932 | 1,0758333 | 0,10726 | O15056;H7 | SYNJ2 |
| 0,484107 | 1,34063 | 0,522008 | 0,522008 | 0,4839439 | 0,107412 | Q9Y2I7;E9P | PIKFYVE |
| 0,251605 | 1,16926 | 0,824966 | 0,824966 | 0,463568 | 0,107583 | P09622;E9F | DLD |
| 1,53821 | 1,25843 | 0,310366 | 1,25843 | 0,6435194 | 0,108205 | A0A087WV | LLGL1 |
| -1,45922 | -2,82159 | -0,766351 | -1,45922 | 1,0456359 | 0,108253 | F6U0I4;H9K | AGBL2 |
| -0,219894 | -0,438985 | -0,830378 | -0,438985 | 0,309268 | 0,108687 | A2VDJ0;H0 | TMEM131L |
| 3,75201 | 1,92269 | 1,02099 | 1,92269 | 1,3915186 | 0,108826 | Q9UBD5 | ORC3 |
| 0,959227 | 0,95996 | 0,19702 | 0,959227 | 0,4402722 | 0,109024 | O00487 | PSMD14 |
| 1,7305 | 6,01149 | 2,89571 | 2,89571 | 2,2133186 | 0,109038 | P00533 | EGFR |
| -0,94228 | -2,02748 | -0,602159 | -0,94228 | 0,7444104 | 0,109341 | P17066 | HSPA6 |
| 0,249096 | 1,27217 | 1,05274 | 1,05274 | 0,5386207 | 0,11009 | Q5JU69 | TOR2A |
| -2,60155 | -0,507794 | -2,14843 | -2,14843 | 1,101576 | 0,11032 | Q16555;C9. | DPYSL2 |
| 3,31186 | 1,07881 | 1,38563 | 1,38563 | 1,2104416 | 0,110355 | Q7RTS7;F8\ | KRT74 |
| 0,304424 | 1,45534 | 1,55618 | 1,45534 | 0,6954219 | 0,110503 | Q8IUG5 | MYO18B |
| -2,02353 | -0,389384 | -1,76961 | -1,76961 | 0,8793873 | 0,110975 | P06733;A0/ | ENO1 |
| -0,393018 | -1,47934 | -0,755748 | -0,755748 | 0,5530603 | 0,111142 | P25705;K7E | ATP5F1A |
| -0,411184 | -1,30878 | -0,561229 | -0,561229 | 0,4808022 | 0,111432 | A4UGR9;AC | XIRP2 |
| 1,18239 | 6,17156 | 5,12688 | 5,12688 | 2,6312917 | 0,111484 | A0A1B0GTU | KIF16B |
| -0,881899 | -0,218906 | -0,481713 | -0,481713 | 0,3338603 | 0,111609 | Q03113 | GNA12 |
| 0,441394 | 0,17328 | 0,74995 | 0,441394 | 0,2885713 | 0,112051 | B4DYB8;F5( | MPI |
| 0,44236 | 2,04328 | 1,30211 | 1,30211 | 0,8011916 | 0,112101 | J3KSM2;Q9 | MARVELD3 |
| 2,48197 | 0,696171 | 3,4816 | 2,48197 | 1,4110843 | 0,112422 | P49796 | RGS3 |
| -0,329188 | -1,14433 | -0,528129 | -0,528129 | 0,4249976 | 0,112814 | Q9BZE9;J3C | ASPSCR1 |

| | | | | | | | |
|---|---|---|---|---|---|---|---|
| 0,273096 | 1,4298 | 1,39856 | 1,39856 | 0,6589903 | 0,112951 | B1AKV2;B1. | UQCC1 |
| 0,252169 | 1,19765 | 0,777432 | 0,777432 | 0,4737121 | 0,113139 | Q9UQB3;E7 | CTNND2 |
| -2,83054 | -0,525505 | -2,51246 | -2,51246 | 1,2491565 | 0,113289 | A0A024RCF | BAG6 |
| -2,26693 | -0,5339 | -1,28094 | -1,28094 | 0,8692562 | 0,113381 | Q9NQW7;C | XPNPEP1 |
| -1,10599 | -2,1505 | -0,552893 | -1,10599 | 0,811302 | 0,113394 | Q9NX78 | TMEM260 |
| 1,97321 | 4,35444 | 1,26673 | 1,97321 | 1,6177826 | 0,113437 | A0A0G2JM: | LILRB5 |
| 0,590621 | 2,26448 | 1,14367 | 1,14367 | 0,8528269 | 0,113659 | A0A0A0MR | FBXL13 |
| 1,12526 | 1,03782 | 0,208138 | 1,03782 | 0,5061506 | 0,113822 | Q9HD67;A0 | MYO10 |
| 2,94255 | 1,01863 | 1,11447 | 1,11447 | 1,0841686 | 0,113953 | G5EA42;Q9 | TMOD2 |
| -2,08967 | -1,81854 | -0,381952 | -1,81854 | 0,9177506 | 0,114236 | P48643;B7z | CCT5 |
| 2,68012 | 2,03047 | 0,502132 | 2,03047 | 1,1181453 | 0,114756 | A0A087WX | ABCA2 |
| -1,25261 | -2,00348 | -0,424735 | -1,25261 | 0,7896854 | 0,11479 | P19087;A0/ | GNAT2 |
| 1,23864 | 1,88249 | 0,381055 | 1,23864 | 0,7532487 | 0,115271 | Q13098;A0 | GPS1 |
| 2,50938 | 0,563958 | 1,44073 | 1,44073 | 0,9742868 | 0,115945 | O43294 | TGFB1I1 |
| 0,185914 | 0,802107 | 0,444684 | 0,444684 | 0,3094099 | 0,116061 | Q07666 | KHDRBS1 |
| -1,93301 | -3,5404 | -0,83114 | -1,93301 | 1,3624677 | 0,116192 | Q9Y5Q6 | INSL5 |
| 0,969745 | 0,322442 | 0,37016 | 0,37016 | 0,3607354 | 0,116994 | Q14789;E7I | GOLGB1 |
| 0,16206 | 0,0461425 | 0,071901 | 0,071901 | 0,0608674 | 0,117262 | P05388;F8\ | RPLP0 |
| -0,618246 | -0,107198 | -0,529028 | -0,529028 | 0,2729684 | 0,117525 | H7BZJ3 | PDIA3 |
| -2,97518 | -1,15089 | -0,969601 | -1,15089 | 1,1092976 | 0,117612 | H0YKD8;P4 | RPL28 |
| -1,18539 | -0,430864 | -2,01162 | -1,18539 | 0,790649 | 0,117831 | Q5T3I3;Q5T | NAXE |
| -2,22575 | -0,395028 | -1,69249 | -1,69249 | 0,9415693 | 0,118153 | Q15102;M0 | PAFAH1B3 |
| -1,83625 | -0,41913 | -2,38145 | -1,83625 | 1,0129307 | 0,118294 | Q15149;E9I | PLEC |
| 1,2608 | 1,35173 | 0,228808 | 1,2608 | 0,6237293 | 0,119253 | P0CJ78 | ZNF865 |
| -0,376904 | -2,14581 | -2,19531 | -2,14581 | 1,0358635 | 0,119285 | Q14562;K7I | DHX8 |
| 2,15568 | 3,92324 | 0,876072 | 2,15568 | 1,5300816 | 0,119683 | Q86UK0 | ABCA12 |
| -1,16179 | -1,20249 | -0,203802 | -1,16179 | 0,5652102 | 0,119767 | P00387;B1/ | CYB5R3 |
| -0,532318 | -0,799111 | -1,87127 | -0,799111 | 0,7086955 | 0,12084 | P15311;E7E | EZR |
| -0,219546 | -0,427554 | -0,883274 | -0,427554 | 0,3394807 | 0,121334 | P61019;H7 | RAB2A |
| -2,02735 | -2,42115 | -0,393979 | -2,02735 | 1,0748955 | 0,121464 | P11766;H0 | ADH5 |
| 0,354579 | 2,18996 | 1,82641 | 1,82641 | 0,9718606 | 0,121801 | O00763;F8 | ACACB |
| 1,78398 | 1,51117 | 4,73431 | 1,78398 | 1,7873399 | 0,122029 | E7ESX4;P1C | SAG |
| 0,348774 | 1,08863 | 0,406743 | 0,406743 | 0,411444 | 0,12249 | Q9H999;E5 | PANK3 |
| -0,509115 | -0,554315 | -1,53707 | -0,554315 | 0,5808818 | 0,12273 | Q6IQ55;A0, | TTBK2 |
| 0,865058 | 1,48859 | 0,30211 | 0,865058 | 0,5934977 | 0,122823 | Q9BYK8 | HELZ2 |
| -0,324628 | -1,66439 | -1,00842 | -1,00842 | 0,6699291 | 0,122846 | V9GYU8 | EBF4 |
| -1,67813 | -1,6084 | -0,268228 | -1,6084 | 0,7946432 | 0,122885 | P10644;K7E | PRKAR1A |
| -0,470861 | -1,40594 | -0,498182 | -0,498182 | 0,5321566 | 0,123358 | P63000;A4I | RAC1 |
| 0,479908 | 0,765579 | 1,768 | 0,765579 | 0,6764659 | 0,123732 | P49458;E9F | SRP9 |
| -0,0719984 | -0,430349 | -0,307904 | -0,307904 | 0,1821443 | 0,124023 | P49207 | RPL34 |
| 0,988986 | 0,195368 | 1,24376 | 0,988986 | 0,5467884 | 0,124377 | M0R268;P0 | SNRPA |
| -2,13274 | -0,327963 | -1,81072 | -1,81072 | 0,9625907 | 0,124527 | Q5SRE5;H7 | NUP188 |
| 0,413366 | 1,96433 | 2,58728 | 1,96433 | 1,1194834 | 0,124636 | Q13905 | RAPGEF1 |
| 1,09675 | 1,27121 | 0,191574 | 1,09675 | 0,5795679 | 0,125507 | Q14118 | DAG1 |
| 3,26791 | 0,615825 | 4,06328 | 3,26791 | 1,8051368 | 0,126154 | O75140;A0 | DEPDC5 |
| -0,592516 | -0,159961 | -0,915947 | -0,592516 | 0,3793034 | 0,126332 | A0A2U3TZH | SPG7 |
| -0,34842 | -0,51927 | -0,0874805 | -0,34842 | 0,2174555 | 0,126622 | Q07065 | CKAP4 |
| 0,732087 | 1,27747 | 0,247965 | 0,732087 | 0,5150562 | 0,127068 | P60983;G3 | GMFB |
| 0,902405 | 1,36555 | 3,29683 | 1,36555 | 1,2700142 | 0,127133 | Q92508 | PIEZO1 |
| -0,477561 | -3,27904 | -2,89859 | -2,89859 | 1,5195618 | 0,127228 | H7BY49 | TIA1 |
| 0,43981 | 0,112239 | 0,66533 | 0,43981 | 0,2781102 | 0,12734 | P62244;I3L | RPS15A |
| 0,483628 | 0,510374 | 1,47529 | 0,510374 | 0,5649737 | 0,127658 | Q86WR0;B | CCDC25 |

| | | | | | | | |
|---|---|---|---|---|---|---|---|
| 0,168585 | 0,32462 | 0,700394 | 0,32462 | 0,273366 | 0,127865 | Q15652 | JMJD1C |
| 0,298765 | 0,87463 | 1,54415 | 0,87463 | 0,6232791 | 0,128163 | P78347;A0/ | GTF2I |
| -0,590365 | -0,300719 | -1,27071 | -0,590365 | 0,4979369 | 0,12906 | Q92945;A0. | KHSRP |
| 1,29005 | 0,909864 | 3,2519 | 1,29005 | 1,256883 | 0,12925 | Q96DN5;E5 | TBC1D31 |
| -0,398145 | -0,618911 | -1,49484 | -0,618911 | 0,5800473 | 0,129594 | P26038 | MSN |
| -0,894603 | -2,69448 | -0,901305 | -0,901305 | 1,0372302 | 0,129657 | Q16696 | CYP2A13 |
| -0,787194 | -2,83431 | -1,1265 | -1,1265 | 1,0971499 | 0,129736 | P50395;Q6I | GDI2 |
| -1,88758 | -1,02491 | -0,37069 | -1,02491 | 0,7608283 | 0,130339 | Q02224;A0. | CENPE |
| 0,391043 | 0,280329 | 0,0587205 | 0,280329 | 0,1692169 | 0,130374 | Q13813;A0. | SPTAN1 |
| 1,8333 | 1,55289 | 0,252578 | 1,55289 | 0,843418 | 0,130385 | Q6ZNJ1;H0' | NBEAL2 |
| -0,393308 | -0,648796 | -1,53127 | -0,648796 | 0,5970751 | 0,130599 | Q9BQ95 | ECSIT |
| -0,423515 | -1,98621 | -0,994142 | -0,994142 | 0,7907623 | 0,130867 | Q8WWZ7;K | ABCA5 |
| -1,24808 | -1,20867 | -0,17115 | -1,20867 | 0,6107071 | 0,130941 | P62913;Q5' | RPL11 |
| -0,429096 | -2,70986 | -3,15773 | -2,70986 | 1,4633247 | 0,130943 | P62837 | UBE2D2 |
| 1,38323 | 0,273858 | 0,720223 | 0,720223 | 0,5582004 | 0,133149 | Q5QJ74 | TBCEL |
| -1,50916 | -2,56243 | -0,442282 | -1,50916 | 1,0600813 | 0,133192 | P17600;A0/ | SYN1 |
| -0,253463 | -0,114044 | -0,527111 | -0,253463 | 0,210137 | 0,133229 | P62879;E7E | GNB2 |
| -1,83201 | -0,887557 | -0,393007 | -0,887557 | 0,7311294 | 0,133233 | Q9BVA1;K7 | TUBB2B |
| -0,997458 | -1,05442 | -0,135022 | -0,997458 | 0,5151591 | 0,133849 | P19623;K7E | SRM |
| 0,882892 | 0,112445 | 0,769433 | 0,769433 | 0,4159517 | 0,13397 | P48147 | PREP |
| 1,32289 | 2,98486 | 0,701092 | 1,32289 | 1,1807041 | 0,133994 | Q9NZV7 | ZIM2 |
| -0,401086 | -2,21049 | -2,95043 | -2,21049 | 1,3115264 | 0,134065 | Q6NXT6 | TAPT1 |
| 0,391506 | 0,97015 | 0,252302 | 0,391506 | 0,3806819 | 0,134127 | P48735;H0' | IDH2 |
| -0,333894 | -2,52468 | -2,6877 | -2,52468 | 1,3144404 | 0,135163 | P41743 | PRKCI |
| -0,413383 | -1,53319 | -0,591188 | -0,591188 | 0,6017959 | 0,135293 | Q9NSI6;H7I | BRWD1 |
| 0,620965 | 0,648614 | 0,0806711 | 0,620965 | 0,320219 | 0,13531 | H0Y4T4;E7E | CEP170 |
| 2,55208 | 0,739344 | 0,916838 | 0,916838 | 0,9992941 | 0,135591 | Q14593 | ZNF273 |
| -2,8749 | -1,35084 | -0,614195 | -1,35084 | 1,1529811 | 0,136293 | Q10713 | PMPCA |
| 1,21775 | 3,3103 | 0,928156 | 1,21775 | 1,2998231 | 0,136298 | F5GZK2;Q9 | COL21A1 |
| -0,158612 | -1,27699 | -1,02227 | -1,02227 | 0,5861679 | 0,136535 | P49591;Q5' | SARS |
| 0,247415 | 0,498378 | 1,10675 | 0,498378 | 0,4418815 | 0,136574 | P55145;A8I | MANF |
| 3,07469 | 0,404517 | 2,25428 | 2,25428 | 1,3677548 | 0,136599 | H0YAR9;Q8 | THEM6 |
| -2,23161 | -0,424202 | -1,15474 | -1,15474 | 0,9092175 | 0,136645 | Q9Y2L6;E9I | FRMD4B |
| -0,305967 | -2,37992 | -2,55582 | -2,37992 | 1,25127 | 0,136746 | Q7Z460;F8' | CLASP1 |
| 0,509745 | 2,90299 | 1,60267 | 1,60267 | 1,1981193 | 0,136905 | P50402;Q5I | EMD |
| 0,194125 | 0,720121 | 0,27283 | 0,27283 | 0,2837063 | 0,137005 | O75369;E7I | FLNB |
| -1,01673 | -0,233763 | -1,58233 | -1,01673 | 0,6771969 | 0,137059 | Q93045;E5I | STMN2 |
| 0,780157 | 2,56701 | 4,56692 | 2,56701 | 1,8943802 | 0,137348 | Q96CS2 | HAUS1 |
| 0,925139 | 3,27214 | 1,18125 | 1,18125 | 1,2874929 | 0,137356 | P35913;H7( | PDE6B |
| -0,428626 | -3,11408 | -3,61612 | -3,11408 | 1,7138565 | 0,137382 | Q6P2E9 | EDC4 |
| 2,75881 | 0,472959 | 3,72045 | 2,75881 | 1,6681359 | 0,137878 | Q96NY7 | CLIC6 |
| 2,75991 | 0,995833 | 0,774473 | 0,995833 | 1,0880355 | 0,138091 | O00584;A0. | RNASET2 |
| -0,310695 | -2,64019 | -2,20657 | -2,20657 | 1,2388782 | 0,138126 | P13861;H7( | PRKAR2A |
| 3,39102 | 0,648239 | 1,71413 | 1,71413 | 1,3826865 | 0,138232 | Q9Y250 | LZTS1 |
| 1,27977 | 4,95189 | 1,91509 | 1,91509 | 1,962577 | 0,138765 | O00560;G5 | SDCBP |
| 0,878021 | 3,62613 | 1,48695 | 1,48695 | 1,4433173 | 0,138771 | P40692;H0' | MLH1 |
| 1,59553 | 0,376172 | 2,5505 | 1,59553 | 1,0898397 | 0,138852 | P26368;B5E | U2AF2 |
| 5,3207 | 2,98027 | 0,88523 | 2,98027 | 2,2188661 | 0,139356 | Q9Y6I9;C9J | TEX264 |
| 2,88345 | 0,552059 | 1,43318 | 1,43318 | 1,1772171 | 0,139587 | O00629;H7 | KPNA4 |
| -0,0712199 | -0,470941 | -0,603223 | -0,470941 | 0,2769786 | 0,139613 | Q7Z7A1;Q5 | CNTRL |
| 0,409819 | 0,595014 | 1,57432 | 0,595014 | 0,6257529 | 0,140351 | A6NJL1 | ZSCAN5B |
| 0,959471 | 1,86779 | 0,339364 | 0,959471 | 0,7687286 | 0,14048 | P19827 | ITIH1 |

| | | | | | | | |
|---|---|---|---|---|---|---|---|
| 0,694029 | 1,61842 | 0,364704 | 0,694029 | 0,6499689 | 0,140502 | M0QXZ5;Q9 | ZNF428 |
| 1,25582 | 0,133588 | 1,14873 | 1,14873 | 0,6193257 | 0,141635 | Q92973;S4I | TNPO1 |
| 0,552001 | 2,62405 | 1,17869 | 1,17869 | 1,0626376 | 0,141645 | Q9HBR0;H0 | SLC38A10 |
| -1,74672 | -0,597902 | -0,495206 | -0,597902 | 0,6948161 | 0,142247 | F8VVB5 | NAP1L1 |
| 0,320757 | 3,04731 | 2,9385 | 2,9385 | 1,5437244 | 0,142351 | I1E4Y6;Q6Y | GIGYF2 |
| 0,809305 | 2,20677 | 4,41948 | 2,20677 | 1,8203643 | 0,142387 | Q5JQF8 | PABPC1L2A |
| -2,62659 | -0,372175 | -1,61283 | -1,61283 | 1,1291089 | 0,142407 | Q99627;E9I | COPS8 |
| 2,28265 | 0,302013 | 1,48312 | 1,48312 | 0,9964257 | 0,142513 | Q15418;E9I | RPS6KA1 |
| 6,51052 | 1,42315 | 10,3818 | 6,51052 | 4,4930604 | 0,142847 | P61225 | RAP2B |
| -0,96829 | -1,86255 | -0,32212 | -0,96829 | 0,7735375 | 0,142867 | O00408;F5I | PDE2A |
| 1,33663 | 0,263001 | 0,625374 | 0,625374 | 0,5461805 | 0,142993 | S4R3N1;Q9 | HSPE1-MOB4 |
| 4,47514 | 5,26568 | 0,544329 | 4,47514 | 2,5287475 | 0,143355 | J3KTN3 | POLI |
| 0,317515 | 3,09548 | 2,53133 | 2,53133 | 1,4683512 | 0,144425 | A0A0G2JM | PCMTD2 |
| -1,89979 | -0,281608 | -2,58607 | -1,89979 | 1,1832187 | 0,145511 | Q9UK73;H3 | FEM1B |
| 0,423659 | 0,112698 | 0,752714 | 0,423659 | 0,3200506 | 0,145598 | Q9Y3I0 | RTCB |
| -2,27701 | -0,702156 | -0,69221 | -0,702156 | 0,9121271 | 0,145749 | P19022;C9J | CDH2 |
| -0,0920381 | -0,34118 | -0,61099 | -0,34118 | 0,2595445 | 0,145855 | J3KQJ9;Q8N | VWDE |
| -0,125817 | -0,804148 | -0,435633 | -0,435633 | 0,3395885 | 0,145964 | Q6P5Z2 | PKN3 |
| -0,510062 | -0,339482 | -1,36884 | -0,510062 | 0,5516906 | 0,14598 | A0A0U1RRI | EXOC6B |
| 0,203186 | 1,44883 | 1,93552 | 1,44883 | 0,8934463 | 0,146306 | Q13509;G3 | TUBB3 |
| 0,907231 | 5,46868 | 7,98773 | 5,46868 | 3,5890086 | 0,147074 | Q12879;F5( | GRIN2A |
| 3,41797 | 3,69236 | 0,339305 | 3,41797 | 1,8617396 | 0,147114 | Q9Y4G8;A0 | RAPGEF2 |
| 0,326768 | 0,0530656 | 0,473825 | 0,326768 | 0,2135327 | 0,147326 | P17948;H9I | FLT1 |
| -0,275455 | -3,03214 | -2,73895 | -2,73895 | 1,5140498 | 0,147569 | F8VWP7;F8 | CAPS2 |
| 0,23417 | 1,86928 | 1,17166 | 1,17166 | 0,8204822 | 0,147681 | P20336;S4F | RAB3A |
| -0,523044 | -0,200082 | -0,123221 | -0,200082 | 0,2121597 | 0,147827 | Q8IZT6;Q5\ | ASPM |
| -2,04179 | -0,190239 | -1,68846 | -1,68846 | 0,9830013 | 0,14788 | Q16891;B9. | IMMT |
| 0,088894 | 0,863481 | 0,982022 | 0,863481 | 0,4850626 | 0,147901 | Q8TEP8;A0. | CEP192 |
| -0,760521 | -0,37717 | -1,80883 | -0,760521 | 0,7411208 | 0,148616 | P35580;E7E | MYH10 |
| -2,55089 | -0,564433 | -1,01957 | -1,01957 | 1,040681 | 0,148763 | J3KTF0 | FASN |
| -0,283086 | -1,38242 | -0,583846 | -0,583846 | 0,568142 | 0,1496 | P10606 | COX5B |
| -0,0916377 | -0,922078 | -0,670046 | -0,670046 | 0,4257753 | 0,149871 | P26640;A0/ | VARS |
| 0,895751 | 0,093254 | 0,622451 | 0,622451 | 0,4079918 | 0,150161 | Q9UL54 | TAOK2 |
| -0,0881916 | -0,74256 | -0,969521 | -0,74256 | 0,4576117 | 0,151101 | O95071;E7I | UBR5 |
| 0,337401 | 1,09652 | 2,15307 | 1,09652 | 0,9118857 | 0,151129 | Q9BVS4 | RIOK2 |
| -0,0634888 | -0,668016 | -0,776113 | -0,668016 | 0,3840511 | 0,151614 | P59998;F8\ | ARPC4 |
| -2,90761 | -1,2848 | -0,547009 | -1,2848 | 1,2076344 | 0,151677 | P40926;G3) | MDH2 |
| -3,01253 | -0,295018 | -3,59055 | -3,01253 | 1,7597116 | 0,151951 | E5RFH5;F5H | ASH2L |
| -1,77625 | -0,155363 | -1,38071 | -1,38071 | 0,8451037 | 0,151987 | O60888;C9I | CUTA |
| -0,458453 | -0,235511 | -1,1315 | -0,458453 | 0,4664568 | 0,15235 | A0A0A0MR | CYP4F12 |
| -0,0307848 | -0,374647 | -0,307764 | -0,307764 | 0,1823148 | 0,152446 | P04179;F5( | SOD2 |
| -0,626424 | -0,657874 | -2,18056 | -0,657874 | 0,8883412 | 0,153151 | Q9UL01;A0 | DSE |
| 3,20091 | 3,71966 | 0,289557 | 3,20091 | 1,8489044 | 0,153194 | Q5JPT1;Q5. | SH3KBP1 |
| -2,04864 | -2,6929 | -0,228571 | -2,04864 | 1,278061 | 0,153864 | Q01518;Q5 | CAP1 |
| 0,131664 | 1,65078 | 1,31889 | 1,31889 | 0,7986833 | 0,154219 | Q6UB99;X5 | ANKRD11 |
| -2,79714 | -2,05163 | -0,245554 | -2,05163 | 1,3120142 | 0,154234 | P23528;E9F | CFL1 |
| 3,12847 | 0,57308 | 5,09165 | 3,12847 | 2,2657438 | 0,154351 | Q9Y5G4 | PCDHGA9 |
| 0,401065 | 0,282967 | 1,1584 | 0,401065 | 0,4750239 | 0,154496 | Q13619;A0. | CUL4A |
| -0,16605 | -1,91211 | -1,38277 | -1,38277 | 0,8952964 | 0,155302 | Q8NDM7 | CFAP43 |
| -2,22719 | -0,281422 | -1,24708 | -1,24708 | 0,9728929 | 0,155637 | P00558;F5H | PGK1 |
| -0,101976 | -1,36451 | -1,47746 | -1,36451 | 0,7636214 | 0,155957 | H0YDG0 | NPEPPS |
| -1,97699 | -0,224141 | -1,1804 | -1,1804 | 0,8776357 | 0,156099 | Q13263;M( | TRIM28 |

| | | | | | | | |
|---|---|---|---|---|---|---|---|
| -1,16512 | -0,0913894 | -0,893405 | -0,893405 | 0,5582645 | 0,15622 | J3KR69;Q9I | TTC12 |
| -0,366335 | -0,153315 | -0,845682 | -0,366335 | 0,3546179 | 0,156279 | P35555;F6U | FBN1 |
| -1,52481 | -1,19219 | -0,113242 | -1,19219 | 0,7379351 | 0,157216 | P11021;O9! | HSPA5 |
| 1,58007 | 0,131651 | 1,90277 | 1,58007 | 0,9433023 | 0,157447 | Q14152 | EIF3A |
| 1,3705 | 0,272687 | 0,545996 | 0,545996 | 0,5715036 | 0,157522 | Q9H0X9;H0 | OSBPL5 |
| 0,57438 | 0,298011 | 1,47329 | 0,57438 | 0,6145052 | 0,158379 | Q96NL3 | ZNF599 |
| 2,68432 | 0,168237 | 2,72676 | 2,68432 | 1,4650663 | 0,158956 | P43155;A6I | CRAT |
| -0,307873 | -3,22826 | -2,04856 | -2,04856 | 1,4691462 | 0,159404 | Q86UQ4;A( | ABCA13 |
| -0,288179 | -0,558927 | -1,4395 | -0,558927 | 0,6019764 | 0,159587 | Q9NZI8 | IGF2BP1 |
| 0,128153 | 1,26038 | 1,73872 | 1,26038 | 0,8271108 | 0,160734 | Q8NBJ4;C9. | GOLM1 |
| -0,605874 | -1,32227 | -0,206937 | -0,605874 | 0,5651462 | 0,160932 | P50993;B1/ | ATP1A2 |
| -2,85973 | -0,214901 | -2,02913 | -2,02913 | 1,3525558 | 0,161228 | P26358;K7I | DNMT1 |
| -0,862422 | -1,10325 | -0,0688119 | -0,862422 | 0,5412759 | 0,162203 | P25398 | RPS12 |
| 1,34078 | 0,76571 | 0,143219 | 0,76571 | 0,598937 | 0,162371 | P47756;B1/ | CAPZB |
| -0,949808 | -0,215109 | -1,78466 | -0,949808 | 0,7853079 | 0,162385 | P16949;B5I | STMN1 |
| -0,437299 | -1,27935 | -2,82609 | -1,27935 | 1,2115952 | 0,162822 | Q86WI3 | NLRC5 |
| 1,84432 | 4,06061 | 0,623963 | 1,84432 | 1,7422091 | 0,16295 | Q08AM6;H. | VAC14 |
| 0,147543 | 1,07005 | 1,69021 | 1,07005 | 0,7762559 | 0,163054 | P31689 | DNAJA1 |
| -0,105519 | -0,462663 | -0,875927 | -0,462663 | 0,3855445 | 0,163074 | Q9UEY8 | ADD3 |
| 0,272478 | 0,389752 | 1,17463 | 0,389752 | 0,490521 | 0,163136 | P43686 | PSMC4 |
| 1,83925 | 1,87335 | 0,0930331 | 1,83925 | 1,0181654 | 0,163603 | Q9H0W8;N | SMG9 |
| -3,06212 | -0,151451 | -2,93834 | -2,93834 | 1,6459073 | 0,163606 | Q99250;A0. | SCN2A |
| 4,18383 | 0,454105 | 2,33157 | 2,33157 | 1,8648767 | 0,163636 | Q5T035 | C9orf129 |
| 0,781081 | 0,24448 | 1,67796 | 0,781081 | 0,7242464 | 0,163931 | P42765;A0/ | ACAA2 |
| 5,35864 | 0,34408 | 3,94077 | 3,94077 | 2,5849676 | 0,164084 | Q8TBB0 | THAP6 |
| 0,787786 | 1,48588 | 3,97672 | 1,48588 | 1,6763512 | 0,164222 | P37235 | HPCAL1 |
| -0,193921 | -0,440775 | -1,09009 | -0,440775 | 0,4629013 | 0,164393 | Q07021;I3L | C1QBP |
| -1,59051 | -2,2197 | -0,146798 | -1,59051 | 1,0627878 | 0,164582 | Q8N9H8 | EXD3 |
| -0,562676 | -0,181329 | -1,22889 | -0,562676 | 0,5301966 | 0,16473 | Q13042 | CDC16 |
| 3,13119 | 1,9903 | 0,258358 | 1,9903 | 1,446514 | 0,164857 | Q6ZNA1;A0 | ZNF836 |
| 8,2139 | 2,23343 | 2,30915 | 2,30915 | 3,4311764 | 0,16495 | Q969P6;E5I | TOP1MT |
| -1,25413 | -0,221453 | -2,19913 | -1,25413 | 0,9891624 | 0,165143 | Q13393;C9 | PLD1 |
| -2,95089 | -0,134576 | -2,9524 | -2,95089 | 1,6264357 | 0,165324 | P98164 | LRP2 |
| -1,90591 | -0,0928251 | -1,61956 | -1,61956 | 0,9746959 | 0,16533 | H0Y7W5 | RFX3 |
| -1,32653 | -0,220767 | -2,3015 | -1,32653 | 1,0410514 | 0,166372 | Q9P2M7;A( | CGN |
| 3,22601 | 2,54008 | 0,165258 | 2,54008 | 1,6061591 | 0,166658 | Q9HA77 | CARS2 |
| -0,0670597 | -0,887032 | -1,18909 | -0,887032 | 0,5805953 | 0,166763 | A6QL64;A0. | ANKRD36 |
| -0,905586 | -0,234478 | -1,84763 | -0,905586 | 0,8103592 | 0,167072 | Q9NYC9;E7 | DNAH9 |
| -1,23477 | -0,188948 | -0,535915 | -0,535915 | 0,5326863 | 0,167633 | O60256;E7I | PRPSAP2 |
| 0,135587 | 3,39932 | 3,04909 | 3,04909 | 1,7917921 | 0,167929 | Q9Y5H0 | PCDHGA3 |
| -0,969571 | -0,0516821 | -0,723603 | -0,723603 | 0,4751312 | 0,16808 | Q92841;A0. | DDX17 |
| 2,04163 | 0,876012 | 0,315654 | 0,876012 | 0,8804979 | 0,168097 | Q7Z4G4;K7 | TRMT11 |
| -1,30571 | -0,663585 | -0,152184 | -0,663585 | 0,5779962 | 0,168217 | A0A2R8Y79 | ACTB |
| -0,561093 | -1,89855 | -0,469229 | -0,561093 | 0,8000197 | 0,168876 | P18206;A0/ | VCL |
| -1,36203 | -2,33186 | -0,204465 | -1,36203 | 1,0650772 | 0,168934 | C9IZY8;Q96 | FMNL2 |
| -1,05685 | -2,00163 | -0,205068 | -1,05685 | 0,8986821 | 0,170966 | P80404;H3I | ABAT |
| -0,751218 | -0,493997 | -2,2993 | -0,751218 | 0,9765449 | 0,171096 | O75915;C9. | ARL6IP5 |
| -0,711358 | -0,0229062 | -0,776596 | -0,711358 | 0,4175864 | 0,171925 | P62310 | LSM3 |
| 0,205823 | 3,21323 | 4,40763 | 3,21323 | 2,1651124 | 0,172154 | Q58FF3 | HSP90B2P |
| -2,57706 | -0,793595 | -0,581489 | -0,793595 | 1,0960566 | 0,172813 | Q96R06;J3I | SPAG5 |
| -0,0824262 | -1,96598 | -1,4346 | -1,4346 | 0,9711259 | 0,174213 | E9PR30;P6. | FAU |
| -1,99641 | -0,348736 | -0,741316 | -0,741316 | 0,8606402 | 0,174236 | P11142;E9I | HSPA8 |

| | | | | | | | |
|---|---|---|---|---|---|---|---|
| -2,09297 | -0,142214 | -1,27745 | -1,27745 | 0,9797349 | 0,174306 | P61163;R4( | ACTR1A |
| -0,456658 | -0,547293 | -1,9129 | -0,547293 | 0,8158572 | 0,175048 | O15061;A0. | SYNM |
| 0,200746 | 0,779956 | 1,67647 | 0,779956 | 0,7435257 | 0,175155 | E7ENS9;H7I | CLEC4M |
| -0,637701 | -0,217917 | -1,52564 | -0,637701 | 0,6676818 | 0,175694 | Q15029;K7I | EFTUD2 |
| 0,968941 | 0,0164614 | 0,995826 | 0,968941 | 0,5578374 | 0,176797 | Q6W2J9;H7 | BCOR |
| -1,96056 | -1,06034 | -0,165852 | -1,06034 | 0,8973555 | 0,176824 | C9JPM4;P1 | ARF4 |
| 1,72151 | 1,41296 | 0,0409265 | 1,41296 | 0,8946172 | 0,17696 | P38117;M0 | ETFB |
| 0,0644095 | 0,68489 | 1,10024 | 0,68489 | 0,5212895 | 0,177068 | Q96HN2;H( | AHCYL2 |
| 0,643192 | 2,28585 | 0,538313 | 0,643192 | 0,9800689 | 0,177829 | O95573 | ACSL3 |
| 0,33691 | 2,71042 | 1,19986 | 1,19986 | 1,2013897 | 0,178028 | Q16531;F5( | DDB1 |
| 0,542382 | 0,177075 | 1,29428 | 0,542382 | 0,5696412 | 0,178038 | Q16186;A0. | ADRM1 |
| -0,153761 | -0,13309 | -0,556088 | -0,153761 | 0,2384749 | 0,178068 | P36776;K7E | LONP1 |
| -1,10627 | -0,112443 | -1,83371 | -1,10627 | 0,8640622 | 0,178223 | Q9UL58 | ZNF215 |
| -1,82146 | -0,254802 | -0,750276 | -0,750276 | 0,8007649 | 0,178438 | Q99996;A0. | AKAP9 |
| 1,22262 | 0,449945 | 0,203185 | 0,449945 | 0,5318466 | 0,178661 | O94988;D6 | FAM13A |
| -0,0371059 | -2,56632 | -2,25943 | -2,25943 | 1,3802071 | 0,178932 | Q7KZN9 | COX15 |
| 0,542548 | 0,414176 | 1,85915 | 0,542548 | 0,7997781 | 0,179119 | O95782 | AP2A1 |
| -0,15585 | -1,70851 | -2,7818 | -1,70851 | 1,3202473 | 0,179246 | P61244;G3' | MAX |
| 0,932091 | 6,68396 | 2,73006 | 2,73006 | 2,942505 | 0,179479 | C9JYJ8;C9JZ | REPIN1 |
| 0,603849 | 0,658197 | 0,00713861 | 0,603849 | 0,3612234 | 0,179671 | Q5SW79;H( | CEP170 |
| -1,27716 | -0,339408 | -0,312617 | -0,339408 | 0,5493086 | 0,17979 | Q01844;A0. | EWSR1 |
| -1,83471 | -0,35615 | -0,590236 | -0,590236 | 0,7947381 | 0,180759 | P22695;H3I | UQCRC2 |
| -2,07744 | -3,2793 | -0,155559 | -2,07744 | 1,5756402 | 0,180832 | H0YEF6 | TTC12 |
| -0,233783 | -1,3643 | -2,6836 | -1,3643 | 1,2261202 | 0,181324 | Q9UMR2;H | DDX19B |
| 0,958985 | 2,2495 | 0,279484 | 0,958985 | 1,0006759 | 0,181824 | E9PKC0;Q6 | PLEKHA7 |
| -0,454486 | -1,18621 | -3,078 | -1,18621 | 1,3538288 | 0,181837 | P62942;A0/ | FKBP1A |
| -0,463148 | -1,12346 | -0,148015 | -0,463148 | 0,4977974 | 0,181902 | Q9P0X4 | CACNA1I |
| -2,54555 | -0,289409 | -1,13335 | -1,13335 | 1,1399355 | 0,182169 | H0YG02 | CHN2 |
| -0,664665 | -1,69583 | -0,242146 | -0,664665 | 0,7477768 | 0,18222 | F5GXQ8 | SYNE1 |
| 0,319959 | 3,36207 | 1,62864 | 1,62864 | 1,5259896 | 0,182248 | Q96MZ0;A( | GDAP1L1 |
| -0,00542685 | -2,16945 | -2,2582 | -2,16945 | 1,2757912 | 0,182668 | M0QYA8;Q | VRK3 |
| 0,576582 | 0,0182541 | 0,392732 | 0,392732 | 0,284536 | 0,182978 | A0A1W2PR | ADGRV1 |
| 1,08749 | 2,04728 | 0,152757 | 1,08749 | 0,9472891 | 0,183005 | I3L119 | ZNF785 |
| -0,0951594 | -2,29837 | -1,45823 | -1,45823 | 1,1119003 | 0,183502 | Q7Z3U7;A0 | MON2 |
| -0,241393 | -0,82713 | -0,174561 | -0,241393 | 0,3590266 | 0,183639 | Q9BY12;H3 | SCAPER |
| 1,0864 | 0,643851 | 0,0571868 | 0,643851 | 0,5162855 | 0,18366 | P46459;I3L( | NSF |
| -0,0475031 | -3,06133 | -3,88619 | -3,06133 | 2,0206873 | 0,183692 | P02549 | SPTA1 |
| 0,00516988 | -3,28203 | -3,3207 | -3,28203 | 1,9091267 | 0,184157 | P28340;M0 | POLD1 |
| -1,28371 | -1,25733 | 0,0030818 | -1,25733 | 0,7354326 | 0,184539 | Q9P2D0;E7 | IBTK |
| -0,243421 | -0,340746 | -1,16932 | -0,340746 | 0,5088051 | 0,184911 | P51003;G3) | PAPOLA |
| -2,88443 | -1,85243 | -0,102557 | -1,85243 | 1,4062893 | 0,185309 | P08865;C9J | RPSA |
| 3,10589 | 1,19625 | 0,433402 | 1,19625 | 1,3766417 | 0,185416 | Q92824 | PCSK5 |
| -0,00176033 | 0,871018 | 0,77855 | 0,77855 | 0,47944 | 0,185654 | Q5S007;E9I | LRRK2 |
| -0,557915 | -0,655849 | -0,00015125 | -0,557915 | 0,3537021 | 0,186047 | Q9NQW1 | SEC31B |
| 6,92065 | 4,06595 | 0,33682 | 4,06595 | 3,3015789 | 0,186233 | Q9Y2G4;E5 | ANKRD6 |
| -0,781011 | -0,260598 | -0,132899 | -0,260598 | 0,3433137 | 0,186925 | P30040;F8\ | ERP29 |
| -1,41035 | -0,0987911 | -0,733157 | -0,733157 | 0,655896 | 0,18712 | P63244;J3K | RACK1 |
| 3,52601 | 0,612133 | 1,15521 | 1,15521 | 1,5495324 | 0,187328 | P23921;E9F | RRM1 |
| 0,951217 | 0,368311 | 0,124899 | 0,368311 | 0,4246234 | 0,188499 | Q9NRN7;E9 | AASDHPPT |
| -1,88834 | -0,349517 | -0,581058 | -0,581058 | 0,8297162 | 0,188843 | Q8WWI1;J3 | LMO7 |
| -0,0553331 | -1,68545 | -2,54355 | -1,68545 | 1,2639119 | 0,189473 | F5H2F4 | MTHFD1 |
| -0,055331 | -1,68545 | -2,54355 | -1,68545 | 1,2639131 | 0,189474 | P11586;V9( | MTHFD1 |

| | | | | | | | |
|---|---|---|---|---|---|---|---|
| 1,76891 | 1,94289 | -0,0245367 | 1,76891 | 1,0891501 | 0,189826 | Q9ULE0 | WWC3 |
| -0,273953 | -0,851612 | -2,19118 | -0,851612 | 0,9835219 | 0,19091 | O95757;E9I | HSPA4L |
| 1,47364 | -0,00203896 | 1,13495 | 1,13495 | 0,7729902 | 0,190933 | Q8TB36 | GDAP1 |
| -2,83706 | -0,617044 | -0,746844 | -0,746844 | 1,2459482 | 0,190962 | A0A2R8YEC | KCNQ1 |
| 1,55595 | 0,286814 | 0,469634 | 0,469634 | 0,6860772 | 0,191064 | Q9NPI1 | BRD7 |
| -0,015741 | 1,27369 | 1,07815 | 1,07815 | 0,6949179 | 0,191793 | E7EMS2;G3 | NPC2 |
| 0,869328 | -0,0142229 | 0,779705 | 0,779705 | 0,4863154 | 0,191798 | Q6FI81;H3E | CIAPIN1 |
| 0,0638938 | 1,44257 | 0,813644 | 0,813644 | 0,6902199 | 0,191817 | O95292;E5I | VAPB |
| 0,836375 | 1,13112 | -3,26E-05 | 0,836375 | 0,5867933 | 0,192527 | Q9BZF9;F5I | UACA |
| -1,71732 | -1,56539 | 0,0335796 | -1,56539 | 0,9700031 | 0,192804 | P04792;F8V | HSPB1 |
| 0,588063 | 0,687619 | -0,011176 | 0,588063 | 0,378002 | 0,193172 | Q92522 | H1FX |
| 2,50376 | -0,0479218 | 2,20604 | 2,20604 | 1,3952336 | 0,193504 | F5H829;Q9 | CHFR |
| -0,191941 | -3,12255 | -1,57199 | -1,57199 | 1,466131 | 0,194213 | A0A1W2PP | hCG_1809904 |
| 0,0345204 | -2,08957 | -2,5967 | -2,08957 | 1,3959621 | 0,194266 | E7ERI8;E7E | CLASP2 |
| 3,48814 | 5,44175 | 0,0907649 | 3,48814 | 2,7077602 | 0,19434 | O95398 | RAPGEF3 |
| 1,75403 | 0,959359 | 0,077951 | 0,959359 | 0,8384135 | 0,194518 | A0A1B0GV | CLCN2 |
| -2,47772 | -1,17467 | -0,179488 | -1,17467 | 1,1525477 | 0,194911 | O60506;F6I | SYNCRIP |
| 0,328175 | 0,347032 | 1,44094 | 0,347032 | 0,6370814 | 0,195172 | Q8IZH2;H7 | XRN1 |
| -0,0483442 | -1,43874 | -2,32941 | -1,43874 | 1,1496198 | 0,19533 | Q9C0E4;A0 | GRIP2 |
| 4,08982 | 5,18643 | -0,0702193 | 4,08982 | 2,7731102 | 0,195333 | Q86UE8;J3I | TLK2 |
| -2,29137 | -0,328364 | -0,788181 | -0,788181 | 1,0266776 | 0,195367 | P62241;Q5. | RPS8 |
| -2,26409 | -3,40992 | -0,0263484 | -2,26409 | 1,7208995 | 0,19596 | O60443 | GSDME |
| 1,79764 | 1,02431 | 0,0595226 | 1,02431 | 0,8708144 | 0,196259 | O75030 | MITF |
| -0,0352414 | -1,06698 | -1,74205 | -1,06698 | 0,8595929 | 0,196266 | P31946;A0 | YWHAB |
| -0,0594466 | 1,91342 | 2,03078 | 1,91342 | 1,1743809 | 0,196348 | F5H4N6;Q9 | GRIP1 |
| -0,78444 | -0,22283 | -0,144757 | -0,22283 | 0,3489736 | 0,196928 | P27694;I3L | RPA1 |
| 0,393324 | 0,518466 | -0,00665438 | 0,393324 | 0,2742853 | 0,197032 | O14531;Q5 | DPYSL4 |
| 1,63313 | 1,22438 | -0,0200152 | 1,22438 | 0,8610541 | 0,197431 | Q5JWF2;P6 | GNAS |
| -2,46521 | -0,366442 | -0,811064 | -0,811064 | 1,1059472 | 0,197572 | Q93009;H3 | USP7 |
| 0,0597788 | 0,557505 | 0,219576 | 0,219576 | 0,2541202 | 0,197622 | Q9UQ35;I3 | SRRM2 |
| -0,597412 | -0,328535 | -0,0219472 | -0,328535 | 0,2879383 | 0,197723 | P27448;A0 | MARK3 |
| -1,72553 | -0,0300274 | -2,72157 | -1,72553 | 1,3608347 | 0,197898 | Q96JF6;I3L | ZNF594 |
| 1,0375 | 1,38802 | -0,0181035 | 1,0375 | 0,731932 | 0,197974 | A6NKB5;H0 | PCNX2 |
| -1,24299 | -0,0478773 | -0,67451 | -0,67451 | 0,5977921 | 0,198095 | A0A2R8Y7Y | ANKRD44 |
| 1,29512 | -0,0465285 | 1,29578 | 1,29512 | 0,7747917 | 0,198425 | P51531;F6 | SMARCA2 |
| 0 | 0,0868616 | 0,128694 | 0,0868616 | 0,0656468 | 0,198458 | E9PMP2 | PAK1 |
| -2,42781 | -0,679302 | -0,446109 | -0,679302 | 1,0831126 | 0,19872 | O75874;C9. | IDH1 |
| 5,29215 | 0,182761 | 2,89416 | 2,89416 | 2,556296 | 0,199303 | Q9P253;H0 | VPS18 |
| 0,0495052 | -1,28307 | -1,3256 | -1,28307 | 0,7819292 | 0,199393 | C9JXG5;F8V | THOC5 |
| 0,189055 | 0,543194 | 1,51723 | 0,543194 | 0,6877753 | 0,199588 | Q9H3N1;G3 | TMX1 |
| -0,742973 | -0,111462 | -1,62169 | -0,742973 | 0,7584786 | 0,200123 | Q8NC51 | SERBP1 |
| 4,46676 | 3,59396 | -0,118699 | 3,59396 | 2,4348871 | 0,200373 | Q9H361 | PABPC3 |
| 2,3248 | 0,132771 | 1,12236 | 1,12236 | 1,0977355 | 0,200423 | P42338;H0 | PIK3CB |
| 0,0408812 | -1,02675 | -0,973618 | -0,973618 | 0,6016461 | 0,200806 | P52701;A0 | MSH6 |
| 2,81697 | 0,581629 | 0,698283 | 0,698283 | 1,2582522 | 0,200882 | B4DDG0;B4 | TSPAN7 |
| -2,72221 | 0,103075 | -2,47556 | -2,47556 | 1,5648445 | 0,200908 | G3XAJ6;Q1 | RFTN1 |
| -1,84128 | 0,062367 | -1,58119 | -1,58119 | 1,0322143 | 0,200951 | Q6AI08;K7E | HEATR6 |
| -0,346809 | -0,922583 | -2,67365 | -0,922583 | 1,2118815 | 0,201092 | Q9Y228;E2 | TRAF3IP3 |
| 0,121933 | 0,113077 | 0,519574 | 0,121933 | 0,2321769 | 0,201413 | Q9UN86;D | G3BP2 |
| 1,61713 | 2,5324 | 0,00538862 | 1,61713 | 1,2794019 | 0,201635 | Q86XA9;F5 | HEATR5A |
| -0,716827 | -0,171011 | -0,1529 | -0,171011 | 0,3204832 | 0,201648 | P60866;E5F | RPS20 |
| -0,389867 | -0,430365 | -1,82406 | -0,430365 | 0,816592 | 0,20247 | Q08211 | DHX9 |

| | | | | | | | |
|---|---|---|---|---|---|---|---|
| 0,0610264 | -1,37572 | -1,61016 | -1,37572 | 0,9048081 | 0,202976 | P08237;A0/ | PFKM |
| -2,0415 | -0,51447 | -0,400928 | -0,51447 | 0,9161686 | 0,203436 | Q15027;I3L | ACAP1 |
| 0,00544092 | -1,01987 | -1,58047 | -1,01987 | 0,8042231 | 0,203515 | B5BU24 | YWHAB |
| 3,37774 | 1,16992 | 0,41098 | 1,16992 | 1,5412183 | 0,20435 | A0A1B0GU! | UNC13B |
| -0,039723 | -0,57663 | -1,1215 | -0,57663 | 0,5408934 | 0,204753 | P51812;B1/ | RPS6KA3 |
| -1,79519 | -0,701505 | -0,167919 | -0,701505 | 0,8295452 | 0,204825 | P10515;E9F | DLAT |
| -0,089626 | -1,1376 | -0,474637 | -0,474637 | 0,5300948 | 0,204975 | Q9BUF5;K7 | TUBB6 |
| -0,439425 | -0,610502 | 0,0140872 | -0,439425 | 0,3227621 | 0,205083 | P78559 | MAP1A |
| 1,32793 | -0,0102581 | 0,861501 | 0,861501 | 0,679248 | 0,205182 | Q16851;A0 | UGP2 |
| -0,641735 | -3,12468 | -0,740113 | -0,740113 | 1,4059904 | 0,205454 | Q9H7Z3;G3 | NRDE2 |
| -1,68053 | 0,0895941 | -1,76527 | -1,68053 | 1,0473014 | 0,20551 | Q9NUL3;AC | STAU2 |
| -1,33049 | -0,3755 | -0,220285 | -0,3755 | 0,6012006 | 0,205563 | Q5VTE0;P6 | EEF1A1P5 |
| 0,0500469 | 2,58474 | 1,41881 | 1,41881 | 1,2686984 | 0,206385 | A0A0G2JLC | NLRP2 |
| 0,16438 | 2,2204 | 0,929101 | 0,929101 | 1,0391879 | 0,206953 | C9IZS8;Q03 | ZNF92 |
| 0,0926054 | -1,94077 | -2,51509 | -1,94077 | 1,3701906 | 0,207369 | P21399 | ACO1 |
| -0,364413 | -0,1003 | -0,98982 | -0,364413 | 0,4568252 | 0,207407 | E9PG32;Q6 | DNAH12 |
| 0,913025 | 0,0910098 | 0,338299 | 0,338299 | 0,4217367 | 0,207512 | A0A1C7CY> | ELMSAN1 |
| 0,52481 | 0,765191 | 2,93567 | 0,765191 | 1,3279689 | 0,207587 | Q9Y5J7 | TIMM9 |
| -0,0511832 | 2,93487 | 1,95697 | 1,95697 | 1,5223601 | 0,207805 | Q16587;F8' | ZNF74 |
| -0,814348 | -0,166762 | -2,0389 | -0,814348 | 0,9507713 | 0,208114 | P46109 | CRKL |
| 1,78032 | 0,867762 | 5,94219 | 1,78032 | 2,7050491 | 0,208183 | Q9UHY7 | ENOPH1 |
| 2,10508 | 0,318777 | 0,615472 | 0,615472 | 0,9572389 | 0,208233 | J3QSF4 | RAB15 |
| -2,26382 | -0,998384 | -0,130875 | -0,998384 | 1,0726412 | 0,209336 | H0Y8R5 | RACK1 |
| 0,429431 | 0,104335 | 1,14642 | 0,429431 | 0,5331825 | 0,210469 | Q2LD37 | KIAA1109 |
| -1,65638 | -0,767419 | -0,0744603 | -0,767419 | 0,792981 | 0,210542 | P61604;B8 | HSPE1 |
| -0,0828447 | 1,60041 | 1,31849 | 1,31849 | 0,9015324 | 0,210981 | Q15185;A0 | PTGES3 |
| -2,67322 | 0,0675676 | -1,79515 | -1,79515 | 1,3995619 | 0,211114 | E9PLG0;E9F | GEN1 |
| -1,00946 | 0,0189488 | -1,63664 | -1,00946 | 0,8358582 | 0,211243 | Q14157;F8' | UBAP2L |
| -0,20335 | 3,1795 | 2,95547 | 2,95547 | 1,8917367 | 0,211957 | P49720;A0/ | PSMB3 |
| 2,58226 | 0,263477 | 5,69206 | 2,58226 | 2,7238797 | 0,212064 | Q16650;H7 | TBR1 |
| 0,00497422 | 0,565235 | 1,0135 | 0,565235 | 0,5052982 | 0,212084 | A0A0G2JP8 | CCHCR1 |
| 3,96201 | -0,107137 | 2,65689 | 2,65689 | 2,0777047 | 0,212093 | Q9NTJ3;E9I | SMC4 |
| -0,0212921 | 1,1099 | 1,83433 | 1,1099 | 0,9352119 | 0,212923 | O60271;A0 | SPAG9 |
| -0,188786 | 2,55274 | 3,00192 | 2,55274 | 1,7271524 | 0,214721 | Q5FVE4;K7I | ACSBG2 |
| 0,00645549 | 1,20183 | 0,650297 | 0,650297 | 0,598281 | 0,214744 | P49748;G3' | ACADVL |
| -2,26259 | -0,253228 | -0,735157 | -0,735157 | 1,0490347 | 0,215474 | A0A0A0MR | APP |
| 6,8349 | 2,29623 | 0,714243 | 2,29623 | 3,1771227 | 0,215492 | Q9BZ67 | FRMD8 |
| 0,0783551 | -1,03631 | -1,24008 | -1,03631 | 0,7097267 | 0,215669 | Q8WUD1;C | RAB2B |
| 0,344167 | 1,24884 | 3,591 | 1,24884 | 1,6756129 | 0,215983 | Q92930;H0 | RAB8B |
| -2,75041 | -0,0746802 | -1,30815 | -1,30815 | 1,3392219 | 0,216717 | F2Z2K5;Q9 | JAKMIP1 |
| 0,00159048 | -0,612754 | -0,337277 | -0,337277 | 0,3077168 | 0,217121 | P31327;Q5I | CPS1 |
| 0,0336142 | -1,45864 | -0,893252 | -0,893252 | 0,7533887 | 0,217618 | F8W9U4 | MAP4 |
| 2,3276 | 4,56403 | 0,0494264 | 2,3276 | 2,257334 | 0,217839 | Q8NFZ4 | NLGN2 |
| 1,18788 | 0,764376 | 4,7196 | 1,18788 | 2,171643 | 0,218099 | P33316;H0' | DUT |
| -0,344371 | -0,0549874 | -0,870394 | -0,344371 | 0,4133866 | 0,218166 | O75410;R4 | TACC1 |
| 3,90599 | 4,21285 | -0,326642 | 3,90599 | 2,5369379 | 0,218175 | Q6PCB5;C9 | RSBN1L |
| -2,44587 | 0,212265 | -2,78024 | -2,44587 | 1,6397444 | 0,219546 | Q96JI7;C4B | SPG11 |
| -0,123317 | 2,25702 | 1,64381 | 1,64381 | 1,2359051 | 0,219668 | Q9NXZ1;F5 | SAGE1 |
| -0,0288093 | 3,7452 | 2,05658 | 2,05658 | 1,8904776 | 0,219945 | Q86U06;G3 | RBM23 |
| 1,49693 | 2,45838 | -0,0676024 | 1,49693 | 1,2749336 | 0,220382 | Q13564;H3 | NAE1 |
| 4,24949 | 0,493032 | 1,29066 | 1,29066 | 1,9791354 | 0,220477 | Q8TC12;G3 | RDH11 |
| -2,27331 | -0,768528 | -0,207386 | -0,768528 | 1,0682767 | 0,221165 | Q9NYB9;AC | ABI2 |

| | | | | | | | |
|---|---|---|---|---|---|---|---|
| 0,401568 | 0,377875 | 1,93745 | 0,401568 | 0,89366 | 0,221302 | A0A0A0MR | CDC42BPA |
| 1,31447 | 1,80407 | -0,10519 | 1,31447 | 0,9916666 | 0,221456 | Q16537;B5 | PPP2R5E |
| 0,0843006 | -3,44235 | -2,03264 | -2,03264 | 1,7751049 | 0,221641 | P38405;K7E | GNAL |
| -1,74292 | -0,141456 | -0,613967 | -0,613967 | 0,8228496 | 0,221704 | O75396;A0. | SEC22B |
| -2,43517 | -0,681806 | -0,312629 | -0,681806 | 1,1340016 | 0,222908 | Q8N1G4;J3 | LRRC47 |
| -2,14368 | -0,451411 | -0,398282 | -0,451411 | 0,9927245 | 0,223829 | O14686;H0 | KMT2D |
| -0,146868 | -1,12306 | -2,8491 | -1,12306 | 1,3683459 | 0,224351 | Q8N9K5 | ZNF565 |
| 0,189161 | -2,22585 | -3,1445 | -2,22585 | 1,7218929 | 0,224476 | P61201;B4I | COPS2 |
| 1,86399 | 0,314163 | 0,423832 | 0,423832 | 0,8648744 | 0,224526 | Q14676;A2 | MDC1 |
| 0,0497953 | 0,185138 | 0,560232 | 0,185138 | 0,2644361 | 0,22468 | A0MZ66 | SHTN1 |
| 1,38331 | 2,54822 | -0,0400498 | 1,38331 | 1,2962837 | 0,225194 | E7ESC5;F5G | CCDC74B |
| 1,40108 | -0,137036 | 1,76234 | 1,40108 | 1,0086244 | 0,225351 | P21246;C9J | PTN |
| 0,181873 | -1,78393 | -1,89183 | -1,78393 | 1,1673523 | 0,226128 | Q96FZ7;I3L | CHMP6 |
| -2,80564 | 0,275754 | -2,73145 | -2,73145 | 1,7580183 | 0,226152 | P07384;E9F | CAPN1 |
| 4,64317 | 3,11307 | -0,259689 | 3,11307 | 2,5084768 | 0,226596 | P35659;B4I | DEK |
| 0,152394 | 1,44106 | 0,429085 | 0,429085 | 0,6783941 | 0,227339 | O14514;A0. | ADGRB1 |
| 0,143986 | 4,43561 | 1,86011 | 1,86011 | 2,1601049 | 0,227356 | Q9UBK8;H0 | MTRR |
| -2,3016 | -0,597387 | -0,309879 | -0,597387 | 1,0765652 | 0,227413 | A0A087WS | TXNRD1 |
| 0,302416 | 3,46997 | 1,1252 | 1,1252 | 1,6435895 | 0,227502 | P63208;E7E | SKP1 |
| 1,62874 | 0,669408 | 0,0591401 | 0,669408 | 0,7912426 | 0,227562 | Q9UPM6;H | LHX6 |
| 1,8829 | 0,0935618 | 0,724424 | 0,724424 | 0,9075411 | 0,227895 | Q92599;A0. | SEPT8 |
| 1,39988 | 0,909871 | -0,0748542 | 0,909871 | 0,7510696 | 0,22794 | P15170;H3I | GSPT1 |
| -0,0583204 | 0,577229 | 0,55009 | 0,55009 | 0,3593565 | 0,22803 | P52888;K7E | THOP1 |
| 3,32849 | 0,748854 | 0,540806 | 0,748854 | 1,5528999 | 0,228122 | P35637;H3I | FUS |
| 4,20824 | 2,99661 | -0,290959 | 2,99661 | 2,3280517 | 0,228549 | Q13029 | PRDM2 |
| 0,0403334 | -1,35617 | -0,759525 | -0,759525 | 0,7007116 | 0,229395 | O94973;A0. | AP2A2 |
| 1,6779 | -0,0852178 | 1,0549 | 1,0549 | 0,8941087 | 0,229463 | O14773;A0. | TPP1 |
| 2,64181 | 2,06264 | -0,223001 | 2,06264 | 1,5147461 | 0,229738 | P67936;A0/ | TPM4 |
| 1,94654 | 6,88684 | 0,764319 | 1,94654 | 3,2478063 | 0,2301 | O43347 | MSI1 |
| 0,136104 | -1,32512 | -1,83823 | -1,32512 | 1,0244065 | 0,230102 | P62888;E5F | RPL30 |
| 1,56789 | 4,76626 | 0,372518 | 1,56789 | 2,27169 | 0,230403 | Q7Z4W1;J3 | DCXR |
| 0,114273 | -1,05627 | -1,03471 | -1,03471 | 0,6696763 | 0,230461 | P12956;B1/ | XRCC6 |
| 4,44859 | 1,05226 | 0,651052 | 1,05226 | 2,0863573 | 0,230791 | P51843;A6I | NR0B1 |
| 0,0488334 | -1,70182 | -0,92792 | -0,92792 | 0,8772833 | 0,231507 | P09960;B4I | LTA4H |
| 2,49827 | 0,0317434 | 1,10779 | 1,10779 | 1,2365991 | 0,231522 | Q14687;H0 | GSE1 |
| -0,254943 | 2,2517 | 2,87079 | 2,2517 | 1,6551303 | 0,231619 | Q8N668;H0 | COMMD1 |
| -2,88873 | -3,43176 | 0,336495 | -2,88873 | 2,03702 | 0,231968 | M0QXH3 | CPAMD8 |
| -0,61495 | -2,15687 | -0,226534 | -0,61495 | 1,0209947 | 0,232067 | O00571;A0. | DDX3X |
| -1,90479 | -1,47276 | 0,168254 | -1,47276 | 1,0937008 | 0,23232 | Q8NEV1 | CSNK2A3 |
| -1,49471 | -0,675524 | -0,0102582 | -0,675524 | 0,7435547 | 0,232515 | P62701 | RPS4X |
| -0,127864 | -0,578467 | -0,0902931 | -0,127864 | 0,2716519 | 0,232516 | P49755;G3 | TMED10 |
| 0,052842 | -0,997509 | -0,609457 | -0,609457 | 0,5311091 | 0,233198 | O60333;A0. | KIF1B |
| -0,90008 | 0,0401837 | -0,522688 | -0,522688 | 0,4731711 | 0,233656 | P46777;A0/ | RPL5 |
| 0,101558 | -1,03801 | -0,843725 | -0,843725 | 0,609634 | 0,23386 | P0DMV8;A( | HSPA1A |
| 0,0599613 | 1,9127 | 0,757363 | 0,757363 | 0,935754 | 0,234143 | Q6P158;H7 | DHX57 |
| 0,269022 | 0,193613 | 1,24364 | 0,269022 | 0,5856796 | 0,234593 | P51991;H7( | HNRNPA3 |
| 0,122253 | 3,26533 | 1,25096 | 1,25096 | 1,5921997 | 0,234596 | J3QLE5;P14 | SNRPN |
| 0,161035 | 0,542742 | 1,81399 | 0,542742 | 0,8654509 | 0,235042 | P42345;B1/ | MTOR |
| -0,295869 | -0,094251 | -1,0108 | -0,295869 | 0,4816358 | 0,2351 | Q9Y617 | PSAT1 |
| -2,47088 | -2,74843 | 0,306463 | -2,47088 | 1,6893312 | 0,235155 | Q99623;J3K | PHB2 |
| -2,1552 | 0,0410801 | -1,07919 | -1,07919 | 1,0982144 | 0,235203 | P53992;G5I | SEC24C |
| 0,0903724 | -0,798543 | -0,727255 | -0,727255 | 0,4939243 | 0,235373 | Q3L8U1;H3 | CHD9 |

| | | | | | | | |
|---|---|---|---|---|---|---|---|
| 3,97374 | -0,123497 | 2,10883 | 2,10883 | 2,0513623 | 0,235508 | Q9NSD4;A6 | ZNF275 |
| -0,472653 | -0,167937 | -1,67976 | -0,472653 | 0,7995382 | 0,235817 | P28074;H0Y | PSMB5 |
| 0,204464 | 4,35808 | 1,57474 | 1,57474 | 2,1164894 | 0,236076 | P07099 | EPHX1 |
| 0,0087494 | -1,62969 | -0,756208 | -0,756208 | 0,8198185 | 0,236092 | P39748;I3L | FEN1 |
| 0,333156 | -2,83503 | -2,64536 | -2,64536 | 1,7769325 | 0,236411 | Q5T1H1 | EYS |
| 0,375633 | -3,44696 | -2,94677 | -2,94677 | 2,0776898 | 0,236429 | P55286;J3K | CDH8 |
| 0,4381 | 0,84672 | -0,0236578 | 0,4381 | 0,4354592 | 0,236469 | Q52M93;F2 | ZNF585B |
| -2,68879 | 0,311029 | -3,90164 | -2,68879 | 2,1685825 | 0,236529 | Q2M2I8 | AAK1 |
| -2,88567 | 0,0884435 | -1,50901 | -1,50901 | 1,4884221 | 0,2368 | P54652 | HSPA2 |
| -0,106869 | 2,04057 | 3,99362 | 2,04057 | 2,0510123 | 0,237154 | P18505;D6I | GABRB1 |
| -2,71749 | -0,671982 | -0,333791 | -0,671982 | 1,2897349 | 0,237497 | Q9H078;A0 | CLPB |
| 0,0789746 | 0,361193 | 1,12975 | 0,361193 | 0,5438223 | 0,2375 | Q09028;H0 | RBBP4 |
| 1,34685 | 2,78247 | -0,0483863 | 1,34685 | 1,4154762 | 0,237913 | Q9H4M9;A | EHD1 |
| 2,70644 | 2,66458 | -0,340033 | 2,66458 | 1,7469235 | 0,238265 | F8W914 | RTN4 |
| 1,93894 | 0,0448011 | 0,764902 | 0,764902 | 0,9560921 | 0,238825 | P49750;H0Y | YLPM1 |
| 1,21145 | 0,411348 | 0,0658206 | 0,411348 | 0,5876534 | 0,238978 | Q96PU8;F5 | QKI |
| 2,72668 | 3,61401 | -0,35667 | 2,72668 | 2,0840946 | 0,239226 | C9J4M0;Q9 | TBCCD1 |
| 4,49496 | 0,847824 | 0,768945 | 0,847824 | 2,1288107 | 0,239262 | A0A2R8YD5 | CSNK2A1 |
| -0,134956 | 1,03177 | 1,06296 | 1,03177 | 0,6827915 | 0,239342 | Q9H2K8;G3 | TAOK3 |
| -1,49601 | -1,24427 | 0,167378 | -1,24427 | 0,8965659 | 0,239399 | P52597;A0A | HNRNPF |
| 0,288343 | 0,329342 | 1,72144 | 0,329342 | 0,8158211 | 0,239681 | Q9NQ89;F5 | C12orf4 |
| -0,887493 | 0,118122 | -0,938309 | -0,887493 | 0,5958034 | 0,239796 | Q14980;A0 | NUMA1 |
| -1,98781 | -1,57962 | 0,214439 | -1,57962 | 1,1715494 | 0,240263 | G3V2S6;Q9 | ATP6V1D |
| 0,0625961 | -2,84061 | -1,36855 | -1,36855 | 1,4516511 | 0,240889 | O75165 | DNAJC13 |
| -0,69932 | 0,0834926 | -1,08758 | -0,69932 | 0,596511 | 0,240982 | Q9Y2K5;A0 | R3HDM2 |
| 0,0928658 | 3,39394 | 1,28549 | 1,28549 | 1,6715763 | 0,241056 | Q9GZM8;A | NDEL1 |
| -0,03585 | 1,04923 | 2,26235 | 1,04923 | 1,1496943 | 0,241705 | F5H2A4;F5I | HMGA2 |
| -0,0881532 | 1,61328 | 0,905961 | 0,905961 | 0,8547356 | 0,242267 | Q9UF12;K7 | PRODH2 |
| -0,0728007 | 1,04373 | 0,633395 | 0,633395 | 0,5647607 | 0,242668 | Q14257;H0 | RCN2 |
| 0,339846 | -2,37964 | -2,77796 | -2,37964 | 1,6968096 | 0,24283 | Q9NS86 | LANCL2 |
| -2,52739 | -0,774934 | -0,171437 | -0,774934 | 1,22378 | 0,242915 | P06576;H0Y | ATP5F1B |
| -1,50057 | 0,0854748 | -3,08806 | -1,50057 | 1,5867675 | 0,242984 | Q9BRZ2;C9 | TRIM56 |
| -1,05674 | -1,83183 | 0,112258 | -1,05674 | 0,9786725 | 0,243113 | Q86US8;I3L | SMG6 |
| 2,13493 | 3,41733 | -0,260206 | 2,13493 | 1,8666145 | 0,243307 | Q99698 | LYST |
| 0,0312166 | -0,276061 | -0,466139 | -0,276061 | 0,2509687 | 0,243553 | P62714;E5F | PPP2CB |
| 0,0312185 | -0,276048 | -0,466151 | -0,276048 | 0,2509742 | 0,243561 | B3KUN1;P6 | PPP2CA |
| -0,186023 | 1,47617 | 1,27779 | 1,27779 | 0,9078352 | 0,244047 | Q8NBS9 | TXNDC5 |
| 0,286137 | -1,97761 | -2,1788 | -1,97761 | 1,368755 | 0,244167 | P31930 | UQCRC1 |
| -0,788998 | -2,77599 | -0,223786 | -0,788998 | 1,3404817 | 0,2443 | O00154;K7I | ACOT7 |
| -1,861 | -0,377961 | -0,271464 | -0,377961 | 0,8885729 | 0,244438 | P05091;S4F | ALDH2 |
| 2,57029 | -0,273372 | 1,90548 | 1,90548 | 1,4874913 | 0,244446 | P62273;A0A | RPS29 |
| 0,256132 | -1,90135 | -1,77066 | -1,77066 | 1,2096621 | 0,244598 | Q9Y4B5;J3 | MTCL1 |
| -0,0421832 | -0,165002 | -0,56743 | -0,165002 | 0,2747474 | 0,245111 | O60264 | SMARCA5 |
| 3,60781 | -0,114293 | 1,74609 | 1,74609 | 1,8610515 | 0,245569 | Q2M243;J3 | CCDC27 |
| -1,452 | -0,0922464 | -0,441635 | -0,441635 | 0,7061444 | 0,245929 | Q9HC38;I3I | GLOD4 |
| -0,112995 | 0,797061 | 0,774423 | 0,774423 | 0,5190095 | 0,246178 | B4E2Q0;H0 | ATP2C1 |
| 1,54228 | 1,69329 | -0,230579 | 1,54228 | 1,0698212 | 0,246324 | O75475 | PSIP1 |
| -0,667489 | -0,73489 | 0,100272 | -0,667489 | 0,4639496 | 0,246591 | O00311;B1 | CDC7 |
| -1,65809 | -3,00193 | 0,182158 | -1,65809 | 1,5984803 | 0,247201 | P13798;C9J | APEH |
| 0,17393 | 1,18354 | 0,229114 | 0,229114 | 0,5676394 | 0,247933 | P49903;Q5 | SEPHS1 |
| 4,57752 | 0,00762012 | 1,84061 | 1,84061 | 2,2998012 | 0,248051 | Q96CS3 | FAF2 |
| -0,151031 | 1,95055 | 1,16716 | 1,16716 | 1,0620711 | 0,248138 | Q8TE73 | DNAH5 |

| | | | | | | | |
|---|---|---|---|---|---|---|---|
| 1,63479 | 1,43233 | -0,222735 | 1,43233 | 1,0190379 | 0,248378 | Q70Z35 | PREX2 |
| 0,434612 | -2,85238 | -2,95454 | -2,85238 | 1,9279136 | 0,248924 | A0A024R4E | HDLBP |
| -1,3832 | 0,190313 | -2,24704 | -1,3832 | 1,2357759 | 0,249274 | F8W6K2 | CHN1 |
| 0,104589 | 1,3957 | 0,386118 | 0,386118 | 0,6789062 | 0,249866 | P67809;H0Y | YBX1 |
| -0,876346 | -0,137422 | -2,79385 | -0,876346 | 1,3710969 | 0,250047 | P15586;F6S | GNS |
| -0,975332 | -0,694183 | 0,110053 | -0,694183 | 0,5633088 | 0,251074 | P43243;A0A | MATR3 |
| 0,408634 | -2,9599 | -2,4856 | -2,4856 | 1,823393 | 0,251792 | P00338;F5C | LDHA |
| -0,0515624 | 2,35615 | 1,0259 | 1,0259 | 1,2060659 | 0,251901 | Q9ULJ3;E7E | ZBTB21 |
| 2,23085 | 2,32458 | -0,356445 | 2,23085 | 1,5215549 | 0,252113 | G5EA30;Q9 | CELF1 |
| -1,1306 | 0,185757 | -1,58765 | -1,1306 | 0,9207481 | 0,253215 | Q5T123;Q9 | SH3BGRL3 |
| -1,78185 | 0,276006 | -1,69529 | -1,69529 | 1,163921 | 0,253235 | P41252;A0A | IARS |
| -2,83692 | -1,65557 | 0,234972 | -1,65557 | 1,5495299 | 0,253558 | Q96Q35 | ALS2CR12 |
| 0,0593964 | 3,53661 | 1,25609 | 1,25609 | 1,7665343 | 0,253671 | E9PBD5;G3 | ADAMTS20 |
| 0,660277 | 3,34155 | 0,437388 | 0,660277 | 1,616223 | 0,253673 | A0A1W2PC | RPS10-NUDT3 |
| 0,31479 | -2,67301 | -1,89261 | -1,89261 | 1,5496552 | 0,254104 | J3KSW6;Q8 | DHX40 |
| 0,0781254 | -0,679222 | -1,32441 | -0,679222 | 0,7020147 | 0,254135 | P35241;A0A | RDX |
| 1,62244 | 5,85501 | 0,386237 | 1,62244 | 2,8679355 | 0,254242 | Q71F56;H0 | MED13L |
| 0,0422807 | -1,08047 | -2,62553 | -1,08047 | 1,3394647 | 0,255056 | Q96NB2;AC | SFXN2 |
| 0,24625 | 2,87828 | 0,714708 | 0,714708 | 1,4040465 | 0,255153 | Q9P0I2;S4R | EMC3 |
| -1,83835 | -0,219135 | -0,379812 | -0,379812 | 0,8920956 | 0,255433 | P04080;A0A | CSTB |
| -2,84791 | 0,443901 | -2,59625 | -2,59625 | 1,8322059 | 0,255799 | O14578;H0 | CIT |
| -1,74263 | 0,289315 | -2,72455 | -1,74263 | 1,5371158 | 0,257154 | G3XAL9;P5 | SLC12A2 |
| -0,23958 | -0,636135 | 0,00114048 | -0,23958 | 0,3217976 | 0,25718 | Q14344 | GNA13 |
| 1,68649 | 1,31345 | -0,234156 | 1,31345 | 1,018424 | 0,257429 | Q86XP3;A0 | DDX42 |
| -0,0615863 | -0,971123 | -2,90385 | -0,971123 | 1,4515023 | 0,257882 | Q8IUR6;E5I | CREBRF |
| 0,999864 | 1,50782 | -0,171875 | 0,999864 | 0,8614297 | 0,257945 | Q9NQ48;H7 | LZTFL1 |
| -0,507223 | 2,84633 | 3,29508 | 2,84633 | 2,0778676 | 0,257948 | A0AVT1;H0 | UBA6 |
| -1,79106 | 0,195626 | -1,15313 | -1,15313 | 1,0143158 | 0,258164 | P62249;M0 | RPS16 |
| -1,85863 | -2,80856 | 0,32156 | -1,85863 | 1,6048492 | 0,258405 | Q8N465;B5 | D2HGDH |
| -0,226526 | 1,3513 | 2,2238 | 1,3513 | 1,2419668 | 0,259843 | O43396;K7I | TXNL1 |
| 3,39679 | 0,0894412 | 1,08663 | 1,08663 | 1,6965544 | 0,259942 | E9PC15;Q5 | AGK |
| -3,02345 | -2,95403 | 0,522113 | -2,95403 | 2,0272891 | 0,260492 | Q69YN4 | VIRMA |
| 0,265029 | -1,59308 | -2,69945 | -1,59308 | 1,4980409 | 0,260798 | A0A0A0MS | EPB41L3 |
| 0,241283 | 0,218362 | -0,0400541 | 0,218362 | 0,1562342 | 0,261155 | P00505 | GOT2 |
| 2,6605 | 1,53311 | -0,251879 | 1,53311 | 1,4685109 | 0,26134 | P31350;A0A | RRM2 |
| 1,09703 | 3,10036 | 0,00945401 | 1,09703 | 1,5678995 | 0,261472 | Q9NR99;G3 | MXRA5 |
| -0,0442498 | 0,671221 | 0,333334 | 0,333334 | 0,3579189 | 0,261483 | P49006 | MARCKSL1 |
| 0,451516 | -2,63384 | -2,48052 | -2,48052 | 1,7387622 | 0,261649 | Q9BX84 | TRPM6 |
| 0,0320583 | -0,465947 | -1,14024 | -0,465947 | 0,5883541 | 0,262429 | P46776;E9F | RPL27A |
| 0,524293 | -2,76494 | -3,24796 | -2,76494 | 2,0527321 | 0,26264 | Q7Z7L1;K7E | SLFN11 |
| -0,197378 | 1,26367 | 1,03497 | 1,03497 | 0,7858802 | 0,262645 | P33991;E5F | MCM4 |
| -0,57413 | 3,11121 | 3,26096 | 3,11121 | 2,172252 | 0,263228 | P42224;J3K | STAT1 |
| -1,80026 | 0,311115 | -1,6615 | -1,6615 | 1,1809861 | 0,263396 | P14324;A0A | FDPS |
| -0,952872 | 0,134543 | -0,699576 | -0,699576 | 0,5689735 | 0,263398 | Q9H0C2 | SLC25A31 |
| -1,96205 | 0,329788 | -1,73758 | -1,73758 | 1,2633895 | 0,26346 | A0A0B4J22 | ZSCAN9 |
| -1,73986 | -0,421745 | -0,127479 | -0,421745 | 0,858661 | 0,26364 | P55786;E9F | NPEPPS |
| -0,787208 | -0,379133 | -3,77379 | -0,787208 | 1,8533705 | 0,26369 | Q9C0D4;D6 | ZNF518B |
| -0,173959 | 1,43401 | 0,932151 | 0,932151 | 0,8226893 | 0,263792 | E7EW50;J3I | NCOR1 |
| 0,701664 | 0,0298528 | 0,19824 | 0,19824 | 0,3495522 | 0,264403 | A0A087WT | ZNF732 |
| -3,04773 | -2,1239 | 0,410907 | -2,1239 | 1,7907577 | 0,264576 | Q8NI27;A0A | THOC2 |
| -0,16721 | -0,142852 | -1,02416 | -0,16721 | 0,5019396 | 0,264622 | Q9UEU0;H0 | VTI1B |
| 1,13294 | 0,970354 | -0,188747 | 0,970354 | 0,720741 | 0,264847 | P14927;B7Z | UQCRB |

| | | | | | | | |
|---|---|---|---|---|---|---|---|
| -0,0744613 | -1,38796 | -0,368103 | -0,368103 | 0,6893974 | 0,264988 | P32969;A0/ | RPL9 |
| -0,4307 | -0,0621582 | -0,0677923 | -0,067792 | 0,2111701 | 0,265024 | Q9UDY2;A0 | TJP2 |
| 0,355003 | -1,85351 | -2,89017 | -1,85351 | 1,6574751 | 0,265937 | A0A1W2PR | IQSEC2 |
| 0,309153 | 1,18553 | 0,0650452 | 0,309153 | 0,5892243 | 0,26603 | P78563;A0/ | ADARB1 |
| 3,08128 | 0,142828 | 0,839844 | 0,839844 | 1,5353809 | 0,266057 | P19174;A0/ | PLCG1 |
| 1,12496 | 0,151577 | 0,185365 | 0,185365 | 0,5524875 | 0,266165 | O95202 | LETM1 |
| -2,41012 | -0,36624 | -0,353486 | -0,36624 | 1,1837336 | 0,266419 | P36405 | ARL3 |
| 3,09961 | 1,20132 | -0,0810312 | 1,20132 | 1,6002296 | 0,267316 | Q86V15 | CASZ1 |
| 0,112267 | 0,145883 | 0,86919 | 0,145883 | 0,427636 | 0,267425 | P12081;B3I | HARS |
| -1,4971 | 0,00350377 | -0,514423 | -0,514423 | 0,7622023 | 0,267646 | Q9H2U2;D6 | PPA2 |
| -2,23923 | -2,10279 | 0,412133 | -2,10279 | 1,4929378 | 0,267927 | P21796;C9J | VDAC1 |
| 0,141392 | 0,424054 | 0,00048685 | 0,141392 | 0,2157009 | 0,269043 | P78310 | CXADR |
| -1,56086 | 0,299402 | -2,65623 | -1,56086 | 1,4942206 | 0,269277 | Q6ZNG0 | ZNF620 |
| -1,81448 | 0,183767 | -1,00426 | -1,00426 | 1,0050585 | 0,269301 | Q07020;G3 | RPL18 |
| 0,000394145 | -1,30869 | -0,436599 | -0,436599 | 0,6664842 | 0,269775 | Q15907;H3 | RAB11B |
| 4,51395 | 4,85125 | -0,908143 | 4,51395 | 3,2322199 | 0,26998 | Q9NWQ4;G | GPATCH2L |
| -2,18566 | 0,0204625 | -0,758931 | -0,758931 | 1,1187781 | 0,270347 | P21266;A0/ | GSTM3 |
| -1,05577 | -2,44902 | 0,130522 | -1,05577 | 1,291154 | 0,270386 | Q86UV5 | USP48 |
| 4,79519 | 0,837879 | 0,556354 | 0,837879 | 2,3702074 | 0,270652 | P09417;B7Z | QDPR |
| -0,170402 | -1,1433 | -0,160556 | -0,170402 | 0,5645667 | 0,270656 | Q13363;D6 | CTBP1 |
| -0,00515767 | 3,29784 | 1,09683 | 1,09683 | 1,6816963 | 0,270805 | Q13423;D6 | NNT |
| 0,196969 | -1,68792 | -0,997938 | -0,997938 | 0,9536495 | 0,270846 | O00443;A0 | PIK3C2A |
| -2,44495 | -0,995019 | 0,106792 | -0,995019 | 1,2798226 | 0,271561 | O95271;E7I | TNKS |
| 0,296196 | -2,03924 | -1,3957 | -1,3957 | 1,2062972 | 0,271881 | P26639 | TARS |
| 0,941632 | 0,26511 | 0,0255281 | 0,26511 | 0,4751014 | 0,272971 | Q8IV36 | HID1 |
| -1,09243 | -0,0529438 | -0,273449 | -0,273449 | 0,5477049 | 0,273395 | P12111;C9J | COL6A3 |
| -0,902297 | 0,0156822 | -0,316173 | -0,316173 | 0,4648217 | 0,273774 | Q9HDC9;H0 | APMAP |
| -2,21333 | -2,09466 | 0,436151 | -2,09466 | 1,4965982 | 0,273846 | O15294 | OGT |
| -0,0770209 | 2,14204 | 0,821242 | 0,821242 | 1,1162149 | 0,274023 | P20338 | RAB4A |
| 0,167637 | 1,65816 | 0,304004 | 0,304004 | 0,8240139 | 0,274168 | Q6TFL3 | CCDC171 |
| 1,95766 | -0,325687 | 4,17084 | 1,95766 | 2,2483547 | 0,274667 | Q8IWG1 | WDR63 |
| 0,024846 | -0,336481 | -0,92487 | -0,336481 | 0,4793606 | 0,274857 | Q01780;K7I | EXOSC10 |
| 0,425855 | -0,0946388 | 0,588878 | 0,425855 | 0,356998 | 0,275147 | Q8IYM0;A0 | FAM186B |
| 0,587923 | -2,89592 | -2,81806 | -2,81806 | 1,9893024 | 0,275214 | Q9UKE5;C9 | TNIK |
| -0,389864 | -0,0857923 | -1,6291 | -0,389864 | 0,8175129 | 0,275511 | Q9P2D8 | UNC79 |
| 0,66339 | 0,245132 | -0,0217107 | 0,245132 | 0,3453278 | 0,27639 | Q9NSK0;H7 | KLC4 |
| -0,560311 | 0,116658 | -0,996632 | -0,560311 | 0,5609631 | 0,276457 | P12931;J3C | SRC |
| -0,533273 | 0,119799 | -0,663303 | -0,533273 | 0,4196545 | 0,27668 | O60292 | SIPA1L3 |
| -1,34001 | -2,53182 | 0,26817 | -1,34001 | 1,4051452 | 0,276848 | Q9Y6E2;E7I | BZW2 |
| 2,44102 | 1,53051 | -0,340137 | 1,53051 | 1,4179318 | 0,277328 | P52294;C9J | KPNA1 |
| -0,405829 | 0,0828967 | -0,755465 | -0,405829 | 0,4210995 | 0,277348 | Q13164;C9 | MAPK7 |
| 0,646714 | -2,8355 | -3,82813 | -2,8355 | 2,3500127 | 0,277419 | H3BR42;H3 | BLOC1S6 |
| 0,343878 | -2,44006 | -1,53985 | -1,53985 | 1,420629 | 0,277544 | O60268;H3 | KIAA0513 |
| 2,2589 | 0,431986 | 0,197056 | 0,431986 | 1,1287166 | 0,277658 | Q8N7H5;M | PAF1 |
| -1,37501 | 0,253442 | -1,11841 | -1,11841 | 0,8755643 | 0,277696 | O75347;E5I | TBCA |
| 2,88584 | -0,260383 | 1,39693 | 1,39693 | 1,5738625 | 0,278046 | Q8NEN0 | ARMC2 |
| 3,6446 | 1,98222 | -0,416478 | 1,98222 | 2,0416339 | 0,278546 | P35908 | KRT2 |
| 0,00739664 | -1,68412 | -0,522804 | -0,522804 | 0,8651586 | 0,279865 | Q9H1B7 | IRF2BPL |
| 0,287097 | -0,00851142 | 0,901774 | 0,287097 | 0,4643691 | 0,279931 | P22626;A0/ | HNRNPA2B1 |
| 0,206332 | 2,1178 | 0,368473 | 0,368473 | 1,0598856 | 0,280121 | Q8N944;C9 | AMER3 |
| 0,816536 | 0,126427 | 3,31298 | 0,816536 | 1,6764349 | 0,280364 | P63151;Q6 | PPP2R2A |
| -0,387099 | -1,31036 | -0,004972 | -0,387099 | 0,6711271 | 0,280639 | P01116;G3 | KRAS |

| | | | | | | | |
|---|---|---|---|---|---|---|---|
| -1,49971 | -0,940519 | 0,218008 | -0,940519 | 0,8761121 | 0,280669 | Q92614;A0 | MYO18A |
| -0,194819 | 0,85522 | 1,46149 | 0,85522 | 0,838004 | 0,281269 | P53597 | SUCLG1 |
| -0,539215 | 2,36305 | 4,17996 | 2,36305 | 2,3802982 | 0,282617 | P42025 | ACTR1B |
| 1,87142 | 0,396212 | 0,111738 | 0,396212 | 0,9446026 | 0,283083 | Q92623 | TTC9 |
| -0,0348169 | -0,480704 | -1,81429 | -0,480704 | 0,9259042 | 0,283451 | P04899 | GNAI2 |
| -0,528953 | -1,14341 | 0,103845 | -0,528953 | 0,62365 | 0,283616 | P09874;Q5Y | PARP1 |
| -0,527315 | 0,125536 | -0,619921 | -0,527315 | 0,4063038 | 0,283677 | Q9NTZ6 | RBM12 |
| -0,277928 | 1,33479 | 1,18155 | 1,18155 | 0,8901702 | 0,283682 | A0A087X08 | RYR3 |
| 5,25567 | 0,0858188 | 1,4 | 1,4 | 2,6870251 | 0,284471 | O75663 | TIPRL |
| 0,110314 | -0,870388 | -2,3804 | -0,870388 | 1,2546958 | 0,285298 | P07951;A7X | TPM2 |
| 2,47046 | -0,0409293 | 0,775415 | 0,775415 | 1,2810589 | 0,285464 | H3BMM9;H | RNPS1 |
| 0,976237 | -0,241596 | 1,2765 | 0,976237 | 0,8039372 | 0,285489 | M0QZR4;Q | ARHGEF1 |
| -1,21762 | -0,496712 | 0,0837202 | -0,496712 | 0,6519325 | 0,285547 | Q8N163;H0 | CCAR2 |
| -0,671374 | 3,93187 | 2,66192 | 2,66192 | 2,3774453 | 0,286965 | Q9UJX5;D6 | ANAPC4 |
| -0,725312 | -0,0859752 | -0,098988 | -0,098988 | 0,3654227 | 0,286974 | P18669 | PGAM1 |
| -0,0207265 | -1,62747 | -0,430746 | -0,430746 | 0,8348542 | 0,287092 | P31948;F5H | STIP1 |
| 0,188242 | -0,0474817 | 0,249287 | 0,188242 | 0,1567183 | 0,287281 | P62917;E9F | RPL8 |
| 0,429869 | 0,113259 | 2,07453 | 0,429869 | 1,0529115 | 0,287666 | C9JVB6;Q7I | TRMT10C |
| 1,90708 | -0,485275 | 2,71987 | 1,90708 | 1,6661805 | 0,28772 | F8W6N3;Q | BAP1 |
| 0,273186 | 0,122878 | 1,54444 | 0,273186 | 0,7809735 | 0,287862 | Q06323;H0 | PSME1 |
| 2,79318 | -0,139749 | 1,00128 | 1,00128 | 1,4784522 | 0,289668 | P04275 | VWF |
| 0,356594 | 0,845521 | -0,0702733 | 0,356594 | 0,4582475 | 0,28996 | P62753;A2A | RPS6 |
| 0,172655 | 2,3205 | 0,412849 | 0,412849 | 1,1768647 | 0,290054 | Q13308 | PTK7 |
| 0,54635 | -2,10835 | -3,36272 | -2,10835 | 1,9959001 | 0,290319 | O95450 | ADAMTS2 |
| -0,351811 | -0,0154094 | -1,36291 | -0,351811 | 0,7013374 | 0,290393 | O15068;A2 | MCF2L |
| -0,433662 | -0,273059 | -2,85096 | -0,433662 | 1,4442239 | 0,290896 | Q9UI46;A0A | DNAI1 |
| 0,653255 | -0,171701 | 0,962038 | 0,653255 | 0,5861262 | 0,29096 | O15083;H7 | ERC2 |
| -0,308575 | 1,68632 | 1,17273 | 1,17273 | 1,0358286 | 0,291058 | P23142;B1A | FBLN1 |
| 0,310312 | 0,171634 | 1,9449 | 0,310312 | 0,9862033 | 0,291267 | Q6ULP2 | AFTPH |
| 4,54348 | 2,57692 | -0,662721 | 2,57692 | 2,6289149 | 0,291897 | Q9HC62 | SENP2 |
| 0,384256 | 0,0239051 | 1,54715 | 0,384256 | 0,796079 | 0,29193 | Q96NB3;J3 | ZNF830 |
| -2,48362 | -0,173728 | -0,443942 | -0,443942 | 1,2628607 | 0,291989 | Q6ZU15 | SEPT14 |
| 0,786267 | -0,190601 | 0,795285 | 0,786267 | 0,5666162 | 0,292123 | P35611;E7E | ADD1 |
| 0,079426 | -0,297076 | -0,413143 | -0,297076 | 0,2575039 | 0,292871 | P61254 | RPL26 |
| -0,40121 | 0,0864445 | -0,92842 | -0,40121 | 0,5075607 | 0,292916 | O15173 | PGRMC2 |
| -0,0814369 | 2,18596 | 0,71222 | 0,71222 | 1,1505716 | 0,29309 | Q8NE71;H0 | ABCF1 |
| 0,997367 | 5,56775 | 0,374661 | 0,997367 | 2,8356168 | 0,293201 | B0QY60 | SUN2 |
| -1,3358 | -2,94707 | 0,304351 | -1,3358 | 1,6257319 | 0,293222 | Q9P2D1 | CHD7 |
| -0,429915 | 1,62843 | 2,79308 | 1,62843 | 1,6320177 | 0,293426 | C9JPR7;Q1 | CLEC5A |
| -2,0523 | -0,0204681 | -0,517508 | -0,517508 | 1,0591645 | 0,293458 | P05026;V9 | ATP1B1 |
| -0,92847 | -2,65144 | 0,144841 | -0,92847 | 1,4106623 | 0,294983 | P18858;F5 | LIG1 |
| 0,154604 | -0,835923 | -2,22275 | -0,835923 | 1,1941695 | 0,29545 | Q00169;F5 | PITPNA |
| 0,579471 | 0,390476 | -0,107312 | 0,390476 | 0,3547729 | 0,295502 | P09497;H0 | CLTB |
| 0,363867 | -1,92195 | -1,31146 | -1,31146 | 1,1835243 | 0,296517 | Q9Y5K5;Q5 | UCHL5 |
| 0,554943 | -0,152517 | 0,928104 | 0,554943 | 0,548861 | 0,296577 | O75054 | IGSF3 |
| 0,808756 | 0,183042 | 0,0185858 | 0,183042 | 0,4169198 | 0,296682 | P41219;H7 | PRPH |
| 1,54544 | 0,474666 | -0,0503168 | 0,474666 | 0,8132859 | 0,296893 | Q7Z2Z1;H0 | TICRR |
| 0,22686 | -1,05913 | -0,823712 | -0,823712 | 0,6847011 | 0,297402 | A0A2U3TZH | EEF1A2 |
| 0,575497 | -2,08657 | -3,61719 | -2,08657 | 2,1216355 | 0,297611 | Q13342;U3 | SP140 |
| 1,47876 | -0,356216 | 3,32511 | 1,47876 | 1,8406659 | 0,297728 | A0A1B0GU | KIF1BP |
| -0,151033 | 1,04819 | 0,558276 | 0,558276 | 0,602947 | 0,298092 | K7EP73;O1 | GAPDHS |
| 0,205139 | -0,773596 | -1,54931 | -0,773596 | 0,8791801 | 0,29884 | J3KTH2;Q9 | TAF4B |

| | | | | | | | |
|---|---|---|---|---|---|---|---|
| 0,732403 | -0,203766 | 1,32744 | 0,732403 | 0,7719103 | 0,299474 | O43602;A8 | DCX |
| 0,797988 | 0,127271 | 3,87409 | 0,797988 | 1,9979538 | 0,299829 | B5MEG5;E7 | USP19 |
| -2,47798 | -2,33582 | 0,620358 | -2,33582 | 1,749233 | 0,300547 | Q12767;C9. | TMEM94 |
| -2,23414 | 0,33371 | -1,19613 | -1,19613 | 1,2917513 | 0,300566 | B7ZM87;O7 | SRGAP2 |
| -0,165161 | -0,365198 | 0,0425221 | -0,165161 | 0,203872 | 0,301211 | Q9P2B2 | PTGFRN |
| -0,319773 | -0,96446 | 0,0528762 | -0,319773 | 0,5146944 | 0,301279 | Q71F23;Q0 | CENPU |
| -2,34154 | 0,0301491 | -0,619062 | -0,619062 | 1,2256505 | 0,301498 | Q8N1I0;H0' | DOCK4 |
| -0,500394 | 1,72142 | 2,86109 | 1,72142 | 1,7095262 | 0,301958 | G3XAG1;Q9 | ZNF512 |
| -0,323781 | 1,1468 | 1,38858 | 1,1468 | 0,9267548 | 0,302179 | Q9NZN3 | EHD3 |
| -0,25211 | 3,26941 | 1,20387 | 1,20387 | 1,7695308 | 0,302317 | P53779;A0/ | MAPK10 |
| 0,373448 | 3,70009 | 0,451988 | 0,451988 | 1,8983713 | 0,302553 | K7EK91;Q9 | DNAH17 |
| -0,662425 | 4,40639 | 2,32541 | 2,32541 | 2,547892 | 0,302821 | Q5VT97 | SYDE2 |
| -2,30825 | -0,44034 | -0,0899887 | -0,44034 | 1,1925125 | 0,303089 | P51149;C9J | RAB7A |
| 0,841936 | -0,248956 | 1,21139 | 0,841936 | 0,7592927 | 0,303685 | O95995;H3 | GAS8 |
| -0,979697 | -0,677319 | 0,200625 | -0,677319 | 0,6131039 | 0,303828 | Q04637;E7I | EIF4G1 |
| 2,55625 | 0,129459 | 0,446993 | 0,446993 | 1,3190342 | 0,303894 | O15020;A4 | SPTBN2 |
| 0,0372015 | -0,211104 | -0,643471 | -0,211104 | 0,344459 | 0,304208 | Q5JPF3;E9F | ANKRD36C |
| 2,77092 | -0,582827 | 1,96551 | 1,96551 | 1,7507296 | 0,304272 | P62314;J3C | SNRPD1 |
| 5,17471 | -1,47636 | 6,0535 | 5,17471 | 4,1171956 | 0,304868 | C9JLV4;O14 | APAF1 |
| -0,563998 | 0,168532 | -0,954816 | -0,563998 | 0,5702704 | 0,304985 | Q9H0Q0;C9 | FAM49A |
| -0,358297 | 0,00451949 | -1,49078 | -0,358297 | 0,7799652 | 0,305421 | Q6ZU80;H0 | CEP128 |
| 0,4649 | 3,2734 | 0,248726 | 0,4649 | 1,6873577 | 0,305732 | H7C141 | OPA1 |
| 4,42102 | -0,848168 | 2,79165 | 2,79165 | 2,6977603 | 0,306295 | Q9UI17;E5F | DMGDH |
| -0,156414 | 0,0350642 | -0,436533 | -0,156414 | 0,2371829 | 0,307374 | Q6SZW1;J3 | SARM1 |
| 0,053623 | -0,213829 | -0,548369 | -0,213829 | 0,3016184 | 0,307816 | P36871 | PGM1 |
| 0,755926 | 8,08921 | 0,951302 | 0,951302 | 4,1786153 | 0,308557 | O00479 | HMGN4 |
| -0,0959958 | -0,50478 | -0,0140871 | -0,095996 | 0,2628666 | 0,309382 | Q14679;E7I | TTLL4 |
| -1,67184 | 0,500737 | -2,04037 | -1,67184 | 1,373143 | 0,309428 | Q502W6;F8 | VWA3B |
| 0,610391 | 5,69903 | 0,580689 | 0,610391 | 2,9465388 | 0,30949 | Q07866;B5 | KLC1 |
| -0,0352474 | -0,477094 | -2,32125 | -0,477094 | 1,2125694 | 0,309725 | O14513;A0. | NCKAP5 |
| 0,0901764 | -0,295567 | -0,380483 | -0,295567 | 0,2508416 | 0,309913 | P17483 | HOXB4 |
| 0,405784 | 0,0210635 | 1,93298 | 0,405784 | 1,0112502 | 0,310232 | A6NMH8;E9 | CD81 |
| 2,36987 | -0,651991 | 2,31971 | 2,31971 | 1,7303741 | 0,310265 | Q9P2E9;A0. | RRBP1 |
| -2,82321 | -0,783817 | 0,105095 | -0,783817 | 1,5013471 | 0,310395 | P30044 | PRDX5 |
| 0,508155 | -1,64505 | -3,13541 | -1,64505 | 1,8318038 | 0,310431 | Q8NA03;A0 | FSIP1 |
| -1,28732 | 0,352995 | -3,14665 | -1,28732 | 1,7509643 | 0,310677 | P57678;I3L | GEMIN4 |
| -1,98756 | 0,5053 | -1,68786 | -1,68786 | 1,361012 | 0,310905 | P0DME0;A0 | SETSIP |
| 1,23918 | 0,131535 | 0,123517 | 0,131535 | 0,6418263 | 0,311083 | Q9UIV1 | CNOT7 |
| -0,399551 | -1,12814 | 0,0951927 | -0,399551 | 0,6153801 | 0,311124 | Q9C0C9;K7 | UBE2O |
| -0,643117 | 2,13167 | 2,50213 | 2,13167 | 1,7189756 | 0,312103 | Q9Y490 | TLN1 |
| 0,163626 | -1,00797 | -0,521034 | -0,521034 | 0,5885722 | 0,312373 | P08559;Q5. | PDHA1 |
| 0,683892 | -3,02333 | -2,15554 | -2,15554 | 1,9390259 | 0,312631 | F5GWH7 | MDM2 |
| 0,449052 | -1,76166 | -1,4708 | -1,4708 | 1,2012271 | 0,312792 | P20042;B5E | EIF2S2 |
| 0,94618 | 1,611 | -0,30495 | 0,94618 | 0,9728118 | 0,313099 | P27708;F8\ | CAD |
| 0,266755 | -1,42103 | -0,825869 | -0,825869 | 0,856024 | 0,313411 | E7ESY4;H0Y | MTA1 |
| 1,63462 | -0,216374 | 0,722004 | 0,722004 | 0,9255269 | 0,313524 | Q8IV08;M0 | PLD3 |
| -0,535649 | 1,82322 | 1,93545 | 1,82322 | 1,3954204 | 0,313957 | Q01658 | DR1 |
| 0,455922 | 1,75121 | -0,0560348 | 0,455922 | 0,9314867 | 0,314019 | Q9Y2B0 | CNPY2 |
| -0,104424 | -0,377126 | 0,0165266 | -0,104424 | 0,2016423 | 0,314513 | P49756 | RBM25 |
| 1,87329 | 0,262477 | 0,112181 | 0,262477 | 0,9762865 | 0,315086 | O95626 | ANP32D |
| 0,417453 | -1,35908 | -1,54492 | -1,35908 | 1,0833216 | 0,316233 | Q9P2N2;E9 | ARHGAP28 |
| -2,53041 | 0,775524 | -2,85932 | -2,53041 | 2,0103678 | 0,316248 | O60493 | SNX3 |

| | | | | | | | |
|---|---|---|---|---|---|---|---|
| 0,141126 | 2,58359 | 0,36777 | 0,36777 | 1,3494972 | 0,316823 | Q9Y597 | KCTD3 |
| 1,05898 | 0,348667 | -0,0851835 | 0,348667 | 0,5776217 | 0,317155 | Q96T58;F6\ | SPEN |
| 3,28787 | 5,76999 | -1,10426 | 3,28787 | 3,4810688 | 0,317901 | A6NGH7 | CCDC160 |
| 0,402757 | 0,0067332 | 0,0743264 | 0,0743264 | 0,2118452 | 0,31806 | O75955;A0. | FLOT1 |
| -0,817979 | 4,53657 | 2,42815 | 2,42815 | 2,6973439 | 0,318861 | D6RDN9;Q: | PDK2 |
| 0,0171745 | -2,4302 | -0,517298 | -0,517298 | 1,2867593 | 0,319105 | Q9UPP1;H( | PHF8 |
| 1,89188 | 1,29023 | -0,433544 | 1,29023 | 1,2069919 | 0,319119 | Q9GZP4;X6 | PITHD1 |
| -0,819059 | 3,47818 | 2,43216 | 2,43216 | 2,240939 | 0,319965 | Q8N103 | TAGAP |
| -1,4259 | -0,19833 | -0,0726208 | -0,19833 | 0,7476737 | 0,320357 | P62826;B5I | RAN |
| 0,153532 | 1,28044 | 0,0872711 | 0,153532 | 0,6705674 | 0,320504 | P61266 | STX1B |
| -0,0167751 | 0,60107 | 2,89482 | 0,60107 | 1,5340799 | 0,320618 | O75038;B9 | PLCH2 |
| -0,80938 | 2,81741 | 2,61051 | 2,61051 | 2,0368301 | 0,320678 | I3L4C2;Q9L | BAIAP2 |
| -2,95695 | -1,35677 | 0,450184 | -1,35677 | 1,7046124 | 0,320838 | A0A0G2JQ6 | LPCAT1 |
| 0,621872 | -1,93797 | -2,26831 | -1,93797 | 1,5819328 | 0,320948 | Q5TA45;C9. | INTS11 |
| -0,141858 | -0,161985 | -1,64228 | -0,161985 | 0,8605177 | 0,321638 | P13804;H0' | ETFA |
| 0,119829 | -2,72274 | -0,700598 | -0,700598 | 1,4630082 | 0,322213 | Q8NDI1;B5 | EHBP1 |
| -0,994295 | 0,263662 | -0,796733 | -0,796733 | 0,6765013 | 0,322259 | P51649;C9J | ALDH5A1 |
| 2,696 | -0,366822 | 1,1258 | 1,1258 | 1,5315747 | 0,322569 | P50440;H0' | GATM |
| 1,18165 | 0,471213 | -0,150883 | 0,471213 | 0,6667544 | 0,323083 | Q96L34;Q6 | MARK4 |
| 0,578364 | -0,0891805 | 0,262915 | 0,262915 | 0,3339399 | 0,323161 | P16144;J3C | ITGB4 |
| 0,0901708 | -0,0228586 | 0,292957 | 0,0901708 | 0,1600195 | 0,323289 | Q99961;M( | SH3GL1 |
| 0,768808 | -2,2105 | -3,22893 | -2,2105 | 2,0774738 | 0,323809 | Q86YA3;G3 | ZGRF1 |
| 0,939781 | -0,280227 | 0,887665 | 0,887665 | 0,6898197 | 0,324674 | Q8N7U6 | EFHB |
| 0,70001 | -3,46631 | -1,96392 | -1,96392 | 2,1099732 | 0,324853 | A0A2R8YE( | NEXMIF |
| 3,83071 | 1,07229 | -0,242964 | 1,07229 | 2,0790059 | 0,324913 | Q9NT99;M( | LRRC4B |
| -0,013469 | 2,83493 | 0,551695 | 0,551695 | 1,5080865 | 0,325694 | O95359;E7I | TACC2 |
| -0,837077 | 0,285455 | -1,06568 | -0,837077 | 0,7231762 | 0,325748 | A0A2U3TZI | CCDC30 |
| -2,43791 | -0,747203 | 0,20055 | -0,747203 | 1,3365501 | 0,3263 | A1A4S6 | ARHGAP10 |
| -0,130044 | 0,375278 | 0,497177 | 0,375278 | 0,3325698 | 0,326412 | O43491;E9I | EPB41L2 |
| 0,0199873 | -2,65615 | -0,522119 | -0,522119 | 1,4147838 | 0,326413 | Q93084 | ATP2A3 |
| -0,0412544 | -0,355708 | -2,17786 | -0,355708 | 1,1535599 | 0,326459 | Q99715;D6 | COL12A1 |
| 1,0942 | -0,101513 | 0,353478 | 0,353478 | 0,6035196 | 0,326713 | Q8WZA9;M | IRGQ |
| 4,08925 | -0,995995 | 2,78757 | 2,78757 | 2,6416362 | 0,327427 | Q86YS7;H0 | C2CD5 |
| -2,84726 | 0,923885 | -3,07032 | -2,84726 | 2,2444362 | 0,327639 | Q8IX07;A0/ | ZFPM1 |
| -1,79115 | 0,0954999 | -0,460344 | -0,460344 | 0,9694891 | 0,327818 | Q15020;F8' | SART3 |
| -0,274893 | -0,269869 | -3,15361 | -0,274893 | 1,6634802 | 0,327913 | Q9NQT8;E7 | KIF13B |
| 1,52007 | 0,304941 | -0,0225846 | 0,304941 | 0,8127725 | 0,32885 | P22033 | MUT |
| -1,67463 | -3,35868 | 0,614263 | -1,67463 | 1,9941302 | 0,329116 | Q3T906;H0 | GNPTAB |
| 1,47574 | 2,1337 | -0,532447 | 1,47574 | 1,3888884 | 0,329215 | P53814;A0/ | SMTN |
| -0,30932 | -1,40005 | 0,0433482 | -0,30932 | 0,752492 | 0,329453 | P23378;A0/ | GLDC |
| 0,469798 | -2,5432 | -1,27303 | -1,27303 | 1,5126653 | 0,329742 | Q96G03;E7 | PGM2 |
| 1,77361 | 1,51856 | -0,518373 | 1,51856 | 1,2561404 | 0,330424 | Q14160;A0 | SCRIB |
| 0,575916 | 0,146222 | 4,22766 | 0,575916 | 2,2426921 | 0,33061 | P00918;E5F | CA2 |
| 3,17705 | 1,29696 | -0,468507 | 1,29696 | 1,8230788 | 0,332284 | Q9NPQ8;E9 | RIC8A |
| -0,0475398 | -0,328564 | -2,23605 | -0,328564 | 1,1907319 | 0,332852 | P31644 | GABRA5 |
| 2,55375 | 0,766674 | -0,228366 | 0,766674 | 1,4097231 | 0,332913 | Q9NQR4;F8 | NIT2 |
| 0,973086 | 2,04259 | -0,367974 | 0,973086 | 1,2078286 | 0,333126 | Q9P0V3;C9 | SH3BP4 |
| 0,448193 | -1,21321 | -2,9297 | -1,21321 | 1,6890214 | 0,333909 | Q6PL18 | ATAD2 |
| -0,901721 | 3,35949 | 2,45167 | 2,45167 | 2,2445225 | 0,333941 | Q8NDV7 | TNRC6A |
| -2,40613 | -0,814461 | 0,27436 | -0,814461 | 1,3480831 | 0,334247 | Q9UL46;A0 | PSME2 |
| -0,0732305 | 0,918686 | 0,254847 | 0,254847 | 0,5053407 | 0,335632 | Q9Y570 | PPME1 |
| 0,550386 | -2,03409 | -1,46475 | -1,46475 | 1,3579665 | 0,336677 | J3KPH8;Q8' | HDAC7 |

| | | | | | | | |
|---|---|---|---|---|---|---|---|
| -0,508942 | 0,173989 | -0,543589 | -0,508942 | 0,4046631 | 0,336709 | P78386 | KRT85 |
| -2,69275 | 0,266152 | -0,822103 | -0,822103 | 1,4965917 | 0,336762 | P59901;A0/ | LILRA4 |
| -2,64538 | -2,70424 | 0,887877 | -2,64538 | 2,0571288 | 0,337073 | P50148;B1/ | GNAQ |
| -0,490986 | 0,19312 | -1,04967 | -0,490986 | 0,6224489 | 0,337763 | O00533;A0. | CHL1 |
| 1,11741 | -0,317715 | 4,31311 | 1,11741 | 2,3705352 | 0,339154 | Q07812;K4. | BAX |
| 0,931307 | -0,25942 | 0,68231 | 0,68231 | 0,6280506 | 0,339262 | Q5H9U9 | DDX60L |
| -1,40365 | 0,456334 | -1,31472 | -1,31472 | 1,0491331 | 0,339278 | Q9NP71;H7 | MLXIPL |
| -1,80412 | -1,01546 | 0,410278 | -1,01546 | 1,1223689 | 0,340918 | Q15181;Q5 | PPA1 |
| -1,12398 | 0,438474 | -2,95959 | -1,12398 | 1,7008608 | 0,341533 | Q5JXI8 | FHL1 |
| -2,47503 | -0,602644 | 0,167023 | -0,602644 | 1,358839 | 0,34172 | P0DPH8;P0 | TUBA3D |
| 0,614375 | -2,06905 | -1,62633 | -1,62633 | 1,4386064 | 0,341782 | O00425;F8\ | IGF2BP3 |
| -1,21275 | -0,0401207 | -0,139141 | -0,139141 | 0,6503205 | 0,341984 | P62195;J3C | PSMC5 |
| -0,92903 | 0,167408 | -0,411982 | -0,411982 | 0,5485143 | 0,34214 | Q14568 | HSP90AA2P |
| -2,16152 | 0,657105 | -1,75436 | -1,75436 | 1,5234604 | 0,342234 | P29401;A0/ | TKT |
| -0,565109 | 1,52642 | 1,80737 | 1,52642 | 1,2962821 | 0,342793 | Q8TBY8;F5I | PMFBP1 |
| -0,405029 | 1,03977 | 1,42898 | 1,03977 | 0,9663089 | 0,342826 | Q9C0I4;C9J | THSD7B |
| 3,67764 | -1,12213 | 2,97942 | 2,97942 | 2,5931967 | 0,343049 | P20701;I3L | ITGAL |
| -2,02597 | -1,23089 | 0,500287 | -1,23089 | 1,2917108 | 0,343111 | P68402;J3K | PAFAH1B2 |
| 4,72539 | 0,182745 | 0,501803 | 0,501803 | 2,5356166 | 0,343193 | P07437;Q5. | TUBB |
| -1,74266 | -2,61582 | 0,696255 | -1,74266 | 1,7166127 | 0,343223 | O95352;C9. | ATG7 |
| 1,06101 | 1,96957 | -0,438379 | 1,06101 | 1,2159953 | 0,343512 | Q96JM2;H3 | ZNF462 |
| -0,00569722 | -0,975701 | -0,139096 | -0,139096 | 0,5257709 | 0,343619 | P50991 | CCT4 |
| -0,347533 | -0,689067 | 0,144059 | -0,347533 | 0,4188092 | 0,34362 | A0A0A0MR | AKAP9 |
| 0,0274248 | -0,0705053 | -0,2004 | -0,070505 | 0,1142855 | 0,34374 | Q96QB1;A0 | DLC1 |
| -1,35674 | -3,21448 | 0,555619 | -1,35674 | 1,8851154 | 0,343791 | H0YD86 | BRCA2 |
| -0,555551 | 0,0878243 | -2,99209 | -0,555551 | 1,6246295 | 0,343889 | Q5M9N0 | CCDC158 |
| -0,214333 | 2,82385 | 0,695714 | 0,695714 | 1,5592575 | 0,345624 | C9IZE1;F8W | EIF2A |
| 0,19381 | 1,43664 | 0,0115288 | 0,19381 | 0,7755423 | 0,346074 | O60486;F5I | PLXNC1 |
| -0,357899 | 0,135045 | -1,1729 | -0,357899 | 0,6605478 | 0,346813 | Q08380 | LGALS3BP |
| -1,30221 | -0,600362 | 0,255197 | -0,600362 | 0,7799667 | 0,346975 | Q15700;B7. | DLG2 |
| -0,363498 | 0,151349 | -0,582607 | -0,363498 | 0,3767775 | 0,347464 | Q12874 | SF3A3 |
| 1,98436 | 0,194252 | 0,0759993 | 0,194252 | 1,0692919 | 0,347614 | A8K3Y2;B5I | MAP2K6 |
| -0,0694585 | 0,37513 | 2,04911 | 0,37513 | 1,1171542 | 0,347733 | E9PR03 | IRF7 |
| 1,27875 | 4,03131 | -0,503153 | 1,27875 | 2,284481 | 0,348389 | P29400 | COL4A5 |
| -1,97895 | -0,560701 | 0,206583 | -0,560701 | 1,1088064 | 0,348394 | A0A2R8Y5C | ATP8B1 |
| -1,70584 | -0,382514 | 0,112465 | -0,382514 | 0,9400735 | 0,348801 | P14618;B4I | PKM |
| -1,70583 | -0,382491 | 0,112468 | -0,382491 | 0,9400725 | 0,348807 | H3BTN5 | PKM |
| 0,839493 | -2,0861 | -2,82867 | -2,0861 | 1,9393265 | 0,348883 | O60664;K7I | PLIN3 |
| 0,628419 | -0,207863 | 0,548678 | 0,548678 | 0,4615338 | 0,349126 | Q6QNY1;A0 | BLOC1S2 |
| -1,5873 | 0,687482 | -3,40588 | -1,5873 | 2,0509136 | 0,349237 | Q5VT06;H0 | CEP350 |
| 1,39183 | 0,226837 | -0,0292663 | 0,226837 | 0,7574422 | 0,34942 | P45974;F5H | USP5 |
| 0,154266 | 0,0499771 | 1,5642 | 0,154266 | 0,8457404 | 0,350743 | P05204 | HMGN2 |
| -0,3672 | 0,888228 | 1,26436 | 0,888228 | 0,8543577 | 0,350968 | Q9UK10;K7 | ZNF225 |
| 1,82699 | 2,42355 | -0,740208 | 1,82699 | 1,6810589 | 0,351252 | O14647;B7. | CHD2 |
| 0,309235 | -0,135431 | 0,758601 | 0,309235 | 0,4470181 | 0,351672 | A0A087X0T | CADM1 |
| -1,2232 | -1,10373 | 0,418449 | -1,10373 | 0,9152699 | 0,351794 | Q14194;E9I | CRMP1 |
| -1,07556 | 0,00984692 | -0,154243 | -0,154243 | 0,5850725 | 0,351797 | Q8IVM0 | CCDC50 |
| 0,676243 | -0,122695 | 3,98502 | 0,676243 | 2,1779053 | 0,352018 | Q5VYS8;Q5 | TUT7 |
| -0,98376 | 2,26419 | 3,85297 | 2,26419 | 2,4653383 | 0,352322 | Q8NEM7;B4 | SUPT20H |
| 2,14143 | 0,0929793 | 0,176064 | 0,176064 | 1,1594335 | 0,352904 | Q16623;A0 | STX1A |
| 0,492594 | -0,0571522 | 0,142599 | 0,142599 | 0,2782738 | 0,353225 | C9J1P0;C9J. | TMEM169 |
| 10,6617 | 1,36425 | 0,0148159 | 1,36425 | 5,7968338 | 0,353245 | Q14576;K7I | ELAVL3 |

| | | | | | | | |
|---|---|---|---|---|---|---|---|
| -3,23703 | 1,201 | -3,35983 | -3,23703 | 2,5984726 | 0,35335 | Q9GZT4;V9 | SRR |
| -1,37935 | 3,82895 | 3,71458 | 3,71458 | 2,9745474 | 0,354118 | P47755;F8V | CAPZA2 |
| -1,55257 | -1,99706 | 0,63266 | -1,55257 | 1,407612 | 0,354124 | P08574 | CYC1 |
| -2,08491 | 0,00972328 | -0,277156 | -0,277156 | 1,1356175 | 0,35428 | Q13200;C9. | PSMD2 |
| 0,709642 | 2,96996 | -0,253303 | 0,709642 | 1,6545757 | 0,354393 | M0QXB4;O | COPE |
| -0,733613 | 0,0599306 | -0,170817 | -0,170817 | 0,4081861 | 0,354737 | Q8NEV4;A0 | MYO3A |
| -0,0893552 | 0,222018 | 0,797693 | 0,222018 | 0,4500388 | 0,355034 | Q8WUM4;( | PDCD6IP |
| -1,08576 | -2,52661 | 0,491813 | -1,08576 | 1,5097275 | 0,355091 | A0A087WS | NUCB2 |
| -1,03256 | 2,29887 | 6,01811 | 2,29887 | 3,5271121 | 0,355401 | L7N2F4;K7I | AARSD1 |
| -1,60512 | -2,03746 | 0,656856 | -1,60512 | 1,4469964 | 0,355743 | Q9UNW9;F | NOVA2 |
| 0,961486 | 0,591792 | -0,261495 | 0,591792 | 0,6272234 | 0,35645 | Q9BTV4 | TMEM43 |
| -2,68651 | -0,689427 | 0,271165 | -0,689427 | 1,508803 | 0,356771 | Q04727 | TLE4 |
| 1,04614 | -2,48815 | -3,3329 | -2,48815 | 2,3231014 | 0,357205 | P40424 | PBX1 |
| 0,937968 | -2,61654 | -2,42474 | -2,42474 | 1,9991298 | 0,35773 | A0A0A0MR | MYEF2 |
| 1,04309 | -0,0230219 | 0,151963 | 0,151963 | 0,5717403 | 0,35821 | O43684;J3( | BUB3 |
| 0,882996 | 0,944476 | -0,341161 | 0,882996 | 0,725167 | 0,35827 | O75962;E7I | TRIO |
| 0,717573 | 2,39536 | -0,31258 | 0,717573 | 1,3668165 | 0,358417 | A2RUR9;C9 | CCDC144A |
| -1,50285 | 0,613392 | -1,82052 | -1,50285 | 1,3230849 | 0,358525 | Q13884 | SNTB1 |
| -0,632094 | 0,0541529 | -0,144213 | -0,144213 | 0,3531552 | 0,359146 | P01024;M0 | C3 |
| 0,587 | -1,56945 | -1,55126 | -1,55126 | 1,2398093 | 0,359374 | Q15349;F2. | RPS6KA2 |
| -0,130703 | 2,45883 | 0,448199 | 0,448199 | 1,3591324 | 0,359542 | P50914;E7E | RPL14 |
| 0,202668 | -0,566447 | -2,65429 | -0,566447 | 1,4783344 | 0,359762 | Q86X76 | NIT1 |
| -0,936555 | -2,11254 | 0,440587 | -0,936555 | 1,2778836 | 0,35981 | F8VNV8;P5 | CACNB3 |
| 1,24145 | -0,0488242 | 0,202858 | 0,202858 | 0,6839621 | 0,359991 | Q14195;H0 | DPYSL3 |
| -0,088212 | 0,188836 | 0,513086 | 0,188836 | 0,3009576 | 0,360196 | Q9H251;A0 | CDH23 |
| -1,98456 | 0,499328 | -1,07689 | -1,07689 | 1,2568497 | 0,360318 | P29728;A0 | OAS2 |
| 0,823552 | 1,82444 | -0,389749 | 0,823552 | 1,1087913 | 0,360663 | Q9H0E9;B5 | BRD8 |
| 0,150481 | -0,0653921 | 0,208832 | 0,150481 | 0,1444558 | 0,361033 | Q9BR84;A0 | ZNF559 |
| -0,246848 | 0,0328854 | -0,0732899 | -0,07329 | 0,1412128 | 0,361125 | Q9Y333 | LSM2 |
| -0,258422 | 0,669083 | 2,96735 | 0,669083 | 1,6607178 | 0,361146 | A0A2R8Y8( | AUTS2 |
| 0,970365 | -2,6991 | -2,40568 | -2,40568 | 2,039148 | 0,362366 | Q14590;K7I | ZNF235 |
| -0,265103 | 0,654177 | 2,80506 | 0,654177 | 1,5757154 | 0,362444 | Q6P2I3 | FAHD2B |
| 0,0949326 | -0,817451 | -0,216585 | -0,216585 | 0,4637756 | 0,362853 | P35606;D6I | COPB2 |
| 1,81723 | -0,494143 | 1,06075 | 1,06075 | 1,1784453 | 0,363236 | K7N7B3;Q5 | SYCP2L |
| -0,237472 | -2,93122 | -0,0752077 | -0,237472 | 1,6041308 | 0,363357 | O14744;G3 | PRMT5 |
| 0,00897629 | 0,0722599 | 0,762247 | 0,0722599 | 0,4178325 | 0,364014 | Q13428;J3k | TCOF1 |
| -0,403936 | 0,159913 | -1,89862 | -0,403936 | 1,0637641 | 0,36486 | Q7L591;D6 | DOK3 |
| -0,561307 | 1,57076 | 1,35344 | 1,35344 | 1,1732571 | 0,364907 | Q8NAP3;D6 | ZBTB38 |
| 3,16641 | 0,968109 | -0,457676 | 0,968109 | 1,825714 | 0,364914 | A8MXL6;P5 | SEC13 |
| 0,915843 | -2,73459 | -2,11038 | -2,11038 | 1,9524909 | 0,365207 | Q8NCN4 | RNF169 |
| 0,672224 | -0,328744 | 1,84109 | 0,672224 | 1,0859991 | 0,365356 | Q5T2S8;Q5 | ARMC4 |
| 0,0312849 | 1,85917 | 0,160214 | 0,160214 | 1,0201502 | 0,365622 | Q9NRW1;J3 | RAB6B |
| -0,747625 | -2,0193 | 0,367016 | -0,747625 | 1,1940188 | 0,365664 | Q13123;A0 | IK |
| -2,78273 | -0,498474 | 0,170832 | -0,498474 | 1,548619 | 0,365939 | P42695;G3 | NCAPD3 |
| -0,769405 | -1,95532 | 0,382123 | -0,769405 | 1,1687637 | 0,366722 | P48995 | TRPC1 |
| 0,99525 | -0,494909 | 2,47208 | 0,99525 | 1,4834995 | 0,366852 | Q8NDH2 | CCDC168 |
| 0,105083 | -0,229388 | -0,339349 | -0,229388 | 0,2314743 | 0,366967 | Q01469;I6L | FABP5 |
| 0,244086 | -0,0628381 | 1,66714 | 0,244086 | 0,9230483 | 0,367074 | Q86YW9;F8 | MED12L |
| -0,214189 | 0,485866 | 0,636811 | 0,485866 | 0,4540672 | 0,367397 | P54727;Q5 | RAD23B |
| -0,25401 | 1,22349 | 0,507102 | 0,507102 | 0,7388628 | 0,367839 | Q9UHV9 | PFDN2 |
| -1,37811 | 3,70748 | 3,32218 | 3,32218 | 2,8315016 | 0,368312 | Q5VZM2 | RRAGB |
| 4,86567 | -1,1171 | 2,23452 | 2,23452 | 2,9986064 | 0,368439 | Q9Y3R5 | DOP1B |

| | | | | | | | |
|---|---|---|---|---|---|---|---|
| -2,38648 | 0,663568 | -1,3758 | -1,3758 | 1,5536671 | 0,368599 | P29992;K7E | GNA11 |
| 0,918263 | -1,83617 | -3,84949 | -1,83617 | 2,3934573 | 0,369096 | Q5T7W0;B5 | ZNF618 |
| -2,31006 | 0,173403 | -0,438355 | -0,438355 | 1,2939034 | 0,369426 | P08670;B0Y | VIM |
| 0,00684361 | 0,24202 | 2,57608 | 0,24202 | 1,4203356 | 0,369652 | P24821;F5H | TNC |
| 0,256444 | -0,559422 | -0,791124 | -0,559422 | 0,5502608 | 0,369766 | P52948;H0Y | NUP98 |
| 1,13975 | -2,66443 | -3,1334 | -2,66443 | 2,3434849 | 0,369893 | Q02818;C9. | NUCB1 |
| -0,250416 | 0,0687326 | -1,76395 | -0,250416 | 0,9790613 | 0,369974 | O95678 | KRT75 |
| -0,200942 | 0,0529847 | -1,45069 | -0,200942 | 0,804921 | 0,370195 | P78352;C9J | DLG4 |
| 0,215755 | 1,33644 | -0,0736355 | 0,215755 | 0,7447589 | 0,370349 | P38159;H0Y | RBMX |
| -0,342911 | 1,5854 | 0,672325 | 0,672325 | 0,9646064 | 0,37039 | A0A1W2PP | SCN1A |
| 4,18337 | 0,0165209 | 0,374017 | 0,374017 | 2,309459 | 0,371264 | Q9UPV9;A0 | TRAK1 |
| 0,182299 | -0,0621557 | 1,15896 | 0,182299 | 0,6461099 | 0,37142 | Q96CW5 | TUBGCP3 |
| -0,204736 | -0,0329357 | -2,57644 | -0,204736 | 1,4214962 | 0,371423 | P52292;J3K | KPNA2 |
| 3,77656 | 0,773948 | -0,339261 | 0,773948 | 2,1289628 | 0,371732 | Q8TES7;A0 | FBF1 |
| 4,27243 | 0,725933 | -0,273565 | 0,725933 | 2,3889575 | 0,371791 | A0A0A0MR | NOLC1 |
| -0,247912 | 0,610274 | 0,624008 | 0,610274 | 0,4994858 | 0,372368 | P51858;H3I | HDGF |
| 0,657344 | -2,18247 | -1,36007 | -1,36007 | 1,4612121 | 0,372416 | P05387;H0Y | RPLP2 |
| -0,146548 | -0,0375576 | -2,07936 | -0,146548 | 1,1486658 | 0,373188 | A0RZB6;D6 | SIL1 |
| 1,59381 | 5,33255 | -0,820304 | 1,59381 | 3,1001004 | 0,373359 | A0A087WX | ASNA1 |
| 0,769402 | 3,00659 | -0,382131 | 0,769402 | 1,7231013 | 0,373361 | P47985;P0( | UQCRFS1 |
| -0,789861 | 1,65503 | 2,47606 | 1,65503 | 1,6989125 | 0,373929 | O43488;H3 | AKR7A2 |
| -0,341117 | 0,17986 | -1,00873 | -0,341117 | 0,5958006 | 0,374503 | P12235;V9( | SLC25A4 |
| 4,93515 | 0,173377 | 0,244026 | 0,244026 | 2,7290449 | 0,374968 | E9PDF6;O4 | MYO1B |
| 1,14295 | -3,29491 | -2,48048 | -2,48048 | 2,3624543 | 0,375059 | Q96JG6;H7 | VPS50 |
| -0,238613 | 2,68133 | 0,525704 | 0,525704 | 1,5142089 | 0,375152 | H0YMX3;Q9 | LINGO1 |
| 0,566214 | -1,23744 | -1,60367 | -1,23744 | 1,1615858 | 0,375529 | K7ESM5 | TUBB6 |
| -0,0461771 | -0,0405241 | -1,04467 | -0,046177 | 0,5781189 | 0,375812 | O43175;A0 | PHGDH |
| 0,223847 | 2,45556 | -0,0167451 | 0,223847 | 1,363251 | 0,376556 | Q9GZV7;Q5 | HAPLN2 |
| -1,43659 | 0,535909 | -1,20386 | -1,20386 | 1,0779388 | 0,37671 | Q5T200;E5I | ZC3H13 |
| 0,334439 | -0,0951014 | 2,73785 | 0,334439 | 1,5267887 | 0,377181 | O00410;E7I | IPO5 |
| -0,383026 | -0,864115 | 0,204716 | -0,383026 | 0,5353016 | 0,377691 | P07737;K7E | PFN1 |
| 0,906888 | -0,36035 | 0,84359 | 0,84359 | 0,7140694 | 0,377807 | Q8TF46 | DIS3L |
| -1,76597 | -2,9487 | 0,892848 | -1,76597 | 1,9674711 | 0,378643 | O94819 | KBTBD11 |
| -0,152299 | -0,664148 | 0,0783988 | -0,152299 | 0,3800409 | 0,378738 | P62333;A0 | PSMC6 |
| -0,162499 | 0,429071 | 3,05397 | 0,429071 | 1,7120029 | 0,379221 | Q75QN2;J3 | INTS8 |
| -0,802799 | 0,135467 | -0,24719 | -0,24719 | 0,4717822 | 0,379442 | Q12906;K7I | ILF3 |
| 1,30204 | -0,357563 | 0,67739 | 0,67739 | 0,8382121 | 0,380138 | Q01814;A0 | ATP2B2 |
| 0,219803 | -2,01511 | -0,429247 | -0,429247 | 1,1497147 | 0,380145 | E7EUS2;Q8 | CCDC110 |
| 0,0210013 | 0,0235382 | 0,593409 | 0,0235382 | 0,3297498 | 0,380191 | Q5VIR6;F6V | VPS53 |
| 1,03444 | -0,138437 | 0,25636 | 0,25636 | 0,5967849 | 0,38092 | A0A075B6F | DNAH8 |
| -0,017369 | 0,856416 | 0,0834582 | 0,0834582 | 0,4780394 | 0,381154 | Q10567;C9. | AP1B1 |
| -1,85556 | -1,07055 | 0,554738 | -1,07055 | 1,2293181 | 0,381303 | Q99832;F8Y | CCT7 |
| 0,65526 | -1,25197 | -2,25329 | -1,25197 | 1,4776012 | 0,381345 | Q9BPU6;E7 | DPYSL5 |
| -0,349747 | 1,3402 | 0,647929 | 0,647929 | 0,8495604 | 0,381404 | Q15276;A0 | RABEP1 |
| 0,205341 | -0,383136 | -1,6771 | -0,383136 | 0,9630016 | 0,381869 | Q13618;H7 | CUL3 |
| 1,16009 | -3,06378 | -2,51011 | -2,51011 | 2,2955753 | 0,382548 | Q96E40 | SPACA9 |
| 0,508333 | -2,44255 | -0,902061 | -0,902061 | 1,4759194 | 0,382754 | Q5T890 | ERCC6L2 |
| 0,014813 | -0,0914396 | -1,06048 | -0,09144 | 0,5925347 | 0,383281 | E9PKF4;J3K | FHL2 |
| 0,993914 | -1,92542 | -3,12915 | -1,92542 | 2,1201861 | 0,384042 | Q9H0U4;E9 | RAB1B |
| 0,193621 | -0,373387 | -2,00023 | -0,373387 | 1,1387931 | 0,384228 | Q86SQ0;A0 | PHLDB2 |
| -1,11659 | 0,376943 | -0,748103 | -0,748103 | 0,778048 | 0,384657 | Q9BTE6;C9. | AARSD1 |
| 0,078496 | -0,029249 | 0,667205 | 0,078496 | 0,3748856 | 0,384864 | Q8N7X0;H0 | ADGB |

| | | | | | | | |
|---|---|---|---|---|---|---|---|
| -0,31875 | 0,181266 | -1,27709 | -0,31875 | 0,7410841 | 0,385331 | Q0VDD8;H7 | DNAH14 |
| 0,612457 | 0,810509 | -0,296366 | 0,612457 | 0,590248 | 0,385351 | Q96DG6 | CMBL |
| -0,514523 | -1,43363 | 0,297195 | -0,514523 | 0,8659676 | 0,385792 | Q9Y371;A0 | SH3GLB1 |
| 0,0286452 | -1,38353 | -0,122432 | -0,122432 | 0,7753957 | 0,386043 | P10636;A0 | MAPT |
| -1,19464 | -1,39993 | 0,551762 | -1,19464 | 1,0724711 | 0,386138 | P18124;A8I | RPL7 |
| 1,21913 | -0,714068 | 4,02761 | 1,21913 | 2,3842652 | 0,38688 | Q9H0B3;A0 | IQCN |
| 0,0664653 | 1,17599 | 0,00819445 | 0,0664653 | 0,658051 | 0,386989 | Q92526;J3K | CCT6B |
| -0,191584 | 0,469191 | 0,42255 | 0,42255 | 0,3687726 | 0,387375 | P37108;H0Y | SRP14 |
| -1,12978 | 3,29197 | 2,21345 | 2,21345 | 2,3055103 | 0,387519 | Q1MSJ5 | CSPP1 |
| -0,193551 | 0,0259852 | -2,67197 | -0,193551 | 1,4983168 | 0,388075 | Q9NVP4 | DZANK1 |
| 0,236184 | -0,128161 | 1,32953 | 0,236184 | 0,7586189 | 0,388113 | P55795 | HNRNPH2 |
| 2,00649 | -0,985614 | 2,66308 | 2,00649 | 1,9449406 | 0,388283 | D6RBV2;D6 | LMAN2 |
| -0,33944 | 0,699325 | 0,900457 | 0,699325 | 0,6654361 | 0,388305 | O15371;B0 | EIF3D |
| 0,783773 | -0,203721 | 0,355799 | 0,355799 | 0,4952051 | 0,389153 | P55010;H0Y | EIF5 |
| 0,885057 | 1,43708 | -0,469741 | 0,885057 | 0,9811705 | 0,389533 | Q96GD0;B1 | PDXP |
| -0,248526 | 1,51008 | 0,414454 | 0,414454 | 0,8881285 | 0,3897 | O75323;H7 | NIPSNAP2 |
| 0,58922 | -0,318762 | 1,01505 | 0,58922 | 0,6812754 | 0,389742 | A0A0B4J2C | TPT1 |
| 5,87167 | 1,15094 | -0,669469 | 1,15094 | 3,3760352 | 0,390775 | G3V0E5;P0 | TFRC |
| -2,66088 | 0,106316 | -0,268903 | -0,268903 | 1,501095 | 0,390958 | Q13554;H7 | CAMK2B |
| -0,159997 | 0,0104492 | -2,68319 | -0,159997 | 1,5083792 | 0,391556 | Q8WXX0 | DNAH7 |
| -0,0742746 | 0,197106 | 0,149127 | 0,149127 | 0,1448319 | 0,391608 | Q8N8E3;F5 | CEP112 |
| -2,63341 | 0,00606078 | -0,151435 | -0,151435 | 1,4805298 | 0,391784 | Q9UQM7 | CAMK2A |
| 4,86931 | -0,703029 | 1,15803 | 1,15803 | 2,8369027 | 0,391801 | P48382;F8V | RFX5 |
| 1,09874 | -2,4094 | -2,58589 | -2,4094 | 2,0782482 | 0,392186 | P19012 | KRT15 |
| -1,95601 | -0,932199 | 0,539775 | -0,932199 | 1,2545809 | 0,392806 | O75390;B4 | CS |
| 0,526915 | -0,319385 | 1,3911 | 0,526915 | 0,8552581 | 0,393362 | A0A0A0MT | TCEA2 |
| -0,765398 | -1,78868 | 0,455676 | -0,765398 | 1,1236297 | 0,393701 | P50570;K7E | DNM2 |
| -2,73672 | 0,600053 | -0,9799 | -0,9799 | 1,6691676 | 0,39378 | P62937;C9J | PPIA |
| -0,519459 | -1,2866 | 0,313132 | -0,519459 | 0,8000891 | 0,394024 | F8W079 | ATP5F1B |
| 2,09448 | -0,0769994 | 0,190707 | 0,190707 | 1,1840144 | 0,394219 | Q13576;F5I | IQGAP2 |
| 1,87806 | 1,87784 | -0,834927 | 1,87784 | 1,5662803 | 0,394238 | Q8WVS4;H7 | WDR60 |
| 0,135723 | -0,221444 | -1,09537 | -0,221444 | 0,6333647 | 0,394263 | Q8N782;J3I | ZNF525 |
| 0,438311 | -1,35569 | -0,792761 | -0,792761 | 0,9175027 | 0,394444 | O15357;A0 | INPPL1 |
| -0,473807 | 2,09535 | 0,772095 | 0,772095 | 1,2847726 | 0,394614 | P10768;X6F | ESD |
| -0,970318 | 3,33256 | 1,6792 | 1,6792 | 2,1705723 | 0,394852 | P52179;A8I | MYOM1 |
| -0,156992 | -1,34589 | 0,0815663 | -0,156992 | 0,7646372 | 0,395495 | P04843;B7Z | RPN1 |
| -0,098845 | 2,32652 | 0,221071 | 0,221071 | 1,3176783 | 0,395583 | A0A2R8Y4H | LRRC9 |
| -1,16668 | 4,80939 | 1,90424 | 1,90424 | 2,9884182 | 0,396044 | A0A1B0GV( | FARP1 |
| -0,0944756 | 0,0475657 | -0,879742 | -0,094476 | 0,4994528 | 0,396213 | Q969Q0 | RPL36AL |
| -1,31323 | -0,060316 | -0,00105794 | -0,060316 | 0,7410691 | 0,396304 | P98172 | EFNB1 |
| -0,299831 | 3,29857 | 0,504204 | 0,504204 | 1,888714 | 0,39635 | F5H837;Q0 | RBL2 |
| 0,561405 | -1,2023 | -1,30273 | -1,2023 | 1,0484704 | 0,396536 | Q9BRP8 | PYM1 |
| 0,257449 | -1,81606 | -0,40106 | -0,40106 | 1,0595045 | 0,397398 | Q01968 | OCRL |
| 0,556086 | -1,19161 | -1,27822 | -1,19161 | 1,0349413 | 0,397497 | P17987;E7E | TCP1 |
| -0,12001 | 0,0650294 | -1,05215 | -0,12001 | 0,5987785 | 0,397529 | Q9HAV0;C9 | GNB4 |
| 0,513888 | -0,798043 | -2,86418 | -0,798043 | 1,7030085 | 0,397589 | O15195;E9I | VILL |
| -2,23092 | 0,306225 | -0,476226 | -0,476226 | 1,2992488 | 0,397747 | F5GZQ3;P5 | HADHB |
| 0,0718705 | -0,170803 | -2,13072 | -0,170803 | 1,2077231 | 0,398114 | Q9NZ71;F6 | RTEL1 |
| 1,47239 | -2,83506 | -3,85906 | -2,83506 | 2,8292243 | 0,398223 | O94823;A0 | ATP10B |
| 2,4276 | -0,615419 | 0,993888 | 0,993888 | 1,5223536 | 0,398722 | A0A087WZ | ZNF568 |
| 0,765954 | -2,17666 | -1,39951 | -1,39951 | 1,5249139 | 0,3988 | O15067;J3( | PFAS |
| 1,61151 | -0,767372 | 1,77768 | 1,61151 | 1,4238435 | 0,399113 | Q9BV35 | SLC25A23 |

| | | | | | | | |
|---|---|---|---|---|---|---|---|
| -0,288926 | 0,139504 | -0,32813 | -0,288926 | 0,259413 | 0,399211 | O60701;E7I | UGDH |
| 1,6846 | -0,710575 | 1,46047 | 1,46047 | 1,3229093 | 0,399343 | M0QWZ7;C | SARS2 |
| -0,195809 | 0,309927 | 2,00786 | 0,309927 | 1,1543326 | 0,399759 | P16401 | HIST1H1B |
| -2,01564 | 0,707391 | -1,27887 | -1,27887 | 1,4084839 | 0,400064 | P68133;A6I | ACTA1 |
| -0,434313 | -1,56407 | 0,287578 | -0,434313 | 0,9332808 | 0,400839 | Q92499;A0 | DDX1 |
| -0,355469 | 0,23679 | -1,68192 | -0,355469 | 0,9824876 | 0,400926 | Q5H9L2 | TCEAL5 |
| -2,10003 | -1,768 | 0,878635 | -1,768 | 1,6323483 | 0,401208 | P08779;K7E | KRT16 |
| -1,55484 | 0,0136928 | -0,0731235 | -0,073124 | 0,8816004 | 0,401267 | P49792 | RANBP2 |
| -1,27903 | 0,648562 | -1,58992 | -1,27903 | 1,2126462 | 0,401275 | F2Z2Z2;Q5T | DHX35 |
| -1,819 | 5,88033 | 3,06871 | 3,06871 | 3,8960364 | 0,401475 | P15313;C9J | ATP6V1B1 |
| 1,35132 | -0,55992 | 1,11011 | 1,11011 | 1,0408347 | 0,40214 | A6NMY6 | ANXA2P2 |
| -1,32474 | -1,36848 | 0,620005 | -1,32474 | 1,1356363 | 0,402415 | Q9UP95;I3I | SLC12A4 |
| 1,42545 | 2,37846 | -0,814559 | 1,42545 | 1,6391684 | 0,402815 | P11216;H0Y | PYGB |
| -1,09 | 8,12426 | 1,5994 | 1,5994 | 4,7383059 | 0,403153 | Q6PGP7 | TTC37 |
| -0,417502 | -0,718001 | 0,241375 | -0,417502 | 0,4907173 | 0,403157 | O00534;B4 | VWA5A |
| 0,0913648 | -0,822055 | -0,135761 | -0,135761 | 0,475556 | 0,403179 | Q16181;E7I | SEPT7 |
| 1,41822 | 1,16456 | -0,590464 | 1,16456 | 1,0938665 | 0,403311 | O14525 | ASTN1 |
| 0,906885 | -1,79289 | -2,17088 | -1,79289 | 1,6785062 | 0,403344 | O95834;K7I | EML2 |
| -1,856 | 0,390999 | -0,585209 | -0,585209 | 1,1267132 | 0,403672 | Q9P1U1;A0 | ACTR3B |
| -2,666 | 0,103733 | -0,199626 | -0,199626 | 1,5191252 | 0,404001 | Q13557;D6 | CAMK2D |
| -1,32545 | 3,15192 | 2,61022 | 2,61022 | 2,4436919 | 0,404533 | Q9GZR1 | SENP6 |
| -0,321241 | 0,207722 | -2,77647 | -0,321241 | 1,5923438 | 0,404669 | P09972;A8I | ALDOC |
| -0,404345 | 0,260566 | -1,00611 | -0,404345 | 0,6336003 | 0,404685 | Q8NCQ5;J3 | FBXO15 |
| 0,753063 | -1,62923 | -1,60468 | -1,60468 | 1,3683856 | 0,405083 | Q13085;Q5 | ACACA |
| -2,05583 | -2,29749 | 1,01107 | -2,05583 | 1,8443989 | 0,405266 | J3KQB2;Q9 | TTLL3 |
| -1,7313 | 4,50468 | 3,17952 | 3,17952 | 3,285312 | 0,405295 | H0Y360;Q0 | AMPD2 |
| 0,813875 | -1,76573 | -1,71063 | -1,71063 | 1,4736872 | 0,406418 | P00492 | HPRT1 |
| 0,431276 | -1,67818 | -0,656966 | -0,656966 | 1,0549055 | 0,406824 | Q9HAV4;H( | XPO5 |
| 5,2595 | 3,04068 | -1,79473 | 3,04068 | 3,6070882 | 0,407093 | E7EVM7;Q9 | PIEZO2 |
| 2,38159 | -0,783604 | 1,3034 | 1,3034 | 1,6091681 | 0,407196 | Q6PCE3 | PGM2L1 |
| -2,28344 | 0,224663 | -0,31896 | -0,31896 | 1,3194246 | 0,407395 | Q9C040;A0 | TRIM2 |
| 0,393473 | -0,603633 | -1,47077 | -0,603633 | 0,9328763 | 0,407442 | Q9H9B1;A0 | EHMT1 |
| -0,796351 | 2,10536 | 1,42382 | 1,42382 | 1,5173213 | 0,407611 | P41223;C9J | BUD31 |
| 0,0406452 | 0,00621698 | 1,7823 | 0,0406452 | 1,0156293 | 0,407627 | Q8TF09;H3 | DYNLRB2 |
| -0,0686825 | -1,35659 | 0,0319113 | -0,068683 | 0,7742481 | 0,407922 | O95336;M( | PGLS |
| -0,275233 | 5,96169 | 0,449542 | 0,449542 | 3,4109696 | 0,408078 | E5RHN2;E5 | CCNE2 |
| 4,36225 | 8,4096 | -2,68462 | 4,36225 | 5,6142844 | 0,408548 | H0YJ82 | ABCD4 |
| -0,73022 | -1,2143 | 0,428369 | -0,73022 | 0,8440995 | 0,408661 | P48668;P0 | KRT6C |
| 1,29587 | 0,733263 | -0,439773 | 0,733263 | 0,8855315 | 0,408954 | Q9H2D6;F6 | TRIOBP |
| 0,725398 | 0,113069 | -0,0825959 | 0,113069 | 0,4215222 | 0,4093 | Q58FF6 | HSP90AB4P |
| 0,0605704 | 2,03563 | -0,0140688 | 0,0605704 | 1,1624469 | 0,409736 | Q86YV5 | PRAG1 |
| -0,58535 | 0,20226 | -0,338409 | -0,338409 | 0,4028301 | 0,409755 | O00567;H0 | NOP56 |
| 0,0697031 | 0,212225 | -0,0484534 | 0,0697031 | 0,1305288 | 0,410269 | A0A1B0GXI | VPS45 |
| 0,392017 | -0,258629 | 0,931125 | 0,392017 | 0,5957477 | 0,410663 | P26378;B1/ | ELAVL4 |
| 0,292824 | -0,389728 | -2,41443 | -0,389728 | 1,4079844 | 0,411355 | Q5VUJ9;H7 | EFCAB2 |
| -0,109821 | 1,52425 | 0,15063 | 0,15063 | 0,8779574 | 0,411576 | Q9P0L2;A0 | MARK1 |
| -2,26114 | 0,722569 | -1,14435 | -1,14435 | 1,5074883 | 0,4122 | P22392;Q3 | NME2 |
| -0,0577025 | 1,2964 | 0,0851041 | 0,0851041 | 0,744001 | 0,412292 | Q9NX14 | NDUFB11 |
| 0,291468 | 2,04938 | -0,222528 | 0,291468 | 1,1913586 | 0,412558 | P08183;E7E | ABCB1 |
| -1,06736 | 1,88969 | 2,70857 | 1,88969 | 1,9863013 | 0,412654 | P49916;K7E | LIG3 |
| 1,23866 | -2,52554 | -2,63201 | -2,52554 | 2,2046399 | 0,412669 | Q9BRJ6;C9J | C7orf50 |
| 0,068146 | -2,30854 | -0,111206 | -0,111206 | 1,3234475 | 0,412817 | Q14204;A0 | DYNC1H1 |

| | | | | | | | |
|---|---|---|---|---|---|---|---|
| -0,0148016 | 0,0516681 | 0,0222702 | 0,0222702 | 0,0333086 | 0,413132 | P36551 | CPOX |
| -2,18981 | 1,07841 | -2,28968 | -2,18981 | 1,9163884 | 0,413279 | O75061;S4I | DNAJC6 |
| 0,369979 | -1,0916 | -0,596605 | -0,596605 | 0,7433615 | 0,413585 | P10155;G5I | TROVE2 |
| -2,90858 | 1,38944 | -2,85053 | -2,85053 | 2,4648763 | 0,413707 | F5H7N0;P2 | GABRB3 |
| 0,960452 | -3,0483 | -1,49388 | -1,49388 | 2,0211408 | 0,413845 | Q53FP2 | TMEM35A |
| -0,813685 | 1,23989 | 2,69806 | 1,23989 | 1,7642649 | 0,414121 | A2A2F0;Q8 | RALGAPB |
| 2,83348 | -0,851028 | 1,29225 | 1,29225 | 1,8504337 | 0,414375 | Q8TDM6 | DLG5 |
| 3,09376 | -0,722408 | 1,00769 | 1,00769 | 1,9108491 | 0,414668 | O15169;H0 | AXIN1 |
| 0,825322 | -3,31089 | -1,17119 | -1,17119 | 2,068519 | 0,414784 | F8VR40 | IRAK4 |
| -3,19727 | 1,51266 | -3,04575 | -3,04575 | 2,6766117 | 0,414896 | O75334;G3 | PPFIA2 |
| -2,87441 | -1,93645 | 1,12041 | -1,93645 | 2,088969 | 0,415039 | O15360;H3 | FANCA |
| 0,64993 | -0,347375 | 0,795866 | 0,64993 | 0,6222157 | 0,415323 | Q9Y2X3;H7 | NOP58 |
| -1,80973 | 0,729149 | -1,27513 | -1,27513 | 1,3384592 | 0,416484 | P48506;A0 | GCLC |
| -2,54636 | -1,01097 | 0,700786 | -1,01097 | 1,6243711 | 0,416805 | Q99755;A6 | PIP5K1A |
| 0,579506 | -1,24298 | -1,1284 | -1,1284 | 1,0207454 | 0,417484 | P10586;A2 | PTPRF |
| 1,37846 | -0,249129 | 0,319699 | 0,319699 | 0,8259929 | 0,41774 | Q86SX6 | GLRX5 |
| -0,0739118 | 0,0912069 | 0,495786 | 0,0912069 | 0,2931166 | 0,418586 | Q86UL8;A0 | MAGI2 |
| -1,79882 | 0,336997 | -0,430883 | -0,430883 | 1,0818661 | 0,418796 | Q07092 | COL16A1 |
| -0,872257 | 0,418303 | -0,825016 | -0,825016 | 0,7318491 | 0,419213 | O14950;J3 | MYL12B |
| 0,38046 | -0,796056 | -0,747694 | -0,747694 | 0,6657402 | 0,419262 | Q8N8N7;G | PTGR2 |
| 0,72116 | -0,445817 | 1,21345 | 0,72116 | 0,8521885 | 0,419337 | Q52LW3;F8 | ARHGAP29 |
| -2,25684 | 1,13921 | -2,34967 | -2,25684 | 1,98805 | 0,419988 | O75746 | SLC25A12 |
| 0,543232 | -0,753998 | -2,05474 | -0,753998 | 1,2989864 | 0,419991 | P10827;J3K | THRA |
| -0,416203 | 0,189655 | -0,354651 | -0,354651 | 0,333447 | 0,420224 | P42167;G5I | TMPO |
| -0,416189 | 0,189674 | -0,354652 | -0,354652 | 0,3334535 | 0,420253 | P42166 | TMPO |
| 0,913918 | 0,102722 | -0,0888793 | 0,102722 | 0,5323458 | 0,420273 | Q6WCQ1;J | MPRIP |
| 0,285961 | -2,6246 | -0,332133 | -0,332133 | 1,5334498 | 0,420517 | Q8N608 | DPP10 |
| -0,482232 | 0,352309 | -1,33079 | -0,482232 | 0,8415592 | 0,421835 | Q9Y3L5;F6 | RAP2C |
| -0,362158 | 1,00808 | 0,565821 | 0,565821 | 0,6993199 | 0,422496 | A0A0A0MT | CTNNA2 |
| 1,58563 | -3,1323 | -3,20729 | -3,1323 | 2,7458019 | 0,422804 | O14818;H0 | PSMA7 |
| 0,404579 | 2,54414 | -0,350275 | 0,404579 | 1,5013974 | 0,422954 | G3V180;G3 | DPP3 |
| 1,72442 | 0,230493 | -0,204756 | 0,230493 | 1,0118448 | 0,423179 | D6RAS9;O1 | RASGRF2 |
| -2,7964 | 1,57637 | -3,59083 | -2,7964 | 2,7824507 | 0,423327 | F8VQS4;F8 | SLC16A7 |
| 0,835035 | 2,99904 | -0,656593 | 0,835035 | 1,8380934 | 0,423399 | Q05BV3;H0 | EML5 |
| 0,628106 | 0,727663 | -0,338799 | 0,628106 | 0,5890894 | 0,42392 | A0A1W2PC | FRMD4A |
| 8,96751 | 0,651049 | -0,64403 | 0,651049 | 5,2157207 | 0,425187 | Q9UHB7;C | AFF4 |
| -0,444057 | 3,36378 | 0,490513 | 0,490513 | 1,9844692 | 0,425681 | P98082 | DAB2 |
| 2,16447 | 0,099813 | -0,107521 | 0,099813 | 1,2561674 | 0,426023 | A0A1B0GW | ALDH7A1 |
| -0,272649 | 1,03908 | 0,359275 | 0,359275 | 0,6560101 | 0,426235 | Q9P2R6;H7 | RERE |
| -1,79362 | -0,544776 | 0,429031 | -0,544776 | 1,114158 | 0,426743 | P56385 | ATP5ME |
| 0,0571625 | -0,605958 | -0,058283 | -0,058283 | 0,3542608 | 0,426761 | Q86SQ7;A0 | SDCCAG8 |
| 0,458322 | -0,52491 | -2,67503 | -0,52491 | 1,6024802 | 0,427388 | Q9BX68 | HINT2 |
| 0,21602 | -3,42533 | -0,198163 | -0,198163 | 1,9935556 | 0,427751 | M0QZ12;Q | GRAMD1A |
| 0,37072 | -0,991745 | -0,570338 | -0,570338 | 0,6975535 | 0,42805 | P62266;D6I | RPS23 |
| 1,10094 | -0,55206 | 1,06094 | 1,06094 | 0,9430251 | 0,428239 | P05787;F8 | KRT8 |
| 1,28522 | -2,84826 | -2,24736 | -2,24736 | 2,2333032 | 0,428444 | Q96J66;H3I | ABCC11 |
| 3,2113 | -0,164357 | 0,136884 | 0,136884 | 1,8680578 | 0,428855 | Q9BPW8;H | NIPSNAP1 |
| 0,104785 | -0,122497 | 2,18156 | 0,104785 | 1,2697329 | 0,428894 | Q5ST30 | VARS2 |
| -0,392873 | -1,66527 | 0,3348 | -0,392873 | 1,0123226 | 0,429303 | B7WNT5;P | TCF7 |
| -0,864666 | 2,32843 | 1,31284 | 1,31284 | 1,6314011 | 0,429391 | O95248 | SBF1 |
| -0,374048 | -2,57357 | 0,35585 | -0,374048 | 1,5249125 | 0,429925 | P63104;E7E | YWHAZ |
| 0,751842 | 0,522616 | -0,317874 | 0,522616 | 0,563214 | 0,430192 | B7ZAP0;F5I | RABGAP1L |

| | | | | | | | |
|---|---|---|---|---|---|---|---|
| 1,11884 | -0,77667 | 2,25427 | 1,11884 | 1,5312716 | 0,430833 | P43034;I3L | PAFAH1B1 |
| 0,449927 | -0,744643 | -1,04789 | -0,744643 | 0,7918767 | 0,430864 | Q99447;I3L | PCYT2 |
| -0,00088279 | -1,88021 | 0,0277643 | -0,000883 | 1,0933936 | 0,430966 | H0Y786;H7 | NEB |
| -0,333676 | 1,16377 | 0,438637 | 0,438637 | 0,7488469 | 0,431143 | Q7KZF4;H7 | SND1 |
| -2,2705 | 1,16199 | -2,2285 | -2,2285 | 1,9697366 | 0,431167 | P60900;G3 | PSMA6 |
| 0,80675 | -1,2167 | -2,14863 | -1,2167 | 1,5109111 | 0,431335 | Q8TC05 | MDM1 |
| 2,161 | 1,78467 | -1,01129 | 1,78467 | 1,7331301 | 0,431401 | P52737;C9J | ZNF136 |
| -0,488277 | -0,651187 | 0,288926 | -0,488277 | 0,5023937 | 0,431432 | Q13404;I3L | UBE2V1 |
| -0,969983 | 1,74266 | 2,01233 | 1,74266 | 1,6495123 | 0,432477 | Q9HD45 | TM9SF3 |
| 2,39697 | 0,0650018 | -0,104249 | 0,0650018 | 1,397785 | 0,432848 | Q9Y620;E5I | RAD54B |
| 0,12627 | 1,05898 | -0,130161 | 0,12627 | 0,625801 | 0,433024 | Q9Y5X3 | SNX5 |
| 0,140918 | -0,0984093 | -2,63858 | -0,098409 | 1,5403113 | 0,433153 | Q66K74;M( | MAP1S |
| -1,234 | -0,217356 | 0,205596 | -0,217356 | 0,73992 | 0,433559 | Q12802;H0 | AKAP13 |
| -0,0831592 | -0,635277 | 0,0844691 | -0,083159 | 0,3766005 | 0,433615 | Q6UWP8;K | SBSN |
| -2,56265 | -0,00847113 | 0,0582442 | -0,008471 | 1,4942873 | 0,434013 | A0A087WV | EEF1A1 |
| 1,67396 | -3,44868 | -2,97851 | -2,97851 | 2,8316067 | 0,434704 | Q9UHC1 | MLH3 |
| 5,23052 | -1,67606 | 2,25815 | 2,25815 | 3,4644345 | 0,434897 | H7BYX7;Q6 | PLB1 |
| 2,48827 | -0,834612 | 1,14687 | 1,14687 | 1,6716843 | 0,435476 | Q6AW86;M | ZNF324B |
| -0,0733084 | 0,0638761 | 0,713278 | 0,0638761 | 0,4201707 | 0,435505 | P62857 | RPS28 |
| 6,84063 | 2,59092 | -2,01421 | 2,59092 | 4,4286087 | 0,435568 | P17040 | ZSCAN20 |
| 4,06303 | 5,67084 | -2,4967 | 4,06303 | 4,3267351 | 0,436075 | P21579;J3K | SYT1 |
| 1,07329 | -0,330906 | 0,432864 | 0,432864 | 0,7030003 | 0,436283 | Q86W92;A( | PPFIBP1 |
| -1,29687 | 0,31567 | -0,371089 | -0,371089 | 0,8092171 | 0,436433 | D6RG30;Q2 | ASTE1 |
| -2,1543 | 0,917731 | -1,45187 | -1,45187 | 1,6096475 | 0,436641 | A0A087WV | BICRA |
| -1,43407 | 0,87574 | -1,95995 | -1,43407 | 1,5084711 | 0,436819 | P15121;E9F | AKR1B1 |
| 0,200468 | 2,67738 | -0,24888 | 0,200468 | 1,5758598 | 0,437085 | Q92618;A0 | ZNF516 |
| 0,435117 | 0,277477 | -0,179857 | 0,277477 | 0,319426 | 0,437197 | O14910;H0 | LIN7A |
| -0,169261 | -1,05859 | 0,169857 | -0,169261 | 0,6344274 | 0,437232 | Q5JTJ3 | COA6 |
| 1,32482 | -0,439435 | 0,590087 | 0,590087 | 0,8862227 | 0,437864 | O76070 | SNCG |
| -2,86473 | 1,59437 | -3,15766 | -2,86473 | 2,6630549 | 0,438359 | O15144;H7 | ARPC2 |
| 4,2052 | 0,0582669 | -0,169619 | 0,0582669 | 2,4626553 | 0,438448 | P61018;M0 | RAB4B |
| -1,18316 | 4,44 | 1,40678 | 1,40678 | 2,8144905 | 0,439693 | P57740;H0 | NUP107 |
| -0,466364 | -0,0154296 | 0,0282483 | -0,01543 | 0,2738281 | 0,439852 | K7EJR9;K7E | KCNJ16 |
| -0,96425 | 0,186791 | -0,193149 | -0,193149 | 0,5864933 | 0,440172 | B7Z2U2;Q6 | TOM1L2 |
| -0,0338795 | 0,009905 | 0,798578 | 0,009905 | 0,4684918 | 0,440529 | Q96D15;M( | RCN3 |
| 0,54646 | -0,752067 | -1,51786 | -0,752067 | 1,0435539 | 0,440962 | P02545;Q5 | LMNA |
| 0,0328886 | -0,884766 | -0,00502902 | -0,005029 | 0,5192085 | 0,441224 | Q99962 | SH3GL2 |
| 1,6758 | -2,90346 | -3,37597 | -2,90346 | 2,7902589 | 0,441344 | Q6ZQQ6;E7 | WDR87 |
| 0,791922 | -1,70154 | -1,29831 | -1,29831 | 1,3384703 | 0,441415 | P62318;H3I | SNRPD3 |
| 1,03979 | -2,50293 | -1,56445 | -1,56445 | 1,8354689 | 0,441438 | Q8IVH8;F8 | MAP4K3 |
| -0,186292 | 0,843694 | 0,20054 | 0,20054 | 0,5202815 | 0,441553 | A0A2R8Y59 | CTCF |
| -2,2606 | 1,24867 | -2,38772 | -2,2606 | 2,0637534 | 0,441945 | Q8N1F7;H3 | NUP93 |
| 0,04446 | 0,00439519 | -1,43835 | 0,0043952 | 0,8447726 | 0,442529 | P14868;C9J | DARS |
| 0,789953 | -2,71279 | -0,955527 | -0,955527 | 1,7513748 | 0,442839 | Q8N3U4;B1 | STAG2 |
| -1,43001 | -2,14362 | 0,931112 | -1,43001 | 1,60925 | 0,443169 | M0R2K1;Q | RSPH6A |
| 3,27255 | 0,434766 | -0,491548 | 0,434766 | 1,9612712 | 0,443741 | Q53GS7 | GLE1 |
| -0,70034 | 1,10376 | 1,54578 | 1,10376 | 1,1899029 | 0,444096 | A6NE02 | BTBD17 |
| -0,100874 | 0,076964 | 0,867386 | 0,076964 | 0,5154158 | 0,444477 | A0A0A0MR | ELAVL2 |
| -0,831792 | -1,9798 | 0,65217 | -0,831792 | 1,3195537 | 0,444481 | Q86UP2;G3 | KTN1 |
| 0,205744 | -0,0945586 | -3,14119 | -0,094559 | 1,851761 | 0,444527 | E9PF63;O7 | ROCK2 |
| -0,460389 | 1,17257 | 0,652577 | 0,652577 | 0,8342303 | 0,444605 | Q15075 | EEA1 |
| -1,06307 | -2,42262 | 0,821091 | -1,06307 | 1,6289107 | 0,444636 | Q9Y4J8;M0 | DTNA |

| | | | | | | | |
|---|---|---|---|---|---|---|---|
| -2,52596 | -2,14273 | 1,26065 | -2,14273 | 2,0843974 | 0,444823 | O15063;A0 | KIAA0355 |
| -1,41053 | -1,2162 | 0,710497 | -1,2162 | 1,1725102 | 0,444993 | P09104;F5H | ENO2 |
| 1,9431 | -0,382443 | 0,376507 | 0,376507 | 1,1859152 | 0,445186 | Q9UBC2;M | EPS15L1 |
| -0,60925 | 0,732417 | 2,02176 | 0,732417 | 1,3155917 | 0,445913 | B1AMU4;B | EXOSC1 |
| -0,38636 | 1,08151 | 0,510272 | 0,510272 | 0,7399216 | 0,446211 | G5E994;Q5 | GPR107 |
| -2,71107 | -0,835795 | 0,735881 | -0,835795 | 1,7257024 | 0,446265 | Q86VH2;F2 | KIF27 |
| -2,68536 | 0,32616 | -0,240519 | -0,240519 | 1,6003971 | 0,447321 | Q8N4V2;H( | SVOP |
| 0,709615 | -0,0875291 | 0,0651456 | 0,0651456 | 0,4231015 | 0,447356 | E9PNZ4;E9I | MACF1 |
| 1,18473 | 2,62629 | -0,917452 | 1,18473 | 1,7821042 | 0,447496 | P30038;Q5 | ALDH4A1 |
| -0,00480406 | 1,8935 | -0,0754079 | -0,004804 | 1,116926 | 0,447547 | A6NCI4 | VWA3A |
| -0,0789112 | 1,31868 | 0,0241405 | 0,0241405 | 0,7788574 | 0,44771 | E7EVD6;Q9 | ADGRL3 |
| 0,706209 | -1,42879 | -1,16331 | -1,16331 | 1,1636009 | 0,448191 | O43166;G3 | SIPA1L1 |
| 0,480831 | -1,678 | -0,551936 | -0,551936 | 1,0797514 | 0,448387 | Q9H3K6;A0 | BOLA2 |
| 2,62144 | 1,45565 | -1,04596 | 1,45565 | 1,8738081 | 0,448928 | Q9BYV8 | CEP41 |
| 0,185971 | -1,22558 | -0,151088 | -0,151088 | 0,7371813 | 0,449505 | O75335;B1 | PPFIA4 |
| -0,138604 | 1,23122 | 0,0918467 | 0,0918467 | 0,7334504 | 0,449572 | Q9UJU6;B4 | DBNL |
| -1,75957 | -0,284466 | 0,31866 | -0,284466 | 1,0691689 | 0,449849 | A0A0C4DGI | SYNE2 |
| 0,553156 | -0,919795 | -1,09078 | -0,919795 | 0,9038202 | 0,450144 | Q8N2N9;A( | ANKRD36B |
| 0,635133 | -0,73262 | -2,12941 | -0,73262 | 1,3822969 | 0,450501 | O75373;M( | ZNF737 |
| 3,35352 | 0,33584 | -0,457031 | 0,33584 | 2,0106108 | 0,451306 | Q99613;B5 | EIF3C |
| 0,656256 | -1,86465 | -0,828241 | -0,828241 | 1,2670729 | 0,451373 | Q13283;E5I | G3BP1 |
| -0,40536 | 10,2998 | -0,0992451 | -0,099245 | 6,0941816 | 0,451385 | Q5MIZ7 | PPP4R3B |
| -0,170353 | 0,345791 | 0,272842 | 0,272842 | 0,279329 | 0,451973 | Q7Z3B4 | NUP54 |
| -0,169531 | 0,200185 | 0,530503 | 0,200185 | 0,3502017 | 0,452561 | P06753 | TPM3 |
| 0,564879 | -0,541791 | -2,5728 | -0,541791 | 1,5913697 | 0,452601 | P18084;V9( | ITGB5 |
| 0,308189 | -0,184932 | -2,75801 | -0,184932 | 1,6464849 | 0,453076 | Q00341;H0 | HDLBP |
| -0,609078 | 0,604252 | 2,55675 | 0,604252 | 1,5972312 | 0,453678 | Q8TDW7;E | FAT3 |
| -0,725145 | -0,757565 | 0,41603 | -0,725145 | 0,6684131 | 0,454129 | Q9BQE3;F5 | TUBA1C |
| -3,13277 | 1,68008 | -2,84844 | -2,84844 | 2,7003661 | 0,454861 | Q01850 | CDR2 |
| 1,50193 | -2,78754 | -2,55709 | -2,55709 | 2,4127544 | 0,454894 | Q8NDA2 | HMCN2 |
| 2,83711 | -0,607487 | 0,555002 | 0,555002 | 1,7523625 | 0,455758 | Q8N823;A0 | ZNF611 |
| 0,631005 | -0,727478 | -1,9575 | -0,727478 | 1,2947837 | 0,456415 | P11413;E7E | G6PD |
| -1,16323 | -2,26587 | 0,890055 | -1,16323 | 1,6016479 | 0,456675 | Q9BQS8;C9 | FYCO1 |
| 0,422785 | -0,338653 | 0,920816 | 0,422785 | 0,6343087 | 0,456904 | Q08174;D6 | PCDH1 |
| -0,341919 | 0,770946 | 0,485896 | 0,485896 | 0,5780715 | 0,457292 | P04075;J3K | ALDOA |
| -0,268259 | 0,253102 | 1,15134 | 0,253102 | 0,7180889 | 0,457409 | Q16566;D6 | CAMK4 |
| -1,05831 | -0,238548 | 0,250643 | -0,238548 | 0,6613969 | 0,457508 | D3YTB5;P5 | IRAK1 |
| 0,233366 | -2,8078 | -0,0723814 | -0,072381 | 1,674549 | 0,457801 | Q9Y3S1;F8 | WNK2 |
| 0,18577 | -0,229214 | -0,506625 | -0,229214 | 0,3484679 | 0,458304 | Q9UGU0;I3 | TCF20 |
| -0,24832 | 3,1113 | 0,0645744 | 0,0645744 | 1,8559583 | 0,458584 | Q9BYP7;A0 | WNK3 |
| 3,47766 | -1,75831 | 2,74427 | 2,74427 | 2,8350917 | 0,459305 | Q9H1J1;A0 | UPF3A |
| -1,05625 | 1,77648 | 1,92305 | 1,77648 | 1,6793883 | 0,459416 | P49454;A0 | CENPF |
| 3,06154 | 0,0915512 | -0,279083 | 0,0915512 | 1,8311183 | 0,460492 | Q9BX66 | SORBS1 |
| 0,34333 | -0,0966436 | 0,0992225 | 0,0992225 | 0,2204271 | 0,460558 | Q9P258 | RCC2 |
| 0,379643 | -1,49104 | -0,364572 | -0,364572 | 0,9418281 | 0,461078 | A0A0A0MT | NLGN4X |
| 1,85983 | 1,01653 | -0,771655 | 1,01653 | 1,3437183 | 0,461274 | A0A087WT | CASP8AP2 |
| -0,656117 | 0,601995 | -1,91938 | -0,656117 | 1,2606884 | 0,461497 | H0YAM0 | CPE |
| -0,941782 | 0,695892 | -1,60463 | -0,941782 | 1,1841836 | 0,462162 | P11532;A0 | DMD |
| -1,48158 | 2,01163 | 3,39138 | 2,01163 | 2,5117048 | 0,462513 | A8K0Z3 | WASHC1 |
| 0,381333 | -1,97732 | -0,298683 | -0,298683 | 1,2140489 | 0,46267 | Q9NR09;H7 | BIRC6 |
| 0,500886 | -0,523135 | -1,66545 | -0,523135 | 1,0837062 | 0,463474 | F8VXC8;F8 | SMARCC2 |
| 0,349654 | -0,41906 | -0,942359 | -0,41906 | 0,6498796 | 0,463595 | P42285;H0 | MTREX |

| | | | | | | | |
|---|---|---|---|---|---|---|---|
| -2,59111 | 4,03538 | 4,96854 | 4,03538 | 4,1216798 | 0,463834 | Q9P0U3;F8 | SENP1 |
| -0,311079 | 2,19381 | 0,181311 | 0,181311 | 1,3270939 | 0,463973 | P54707 | ATP12A |
| -0,585132 | -0,60204 | 0,34465 | -0,585132 | 0,5417568 | 0,464007 | O60925;E5I | PFDN1 |
| 1,96996 | -0,0526196 | -0,0876624 | -0,05262 | 1,1779832 | 0,464484 | Q5VZ46;A0 | KIAA1614 |
| -0,0964834 | 1,37422 | 0 | 0 | 0,8226743 | 0,464503 | P49736;H0Y | MCM2 |
| -0,366945 | 0,778405 | 0,520916 | 0,520916 | 0,6008916 | 0,464872 | O94762;J3K | RECQL5 |
| -1,87639 | -0,897287 | 0,732863 | -0,897287 | 1,3180941 | 0,465696 | Q96FW1;F5 | OTUB1 |
| 8,43105 | 5,3393 | -3,87175 | 5,3393 | 6,4000163 | 0,466104 | H3BLX0;P4( | PBX3 |
| 0,232816 | -0,598409 | -0,283397 | -0,283397 | 0,4196513 | 0,466143 | Q9UHB9 | SRP68 |
| 1,30348 | -3,02188 | -1,71033 | -1,71033 | 2,217805 | 0,466265 | A0A2R8Y5F | CNKSR2 |
| 0,281793 | 0,776315 | -0,258476 | 0,281793 | 0,517564 | 0,466514 | P49961;A0/ | ENTPD1 |
| 2,03348 | -3,23634 | -3,72885 | -3,23634 | 3,1942139 | 0,466772 | P17858 | PFKL |
| -1,03567 | 1,21658 | 2,77652 | 1,21658 | 1,9165436 | 0,46698 | Q9UKX2;J3( | MYH2 |
| -3,03851 | 0,974824 | -1,03036 | -1,03036 | 2,0066672 | 0,467282 | Q9P2G1 | ANKIB1 |
| 2,84156 | 0,92124 | -0,888383 | 0,92124 | 1,8652453 | 0,467491 | Q9H0S4;F5 | DDX47 |
| -0,768092 | 1,49317 | 1,14591 | 1,14591 | 1,2177369 | 0,46863 | Q96SZ5 | ADO |
| -1,02781 | 1,4203 | 2,18635 | 1,4203 | 1,6788339 | 0,468719 | Q9C0F0;A0 | ASXL3 |
| 4,08801 | -0,400421 | 0,0887877 | 0,0887877 | 2,4623538 | 0,469326 | O94956;A0 | SLCO2B1 |
| -0,197312 | 0,0797159 | 1,61008 | 0,0797159 | 0,9734325 | 0,469432 | Q13243;B4 | SRSF5 |
| -0,875471 | 0,645217 | -1,39279 | -0,875471 | 1,0593696 | 0,469714 | E9PQY2;Q9 | PFDN4 |
| -0,768829 | 0,522639 | -1,02345 | -0,768829 | 0,8289665 | 0,469836 | O14497;A0 | ARID1A |
| -0,105728 | -1,99376 | 0,252376 | -0,105728 | 1,2067885 | 0,470083 | O14964;I3L | HGS |
| -2,06811 | 0,283625 | -0,134377 | -0,134377 | 1,2546385 | 0,470379 | E9PRY8 | EEF1D |
| -2,0681 | 0,283627 | -0,134347 | -0,134347 | 1,2546396 | 0,470387 | P29692;E9F | EEF1D |
| 1,48332 | -0,975747 | 1,83129 | 1,48332 | 1,5301171 | 0,470594 | Q96C19 | EFHD2 |
| -1,65998 | 1,04969 | -1,87715 | -1,65998 | 1,6307394 | 0,471396 | D6RDH9;E2 | SNX14 |
| -2,05701 | 1,32455 | -2,41292 | -2,05701 | 2,0627773 | 0,471522 | Q5TAX3;A0 | TUT4 |
| -1,57245 | -3,06165 | 1,27619 | -1,57245 | 2,2041371 | 0,471865 | Q9UPU5 | USP24 |
| 0,623378 | 0,0996727 | -0,132986 | 0,0996727 | 0,3874024 | 0,471947 | Q92804;A0 | TAF15 |
| 0,773321 | -1,01542 | -1,70413 | -1,01542 | 1,2787806 | 0,472245 | Q96CN7;D6 | ISOC1 |
| -0,997784 | 0,541119 | -0,825722 | -0,825722 | 0,8432163 | 0,472523 | P30153;B3I | PPP2R1A |
| -2,46297 | 0,0451504 | 0,163084 | 0,0451504 | 1,483281 | 0,472705 | P17980;R4( | PSMC3 |
| -1,18148 | 0,962758 | -2,2855 | -1,18148 | 1,6516556 | 0,473686 | Q8IVF4;A0/ | DNAH10 |
| 0,743305 | 2,1459 | -0,719271 | 0,743305 | 1,4326901 | 0,474088 | Q99622;U3 | C12orf57 |
| -0,221596 | 0,624036 | 0,238251 | 0,238251 | 0,4233562 | 0,474396 | Q5THJ4;H3 | VPS13D |
| 2,23928 | -3,54122 | -3,89847 | -3,54122 | 3,4451363 | 0,47537 | H0YGI1;Q8I | C12orf56 |
| 0,558288 | 0,714636 | -0,380564 | 0,558288 | 0,5923612 | 0,476138 | P62191;Q5 | PSMC1 |
| 1,45804 | 0,591641 | -0,54026 | 0,591641 | 1,0020853 | 0,47618 | Q6IQ26;H0 | DENND5A |
| -0,780585 | -0,583485 | 0,40665 | -0,583485 | 0,6362314 | 0,476546 | P35268;K7E | RPL22 |
| 1,19225 | -2,1531 | -1,80505 | -1,80505 | 1,8392169 | 0,476793 | A0A2R8YD5 | HSD17B4 |
| -0,49827 | 2,08024 | 0,38768 | 0,38768 | 1,3101133 | 0,476902 | Q9Y696 | CLIC4 |
| 0,42301 | -1,0227 | -0,500884 | -0,500884 | 0,7321145 | 0,476937 | Q9BYX7 | POTEKP |
| 0,213789 | 1,08048 | -0,2675 | 0,213789 | 0,6831108 | 0,476988 | P56192;H0Y | MARS |
| 1,63563 | -0,854152 | 1,21846 | 1,21846 | 1,3334645 | 0,477816 | E5RG77;O9 | PLPBP |
| -2,68135 | 0,347588 | -0,113604 | -0,113604 | 1,6319969 | 0,477865 | Q9ULU8;F1 | CADPS |
| -2,96672 | 0,667991 | -0,481925 | -0,481925 | 1,8577602 | 0,478584 | P46821;D6I | MAP1B |
| -0,265535 | 0,286933 | -0,940266 | -0,265535 | 0,6146137 | 0,479026 | P48449;A0/ | LSS |
| -2,41655 | 0,168558 | 0,0627182 | 0,0627182 | 1,462917 | 0,479345 | O60488 | ACSL4 |
| -2,49051 | 1,59712 | -2,7446 | -2,49051 | 2,436658 | 0,479537 | P00352 | ALDH1A1 |
| 0,399875 | -2,39179 | -0,199886 | -0,199886 | 1,4695549 | 0,479931 | O75582 | RPS6KA5 |
| 0,877902 | -2,41161 | -0,922041 | -0,922041 | 1,6471946 | 0,480085 | Q99733;C9 | NAP1L4 |
| 2,81743 | -0,377604 | 0,119706 | 0,119706 | 1,7191701 | 0,480607 | A0A087WU | PTBP1 |

| | | | | | | | |
|---|---|---|---|---|---|---|---|
| 1,38287 | -1,46588 | 4,60326 | 1,38287 | 3,0364658 | 0,480649 | Q14940;J3K | SLC9A5 |
| 1,80747 | 1,83718 | -1,11881 | 1,80747 | 1,6981301 | 0,480962 | Q9UJS0 | SLC25A13 |
| 1,76357 | -0,753596 | 0,889495 | 0,889495 | 1,2780115 | 0,481257 | P46783 | RPS10 |
| -1,09708 | -0,147745 | 0,229565 | -0,147745 | 0,6835674 | 0,481521 | O00267 | SUPT5H |
| -0,09782 | 1,71143 | -0,0728725 | -0,072873 | 1,0374442 | 0,481548 | Q96H40 | ZNF486 |
| -2,02572 | 0,578586 | -0,495743 | -0,495743 | 1,3087795 | 0,481709 | P83731;C9J | RPL24 |
| 0,59783 | -0,727165 | 2,65766 | 0,59783 | 1,7056549 | 0,482265 | P07602;C9J | PSAP |
| -2,40358 | 5,50492 | 2,86649 | 2,86649 | 4,0265644 | 0,482317 | Q96M27 | PRRC1 |
| 0,203574 | -0,73462 | -0,167704 | -0,167704 | 0,4724844 | 0,483145 | P60660;F8V | MYL6 |
| -0,318282 | 0,510313 | -2,44169 | -0,318282 | 1,5225939 | 0,483495 | A0A0G2JL6 | C2 |
| 3,02955 | -0,928173 | 0,827911 | 0,827911 | 1,9830371 | 0,483583 | P13929;E5F | ENO3 |
| 0,467117 | -1,47662 | -0,426156 | -0,426156 | 0,9729273 | 0,483985 | P20807;F8V | CAPN3 |
| -1,07644 | 0,794263 | -1,53284 | -1,07644 | 1,2331017 | 0,484934 | O75531 | BANF1 |
| -0,449009 | -2,63976 | 0,634142 | -0,449009 | 1,6678849 | 0,484988 | E9PJK2;O75 | EED |
| -1,09866 | 1,92351 | 1,62279 | 1,62279 | 1,664844 | 0,485375 | Q15928 | ZNF141 |
| 0,914655 | -1,66403 | -1,30217 | -1,30217 | 1,396118 | 0,485563 | O95197;F5I | RTN3 |
| -0,334142 | -0,968803 | 0,340948 | -0,334142 | 0,6549795 | 0,485748 | P33993 | MCM7 |
| -2,2841 | 5,55076 | 2,53796 | 2,53796 | 3,9520935 | 0,485748 | P49593;B5M | PPM1F |
| -0,973345 | 0,298271 | -0,260638 | -0,260638 | 0,6373562 | 0,485911 | Q99497;K7I | PARK7 |
| 1,30988 | -0,105803 | -0,0349138 | -0,034914 | 0,7976689 | 0,486527 | F8WF69 | CLTA |
| 0,218747 | -0,298213 | -0,413485 | -0,298213 | 0,3367127 | 0,486968 | J3QRC4;J3C | RPL26 |
| -0,928998 | 1,28595 | 1,72505 | 1,28595 | 1,4226017 | 0,487097 | A0A1W2PP | POLR2B |
| 1,90505 | 2,35899 | -1,32502 | 1,90505 | 2,0087867 | 0,487208 | Q12972;A0 | PPP1R8 |
| 0,991692 | -0,986047 | 2,66797 | 0,991692 | 1,8290799 | 0,487559 | P40939;H0Y | HADHA |
| 0,10404 | -1,65135 | 0,0772102 | 0,0772102 | 1,0058193 | 0,487593 | Q7RTP6;E9I | MICAL3 |
| 6,26019 | 6,24565 | -3,92395 | 6,24565 | 5,8756231 | 0,487852 | Q9C0E8;C9 | LNPK |
| -2,16322 | -2,27037 | 1,39115 | -2,16322 | 2,0837369 | 0,487984 | K7ES00 | H3F3B |
| -0,757702 | -0,0679929 | 0,140027 | -0,067993 | 0,4699093 | 0,488226 | P35221;G3 | CTNNA1 |
| -0,654599 | 0,571413 | -1,30956 | -0,654599 | 0,9548245 | 0,488358 | Q5VUA4 | ZNF318 |
| -1,44074 | 3,11572 | 1,73016 | 1,73016 | 2,3357979 | 0,488575 | O75368 | SH3BGRL |
| 2,72167 | -0,990203 | 0,975525 | 0,975525 | 1,8570187 | 0,488599 | P49902 | NT5C2 |
| -1,60557 | -0,380443 | 0,468527 | -0,380443 | 1,042718 | 0,489219 | Q96BW9;A0 | TAMM41 |
| 0,765711 | -0,53403 | 0,938476 | 0,765711 | 0,8049274 | 0,489628 | P20073 | ANXA7 |
| -0,688514 | 0,8126 | -2,63399 | -0,688514 | 1,7280626 | 0,489967 | Q6NUN0;I3 | ACSM5 |
| -0,33781 | 1,87896 | 0,150427 | 0,150427 | 1,1647797 | 0,49001 | Q8NCX0;E9 | CCDC150 |
| -1,31163 | -0,725611 | 0,610657 | -0,725611 | 0,9852426 | 0,491134 | P61758;B4I | VBP1 |
| 1,00114 | -0,788388 | 1,56774 | 1,00114 | 1,2298228 | 0,491183 | Q6ZNG1 | ZNF600 |
| -1,65222 | -0,427803 | 0,510245 | -0,427803 | 1,0843881 | 0,491221 | P52272;A0 | HNRNPM |
| -1,04913 | 0,639851 | -0,970546 | -0,970546 | 0,9532585 | 0,491257 | Q9NY46;A0 | SCN3A |
| 2,52069 | 0,615744 | -0,755683 | 0,615744 | 1,6454103 | 0,491407 | O15516 | CLOCK |
| 0,294576 | -0,451705 | -0,476082 | -0,451705 | 0,4380722 | 0,491786 | P05062;A0 | ALDOB |
| 7,80334 | -3,21942 | 3,43246 | 3,43246 | 5,5505756 | 0,492101 | Q13011;M0 | ECH1 |
| -0,589793 | 0,695222 | 1,26777 | 0,695222 | 0,9512812 | 0,492287 | F8W8Y7 | ARMCX4 |
| -1,49866 | -2,11053 | 1,12774 | -1,49866 | 1,7204047 | 0,49259 | P29597;E9F | TYK2 |
| 1,01269 | -0,493189 | 0,600999 | 0,600999 | 0,7782896 | 0,493289 | Q92900 | UPF1 |
| -0,879006 | 0,896286 | -2,37258 | -0,879006 | 1,636455 | 0,4934 | Q8TD26;C9 | CHD6 |
| -2,05243 | 0,67133 | -0,580678 | -0,580678 | 1,3633566 | 0,493488 | O60282;A0 | KIF5C |
| -1,02978 | 2,04508 | 1,2892 | 1,2892 | 1,6022789 | 0,493664 | Q5T1J5;Q9 | CHCHD2P9 |
| 0,73612 | 1,27862 | -0,614676 | 0,73612 | 0,9749808 | 0,494259 | B7Z637;Q8 | ARMC8 |
| 0,277678 | 1,31886 | -0,373739 | 0,277678 | 0,8537462 | 0,495233 | Q9H0B6;A8 | KLC2 |
| -0,825986 | -1,42158 | 0,690622 | -0,825986 | 1,0890538 | 0,495929 | Q709C8;A0 | VPS13C |
| 2,0018 | -0,73431 | 0,68722 | 0,68722 | 1,3684033 | 0,496237 | P31040;D6I | SDHA |

| | | | | | | | |
|---|---|---|---|---|---|---|---|
| 5,89495 | 3,81045 | -3,03591 | 3,81045 | 4,6722237 | 0,496496 | Q9H7P9;E7 | PLEKHG2 |
| 1,15107 | 2,07455 | -0,986405 | 1,15107 | 1,5700881 | 0,496839 | Q13488 | TCIRG1 |
| -0,169203 | 0,581076 | -3,54374 | -0,169203 | 2,1971393 | 0,497035 | P37059 | HSD17B2 |
| 0,582437 | -0,614045 | -1,37115 | -0,614045 | 0,984994 | 0,497378 | Q9Y6G9;E9 | DYNC1LI1 |
| 0,493954 | -1,05217 | -0,566947 | -0,566947 | 0,7907224 | 0,497687 | P61964;V9( | WDR5 |
| -0,383728 | 0,169638 | 1,93531 | 0,169638 | 1,2111805 | 0,498175 | P58876;P6z | HIST1H2BD |
| -0,289253 | 0,324741 | -0,915604 | -0,289253 | 0,6201828 | 0,4987 | P62873;B3I | GNB1 |
| 1,90926 | 1,10363 | -0,933872 | 1,10363 | 1,4653699 | 0,498794 | Q5VTH9;H0 | WDR78 |
| -0,092142 | -0,92393 | 0,193578 | -0,092142 | 0,5805646 | 0,499337 | Q9HC56;B7 | PCDH9 |
| 1,06126 | -1,38738 | -1,93332 | -1,38738 | 1,5948562 | 0,499341 | Q01484;I6L | ANK2 |
| 0,83833 | -1,11537 | -1,49783 | -1,11537 | 1,2530538 | 0,499411 | Q14324 | MYBPC2 |
| -0,861465 | 1,1052 | 1,59709 | 1,1052 | 1,3009112 | 0,499783 | P35613;A0/ | BSG |
| -1,46759 | 2,04962 | 2,48213 | 2,04962 | 2,1663379 | 0,499939 | P14923;C9J | JUP |
| 2,41063 | 1,93608 | -1,40642 | 1,93608 | 2,0803595 | 0,500227 | Q9BU68 | PRR15L |
| 0,0397985 | 0,103137 | -1,09894 | 0,0397985 | 0,6764769 | 0,500266 | Q8WYL5;F8 | SSH1 |
| -1,12251 | -1,24803 | 0,77269 | -1,12251 | 1,1321696 | 0,50077 | P11217 | PYGM |
| -1,01449 | 0,19887 | -0,0809077 | -0,080908 | 0,63536 | 0,500838 | P80723 | BASP1 |
| 3,57935 | 9,65174 | -3,75735 | 3,57935 | 6,7144717 | 0,500867 | H7BXF4;Q9 | SMPD4 |
| -0,37187 | 0,509035 | 0,637791 | 0,509035 | 0,5495433 | 0,50107 | P84077;F5I | ARF1 |
| -1,24685 | 2,04366 | 1,76549 | 1,76549 | 1,8247843 | 0,502672 | G3V320 | TMEM260 |
| -0,831182 | -1,25691 | 0,670177 | -0,831182 | 1,0123395 | 0,503614 | P33176 | KIF5B |
| -2,23178 | 3,44874 | 3,3243 | 3,3243 | 3,2443237 | 0,503848 | Q8NFA0;K7 | USP32 |
| -0,724184 | 0,955975 | 1,26799 | 0,955975 | 1,0715289 | 0,503872 | Q13347;Q5 | EIF3I |
| -0,251065 | 0,425942 | -1,67821 | -0,251065 | 1,0741304 | 0,503895 | P62993 | GRB2 |
| -1,32804 | 0,941837 | -1,53863 | -1,32804 | 1,3753428 | 0,503909 | A1L4H1 | SSC5D |
| 1,34216 | -1,54254 | -2,72717 | -1,54254 | 2,0930157 | 0,504125 | O75122;J3I | CLASP2 |
| -1,25295 | 1,27892 | 2,89379 | 1,27892 | 2,0902003 | 0,504617 | P16157 | ANK1 |
| -0,279307 | 0,195504 | 0,945637 | 0,195504 | 0,6176073 | 0,505003 | Q8IXJ9;A0A | ASXL1 |
| 0,00882835 | -1,00195 | 0,12719 | 0,0088284 | 0,6205696 | 0,505019 | A0A087WV | CACNA1A |
| -0,936701 | 1,92445 | 1,05945 | 1,05945 | 1,4673689 | 0,50508 | Q9ULM2 | ZNF490 |
| -0,528174 | -1,21316 | 0,522133 | -0,528174 | 0,8740321 | 0,505144 | A0A087WU | RNPEP |
| 1,0193 | -1,31134 | -1,81787 | -1,31134 | 1,5131639 | 0,505289 | Q8N157 | AHI1 |
| 1,25983 | -1,74019 | -2,08032 | -1,74019 | 1,8381336 | 0,505634 | Q8ND30;E9 | PPFIBP2 |
| -0,263072 | 5,2032 | -0,463916 | -0,263072 | 3,2155008 | 0,505905 | P54819;F8\ | AK2 |
| 1,39822 | -1,01498 | 1,67501 | 1,39822 | 1,4796505 | 0,506182 | P15927;Q5 | RPA2 |
| -1,19542 | -1,74816 | 0,956324 | -1,19542 | 1,4288549 | 0,506246 | P21817 | RYR1 |
| -0,109294 | -0,295458 | 2,89405 | -0,109294 | 1,790144 | 0,506312 | A0A087WV | RUNX1T1 |
| -2,09913 | -0,0244126 | 0,310104 | -0,024413 | 1,3051668 | 0,506615 | P27815 | PDE4A |
| -1,57246 | -2,90844 | 1,41241 | -1,57246 | 2,21224 | 0,507254 | O75445 | USH2A |
| -0,932107 | -1,48549 | 0,780979 | -0,932107 | 1,1816498 | 0,507799 | Q15366;F8 | PCBP2 |
| -0,144709 | -2,33922 | 0,450742 | -0,144709 | 1,4693722 | 0,508153 | A0A075B6I | KAT14 |
| -1,05108 | 2,85583 | 0,893677 | 0,893677 | 1,9534615 | 0,508788 | Q9Y2T4 | PPP2R2C |
| 1,73154 | -1,78122 | -3,84411 | -1,78122 | 2,8190681 | 0,508822 | Q6P2D8 | XRRA1 |
| 0,859682 | -1,06995 | -1,5491 | -1,06995 | 1,275101 | 0,509213 | P23468;F5( | PTPRD |
| -3,21105 | 0,203898 | 0,263693 | 0,203898 | 1,9891072 | 0,50936 | Q05932 | FPGS |
| -2,34403 | 1,18218 | -1,33802 | -1,33802 | 1,8164811 | 0,510178 | Q9ULL4 | PLXNB3 |
| 0,7495 | -1,81938 | -0,701735 | -0,701735 | 1,2880449 | 0,510391 | Q14254;E7I | FLOT2 |
| 1,06532 | -0,0639801 | -0,0931192 | -0,06398 | 0,6605741 | 0,510535 | Q4G0P3;F8 | HYDIN |
| 0,123491 | 0,248749 | -2,49249 | 0,123491 | 1,5477638 | 0,511894 | Q9Y4G6 | TLN2 |
| -1,46313 | 3,56319 | 1,34877 | 1,34877 | 2,5190716 | 0,51211 | F8WBA3;Q: | PRKD1 |
| -0,500435 | -1,84905 | 0,64348 | -0,500435 | 1,2476651 | 0,512579 | H0YNL8;P4 | IREB2 |
| -1,16417 | -0,267685 | 0,374923 | -0,267685 | 0,7730284 | 0,512605 | Q12905;B4 | ILF2 |

| | | | | | | | |
|---|---|---|---|---|---|---|---|
| -2,963 | 1,30653 | -1,28345 | -1,28345 | 2,1508824 | 0,512719 | Q7L311 | ARMCX2 |
| -1,64916 | 3,94819 | 1,5284 | 1,5284 | 2,8072109 | 0,513647 | P01266;E7E | TG |
| 3,62625 | -2,7553 | 4,55449 | 3,62625 | 3,9795067 | 0,513671 | O43572;E7I | AKAP10 |
| 3,66771 | 0,0974618 | -0,631854 | 0,0974618 | 2,3008993 | 0,514098 | Q9BXS5;K7I | AP1M1 |
| -1,1019 | 1,19246 | -2,85212 | -1,1019 | 2,0283813 | 0,514184 | P26583;D6I | HMGB2 |
| 1,89518 | 0,714574 | -0,782837 | 0,714574 | 1,342128 | 0,514253 | Q49AJ0;J3C | FAM135B |
| 2,10624 | -3,11986 | -3,05922 | -3,05922 | 2,9999382 | 0,515227 | A0A0G2JH6 | DIAPH1 |
| 2,22992 | -3,0155 | -3,54734 | -3,0155 | 3,1930659 | 0,515409 | Q9H257;A0 | CARD9 |
| 0,7049 | -0,226974 | 0,157274 | 0,157274 | 0,4683179 | 0,515581 | Q9P0L0;J3C | VAPA |
| 4,76676 | -0,442412 | -0,300376 | -0,300376 | 2,9673646 | 0,515654 | O43283 | MAP3K13 |
| -0,756452 | 0,798833 | 1,55687 | 0,798833 | 1,1793353 | 0,515659 | A2A288;H0 | ZC3H12D |
| -1,63546 | 1,29191 | -2,19618 | -1,63546 | 1,8730845 | 0,5157 | Q12851;C9. | MAP4K2 |
| 0,283683 | -0,186149 | -0,912042 | -0,186149 | 0,6024147 | 0,51675 | P46937;H0Y | YAP1 |
| -0,838902 | 1,11019 | 1,34236 | 1,11019 | 1,1979682 | 0,518146 | Q5VYK3;J3I | ECPAS |
| 1,18395 | -1,44949 | -2,05253 | -1,44949 | 1,7211172 | 0,518186 | Q3V6T2;A0 | CCDC88A |
| -0,32478 | -1,28699 | 0,443976 | -0,32478 | 0,8672828 | 0,518279 | A8MUS3;H7 | RPL23A |
| -1,30511 | 2,39188 | 1,51369 | 1,51369 | 1,9315187 | 0,518327 | Q96AX9;D6 | MIB2 |
| -0,396498 | 0,601526 | -1,88397 | -0,396498 | 1,2507541 | 0,519422 | Q8N8L6 | ARL10 |
| 0,672046 | 0,754297 | -0,492269 | 0,672046 | 0,6971754 | 0,520124 | A0A087WV | DCAF7 |
| 1,01455 | -1,52644 | -1,40632 | -1,40632 | 1,4336242 | 0,520615 | A0A0A0MR | ARHGAP30 |
| 1,50891 | 0,0246711 | -0,262156 | 0,0246711 | 0,9506059 | 0,520761 | Q9P225;H7 | DNAH2 |
| -0,69297 | 0,578459 | 1,73293 | 0,578459 | 1,2134198 | 0,521788 | Q7Z6Z7;A0 | HUWE1 |
| 0,636595 | -1,69172 | -0,495835 | -0,495835 | 1,1643016 | 0,522251 | P61289;K9J | PSME3 |
| -1,73557 | 3,7211 | 1,68155 | 1,68155 | 2,757164 | 0,522826 | F8WEH6;P5 | ETV1 |
| -1,83077 | -1,6345 | 1,20598 | -1,6345 | 1,6994459 | 0,52299 | B5MD58;P3 | SREBF1 |
| -0,770794 | -2,9775 | 1,06169 | -0,770794 | 2,0224822 | 0,523283 | P06744;A0 | GPI |
| 1,34491 | 3,01317 | -1,39983 | 1,34491 | 2,228275 | 0,523497 | O75781;A0 | PALM |
| 0,17747 | -0,196359 | -0,329687 | -0,196359 | 0,2629109 | 0,523987 | P0CJ79 | ZNF888 |
| -0,31909 | -0,103141 | 0,126762 | -0,103141 | 0,2229624 | 0,524164 | B1AJY5;B1A | PSMD10 |
| -0,204291 | 0,328708 | -1,03358 | -0,204291 | 0,6864931 | 0,524397 | Q8NCB2;E7 | CAMKV |
| -1,44881 | -0,236806 | 0,427234 | -0,236806 | 0,9512662 | 0,524817 | A0A0A0MT | DBI |
| 0,469582 | 1,57698 | -0,605755 | 0,469582 | 1,0914067 | 0,525572 | Q9UG01;H( | IFT172 |
| -0,752337 | 0,796493 | 1,44668 | 0,796493 | 1,1296971 | 0,525699 | O00299 | CLIC1 |
| 1,58382 | -0,15994 | -0,113048 | -0,113048 | 0,9935004 | 0,525774 | O15355 | PPM1G |
| -0,521431 | 2,56565 | 0,107947 | 0,107947 | 1,6312822 | 0,525802 | P48741 | HSPA7 |
| -0,725859 | 0,146434 | 3,58901 | 0,146434 | 2,2814586 | 0,525846 | P35052;H7( | GPC1 |
| -2,61568 | -0,318754 | 0,698038 | -0,318754 | 1,6975699 | 0,526331 | C9JJQ8 | TUBA4A |
| 2,39981 | 3,41817 | -2,01743 | 2,39981 | 2,8894871 | 0,52692 | E2QRD4;Q6 | MMS22L |
| 0,872005 | -0,568443 | -2,58921 | -0,568443 | 1,7386968 | 0,527123 | Q8TAT2 | FGFBP3 |
| -1,13687 | 0,608559 | -0,656662 | -0,656662 | 0,9016565 | 0,527222 | P61006 | RAB8A |
| -0,205579 | 0 | 1,20725 | 0 | 0,7633043 | 0,527763 | P55196;J3K | AFDN |
| -0,976125 | -0,707909 | 0,587189 | -0,707909 | 0,8359794 | 0,527829 | P55072;C9I | VCP |
| 0,745837 | -0,320165 | 0,272505 | 0,272505 | 0,5341132 | 0,529194 | Q10571;H7 | MN1 |
| 0,659346 | -0,519911 | 0,81397 | 0,659346 | 0,7295883 | 0,529306 | A2IDC6;Q1 | MRPL28 |
| 0,590671 | -0,880776 | -0,784238 | -0,784238 | 0,8230888 | 0,529731 | Q9NR45;Q5 | NANS |
| -1,95048 | 1,07685 | -1,17762 | -1,17762 | 1,5729312 | 0,530058 | P28070 | PSMB4 |
| 0,000806121 | 0,491997 | -2,81391 | 0,0008061 | 1,7838589 | 0,530879 | Q15052 | ARHGEF6 |
| -2,45672 | -0,862464 | 1,04199 | -0,862464 | 1,7516454 | 0,531199 | A0A0U1RR | DOCK7 |
| 0,74435 | -1,47242 | -0,740152 | -0,740152 | 1,1294565 | 0,531226 | M0QXL5;M | FBL |
| -2,40481 | 0,662997 | -0,297835 | -0,297835 | 1,5691814 | 0,53126 | Q9H936;E9 | SLC25A22 |
| 0,898712 | -2,82624 | -0,516178 | -0,516178 | 1,8803177 | 0,531314 | M0QY97;Q | ZC3H4 |
| 0,6226 | -0,771494 | -0,98575 | -0,771494 | 0,8733263 | 0,531427 | Q08495 | DMTN |

| | | | | | | | |
|---|---|---|---|---|---|---|---|
| -2,53294 | 1,26211 | -1,23401 | -1,23401 | 1,9287405 | 0,531576 | P62258;B4I | YWHAE |
| 0,38979 | -0,056043 | -1,93358 | -0,056043 | 1,2330153 | 0,531913 | Q15691 | MAPRE1 |
| 0,0408646 | -1,90306 | 0,300691 | 0,0408646 | 1,2043581 | 0,532181 | Q9UKG1 | APPL1 |
| -1,79043 | 2,00899 | 3,11822 | 2,00899 | 2,5742581 | 0,532275 | Q86UW7;A | CADPS2 |
| -0,153043 | 0,0748221 | 0,524456 | 0,0748221 | 0,3447458 | 0,532789 | Q15084 | PDIA6 |
| 0,317125 | -0,012614 | -1,71429 | -0,012614 | 1,0901893 | 0,533138 | P53396;K7E | ACLY |
| -0,560946 | 0,340428 | 1,67366 | 0,340428 | 1,1242365 | 0,533306 | Q16352 | INA |
| 6,18381 | -1,99402 | 1,14238 | 1,14238 | 4,1257307 | 0,533345 | Q86UU1;A0 | PHLDB1 |
| -0,309523 | 0,729278 | -2,66522 | -0,309523 | 1,7393022 | 0,533742 | F5H0R1;Q9 | TUT1 |
| 0,224982 | -1,44838 | 0,0408185 | 0,0408185 | 0,9175846 | 0,534369 | O43237;J3K | DYNC1LI2 |
| -1,74544 | -0,185616 | 0,466522 | -0,185616 | 1,1365966 | 0,534446 | O00264 | PGRMC1 |
| 0,358219 | -0,56047 | 1,59164 | 0,358219 | 1,0798838 | 0,534989 | P16402 | HIST1H1D |
| 1,13778 | -0,885487 | -2,74964 | -0,885487 | 1,9442526 | 0,535594 | Q14558;B4 | PRPSAP1 |
| -1,70337 | 1,03121 | -1,19063 | -1,19063 | 1,453582 | 0,536431 | P61353;K7E | RPL27 |
| 0,268009 | 0,118542 | -2,04469 | 0,118542 | 1,2942494 | 0,536535 | O00222;A0 | GRM8 |
| 1,36578 | -0,716391 | 0,710661 | 0,710661 | 1,064667 | 0,537592 | Q9H2M9 | RAB3GAP2 |
| -1,1931 | -0,213895 | 0,388227 | -0,213895 | 0,7981216 | 0,537871 | P11940;E7E | PABPC1 |
| 0,343859 | -2,06457 | 0,0454808 | 0,0454808 | 1,3128769 | 0,538002 | Q6UB98;F5 | ANKRD12 |
| 1,47009 | 0,0780253 | -0,339711 | 0,0780253 | 0,9476046 | 0,538224 | F8VRK6 | ATXN2 |
| -0,494022 | 1,7198 | 0,21481 | 0,21481 | 1,1305195 | 0,538494 | P29317 | EPHA2 |
| -1,44138 | -0,672722 | 0,719192 | -0,672722 | 1,095166 | 0,538658 | O14949 | UQCRQ |
| 1,16399 | 0,52754 | -0,572821 | 0,52754 | 0,8786709 | 0,538804 | Q3KP31 | ZNF791 |
| -1,22843 | 0,741989 | -0,84074 | -0,84074 | 1,0438619 | 0,539309 | P23634;H7I | ATP2B4 |
| 3,27389 | -0,518602 | -0,10918 | -0,10918 | 2,0814972 | 0,539356 | Q13033;G3 | STRN3 |
| 0,663458 | -0,0848202 | -3,1399 | -0,08482 | 2,0149014 | 0,539382 | P35527;K7E | KRT9 |
| -2,55428 | 1,54191 | -1,74045 | -1,74045 | 2,168524 | 0,539872 | A0A2R8YF4 | ARHGEF5 |
| -0,527977 | 2,04738 | 0,17107 | 0,17107 | 1,3317701 | 0,539901 | J3QLH3;J3C | SAP30BP |
| -1,6809 | 1,35589 | -2,03157 | -1,6809 | 1,8627913 | 0,541119 | Q8WU90;H | ZC3H15 |
| -2,47312 | 3,47521 | 3,26286 | 3,26286 | 3,3746404 | 0,54148 | Q05084 | ICA1 |
| 1,39133 | -3,2186 | -1,08441 | -1,08441 | 2,3070728 | 0,541983 | Q2KHM9;F6 | KIAA0753 |
| 1,10214 | -1,86479 | -1,20205 | -1,20205 | 1,5573048 | 0,542116 | P54136;E5F | RARS |
| 1,19904 | -0,438541 | 0,274327 | 0,274327 | 0,8210711 | 0,542479 | P53999;D6I | SUB1 |
| 1,58519 | -3,53999 | -1,28184 | -1,28184 | 2,5686109 | 0,542555 | E9PPP6 | RRP8 |
| 1,35884 | -0,899898 | -3,56241 | -0,899898 | 2,4633842 | 0,542624 | A2A3N6 | PIPSL |
| 1,20497 | -1,55307 | 3,58976 | 1,20497 | 2,5736714 | 0,542707 | Q7Z7M1;A0 | ADGRD2 |
| 0,877272 | 1,48381 | -0,84452 | 0,877272 | 1,2078615 | 0,543848 | P22307;H0Y | SCP2 |
| 0,264481 | -0,603729 | 2,01114 | 0,264481 | 1,3317999 | 0,543909 | Q99259 | GAD1 |
| 0,786409 | -2,18554 | -0,473244 | -0,473244 | 1,4917084 | 0,54396 | Q06203;D6 | PPAT |
| -3,02171 | 4,43272 | 3,74403 | 3,74403 | 4,1194267 | 0,545052 | A0A1B0GW | CABP2 |
| -0,236992 | -0,18443 | 2,06475 | -0,18443 | 1,3140009 | 0,545273 | Q9UIJ7 | AK3 |
| -2,40275 | -1,97491 | 1,61799 | -1,97491 | 2,2082545 | 0,545539 | Q96KK5;P0 | HIST1H2AH |
| -1,41143 | 0,327174 | -0,0557595 | -0,05576 | 0,9135302 | 0,546054 | Q13435;E9I | SF3B2 |
| 2,17827 | -0,465022 | 0,0368212 | 0,0368212 | 1,4038425 | 0,546428 | Q9UBS4;H7 | DNAJB11 |
| -0,378922 | 1,07393 | 0,215133 | 0,215133 | 0,7304351 | 0,546602 | Q02641 | CACNB1 |
| 0,897948 | -0,936031 | -1,54685 | -0,936031 | 1,2723742 | 0,546711 | Q9Y4F9;F5 | RIPOR2 |
| -0,374271 | 0,667041 | -1,90299 | -0,374271 | 1,2926956 | 0,546717 | Q9ULM0 | PLEKHH1 |
| 1,95919 | 0,189046 | -0,544956 | 0,189046 | 1,2873044 | 0,546767 | P46734;Q6I | MAP2K3 |
| 2,37074 | -3,44233 | -2,93294 | -2,93294 | 3,2192205 | 0,547204 | O43824 | GTPBP6 |
| 0,128345 | -0,149148 | 0,305578 | 0,128345 | 0,2291977 | 0,547626 | P30154;H0Y | PPP2R1B |
| 1,57528 | 3,20695 | -1,68667 | 1,57528 | 2,4916588 | 0,54766 | Q02880;E9I | TOP2B |
| -0,969115 | 1,44192 | -3,60927 | -0,969115 | 2,5264609 | 0,547928 | Q9BZ95 | NSD3 |
| 0,340071 | -0,480593 | 1,15346 | 0,340071 | 0,8170292 | 0,54841 | Q6V0I7 | FAT4 |

| | | | | | | | |
|---|---|---|---|---|---|---|---|
| 0,449943 | -0,828578 | -0,432466 | -0,432466 | 0,654493 | 0,548555 | P09496;C9J | CLTA |
| -2,8586 | 1,99335 | -2,47818 | -2,47818 | 2,6981697 | 0,548594 | A0A087WU | OSTC |
| 5,28966 | 0,274633 | -1,30026 | 0,274633 | 3,4413619 | 0,548621 | Q96AE7 | TTC17 |
| 1,25688 | -0,84123 | -3,12876 | -0,84123 | 2,1935017 | 0,549251 | Q9NVE5 | USP40 |
| 2,05455 | -1,50638 | 1,96079 | 1,96079 | 2,0293793 | 0,549415 | D6RCF1;P29 | CNGA1 |
| -0,97276 | 1,59573 | 1,04989 | 1,04989 | 1,3531565 | 0,54943 | Q8WXF7 | ATL1 |
| -3,16555 | 4,09416 | 4,3426 | 4,09416 | 4,2649234 | 0,549524 | Q9UEW8 | STK39 |
| -1,34119 | 0,316464 | -0,0508833 | -0,050883 | 0,8705987 | 0,549658 | Q14141;B1 | SEPT6 |
| 0,721697 | -2,79777 | -0,182011 | -0,182011 | 1,8278189 | 0,549682 | Q9UH03 | SEPT3 |
| 0,721708 | -2,79777 | -0,182 | -0,182 | 1,827825 | 0,549687 | A0A2R8Y4H | SEPT3 |
| 1,87581 | -2,29516 | -2,71287 | -2,29516 | 2,5373036 | 0,549954 | O60462 | NRP2 |
| 0,207701 | -2,18126 | 0,255397 | 0,207701 | 1,39324 | 0,550318 | A1L0T0;E9F | ILVBL |
| 1,96606 | -3,55772 | -1,89234 | -1,89234 | 2,8335163 | 0,551379 | O60318 | MCM3AP |
| -0,323493 | 0,124761 | -0,0765687 | -0,076569 | 0,2245131 | 0,552358 | P23381;H0Y | WARS |
| -1,92202 | 1,22454 | -1,35939 | -1,35939 | 1,6779991 | 0,552484 | O43427;H0 | FIBP |
| -0,511488 | -2,22211 | 0,848842 | -0,511488 | 1,5388021 | 0,552765 | Q9UIG0 | BAZ1B |
| 1,3637 | -2,21345 | -1,4592 | -1,4592 | 1,8856325 | 0,552859 | Q9NP73;A0 | ALG13 |
| -0,564834 | 2,12763 | 0,145424 | 0,145424 | 1,3954073 | 0,552954 | F8VW96;Q1 | CSRP2 |
| -1,26234 | 1,54247 | 1,79191 | 1,54247 | 1,6959571 | 0,553662 | Q9UNA4;X6 | POLI |
| 0,817558 | -0,885947 | 1,64444 | 0,817558 | 1,2902533 | 0,553733 | H0YCT1 | NUP98 |
| -1,57545 | 3,58933 | 1,13889 | 1,13889 | 2,5835134 | 0,554074 | Q6ZSZ6;H0Y | TSHZ1 |
| 0,520087 | -1,71101 | -0,19521 | -0,19521 | 1,1392317 | 0,555134 | P02792 | FTL |
| -1,9035 | 3,85528 | 1,57497 | 1,57497 | 2,9000895 | 0,555323 | Q8NA31;J3 | TERB1 |
| 0,741493 | -1,74309 | -0,50321 | -0,50321 | 1,2422923 | 0,556722 | Q14008;H0 | CKAP5 |
| -1,61081 | 1,02884 | -1,1177 | -1,1177 | 1,4034804 | 0,556805 | O95714;A0 | HERC2 |
| -3,05638 | -3,0405 | 2,33782 | -3,0405 | 3,1097688 | 0,557465 | Q8N841 | TTLL6 |
| -0,93572 | 0,808244 | -1,17818 | -0,93572 | 1,0836726 | 0,558628 | Q00325 | SLC25A3 |
| 0,826461 | -0,809393 | -1,41216 | -0,809393 | 1,1583588 | 0,558768 | P15531 | NME1 |
| -2,89556 | 1,34538 | -1,0057 | -1,0057 | 2,1246459 | 0,55918 | Q4AC99 | ACCSL |
| 2,07786 | -1,82097 | 2,68225 | 2,07786 | 2,4442161 | 0,559324 | Q5GLZ8 | HERC4 |
| 0,142463 | 1,36039 | -0,413297 | 0,142463 | 0,9072101 | 0,559763 | Q16643;D6 | DBN1 |
| 2,20086 | -2,75162 | -2,95439 | -2,75162 | 2,9196112 | 0,559896 | F8W7A7;Q5 | CCDC178 |
| -2,45252 | 1,3563 | -1,23866 | -1,23866 | 1,9456954 | 0,56005 | Q9P2I0 | CPSF2 |
| -2,63996 | 0,0202348 | 0,563067 | 0,0202348 | 1,71419 | 0,560122 | Q96PU5;A0 | NEDD4L |
| 0,360684 | -1,67342 | 0,0102493 | 0,0102493 | 1,0874378 | 0,560722 | A8MX29 | KLC2 |
| 1,53999 | -1,40601 | -2,77441 | -1,40601 | 2,204748 | 0,560765 | P13645;C4A | KRT10 |
| -1,48777 | 2,23751 | 1,64726 | 1,64726 | 2,0022703 | 0,560904 | Q13367;A0 | AP3B2 |
| -0,412449 | -0,0066005 | 0,0968468 | -0,006601 | 0,2691953 | 0,560973 | Q04837;A0 | SSBP1 |
| 0,939945 | -1,18705 | -1,24021 | -1,18705 | 1,2436512 | 0,561265 | Q9UNA1 | ARHGAP26 |
| -0,0195854 | 0,056991 | -0,185726 | -0,019585 | 0,1240821 | 0,561438 | Q13948 | CUX1 |
| 3,76559 | -2,07849 | 1,87689 | 1,87689 | 2,9823226 | 0,561528 | Q96NX9;D6 | DACH2 |
| 1,44762 | -2,68839 | -1,26358 | -1,26358 | 2,1010818 | 0,562449 | Q8IZY2 | ABCA7 |
| 0,70526 | -0,164349 | -2,53772 | -0,164349 | 1,6785921 | 0,563149 | O43852;H0 | CALU |
| -1,61036 | -0,458899 | 0,698749 | -0,458899 | 1,1545559 | 0,563901 | Q14693 | LPIN1 |
| -1,50793 | 2,58719 | 1,42359 | 1,42359 | 2,1102047 | 0,56419 | Q14929 | ZNF169 |
| -2,23137 | -1,90177 | 1,61398 | -1,90177 | 2,1313474 | 0,565418 | Q2TAL8;C9 | QRICH1 |
| -0,0783606 | -0,889706 | 0,266542 | -0,078361 | 0,5935976 | 0,565458 | O95373;E9I | IPO7 |
| 2,71277 | 1,42613 | -1,55423 | 1,42613 | 2,1888078 | 0,565746 | P86790;P86 | CCZ1B |
| -1,36661 | 0,255251 | 0,0647568 | 0,0647568 | 0,8865224 | 0,565831 | P30041 | PRDX6 |
| -2,33824 | 1,87315 | 4,57308 | 1,87315 | 3,4830966 | 0,566177 | Q9NUJ1 | ABHD10 |
| 1,89412 | -2,90704 | -1,99353 | -1,99353 | 2,5494921 | 0,566229 | P81605 | DCD |
| -1,26521 | 0,594434 | -0,427494 | -0,427494 | 0,9313414 | 0,566229 | E9PHG3;Q8 | SMYD1 |

| | | | | | | | |
|---|---|---|---|---|---|---|---|
| -1,05412 | 1,01042 | -1,56293 | -1,05412 | 1,3628 | 0,566323 | P62987;B4I | UBA52 |
| 0,838866 | -0,334476 | -2,50199 | -0,334476 | 1,6949025 | 0,566421 | Q86YS6;C9. | RAB43 |
| -3,13738 | -0,435616 | 1,06394 | -0,435616 | 2,1291347 | 0,566467 | A0A0A0MT | CTAG1A |
| 1,28583 | -0,499785 | 0,268384 | 0,268384 | 0,8957028 | 0,566835 | H0YBS9;Q9 | WWP1 |
| 2,51479 | -3,11213 | -3,27992 | -3,11213 | 3,2982077 | 0,567326 | P61160;F5H | ACTR2 |
| 0,324909 | 0,929733 | -0,445488 | 0,324909 | 0,6892697 | 0,567815 | Q08209;E7I | PPP3CA |
| -2,15373 | 0,141058 | 0,374137 | 0,141058 | 1,3970498 | 0,568137 | Q13867;K7I | BLMH |
| 2,24753 | -3,16415 | -2,55706 | -2,55706 | 2,9647632 | 0,568497 | P35658;A0/ | NUP214 |
| -1,21099 | -0,504881 | 0,629806 | -0,504881 | 0,928676 | 0,569149 | Q07343;E9I | PDE4B |
| 5,90492 | -2,87874 | 2,12561 | 2,12561 | 4,4060448 | 0,569216 | A0A2R8Y54 | DYNC1H1 |
| -1,45798 | 2,24859 | 1,49645 | 1,49645 | 1,9592955 | 0,569806 | P27816 | MAP4 |
| 0,210606 | -0,650699 | 2,04937 | 0,210606 | 1,379207 | 0,569949 | Q9UHJ3 | SFMBT1 |
| -0,109109 | 0,133556 | -0,251729 | -0,109109 | 0,1947953 | 0,569961 | Q96JG9;H3 | ZNF469 |
| -0,585108 | 0,46335 | 1,12667 | 0,46335 | 0,8630799 | 0,570695 | P24666;G5I | ACP1 |
| 1,5 | -2,75181 | -1,25567 | -1,25567 | 2,1567738 | 0,571215 | Q01546 | KRT76 |
| -0,398959 | 0,11711 | 1,2824 | 0,11711 | 0,8613164 | 0,571501 | P07910 | HNRNPC |
| 1,51289 | -0,32725 | -0,0368324 | -0,036832 | 0,989284 | 0,571622 | P51532;Q9I | SMARCA4 |
| 0,871262 | -0,386494 | 0,244641 | 0,244641 | 0,6288794 | 0,572043 | Q9UPN4;I3 | CEP131 |
| 1,7887 | -3,49401 | -1,37521 | -1,37521 | 2,6585292 | 0,572382 | A8MW92;A | PHF20L1 |
| -3,21997 | 3,8718 | 4,20587 | 3,8718 | 4,1942004 | 0,572543 | Q76LX8 | ADAMTS13 |
| 0,801671 | -0,876882 | -1,14677 | -0,876882 | 1,0556832 | 0,572746 | P30419;B7Z | NMT1 |
| 0,685485 | 0,0351919 | -0,193623 | 0,0351919 | 0,4560826 | 0,573323 | Q9ULI0;H7I | ATAD2B |
| 0,0542242 | 0,485132 | -0,159989 | 0,0542242 | 0,3285701 | 0,573629 | E9PM04;Q9 | DCUN1D5 |
| -0,302685 | 0,828444 | -2,44327 | -0,302685 | 1,6616091 | 0,573808 | Q92805 | GOLGA1 |
| 0,215705 | -1,93641 | 0,268881 | 0,215705 | 1,2581557 | 0,573831 | B1AHC2;B1 | ARHGAP8 |
| -1,93507 | 1,86108 | 3,10252 | 1,86108 | 2,6245394 | 0,573832 | P53004;C9J | BLVRA |
| 0,0972968 | -0,820057 | 0,108602 | 0,0972968 | 0,532928 | 0,574287 | Q63HN8;A0 | RNF213 |
| -1,93809 | 3,87786 | 1,42342 | 1,42342 | 2,9197403 | 0,574451 | Q14353;A0 | GAMT |
| 4,10416 | -2,80477 | 2,96551 | 2,96551 | 3,7041861 | 0,574604 | Q96KR1;H0 | ZFR |
| -1,71508 | 1,3902 | -1,7509 | -1,71508 | 1,8032635 | 0,57468 | Q86W50 | METTL16 |
| 5,59983 | 0,58383 | -1,82528 | 0,58383 | 3,7880586 | 0,574851 | P15428 | HPGD |
| -0,821527 | 0,499736 | -0,464446 | -0,464446 | 0,6834825 | 0,574918 | O75083;D6 | WDR1 |
| -2,58183 | 0,376049 | 0,27478 | 0,27478 | 1,679262 | 0,575047 | Q7RTT5 | SSX7 |
| -1,75769 | 0,0304195 | 0,40044 | 0,0304195 | 1,1541069 | 0,575123 | P62424;Q5 | RPL7A |
| 0,799961 | -0,931292 | -1,06357 | -0,931292 | 1,0398303 | 0,575284 | Q9Y277;E5I | VDAC3 |
| -0,928262 | 0,125886 | 0,109933 | 0,109933 | 0,6040601 | 0,576138 | O60566 | BUB1B |
| -0,549637 | 2,46724 | -0,0619438 | -0,061944 | 1,6194733 | 0,576281 | P30837 | ALDH1B1 |
| -0,724899 | 1,08064 | 0,743008 | 0,743008 | 0,9599234 | 0,57665 | Q12797;E5I | ASPH |
| -0,921014 | 0,406927 | -0,244748 | -0,244748 | 0,6640084 | 0,577201 | M0QYS1;P4 | RPL13A |
| 1,6013 | -2,76757 | -1,38165 | -1,38165 | 2,2325542 | 0,577673 | H0YKN8;H0 | TLE3 |
| -1,83764 | -2,67948 | 1,80312 | -1,83764 | 2,3824893 | 0,578316 | B7ZC32 | KIF28P |
| -2,37882 | 2,65423 | 3,23879 | 2,65423 | 3,088442 | 0,578709 | O14939 | PLD2 |
| -0,755659 | 0,0532267 | 2,85981 | 0,0532267 | 1,8974905 | 0,57898 | P24588 | AKAP5 |
| 1,04284 | -0,632227 | -2,31364 | -0,632227 | 1,678241 | 0,579901 | P54920;M0 | NAPA |
| 3,62209 | 6,27807 | -3,90925 | 3,62209 | 5,2845183 | 0,579984 | E9PJT4 | DNHD1 |
| -1,50725 | -0,206742 | 0,539755 | -0,206742 | 1,0359222 | 0,580031 | O43747;H3 | AP1G1 |
| -0,144684 | 2,04141 | -0,383132 | -0,144684 | 1,3363051 | 0,580289 | P28827;E7E | PTPRM |
| -0,256949 | -2,11775 | 0,735179 | -0,256949 | 1,4483383 | 0,580493 | P63241;I3L | EIF5A |
| -1,45212 | -0,261374 | 0,565449 | -0,261374 | 1,01424 | 0,580516 | Q9Y3F4;H0 | STRAP |
| 0,876248 | -1,56164 | -0,715131 | -0,715131 | 1,2377643 | 0,580649 | A0A0D9SF6 | PKP4 |
| 0,401441 | -0,486608 | 0,856151 | 0,401441 | 0,6829341 | 0,581432 | Q9UM54;E7 | MYO6 |
| 0,401434 | -0,486618 | 0,856136 | 0,401434 | 0,6829322 | 0,581446 | A0A0A0MR | MYO6 |

| | | | | | | | |
|---|---|---|---|---|---|---|---|
| 2,8637 | -0,793268 | 0,0836209 | 0,0836209 | 1,9092402 | 0,581648 | Q9P219 | CCDC88C |
| 2,18102 | 0,768785 | -1,09574 | 0,768785 | 1,6435742 | 0,581696 | P31939;H7( | ATIC |
| -0,183772 | 0,146853 | 0,33116 | 0,146853 | 0,2609077 | 0,58179 | Q12765;C9 | SCRN1 |
| -3,12925 | 4,15744 | 3,52284 | 3,52284 | 4,0362703 | 0,58186 | Q9UBF8 | PI4KB |
| -1,44137 | 1,50958 | 2,05024 | 1,50958 | 1,8793509 | 0,581955 | P16520;E9F | GNB3 |
| -2,93105 | -1,91042 | 1,94517 | -1,91042 | 2,5717951 | 0,582275 | Q9P2S2;G5 | NRXN2 |
| 2,62776 | -0,770015 | 0,12447 | 0,12447 | 1,761223 | 0,582489 | Q9Y2H0;A0 | DLGAP4 |
| -2,87804 | 1,697 | -1,45195 | -1,45195 | 2,3409617 | 0,582711 | Q14694;H3 | USP10 |
| -0,697425 | 0,543694 | -0,629879 | -0,629879 | 0,6978792 | 0,583295 | Q9NZZ3 | CHMP5 |
| -0,195785 | 0,810547 | -2,55284 | -0,195785 | 1,7263055 | 0,583348 | P51114;B4I | FXR1 |
| 0,717686 | -1,49344 | -0,466345 | -0,466345 | 1,1064908 | 0,583384 | Q3ZCM7;A( | TUBB8 |
| -3,15612 | 3,04422 | 4,79682 | 3,04422 | 4,1786177 | 0,583811 | Q92621 | NUP205 |
| 1,89671 | -1,45368 | -3,4848 | -1,45368 | 2,7175728 | 0,584385 | Q6ZV73;F8' | FGD6 |
| -2,65124 | -0,184149 | 0,832125 | -0,184149 | 1,7913302 | 0,584688 | A0A1W2PR | MED17 |
| 1,20537 | -2,85466 | -0,621161 | -0,621161 | 2,0334116 | 0,585222 | A0A0A0MR | CACNA1C |
| 1,87074 | -1,32675 | 1,37362 | 1,37362 | 1,7206137 | 0,585862 | Q9Y520;E7I | PRRC2C |
| 0,86679 | -1,02898 | 1,73584 | 0,86679 | 1,4138259 | 0,58631 | Q92542;H0 | NCSTN |
| 1,32157 | -0,460272 | 0,146902 | 0,146902 | 0,9058574 | 0,586328 | P34931;Q5: | HSPA1L |
| 0,273027 | 0,0039189 | -0,0743266 | 0,0039189 | 0,1822069 | 0,586621 | Q9BR76;A0 | CORO1B |
| -2,04352 | 2,93206 | 2,06657 | 2,06657 | 2,6582673 | 0,586732 | P11274;A9l | BCR |
| 3,09364 | -1,79184 | 1,45811 | 1,45811 | 2,4868001 | 0,5873 | Q9HCK8;A0 | CHD8 |
| -3,15797 | 4,12747 | 3,50999 | 3,50999 | 4,0398145 | 0,587605 | Q9C0G6 | DNAH6 |
| -2,65267 | 4,39304 | 2,26353 | 2,26353 | 3,6135362 | 0,587855 | Q8WX94;A( | NLRP7 |
| -1,51041 | -0,481857 | 0,74259 | -0,481857 | 1,1279185 | 0,587879 | P22102;F8\ | GART |
| 0,0454961 | -0,914762 | 0,200159 | 0,0454961 | 0,6040233 | 0,587941 | P24539;Q5( | ATP5PB |
| -0,752916 | 0,897879 | -1,5038 | -0,752916 | 1,228618 | 0,588486 | Q14151 | SAFB2 |
| -2,73417 | -0,916245 | 1,3752 | -0,916245 | 2,0592269 | 0,588825 | P20340;F5H | RAB6A |
| 0,316086 | -0,198055 | 0,175564 | 0,175564 | 0,2657312 | 0,588834 | O00231;J3( | PSMD11 |
| 0,790977 | -0,407861 | 0,2816 | 0,2816 | 0,6016691 | 0,588856 | O75113 | N4BP1 |
| -1,28948 | -0,733026 | 0,817085 | -0,733026 | 1,0916426 | 0,58903 | P51648;J3C | ALDH3A2 |
| 0,730246 | -0,520621 | -1,38181 | -0,520621 | 1,0620025 | 0,589177 | P18074;E7E | ERCC2 |
| 1,10351 | -1,14084 | 1,68136 | 1,10351 | 1,4908516 | 0,589488 | Q5SZK4;Q5 | WASF1 |
| -2,39297 | -1,97535 | 1,81559 | -1,97535 | 2,3186781 | 0,590046 | Q16777;Q6 | HIST2H2AC |
| 1,91475 | -2,44039 | 4,25509 | 1,91475 | 3,3978888 | 0,591089 | Q9Y6M9;E9 | NDUFB9 |
| 0,878587 | -1,87037 | -0,514314 | -0,514314 | 1,3745197 | 0,591664 | Q9P2J8;C9J | ZNF624 |
| 2,14182 | -2,06921 | -3,12624 | -2,06921 | 2,78695 | 0,591676 | Q96J65 | ABCC12 |
| -1,46073 | 1,08106 | -1,13489 | -1,13489 | 1,3830704 | 0,591867 | P52788;H7( | SMS |
| 1,22735 | -1,74616 | -1,21547 | -1,21547 | 1,5859153 | 0,592341 | P30622;J3K | CLIP1 |
| 0,63642 | -2,91909 | 0,170539 | 0,170539 | 1,9323778 | 0,592505 | H0Y325 | SYNE1 |
| -2,55412 | 3,96178 | 2,28968 | 2,28968 | 3,3841609 | 0,592654 | H7BYX6 | NCAM1 |
| 2,54505 | 0,383009 | -0,984455 | 0,383009 | 1,7795966 | 0,592772 | O94885 | SASH1 |
| -0,781677 | -0,972077 | 0,733829 | -0,781677 | 0,9348018 | 0,593115 | Q92734;Q0 | TFG |
| 2,8834 | 0,861058 | -1,40681 | 0,861058 | 2,1462756 | 0,593706 | Q9UPV0;E9 | CEP164 |
| -0,827727 | 0,295197 | -0,0876455 | -0,087646 | 0,5708542 | 0,594567 | Q15019;B5 | SEPT2 |
| -1,44928 | -2,03011 | 1,453 | -1,44928 | 1,8660407 | 0,594714 | P61020;F8\ | RAB5B |
| 1,75289 | 1,96012 | -1,5658 | 1,75289 | 1,9785836 | 0,594933 | Q9ULV4;B4 | CORO1C |
| -1,49604 | 1,37093 | -1,7586 | -1,49604 | 1,7360114 | 0,594979 | P46063;F8\ | RECQL |
| 2,08773 | -1,4363 | -3,93075 | -1,4363 | 3,0238819 | 0,595168 | F8WAI0;Q5 | SYNE1 |
| 0,665161 | -0,437533 | -1,28762 | -0,437533 | 0,9791098 | 0,59575 | Q7Z406;M( | MYH14 |
| -0,547767 | 0,471639 | 0,864104 | 0,471639 | 0,7287657 | 0,596176 | P08133;E5F | ANXA6 |
| -1,42779 | 1,39702 | 2,00846 | 1,39702 | 1,8330859 | 0,596912 | F8VV14 | CACNB3 |
| -0,357662 | 0,974371 | 0,111608 | 0,111608 | 0,6756338 | 0,597199 | Q9Y230;M( | RUVBL2 |

| | | | | | | | |
|---|---|---|---|---|---|---|---|
| 2,76423 | -3,22832 | -3,27817 | -3,22832 | 3,4742802 | 0,597463 | Q92620 | DHX38 |
| 0,781965 | -0,32769 | -1,90317 | -0,32769 | 1,3492851 | 0,5985 | P28838;H0Y | LAP3 |
| 2,0047 | -2,82339 | -1,94106 | -1,94106 | 2,5709266 | 0,598618 | Q9NRC6 | SPTBN5 |
| 1,66815 | -1,10746 | -3,16264 | -1,10746 | 2,4243318 | 0,598673 | F5GYK2;Q9 | STRN4 |
| 0,204287 | -2,83832 | 0,611691 | 0,204287 | 1,8852945 | 0,598854 | Q9UQ16;E5 | DNM3 |
| -0,866378 | 1,48562 | 0,658122 | 0,658122 | 1,1930876 | 0,5995 | P43487;F6V | RANBP1 |
| 1,53434 | -1,15994 | 1,19718 | 1,19718 | 1,4679258 | 0,599506 | Q6ZW49 | PAXIP1 |
| 3,85026 | 5,28937 | -3,87546 | 3,85026 | 4,9286898 | 0,600307 | S4R3L9 | KCTD10 |
| -0,00639992 | 0,436729 | -1,53231 | -0,0064 | 1,0329472 | 0,600696 | Q96RP9;C9 | GFM1 |
| 1,22667 | -0,231829 | -0,12764 | -0,12764 | 0,8136574 | 0,601019 | P36957;Q8 | DLST |
| 0,823436 | -0,783146 | -1,16163 | -0,783146 | 1,0539484 | 0,601605 | Q8NEY1;A0 | NAV1 |
| -0,168955 | 1,03693 | -0,137093 | -0,137093 | 0,6872049 | 0,601726 | P13667;C9J | PDIA4 |
| -1,3791 | -2,77143 | 1,71217 | -1,3791 | 2,2948203 | 0,602045 | O14715;J3K | RGPD8 |
| 0,628972 | -0,902049 | -0,583588 | -0,583588 | 0,8078515 | 0,602715 | O76013 | KRT36 |
| 1,7642 | -3,58423 | -1,01214 | -1,01214 | 2,6748649 | 0,603223 | Q3BBV0;S4 | NBPF1 |
| 0,478704 | -0,460219 | 0,593635 | 0,478704 | 0,5781283 | 0,603229 | P02768;A0/ | ALB |
| -0,780299 | -0,00188804 | 0,224617 | -0,001888 | 0,5271123 | 0,603549 | P30086;O1! | PEBP1 |
| -2,69435 | 0,60954 | 0,184626 | 0,184626 | 1,79744 | 0,603745 | Q96GC6;A0 | ZNF274 |
| 0,825383 | -0,815288 | 1,07522 | 0,825383 | 1,0269892 | 0,603862 | Q14155 | ARHGEF7 |
| -1,90491 | 1,00514 | -0,641477 | -0,641477 | 1,4592236 | 0,604045 | Q96DB9;A8 | FXYD5 |
| -0,372794 | -2,91277 | 1,12831 | -0,372794 | 2,0426747 | 0,604081 | O95613;H7 | PCNT |
| 1,87566 | -0,906398 | 0,499824 | 0,499824 | 1,3910567 | 0,604083 | O00273;K7I | DFFA |
| 0,756009 | -1,86924 | -0,282807 | -0,282807 | 1,3221095 | 0,604138 | D6RJC6 | DTNBP1 |
| -0,36201 | -0,219787 | 1,94909 | -0,219787 | 1,2952116 | 0,604222 | Q8TAE8 | GADD45GIP1 |
| -0,128456 | 2,59015 | -0,633543 | -0,128456 | 1,7338843 | 0,604628 | Q9UJ14 | GGT7 |
| 1,7372 | 1,11696 | -1,21382 | 1,11696 | 1,555943 | 0,60467 | A0A0G2JR6 | KIF26B |
| 4,15347 | -3,34716 | 3,5962 | 3,5962 | 4,1789203 | 0,604902 | Q6P597;K7 | KLC3 |
| 2,7521 | -1,86411 | 1,65019 | 1,65019 | 2,4108748 | 0,605123 | Q9Y4F1;C9. | FARP1 |
| -2,32698 | 0,914387 | -0,310393 | -0,310393 | 1,6367228 | 0,605155 | Q14738;E9I | PPP2R5D |
| -0,429752 | 1,51186 | -0,00957281 | -0,009573 | 1,021532 | 0,606034 | Q9Y2T7 | YBX2 |
| 1,58817 | -0,107173 | -0,366636 | -0,107173 | 1,0616634 | 0,606126 | K7EQJ5;P62 | RPS15 |
| 3,35413 | 2,93926 | -2,73143 | 2,93926 | 3,4000707 | 0,606769 | A0A2R8YGI | AP1S1 |
| 2,56417 | -2,22086 | -3,82557 | -2,22086 | 3,3241636 | 0,606786 | Q8N8A6 | DDX51 |
| -0,724732 | 0,666771 | -0,809742 | -0,724732 | 0,8290153 | 0,607069 | E7EWN3;Q! | SETD5 |
| 2,03716 | -3,14157 | -1,68982 | -1,68982 | 2,6713668 | 0,607282 | Q6P9B6 | TLDC1 |
| -0,697161 | 1,67607 | 0,26759 | 0,26759 | 1,1935092 | 0,607789 | Q9ULT8;A0 | HECTD1 |
| -3,21177 | 2,88395 | 4,60793 | 2,88395 | 4,1084794 | 0,608623 | P38570 | ITGAE |
| -1,29809 | 0,178287 | 0,219299 | 0,178287 | 0,864469 | 0,608652 | H0YK42;Q1 | SNX1 |
| -0,519307 | 0,182236 | -0,0346473 | -0,034647 | 0,359188 | 0,610818 | M0R117;M | RPL18A |
| 0,552267 | -0,0188829 | -0,147964 | -0,018883 | 0,3726476 | 0,611013 | Q8IZS8;B4D | CACNA2D3 |
| -1,38622 | -2,24124 | 1,56373 | -1,38622 | 1,9962908 | 0,611171 | Q8NF50;A2 | DOCK8 |
| 0,0303345 | -0,142688 | 0,397536 | 0,0303345 | 0,275867 | 0,611176 | Q13177;H7 | PAK2 |
| 0,16942 | -0,41996 | 0,973289 | 0,16942 | 0,6993708 | 0,611284 | Q13162;H7 | PRDX4 |
| 2,32095 | -2,80413 | -2,4799 | -2,4799 | 2,8699515 | 0,611595 | P27338 | MAOB |
| 0,919347 | -0,0694144 | -0,213373 | -0,069414 | 0,6166344 | 0,61164 | Q7L2H7;J3K | EIF3M |
| 1,11099 | -0,588755 | -2,26432 | -0,588755 | 1,6876694 | 0,611661 | Q7L2E3;F6F | DHX30 |
| -1,22033 | 0,260375 | 3,38719 | 0,260375 | 2,3522578 | 0,611781 | Q58FF8 | HSP90AB2P |
| 2,83841 | -2,99607 | -3,46648 | -2,99607 | 3,5122187 | 0,611783 | P11047;R4( | LAMC1 |
| 0,688162 | -0,479798 | 0,423464 | 0,423464 | 0,612383 | 0,611818 | P53675;A0/ | CLTCL1 |
| 2,13103 | -2,22185 | -2,63191 | -2,22185 | 2,6394858 | 0,611887 | Q96AC1;H0 | FERMT2 |
| 2,06545 | -2,58474 | -2,12317 | -2,12317 | 2,5619606 | 0,611926 | P60510;H3I | PPP4C |
| 2,59244 | -2,76022 | -3,13567 | -2,76022 | 3,2042466 | 0,612073 | Q4VXU2 | PABPC1L |

| | | | | | | | |
|---|---|---|---|---|---|---|---|
| -2,89385 | 0,666977 | 0,228889 | 0,228889 | 1,9417736 | 0,612716 | Q01105;A0 | SET |
| 0,65983 | -0,485711 | -1,08325 | -0,485711 | 0,8857807 | 0,613544 | Q12756;F8\ | KIF1A |
| 0,0775583 | -1,04609 | 0,247482 | 0,0775583 | 0,7029448 | 0,613738 | P22090;C9J | RPS4Y1 |
| 0,246429 | 0,183819 | -1,36669 | 0,183819 | 0,9137971 | 0,614051 | Q04206;A0 | RELA |
| 0,94448 | 0,0360096 | -3,2283 | 0,0360096 | 2,1944296 | 0,614196 | Q7Z4H8;H0 | KDELC2 |
| -0,112523 | 0,764216 | -0,129374 | -0,112523 | 0,5111194 | 0,61497 | Q14103;H0 | HNRNPD |
| 2,05784 | -1,93363 | -2,75368 | -1,93363 | 2,5740707 | 0,615096 | Q96MT7 | CFAP44 |
| 3,23045 | -3,5142 | -3,76785 | -3,5142 | 3,9692746 | 0,615346 | H0YLX1 | CHP1 |
| 0,442134 | -0,936979 | -0,208468 | -0,208468 | 0,6899232 | 0,615774 | Q9Y6N5;H3 | SQOR |
| -1,97735 | 0,155343 | 0,467495 | 0,155343 | 1,3306067 | 0,616237 | O15145;C9 | ARPC3 |
| 0,513227 | -1,44733 | -0,0878924 | -0,087892 | 1,0044235 | 0,616389 | Q13615;C9 | MTMR3 |
| -2,59883 | -0,715851 | 1,32083 | -0,715851 | 1,9603322 | 0,616516 | A6NHK2;P6 | SNRPE |
| 1,06208 | -1,61086 | 2,81345 | 1,06208 | 2,2280943 | 0,616736 | Q99714;Q5 | HSD17B10 |
| -0,67252 | -1,38853 | 0,881861 | -0,67252 | 1,1607071 | 0,616894 | Q9Y6X9;H7 | MORC2 |
| 0,449201 | -0,655458 | 1,1145 | 0,449201 | 0,8940214 | 0,616899 | P25685;M0 | DNAJB1 |
| -2,25327 | -0,175265 | 0,832307 | -0,175265 | 1,57343 | 0,617356 | H0YJB9 | RTRAF |
| -0,934818 | 1,56686 | -2,9038 | -0,934818 | 2,2406132 | 0,617546 | Q8NE09;A0 | RGS22 |
| 0,173583 | 0,051073 | -0,709934 | 0,051073 | 0,4786688 | 0,617575 | E9PAV3;F8\ | NACA |
| -1,00811 | 0,830658 | 1,48861 | 0,830658 | 1,294062 | 0,61777 | Q13620;K4 | CUL4B |
| -2,80202 | 1,40785 | -0,737472 | -0,737472 | 2,1050641 | 0,617956 | E9PIZ2;Q8N | LARGE2 |
| -0,697004 | 0,728629 | -0,946674 | -0,697004 | 0,9038256 | 0,618023 | P10599 | TXN |
| -0,742882 | -0,284538 | 0,429357 | -0,284538 | 0,5907438 | 0,618032 | Q86TG7;A0 | PEG10 |
| 1,11395 | -0,97694 | -1,56032 | -0,97694 | 1,4061693 | 0,618097 | Q9H0K6 | PUS7L |
| -2,98188 | 3,94236 | 2,79336 | 2,79336 | 3,7107658 | 0,618284 | Q8WZ64;D( | ARAP2 |
| 4,5854 | 0,647086 | -1,92037 | 0,647086 | 3,2768682 | 0,618559 | A0A0U1RRI | FGGY |
| 4,37805 | 1,78252 | -2,59831 | 1,78252 | 3,526047 | 0,618715 | Q15833;R4 | STXBP2 |
| -1,42079 | -2,24357 | 1,61546 | -1,42079 | 2,0325638 | 0,619436 | Q8WXF1;X6 | PSPC1 |
| 0,973688 | -1,0671 | 1,43562 | 0,973688 | 1,3317785 | 0,619504 | Q8WWM7; | ATXN2L |
| -1,61734 | 1,18988 | 2,57896 | 1,18988 | 2,1377153 | 0,61995 | Q9GZX5 | ZNF350 |
| 0,0870562 | -2,5549 | 0,720571 | 0,0870562 | 1,7373343 | 0,620185 | A0A0C4DF\ | LSM14B |
| -2,09156 | 3,6235 | 1,36271 | 1,36271 | 2,8782247 | 0,620189 | Q96BD8 | SKA1 |
| -1,07171 | 1,33384 | 1,0628 | 1,0628 | 1,3175904 | 0,620233 | P49588;H3I | AARS |
| 0,877674 | -1,6965 | -0,474776 | -0,474776 | 1,2876401 | 0,620537 | O95394;A0 | PGM3 |
| 0,980787 | -1,39515 | 2,28484 | 0,980787 | 1,8658312 | 0,621228 | O95716;H0 | RAB3D |
| -0,776434 | 0,248263 | 1,85937 | 0,248263 | 1,3287295 | 0,621434 | Q9BZZ5;H0 | API5 |
| -0,439843 | 0,279794 | 0,76806 | 0,279794 | 0,6076335 | 0,621834 | Q07864;F5I | POLE |
| -2,04805 | -2,47391 | 2,03315 | -2,04805 | 2,4883442 | 0,621974 | P51148;F8\ | RAB5C |
| -1,66602 | 1,15062 | 2,75287 | 1,15062 | 2,2370835 | 0,621981 | J3KPQ0;P2 | FGFR4 |
| -1,37798 | 3,24342 | 0,457556 | 0,457556 | 2,3269282 | 0,622583 | Q86VW0;H | SESTD1 |
| -1,12569 | 0,177339 | 0,193617 | 0,177339 | 0,757047 | 0,623026 | Q562R1 | ACTBL2 |
| -2,78747 | 0,196515 | 0,709145 | 0,196515 | 1,8882652 | 0,623144 | M0R344 | SPHK2 |
| 1,45262 | -2,18149 | -1,1331 | -1,1331 | 1,8704645 | 0,623509 | O94822;H7 | LTN1 |
| -3,40456 | 3,13599 | -3,52523 | -3,40456 | 3,8115003 | 0,623541 | E5RFR7;H0 | TPD52 |
| -0,0257945 | 1,70489 | -0,517552 | -0,025795 | 1,1673576 | 0,623651 | P35269;M0 | GTF2F1 |
| 2,41248 | 0,885132 | -1,39254 | 0,885132 | 1,9148002 | 0,623682 | P78344;D3I | EIF4G2 |
| -0,0473056 | 0,9793 | -0,269543 | -0,047306 | 0,6661978 | 0,62386 | A0A0A0MR | MPP2 |
| -0,536193 | 0,48074 | 0,717081 | 0,48074 | 0,6659211 | 0,624127 | Q86V48;E5 | LUZP1 |
| 0,653605 | -1,40674 | -0,271354 | -0,271354 | 1,0319619 | 0,624385 | Q92616 | GCN1 |
| -0,983167 | 1,30226 | 0,886068 | 0,886068 | 1,2172669 | 0,625265 | Q13362;Q9 | PPP2R5C |
| -0,388043 | 0,207351 | 0,738167 | 0,207351 | 0,5634135 | 0,625458 | P46940;H0 | IQGAP1 |
| -2,34886 | 1,92395 | 3,36416 | 1,92395 | 2,9712429 | 0,625533 | A0A0A0MR | ZNF691 |
| 2,61401 | -2,65593 | -3,10093 | -2,65593 | 3,1788581 | 0,625714 | Q9UKK3 | PARP4 |

| | | | | | | | |
|---|---|---|---|---|---|---|---|
| 1,32608 | -2,50346 | -0,715923 | -0,715923 | 1,9161785 | 0,625913 | A0A0U1RR( | SYTL2 |
| -1,15501 | 0,135182 | 0,249409 | 0,135182 | 0,7799611 | 0,626009 | P12268;H0' | IMPDH2 |
| -0,54406 | 0,763945 | 0,455281 | 0,455281 | 0,6837194 | 0,626099 | P27635;X1\ | RPL10 |
| 0,573195 | -0,284322 | 0,133864 | 0,133864 | 0,4288019 | 0,626632 | Q02252;G3 | ALDH6A1 |
| -1,58736 | 1,10361 | -0,893578 | -0,893578 | 1,3971049 | 0,626636 | Q8NCM8;H | DYNC2H1 |
| 0,195154 | -0,758856 | 1,88061 | 0,195154 | 1,3365177 | 0,626803 | O60447;A0. | EVI5 |
| -0,727769 | 0,828093 | -1,11013 | -0,727769 | 1,0266138 | 0,627359 | P38935;F5( | IGHMBP2 |
| -2,17017 | 1,18649 | -0,669013 | -0,669013 | 1,6814443 | 0,627597 | O00754;M( | MAN2B1 |
| -2,63327 | -2,61713 | 2,39943 | -2,61713 | 2,9009827 | 0,627642 | E9PG46 | AAK1 |
| 0,709571 | -0,380557 | -1,33281 | -0,380557 | 1,0219658 | 0,627817 | O95571;M( | ETHE1 |
| 1,75089 | -2,02625 | -1,79813 | -1,79813 | 2,1179537 | 0,628868 | A0A0G2JRI! | MUC20 |
| -2,67278 | 0,761946 | 0,123013 | 0,123013 | 1,8267474 | 0,628971 | L8EBH5 | EML5 |
| -0,430451 | -1,20582 | 0,699309 | -0,430451 | 0,9580424 | 0,629198 | P49721;A0/ | PSMB2 |
| -0,484151 | 0,363521 | -0,318759 | -0,318759 | 0,4493344 | 0,629241 | P06748;E5I | NPM1 |
| -2,75655 | 0,834382 | 0,0730692 | 0,0730692 | 1,8921388 | 0,629441 | P08134;Q5. | RHOC |
| -1,40277 | 1,73277 | 1,33679 | 1,33679 | 1,7075128 | 0,629803 | Q15056 | EIF4H |
| -0,0665912 | 0,7379 | -2,09579 | -0,066591 | 1,4602884 | 0,630022 | P51570;K7I | GALK1 |
| 2,24784 | 0,669319 | -1,22574 | 0,669319 | 1,7391921 | 0,630986 | Q9Y6K8 | AK5 |
| 0,781533 | 0,274844 | -0,453653 | 0,274844 | 0,6209034 | 0,63158 | Q7KZI7;E9P | MARK2 |
| -2,79936 | -2,63156 | 2,50531 | -2,63156 | 3,0153804 | 0,631742 | P42702 | LIFR |
| -1,39031 | 3,7899 | 0,177156 | 0,177156 | 2,6565468 | 0,63183 | O43592;F8' | XPOT |
| -0,0093344 | -0,386855 | 1,19913 | -0,009334 | 0,8284782 | 0,632088 | Q8IWB6 | TEX14 |
| 2,75008 | -0,739301 | -0,192624 | -0,192624 | 1,8767944 | 0,632225 | Q12840;J3I | KIF5A |
| -2,00646 | 0,98384 | -0,425788 | -0,425788 | 1,4959651 | 0,632405 | Q99700;A0. | ATXN2 |
| 4,13169 | -1,52853 | 0,203243 | 0,203243 | 2,9002823 | 0,632595 | P08729;A0/ | KRT7 |
| 5,23344 | -1,7024 | -0,0293053 | -0,029305 | 3,6194293 | 0,632644 | Q17RW3;Q | ZNF619 |
| 5,7382 | 1,88776 | -3,25979 | 1,88776 | 4,5145502 | 0,632758 | E9PB61;Q8 | ALYREF |
| -0,902359 | -2,0754 | 1,31344 | -0,902359 | 1,7209507 | 0,63277 | O14529 | CUX2 |
| -2,06037 | -1,34323 | 1,55369 | -1,34323 | 1,9134552 | 0,63287 | F8VP71;F8\ | LETMD1 |
| 0,844435 | -0,338658 | 0,0729936 | 0,0729936 | 0,6005952 | 0,633899 | P36542 | ATP5F1C |
| -3,3348 | -1,35741 | 2,06249 | -1,35741 | 2,7305838 | 0,634097 | Q8TBA6;H0 | GOLGA5 |
| 3,05111 | -3,25568 | -3,31414 | -3,25568 | 3,6582196 | 0,634492 | Q16666;H3 | IFI16 |
| 0,838142 | -3,19637 | 0,261344 | 0,261344 | 2,1819628 | 0,634773 | Q9BXP5;H7 | SRRT |
| -0,448897 | 0,296703 | 0,721117 | 0,296703 | 0,592309 | 0,634935 | O00429;G8 | DNM1L |
| 0,201474 | -0,60044 | -0,0005491 | -0,000549 | 0,4170828 | 0,635804 | P61224;A8I | RAP1B |
| 1,46086 | -0,413048 | -0,0885939 | -0,088594 | 1,0014663 | 0,635826 | Q6ZMI0;F8' | PPP1R21 |
| -3,11379 | 2,40187 | -2,09455 | -2,09455 | 2,9348239 | 0,636337 | Q9BZ23;V9 | PANK2 |
| 1,45422 | 3,2351 | -2,09662 | 1,45422 | 2,7143826 | 0,636698 | Q86VV8 | RTTN |
| 1,76712 | -1,17123 | 0,83633 | 0,83633 | 1,5016973 | 0,637174 | O43526;A0. | KCNQ2 |
| 3,95749 | -0,579807 | -0,813871 | -0,579807 | 2,6897254 | 0,637353 | A8MTJ3 | GNAT3 |
| -2,34109 | -0,450403 | 1,13883 | -0,450403 | 1,7421348 | 0,638855 | P30048 | PRDX3 |
| 1,31783 | 1,19373 | -1,17992 | 1,19373 | 1,4076203 | 0,639725 | Q92974;V9 | ARHGEF2 |
| -2,07753 | 1,32862 | -0,885627 | -0,885627 | 1,7284569 | 0,639842 | Q9HC52 | CBX8 |
| -1,14834 | 2,10013 | 0,584448 | 0,584448 | 1,6254437 | 0,640025 | Q8NFF5 | FLAD1 |
| -1,12192 | 1,82914 | -3,01336 | -1,12192 | 2,4404954 | 0,640079 | P32942;K7I | ICAM3 |
| 2,25894 | -2,96785 | -1,9002 | -1,9002 | 2,7615713 | 0,640132 | Q6NUK1;J3 | SLC25A24 |
| 0,564048 | -1,12894 | 2,07873 | 0,564048 | 1,6046608 | 0,640593 | A0A0D9SF( | KCNT1 |
| 0,679785 | -0,578845 | -0,879073 | -0,578845 | 0,8270756 | 0,641448 | E9PKL7 | RAB2A |
| -0,330018 | 0,947491 | -2,01586 | -0,330018 | 1,4863569 | 0,641452 | A0A286YFL | PHGDH |
| -0,908874 | 3,09128 | -0,188501 | -0,188501 | 2,1321794 | 0,643335 | O14924 | RGS12 |
| 0,284326 | 1,65777 | -0,793318 | 0,284326 | 1,2285152 | 0,643353 | P07949 | RET |
| 1,73131 | 1,56983 | -1,5646 | 1,56983 | 1,8580344 | 0,643514 | Q9P0J1;E5I | PDP1 |

| | | | | | | | |
|---|---|---|---|---|---|---|---|
| 2,30936 | -3,22194 | -1,76316 | -1,76316 | 2,8667269 | 0,643925 | Q5T5P2 | KIAA1217 |
| -0,450688 | -1,47907 | 0,844167 | -0,450688 | 1,1641627 | 0,644215 | P78371;F8\ | CCT2 |
| 0,822605 | 0,259598 | -0,47569 | 0,259598 | 0,6510498 | 0,644521 | Q13136;A0. | PPFIA1 |
| 0,184312 | -1,74799 | 4,58907 | 0,184312 | 3,2479228 | 0,644556 | O94760;B4 | DDAH1 |
| -0,915905 | 1,98437 | 0,288675 | 0,288675 | 1,4570512 | 0,644574 | A0A0C4DH( | LTBP4 |
| -2,12632 | -0,247716 | 0,936255 | -0,247716 | 1,544361 | 0,644722 | P61956;A8I | SUMO2 |
| 2,18003 | -3,51503 | -1,34052 | -1,34052 | 2,8739193 | 0,644729 | A0A2R8Y5F | HARS2 |
| 0,111965 | 0,565897 | -1,89683 | 0,111965 | 1,3106192 | 0,645027 | O14618;J3I | CCS |
| 0,676824 | 1,85747 | -1,13328 | 0,676824 | 1,5063746 | 0,645032 | Q8N137 | CNTROB |
| -0,528972 | 0,951989 | 0,265638 | 0,265638 | 0,7411397 | 0,645324 | P47914 | RPL29 |
| 0,602251 | -2,64713 | 0,363328 | 0,363328 | 1,8110043 | 0,645545 | Q7L1Q6;C9 | BZW1 |
| 1,20193 | -1,43199 | -1,10044 | -1,10044 | 1,4345945 | 0,645907 | Q9Y618;C9. | NCOR2 |
| -2,14396 | -1,65632 | 1,80633 | -1,65632 | 2,1537769 | 0,646456 | A0A0A0MR | PCMT1 |
| -1,24534 | -0,150118 | 0,555957 | -0,150118 | 0,9076273 | 0,64674 | P41250;H7( | GARS |
| -3,06549 | 3,18075 | 3,22881 | 3,18075 | 3,6202218 | 0,647149 | F5GX77;Q9 | TRMT112 |
| 1,33343 | -1,30818 | 1,40166 | 1,33343 | 1,5452072 | 0,647242 | Q8NHW5 | RPLP0P6 |
| 0,753628 | -1,24845 | -0,434837 | -0,434837 | 1,0068707 | 0,647285 | Q9UBQ7 | GRHPR |
| -1,84062 | 2,24101 | 1,63043 | 1,63043 | 2,2015411 | 0,647572 | P58304 | VSX2 |
| -0,576388 | -1,06354 | 0,766371 | -0,576388 | 0,9477072 | 0,647806 | Q15126 | PMVK |
| 1,33037 | -1,60068 | -1,19186 | -1,19186 | 1,587442 | 0,64803 | Q9UHD8;K7 | SEPT9 |
| 0,492952 | -0,474613 | 0,497008 | 0,492952 | 0,5597985 | 0,648195 | Q92878;A0. | RAD50 |
| -1,02277 | -1,48884 | 1,19372 | -1,02277 | 1,4333049 | 0,64857 | A0A2R8Y5C | TBCE |
| 1,44742 | -0,328347 | -0,208306 | -0,208306 | 0,9924034 | 0,649151 | P54289 | CACNA2D1 |
| -3,01777 | -3,29742 | 3,03495 | -3,01777 | 3,5780007 | 0,649473 | Q9Y276;C9. | BCS1L |
| 1,24867 | -0,335244 | -2,76701 | -0,335244 | 2,0227026 | 0,649603 | P35251 | RFC1 |
| -2,14556 | 1,64632 | -1,32847 | -1,32847 | 1,9956355 | 0,649784 | P49448 | GLUD2 |
| -1,83267 | 1,82161 | 1,9864 | 1,82161 | 2,1589432 | 0,650086 | Q9UJZ1;A0. | STOML2 |
| -1,12673 | 0,599349 | 1,92668 | 0,599349 | 1,5310383 | 0,650421 | F2Z2X0;O0( | NME4 |
| -0,502437 | -0,451833 | 0,459856 | -0,451833 | 0,5415634 | 0,650762 | P08754 | GNAI3 |
| -2,49349 | 3,27937 | 1,97399 | 1,97399 | 3,0273285 | 0,651195 | P50479;C9J | PDLIM4 |
| -0,82084 | 0,361359 | 1,53168 | 0,361359 | 1,176265 | 0,651236 | Q9UJW0;H9 | DCTN4 |
| -1,2184 | 0,672198 | -0,3156 | -0,3156 | 0,9456174 | 0,651291 | P55060 | CSE1L |
| 0,409085 | -0,313631 | -0,551289 | -0,313631 | 0,5001863 | 0,651302 | Q9H773 | DCTPP1 |
| 0,648563 | -1,06477 | -0,368792 | -0,368792 | 0,8616754 | 0,651409 | Q5TCS8;J3I | AK9 |
| -1,02246 | 0,220284 | 0,163287 | 0,163287 | 0,701624 | 0,651553 | Q5T124;X6I | UBXN11 |
| 1,13893 | -0,238435 | -2,64289 | -0,238435 | 1,9140141 | 0,651637 | P54277;Q3I | PMS1 |
| 0,0293966 | 0,126079 | -0,420101 | 0,0293966 | 0,2914642 | 0,65245 | Q9NUQ9;E5 | FAM49B |
| -0,3273 | 0,720772 | -1,31917 | -0,3273 | 1,0201 | 0,652605 | P29762;B5I | CRABP1 |
| 1,51933 | -1,13618 | -2,07346 | -1,13618 | 1,8636151 | 0,652758 | Q0VGE8;AC | ZNF816 |
| 0,645155 | -1,80208 | 3,61561 | 0,645155 | 2,7130526 | 0,653014 | J3KNN5;Q9 | DDX41 |
| -0,507228 | 0,209062 | 0,963889 | 0,209062 | 0,7356426 | 0,653448 | Q9C0H2 | TTYH3 |
| -2,54929 | -0,967626 | 1,61795 | -0,967626 | 2,1036775 | 0,654211 | Q9HB75 | PIDD1 |
| 2,61828 | -2,47731 | 2,47739 | 2,47739 | 2,9021239 | 0,654369 | Q9NR31;Q5 | SAR1A |
| 0,398019 | -0,451723 | -0,368637 | -0,368637 | 0,4684596 | 0,654596 | Q9Y5S2;A0. | CDC42BPB |
| -0,266301 | -0,990249 | 0,558486 | -0,266301 | 0,7749144 | 0,65484 | O94826 | TOMM70 |
| -1,66133 | 1,47942 | 1,94807 | 1,47942 | 1,962639 | 0,655157 | Q8TB72 | PUM2 |
| 0,9337 | -1,5199 | -0,523658 | -0,523658 | 1,2340005 | 0,655323 | Q9ULB1;A0 | NRXN1 |
| 0,349965 | -0,657456 | 1,10095 | 0,349965 | 0,8823139 | 0,655358 | P00441;H7I | SOD1 |
| -0,977541 | 1,23529 | 0,794758 | 0,794758 | 1,1713051 | 0,6556 | J3QKW1 | AKT2 |
| -2,57668 | 0,631458 | 0,34968 | 0,34968 | 1,7764727 | 0,655744 | Q9BT78;D6 | COPS4 |
| 1,71026 | -0,403767 | -0,248456 | -0,248456 | 1,1782614 | 0,655807 | P17677 | GAP43 |
| -1,22829 | 1,56934 | 0,981687 | 0,981687 | 1,475131 | 0,656237 | Q8N7K0;C9 | ZNF433 |

| | | | | | | | |
|---|---|---|---|---|---|---|---|
| 1,18494 | -0,116305 | -2,9772 | -0,116305 | 2,1292161 | 0,656345 | Q15643;H0 | TRIP11 |
| -1,33297 | -0,908993 | 1,0861 | -0,908993 | 1,2917722 | 0,656881 | P07858;E9F | CTSB |
| 1,91983 | -1,14953 | -2,98205 | -1,14953 | 2,47681 | 0,657492 | A0A0A0MC | KSR1 |
| -3,34391 | 3,43807 | -3,67796 | -3,34391 | 4,0154851 | 0,657658 | Q8N8L2 | ZNF491 |
| 2,41464 | -1,81939 | -3,20837 | -1,81939 | 2,9290077 | 0,657772 | P04264 | KRT1 |
| 1,02039 | 0,341345 | -3,56717 | 0,341345 | 2,4759947 | 0,658255 | Q9H857;H7 | NT5DC2 |
| 1,96468 | -1,84397 | 1,79271 | 1,79271 | 2,1510009 | 0,658656 | Q8NFI3;F8V | ENGASE |
| 0,560451 | 0,735614 | -3,35172 | 0,560451 | 2,3109184 | 0,658658 | A0A2R8Y85 | CUX1 |
| 2,97313 | 0,18895 | -1,2522 | 0,18895 | 2,1479442 | 0,658785 | Q15717 | ELAVL1 |
| -0,214028 | 0,60092 | -1,17873 | -0,214028 | 0,8908745 | 0,658899 | A5PLK6;H3I | RGSL1 |
| 2,98001 | 0,800536 | -1,70213 | 0,800536 | 2,3429283 | 0,659485 | A0A0A0MS | TBXAS1 |
| -2,60547 | 3,07213 | 2,25346 | 2,25346 | 3,0690542 | 0,659756 | G3V0E4;O7 | PMPCB |
| 2,17151 | -1,8214 | -2,627 | -1,8214 | 2,569631 | 0,659834 | Q96PE2 | ARHGEF17 |
| 1,19586 | 2,13805 | -1,60776 | 1,19586 | 1,9484652 | 0,659893 | F6SXL5 | DDX39B |
| 2,02962 | -1,91544 | 1,86059 | 1,86059 | 2,2304884 | 0,660078 | A0A075B78 | TNPO2 |
| 0,766482 | 0,516386 | -3,30097 | 0,516386 | 2,2795804 | 0,660098 | Q9Y5E6;Q9 | PCDHB3 |
| -0,00923756 | 0,593168 | -1,5735 | -0,009238 | 1,1183514 | 0,66025 | P25786;F5C | PSMA1 |
| 1,12283 | -2,37698 | -0,301964 | -0,301964 | 1,7599432 | 0,660476 | Q8IWR0;I3I | ZC3H7A |
| 2,41203 | 3,35061 | -2,82211 | 2,41203 | 3,3261496 | 0,660516 | O60296;Q5 | TRAK2 |
| 0,717107 | -1,20366 | 1,85192 | 0,717107 | 1,544545 | 0,660542 | O95758 | PTBP3 |
| 0,768908 | 0,566063 | -3,42345 | 0,566063 | 2,3640793 | 0,660735 | Q96JJ6;A0A | JPH4 |
| 1,44797 | -0,293113 | -0,272739 | -0,272739 | 0,9993852 | 0,660993 | Q9NVI7;Q5 | ATAD3A |
| -1,37251 | 1,74673 | 1,07388 | 1,07388 | 1,6415038 | 0,661159 | Q13561;F8Y | DCTN2 |
| -1,73665 | 0,586479 | 0,0753795 | 0,0753795 | 1,2207649 | 0,661755 | A0A1B0GW | SPEF2 |
| -2,95114 | 1,11318 | -0,00621087 | -0,006211 | 2,0993785 | 0,662431 | Q9UMS4;F5 | PRPF19 |
| -3,33634 | 1,76269 | -0,661753 | -0,661753 | 2,5505374 | 0,663109 | C9JFW8;Q9 | NAALADL1 |
| -0,705787 | 0,566983 | 0,871145 | 0,566983 | 0,8365776 | 0,663465 | P52565;J3K | ARHGDIA |
| -2,58312 | 1,47981 | -0,675122 | -0,675122 | 2,0327153 | 0,663633 | Q96T76;Q5 | MMS19 |
| 6,77243 | 1,36297 | -3,599 | 1,36297 | 5,1873237 | 0,663766 | Q9Y2X7;J3C | GIT1 |
| 0,184872 | -1,16049 | 0,273629 | 0,184872 | 0,8035934 | 0,664092 | O60240;H0 | PLIN1 |
| -2,76873 | -0,507854 | 1,43825 | -0,507854 | 2,1054517 | 0,664239 | P14174 | MIF |
| 1,25562 | -0,25694 | -2,77674 | -0,25694 | 2,0370386 | 0,664329 | Q99873;E9I | PRMT1 |
| 1,35021 | -0,0413653 | -3,47626 | -0,041365 | 2,4842771 | 0,664471 | Q9C093;A0 | SPEF2 |
| 0,590778 | -0,824108 | -0,399471 | -0,399471 | 0,7260409 | 0,664769 | Q12768;E5I | WASHC5 |
| -3,03967 | 2,16067 | -1,43691 | -1,43691 | 2,6631735 | 0,665443 | O60522 | TDRD6 |
| 0,448424 | -0,587093 | 0,746688 | 0,448424 | 0,7000272 | 0,665798 | Q5JSZ5;Q5J | PRRC2B |
| 0,748837 | 3,69619 | -1,98041 | 0,748837 | 2,8389983 | 0,665947 | O15090;K7I | ZNF536 |
| 0,43554 | -2,48052 | 0,553548 | 0,43554 | 1,7186671 | 0,666065 | P27348;E9F | YWHAQ |
| -1,88126 | 0,762148 | -0,0539738 | -0,053974 | 1,3535531 | 0,666446 | P09936;D6I | UCHL1 |
| -0,529877 | -1,3841 | 0,909549 | -0,529877 | 1,1592001 | 0,666509 | E7EN40 | HNRNPH1 |
| -2,75661 | 2,88016 | -3,00208 | -2,75661 | 3,3275161 | 0,666995 | Q96T23;H0 | RSF1 |
| -1,32003 | -2,32965 | 1,79198 | -1,32003 | 2,1483239 | 0,667113 | E7EQ29;P1 | GLB1 |
| 0,0203178 | 0,476378 | -1,28905 | 0,0203178 | 0,9164398 | 0,667154 | Q86XL3;F5I | ANKLE2 |
| 0,4103 | -0,172446 | -0,731219 | -0,172446 | 0,5708015 | 0,667245 | P50502;Q3I | ST13 |
| 0,578714 | -0,915447 | 1,32009 | 0,578714 | 1,1386967 | 0,667503 | Q9H799 | CPLANE1 |
| 0,742698 | -2,80619 | 0,380511 | 0,380511 | 1,952812 | 0,668105 | A0A286YFK | PHGDH |
| 1,28297 | -0,990947 | -1,60153 | -0,990947 | 1,5200795 | 0,668227 | O43865 | AHCYL1 |
| 1,18795 | -0,431625 | -2,22723 | -0,431625 | 1,7083459 | 0,668384 | P14649;F8V | MYL6B |
| -2,04946 | 0,112733 | 0,692983 | 0,112733 | 1,445266 | 0,668537 | Q15942;H0 | ZYX |
| 0,122469 | 0,310738 | -1,08491 | 0,122469 | 0,7573025 | 0,668539 | Q8WXA3;H | RUFY2 |
| 2,43434 | -2,94258 | -1,95186 | -1,95186 | 2,8615714 | 0,668833 | F5GX29;H0 | CHP1 |
| -2,62981 | -0,72803 | 1,55606 | -0,72803 | 2,0958428 | 0,668837 | O95865;A0 | DDAH2 |

| | | | | | | | |
|---|---|---|---|---|---|---|---|
| 1,34049 | 0,706702 | -1,00445 | 0,706702 | 1,2130177 | 0,668858 | O75534;E9I | CSDE1 |
| 0,375011 | -0,0382937 | -0,89172 | -0,038294 | 0,6459831 | 0,669018 | P83881;H0Y | RPL36A |
| -1,78839 | 0,82133 | -0,164317 | -0,164317 | 1,3178108 | 0,669235 | Q3SXM5;H3 | HSDL1 |
| 1,60356 | -0,986273 | 0,498291 | 0,498291 | 1,2995374 | 0,669265 | Q02386;K7I | ZNF45 |
| 5,47861 | -3,26601 | 1,54613 | 1,54613 | 4,3796779 | 0,669341 | E5RJ52;P53 | AP3M2 |
| -2,28085 | 1,20492 | -0,419727 | -0,419727 | 1,7442214 | 0,669591 | Q8NBT2 | SPC24 |
| 0,0921636 | -0,592077 | 1,33409 | 0,0921636 | 0,9764464 | 0,670689 | Q96A35 | MRPL24 |
| -0,736998 | 0,290399 | 1,31941 | 0,290399 | 1,0282041 | 0,672556 | Q12955;A0 | ANK3 |
| -2,2388 | -1,67279 | 1,97429 | -1,67279 | 2,2866164 | 0,673118 | Q9BZF1;F8V | OSBPL8 |
| 1,7266 | 0,448951 | -1,01379 | 0,448951 | 1,3712364 | 0,673118 | A0A2R8Y54 | PATJ |
| 1,12933 | 1,54815 | -1,35066 | 1,12933 | 1,5667833 | 0,673253 | P31153 | MAT2A |
| -2,15433 | -0,135558 | 0,954373 | -0,135558 | 1,5773091 | 0,673301 | Q9Y5H2 | PCDHGA11 |
| 0,752024 | -3,26043 | 0,58853 | 0,58853 | 2,2708666 | 0,673738 | Q07960 | ARHGAP1 |
| -2,45243 | 2,6498 | -2,76279 | -2,45243 | 3,0393312 | 0,674209 | H0YLC4;H0Y | ZNF710 |
| -1,37811 | 1,04296 | -0,721194 | -0,721194 | 1,2520222 | 0,674332 | Q9H1I8 | ASCC2 |
| -1,49133 | -1,00139 | 1,25672 | -1,00139 | 1,4657695 | 0,674493 | A0A1B0GTJ | ARID1B |
| 4,19616 | 2,93879 | -3,61439 | 2,93879 | 4,1938415 | 0,675802 | A0A1W2PR | GOSR2 |
| -1,08386 | 0,777076 | 1,38573 | 0,777076 | 1,2866246 | 0,676104 | F8VU56;P2 | PTPRB |
| 0,404607 | -2,30594 | 0,553672 | 0,404607 | 1,6096928 | 0,676577 | Q96HP0;K7 | DOCK6 |
| 2,80399 | -2,57187 | -2,9113 | -2,57187 | 3,2062341 | 0,677131 | J3KQ21;Q9I | ANKMY1 |
| -1,02065 | 0,955447 | -0,848236 | -0,848236 | 1,0945287 | 0,677499 | A0A2Q3DQ | CAMK2G |
| 0,29863 | 1,2389 | -0,72014 | 0,29863 | 0,9797821 | 0,677603 | Q15024 | EXOSC7 |
| -1,10198 | 0,767237 | 1,42381 | 0,767237 | 1,3105136 | 0,678723 | A0A087WU | PIBF1 |
| 0,12664 | -0,210485 | 0,299252 | 0,12664 | 0,2592553 | 0,678776 | Q8IWS0;A0 | PHF6 |
| -2,40407 | -0,322293 | 1,21715 | -0,322293 | 1,817366 | 0,678925 | Q9H9B4;D6 | SFXN1 |
| -1,02771 | 0,270183 | 2,03277 | 0,270183 | 1,5361086 | 0,679015 | Q00688 | FKBP3 |
| -0,646064 | 0,18426 | 0,0853473 | 0,0853473 | 0,4535387 | 0,679062 | D6R9L5 | DEK |
| 3,01322 | -3,10353 | -2,76202 | -2,76202 | 3,4371659 | 0,67913 | Q9UQ53 | MGAT4B |
| -0,864997 | 0,573432 | -0,310422 | -0,310422 | 0,7254687 | 0,67915 | E9PK91;Q9I | BCLAF1 |
| 4,05686 | 1,60422 | -2,78543 | 1,60422 | 3,4665401 | 0,679236 | Q9H0W5 | CCDC8 |
| -2,67525 | 1,01384 | 0,0715472 | 0,0715472 | 1,9166897 | 0,679256 | Q86VS8 | HOOK3 |
| -0,127668 | -0,0441477 | 0,413339 | -0,044148 | 0,2912497 | 0,679331 | Q9Y657 | SPIN1 |
| 1,4598 | -0,0458064 | -0,5486 | -0,045806 | 1,0450935 | 0,679753 | F2Z3H1 | UGP2 |
| -1,54259 | 1,4906 | 1,50492 | 1,4906 | 1,7553615 | 0,67987 | P42574;A8I | CASP3 |
| 1,28325 | -0,36611 | -0,171319 | -0,171319 | 0,9013049 | 0,679949 | Q99973;G3 | TEP1 |
| 1,24762 | -1,43975 | -0,998017 | -0,998017 | 1,4410627 | 0,680504 | Q8TAT6 | NPLOC4 |
| -1,88275 | 0,0933085 | 0,6821 | 0,0933085 | 1,343499 | 0,681083 | O15540 | FABP7 |
| -1,92568 | 0,562959 | 0,246263 | 0,246263 | 1,3546806 | 0,681108 | P61313;A0 | RPL15 |
| -0,512092 | 0,974694 | -1,48212 | -0,512092 | 1,2374316 | 0,681196 | P21281;H0Y | ATP6V1B2 |
| -0,0817551 | -1,87492 | 0,825938 | -0,081755 | 1,3744078 | 0,681609 | P20339 | RAB5A |
| 2,08646 | -0,19546 | -0,676753 | -0,19546 | 1,476152 | 0,681655 | B2RPK0 | HMGB1P1 |
| 2,77644 | 4,78642 | -3,85291 | 2,77644 | 4,5208097 | 0,68233 | A1A4V9;H3 | CCDC189 |
| 1,95505 | 2,42351 | -2,26227 | 1,95505 | 2,580755 | 0,682541 | H0YI59 | WDR90 |
| 2,66805 | -0,651667 | -0,487598 | -0,487598 | 1,8710761 | 0,683576 | Q9GZS3;H0 | WDR61 |
| -0,799319 | 0,434 | 1,1821 | 0,434 | 1,0005624 | 0,683833 | Q7Z3V4;S4I | UBE3B |
| -1,2676 | -0,509074 | 0,885391 | -0,509074 | 1,0920367 | 0,683887 | Q4G0M1 | ERFE |
| 0,741605 | -0,528783 | -0,922329 | -0,528783 | 0,8696207 | 0,683986 | Q86XU0;M0 | ZNF677 |
| -0,705334 | 2,41283 | -0,321308 | -0,321308 | 1,7002908 | 0,684201 | A0A087WT | GPX4 |
| -2,5755 | 1,03685 | 0,0225171 | 0,0225171 | 1,8631352 | 0,684728 | Q7Z794 | KRT77 |
| 0,92888 | 0,220063 | -2,72549 | 0,220063 | 1,9379166 | 0,684808 | Q9Y4D1 | DAAM1 |
| 1,20645 | 0,607992 | -0,921915 | 0,607992 | 1,0976266 | 0,684943 | O60673 | REV3L |
| 0,0588421 | 0,376127 | -1,03339 | 0,0588421 | 0,7394118 | 0,686276 | Q9Y4E6;A2 | WDR7 |

| | | | | | | | |
|---|---|---|---|---|---|---|---|
| 1,46586 | -1,13856 | 0,76307 | 0,76307 | 1,347412 | 0,686308 | A0A087WT | KCNE1B |
| 0,467698 | -1,93218 | 0,366603 | 0,366603 | 1,3573281 | 0,686441 | O75891;C9 | ALDH1L1 |
| 1,4675 | -0,0735613 | -0,543165 | -0,073561 | 1,0518374 | 0,686443 | P20648 | ATP4A |
| 1,447 | -0,98438 | 0,529507 | 0,529507 | 1,2278203 | 0,686724 | Q9NYV4;J3 | CDK12 |
| -0,933438 | 1,32861 | 0,531141 | 0,531141 | 1,1473019 | 0,686954 | P62745 | RHOB |
| 0,468206 | 1,51654 | -0,974897 | 0,468206 | 1,2509202 | 0,686988 | O43813;E9I | LANCL1 |
| 3,61292 | -2,88219 | 1,99389 | 1,99389 | 3,3809236 | 0,68748 | P13646;K7E | KRT13 |
| 1,6781 | -0,478632 | -3,14643 | -0,478632 | 2,4167723 | 0,687576 | H3BMF4;Q | SPNS1 |
| 0,44586 | -1,38738 | 0,148969 | 0,148969 | 0,9839784 | 0,687628 | P46013 | MKI67 |
| 3,28596 | 1,49996 | -2,43386 | 1,49996 | 2,926348 | 0,688224 | Q96EY1;I3L | DNAJA3 |
| 0,551651 | 0,448819 | -2,29845 | 0,448819 | 1,6166394 | 0,688528 | Q96CN4 | EVI5L |
| 0,95942 | 0,136178 | -2,58233 | 0,136178 | 1,8534655 | 0,688791 | F5GX23;F5H | PSMD9 |
| -0,901247 | 0,317368 | 1,57384 | 0,317368 | 1,2375918 | 0,689572 | Q9UL62 | TRPC5 |
| -1,51391 | 2,34238 | -3,05066 | -1,51391 | 2,7784124 | 0,689607 | F5H101;F8V | NOL8 |
| 0,240528 | -0,841609 | 0,125746 | 0,125746 | 0,5944144 | 0,689655 | Q9UHF7;E5 | TRPS1 |
| -0,989022 | 1,96952 | -2,9656 | -0,989022 | 2,4837888 | 0,689813 | Q9H972;G3 | C14orf93 |
| -1,4973 | -0,570913 | 1,04297 | -0,570913 | 1,2855468 | 0,690414 | P63267;P62 | ACTG2 |
| 5,8551 | -0,16751 | -2,31719 | -0,16751 | 4,2363365 | 0,691086 | J3KTL3 | MED24 |
| -1,74914 | -2,47064 | 2,21426 | -1,74914 | 2,5224786 | 0,691274 | J3KQW3 | TMEM44 |
| -2,64294 | 1,40054 | -0,368904 | -0,368904 | 2,0269806 | 0,69132 | Q9UL03 | INTS6 |
| -0,130942 | 0,94247 | -0,281294 | -0,130942 | 0,6673851 | 0,691472 | P12955 | PEPD |
| 0,278763 | 0,253218 | -1,20807 | 0,253218 | 0,8511451 | 0,691529 | O94856;X6 | NFASC |
| -0,688932 | 0,853934 | -0,935137 | -0,688932 | 0,969693 | 0,691574 | Q96MU7;J3 | YTHDC1 |
| 0,645963 | -0,151994 | -0,132537 | -0,132537 | 0,4551879 | 0,691636 | P09651;F8V | HNRNPA1 |
| -0,636427 | 0,53765 | 0,668736 | 0,53765 | 0,7186899 | 0,691979 | Q9H9A6 | LRRC40 |
| -1,66938 | -3,38246 | 2,60625 | -1,66938 | 3,0843774 | 0,692034 | P25325;B1A | MPST |
| 3,06102 | -2,83423 | -2,94979 | -2,83423 | 3,4374691 | 0,692296 | B1AKL4;Q9 | EIF4ENIF1 |
| 2,32159 | -1,24908 | -3,33774 | -1,24908 | 2,8618235 | 0,692513 | Q92922 | SMARCC1 |
| -0,744597 | -1,51667 | 1,16754 | -0,744597 | 1,3818676 | 0,692529 | P18615;A0A | NELFE |
| -1,54651 | 3,44824 | 0,109473 | 0,109473 | 2,544182 | 0,692873 | Q05397;E7 | PTK2 |
| -1,10926 | -1,65689 | 1,457 | -1,10926 | 1,6624229 | 0,693933 | P56524;F5H | HDAC4 |
| 0,927717 | -0,204312 | -0,208087 | -0,204312 | 0,6546697 | 0,694059 | Q13409;E7 | DYNC1I2 |
| -0,517869 | -0,978181 | 3,38005 | -0,517869 | 2,3944323 | 0,694171 | Q96BY6;A0 | DOCK10 |
| -2,9175 | 3,30545 | 2,22799 | 2,22799 | 3,3257103 | 0,694256 | Q96MW5;H | COG8 |
| -0,161096 | -0,670906 | 1,90465 | -0,161096 | 1,363861 | 0,694293 | Q9BUT1;D6 | BDH2 |
| -0,596632 | 0,582908 | 0,539417 | 0,539417 | 0,6688066 | 0,694455 | Q15772;B9 | SPEG |
| -3,04759 | 1,41314 | -0,144861 | -0,144861 | 2,2638947 | 0,694479 | Q9Y5G1 | PCDHGB3 |
| -1,57559 | -2,27421 | 2,0331 | -1,57559 | 2,3116961 | 0,694506 | P09132 | SRP19 |
| -0,176435 | -0,485442 | 0,336228 | -0,176435 | 0,4150201 | 0,694935 | O75489 | NDUFS3 |
| -0,440243 | 1,86975 | -0,394819 | -0,394819 | 1,3207576 | 0,695376 | P15907;C9J | ST6GAL1 |
| -2,22616 | 2,49213 | 1,71534 | 1,71534 | 2,5298579 | 0,69546 | Q96K76 | USP47 |
| -1,82752 | -0,82146 | 1,36917 | -0,82146 | 1,6345154 | 0,695525 | O94933;C9 | SLITRK3 |
| 0,168156 | -0,0190597 | -0,0552832 | -0,01906 | 0,1199215 | 0,69577 | P27824;D6 | CANX |
| -3,11167 | 3,10338 | -2,71214 | -2,71214 | 3,478667 | 0,69586 | A0A087X0K | CAB39 |
| 0,974826 | 1,30382 | -1,21049 | 0,974826 | 1,3666015 | 0,69601 | P09429;Q5 | HMGB1 |
| 1,94556 | -1,17621 | -2,57699 | -1,17621 | 2,3152067 | 0,696307 | Q12931;I3L | TRAP1 |
| 0,862035 | -0,917374 | 0,855942 | 0,855942 | 1,0255879 | 0,696356 | P07954 | FH |
| -0,0558535 | -2,63295 | 1,17279 | -0,055854 | 1,9422774 | 0,69639 | D3YTB1;F8V | RPL32 |
| -1,16245 | 3,12139 | -0,207665 | -0,207665 | 2,2489074 | 0,69703 | E7EQM8;P4 | DCC |
| 0,198535 | -0,0385673 | -0,0505944 | -0,038567 | 0,1404918 | 0,697107 | Q05707;J3 | COL14A1 |
| -1,80318 | 2,42027 | -2,76023 | -1,80318 | 2,7565393 | 0,69747 | E9PKF6;H7 | PPP6R3 |
| 1,41521 | -0,638315 | -2,1772 | -0,638315 | 1,8023384 | 0,697661 | Q15147;A0 | PLCB4 |

| | | | | | | | |
|---|---|---|---|---|---|---|---|
| 1,63341 | -1,90224 | -1,18052 | -1,18052 | 1,8681498 | 0,698057 | O43683;C9I | BUB1 |
| 1,60584 | -1,46905 | -1,51973 | -1,46905 | 1,790098 | 0,699212 | Q9NQC3;A( | RTN4 |
| -0,0482927 | -1,29042 | 3,07091 | -0,048293 | 2,2469796 | 0,699797 | O43909;E7I | EXTL3 |
| 2,25413 | -1,01542 | 0,0463132 | 0,0463132 | 1,6679173 | 0,699961 | Q9H1K0 | RBSN |
| -0,690173 | 1,86692 | -0,139968 | -0,139968 | 1,3459228 | 0,700006 | P56537;B7Z | EIF6 |
| -0,268092 | 0,44092 | -0,573499 | -0,268092 | 0,5204193 | 0,700152 | Q4V328;A0 | GRIPAP1 |
| 0,316043 | -0,661439 | 0,981574 | 0,316043 | 0,8264275 | 0,70019 | Q9H582;A0 | ZNF644 |
| -1,4023 | 2,162 | -2,69145 | -1,4023 | 2,5140313 | 0,700688 | Q6PUV4;D6 | CPLX2 |
| 2,24613 | -3,10554 | -1,22227 | -1,22227 | 2,7146785 | 0,701243 | Q6PEY2 | TUBA3E |
| -1,17448 | 0,0673509 | 0,455455 | 0,0673509 | 0,8514162 | 0,701748 | P47897;A0/ | QARS |
| -2,57855 | 0,364065 | 0,806319 | 0,364065 | 1,8399234 | 0,70177 | Q7RTS9 | DYM |
| 0,714665 | -0,649705 | 0,495005 | 0,495005 | 0,7325887 | 0,702116 | Q8NB25;A0 | FAM184A |
| 0,497078 | -1,50415 | 0,184275 | 0,184275 | 1,0765329 | 0,702138 | P52758;H0Y | RIDA |
| 0,569735 | 0,655828 | -2,67798 | 0,569735 | 1,9004096 | 0,702151 | Q5TB80 | CEP162 |
| -2,34988 | -1,20312 | 1,88127 | -1,20312 | 2,1882699 | 0,702263 | P42704;A0/ | LRPPRC |
| 2,22735 | -2,37325 | -1,76245 | -1,76245 | 2,4985696 | 0,702326 | Q9H1A4;H0 | ANAPC1 |
| -0,235374 | 1,02292 | -1,91159 | -0,235374 | 1,4722066 | 0,702418 | A0A1B0GU | LRMP |
| 0,495479 | -1,42279 | 0,147267 | 0,147267 | 1,0219333 | 0,702493 | Q6S8J3;A0/ | POTEE |
| 3,22109 | -1,43577 | 0,0313627 | 0,0313627 | 2,3809376 | 0,702596 | P28482;K7E | MAPK1 |
| 0,0136684 | 0,507664 | -1,18574 | 0,0136684 | 0,8708453 | 0,702621 | Q9Y262;B0 | EIF3L |
| -0,0966573 | -0,43783 | 0,265999 | -0,096657 | 0,3519691 | 0,702665 | P53990;H3I | IST1 |
| 3,43397 | -3,40168 | -2,94721 | -2,94721 | 3,822131 | 0,702727 | Q8N568;A0 | DCLK2 |
| 4,30212 | -0,82253 | -1,14852 | -0,82253 | 3,0571714 | 0,702781 | P17213;A0/ | BPI |
| -0,527873 | 0,0198188 | 1,16606 | 0,0198188 | 0,8644116 | 0,703234 | Q9NVH0;C9 | EXD2 |
| -0,103488 | 2,66287 | -1,08165 | -0,103488 | 1,9421194 | 0,703351 | O75937 | DNAJC8 |
| 3,60951 | -3,35221 | 2,60564 | 2,60564 | 3,763183 | 0,703391 | Q9H2G2 | SLK |
| 0,882887 | -0,216164 | -1,6215 | -0,216164 | 1,2553112 | 0,703457 | O75197;E9I | LRP5 |
| 1,07861 | -2,17446 | -0,147397 | -0,147397 | 1,6428908 | 0,704826 | P20020;E7E | ATP2B1 |
| -1,91361 | -3,46215 | 2,87607 | -1,91361 | 3,3043417 | 0,704918 | Q9H8Y8 | GORASP2 |
| -0,838951 | 0,563196 | 0,997952 | 0,563196 | 0,9599689 | 0,706405 | F5H3U9;F5I | MAGOHB |
| 1,78352 | -2,32478 | -1,03353 | -1,03353 | 2,1008421 | 0,707374 | Q12789 | GTF3C1 |
| -1,01872 | 0,609791 | -0,201229 | -0,201229 | 0,8142576 | 0,707465 | Q9C0B0;K7 | UNK |
| -1,02412 | 1,42325 | 0,527202 | 0,527202 | 1,2382193 | 0,707901 | P07202;A0/ | TPO |
| -0,410447 | 0,93856 | -1,40804 | -0,410447 | 1,1776773 | 0,708241 | A0A087WS | ZNF43 |
| -0,215561 | -0,367494 | 1,2507 | -0,215561 | 0,8936402 | 0,708263 | Q6NVY1 | HIBCH |
| -1,77903 | -0,322434 | 1,04642 | -0,322434 | 1,412952 | 0,708409 | P04406;E7E | GAPDH |
| -2,00854 | 3,00897 | -3,56551 | -2,00854 | 3,4356792 | 0,708444 | A0A0A0MS | EZH1 |
| -1,25356 | 0,963942 | 1,33406 | 0,963942 | 1,3994094 | 0,708536 | A0A067XG | ATP11C |
| -2,44533 | 0,779769 | 0,357755 | 0,357755 | 1,752933 | 0,708635 | P19013 | KRT4 |
| -0,820661 | 0,902825 | -0,82381 | -0,820661 | 0,9959654 | 0,709141 | P22234;E9F | PAICS |
| 0,742137 | 0,611154 | -2,88411 | 0,611154 | 2,0568461 | 0,709282 | P40938 | RFC3 |
| 1,05439 | -0,542042 | 0,0853946 | 0,0853946 | 0,8042827 | 0,709661 | O43707;F5( | ACTN4 |
| 0,303081 | -0,937859 | 1,55883 | 0,303081 | 1,2483518 | 0,710727 | Q9BTU6 | PI4K2A |
| -0,464209 | -0,449532 | 0,503498 | -0,449532 | 0,5545176 | 0,71087 | Q96A46 | SLC25A28 |
| 1,27795 | -2,02974 | -0,470959 | -0,470959 | 1,6547555 | 0,711189 | A0A0B4J23 | DAD1 |
| 0,639848 | -0,473248 | 0,250774 | 0,250774 | 0,5648848 | 0,711211 | P41091 | EIF2S3 |
| -0,161821 | 0,270949 | -0,341811 | -0,161821 | 0,31495 | 0,711238 | Q96HY6;A0 | DDRGK1 |
| -0,127237 | -0,293729 | 0,8974 | -0,127237 | 0,6450309 | 0,711299 | M0R2Z9;M | SUGP2 |
| -0,602856 | 0,270075 | 0,882987 | 0,270075 | 0,7467038 | 0,711935 | P04181 | OAT |
| -2,99784 | 4,39956 | 1,33339 | 1,33339 | 3,7166849 | 0,712274 | P48378 | RFX2 |
| -2,98164 | 2,70562 | -1,95384 | -1,95384 | 3,0307278 | 0,712327 | P12271 | RLBP1 |
| 1,46365 | -1,34477 | 0,985616 | 0,985616 | 1,5025777 | 0,712573 | O15066 | KIF3B |

| | | | | | | | |
|---|---|---|---|---|---|---|---|
| -0,551088 | 1,97684 | -0,386407 | -0,386407 | 1,4143595 | 0,71265 | P84098;J3K | RPL19 |
| 0,707206 | -0,436971 | -0,868539 | -0,436971 | 0,8142854 | 0,712683 | Q5T011 | SZT2 |
| 2,54252 | 3,69997 | -3,4352 | 2,54252 | 3,8293489 | 0,71328 | A8MYR2 | MSRB1 |
| -0,663211 | -0,348894 | 2,133 | -0,348894 | 1,5317413 | 0,713753 | Q14847;C9. | LASP1 |
| 3,58853 | -2,78847 | 1,57664 | 1,57664 | 3,2600618 | 0,714739 | P11215 | ITGAM |
| -0,160836 | -0,958797 | 2,39573 | -0,160836 | 1,752409 | 0,71504 | Q7Z6G8;F8' | ANKS1B |
| 1,78036 | -2,33706 | -0,970375 | -0,970375 | 2,0971218 | 0,715046 | H0YDM0 | PPFIBP2 |
| 0,720108 | -1,70037 | 2,52212 | 0,720108 | 2,1187804 | 0,715216 | A0A1B0GTS | CACNB4 |
| 1,33442 | -3,32285 | 0,219456 | 0,219456 | 2,4317771 | 0,715312 | Q8IWK6;D6 | ADGRA3 |
| 0,126113 | -0,204254 | 0,24831 | 0,126113 | 0,2341255 | 0,715531 | P27797;K7E | CALR |
| 5,98842 | 1,56836 | -3,94428 | 1,56836 | 4,9763551 | 0,715854 | P58546 | MTPN |
| 2,38728 | 3,15526 | -3,07715 | 2,38728 | 3,3983507 | 0,716023 | R4GNB9 | TRIM11 |
| 0,354787 | 0,0274765 | -0,823511 | 0,0274765 | 0,6082349 | 0,716027 | E7EU13;Q9 | ARAP1 |
| -0,240418 | 0,60216 | -0,0421851 | -0,042185 | 0,4405327 | 0,716051 | P13637;A0/ | ATP1A3 |
| 0,454606 | 1,00242 | -3,05286 | 0,454606 | 2,2002922 | 0,716088 | P35579;Q5I | MYH9 |
| 1,50581 | -0,685433 | -2,14181 | -0,685433 | 1,836106 | 0,718103 | Q13642;Q5 | FHL1 |
| -0,50931 | 2,33199 | -0,618982 | -0,50931 | 1,6729838 | 0,718175 | Q9UBB6;H7 | NCDN |
| -1,9139 | -1,27899 | 1,77423 | -1,27899 | 1,9717829 | 0,71818 | A0A087WZ | HMGN3 |
| -0,196283 | -0,0197065 | 0,460421 | -0,019707 | 0,3398435 | 0,718263 | Q92598;A0 | HSPH1 |
| 1,99529 | -1,64064 | -1,92657 | -1,64064 | 2,1864251 | 0,718371 | B0YIW6;P4 | ARCN1 |
| -0,199955 | -0,242805 | 0,913753 | -0,199955 | 0,6557195 | 0,71861 | C9JGT8 | AP2M1 |
| -0,829589 | -1,15662 | 1,10828 | -0,829589 | 1,2242043 | 0,719023 | P09211;A8I | GSTP1 |
| 0,84414 | -0,599955 | 0,276695 | 0,276695 | 0,7275437 | 0,719455 | P61106;X6F | RAB14 |
| 0,180059 | -0,223348 | -0,104989 | -0,104989 | 0,2073638 | 0,719773 | O43765;K7I | SGTA |
| 1,24701 | -1,62135 | -0,670254 | -0,670254 | 1,4610484 | 0,719811 | A0A1W2PP | ADGRV1 |
| -2,59881 | 2,14066 | -1,28453 | -1,28453 | 2,446829 | 0,720799 | A6NK53;K7 | ZNF233 |
| -1,91444 | -0,0688443 | 0,949642 | -0,068844 | 1,4518094 | 0,720891 | P49773;D6I | HINT1 |
| 5,81241 | -2,95835 | 0,301302 | 0,301302 | 4,4332808 | 0,720972 | P54802 | NAGLU |
| 2,57123 | -2,7224 | -1,87054 | -1,87054 | 2,8424609 | 0,72115 | Q86WZ6;K7 | ZNF227 |
| 0,472419 | 0,469164 | -0,530464 | 0,469164 | 0,5780774 | 0,721174 | Q9HCE3;K7 | ZNF532 |
| -1,71932 | -2,23761 | 2,22232 | -1,71932 | 2,4391298 | 0,721184 | Q6UVM3;Q | KCNT2 |
| 0,803428 | -1,52156 | -0,114114 | -0,114114 | 1,1710648 | 0,721359 | Q9H5N1;B4 | RABEP2 |
| 1,20465 | 0,142162 | -0,676669 | 0,142162 | 0,9432856 | 0,721446 | P03951;H0' | F11 |
| -2,52391 | 1,43025 | 3,15691 | 1,43025 | 2,9122858 | 0,722159 | F5GZG1 | RAP1B |
| 2,01495 | -3,34157 | 4,02076 | 2,01495 | 3,8061241 | 0,722383 | O95395;H0 | GCNT3 |
| 1,00295 | 0,0504666 | -2,22769 | 0,0504666 | 1,660033 | 0,722551 | Q8TF20 | ZNF721 |
| -3,13634 | 1,02097 | 0,512038 | 0,512038 | 2,2676309 | 0,722669 | Q8WXG6;A | MADD |
| 3,41356 | -3,1225 | -2,91649 | -2,91649 | 3,7155541 | 0,722831 | Q9UM47 | NOTCH3 |
| 2,0421 | -2,33866 | 2,09298 | 2,0421 | 2,544048 | 0,723004 | Q86XK2 | FBXO11 |
| -1,06509 | 1,88708 | -2,35792 | -1,06509 | 2,1758807 | 0,723091 | Q8N806;G3 | UBR7 |
| -2,95517 | 4,89638 | 0,830425 | 0,830425 | 3,9266092 | 0,723102 | H0YME5;Q | EIF2AK4 |
| -0,942187 | 0,887205 | 0,77957 | 0,77957 | 1,0265401 | 0,723104 | Q13148;A0 | TARDBP |
| 2,56734 | -2,39542 | 1,69719 | 1,69719 | 2,650019 | 0,723298 | Q9Y3P9;B5 | RABGAP1 |
| 0,888589 | -0,544143 | 0,160227 | 0,160227 | 0,7163995 | 0,72361 | A4D0V7;E7 | CPED1 |
| 1,39774 | 0,51673 | -1,0437 | 0,51673 | 1,2363757 | 0,723669 | A0A087WV | POLD2 |
| 5,99101 | 0,298374 | -3,07246 | 0,298374 | 4,5810318 | 0,724418 | Q96QS3 | ARX |
| -1,42975 | 0,718503 | 1,89334 | 0,718503 | 1,685139 | 0,724689 | P11177;F8V | PDHB |
| 1,54298 | 3,22918 | -2,6509 | 1,54298 | 3,0278496 | 0,725015 | F8WEZ9;Q8 | ZNF480 |
| -2,80151 | -0,508361 | 1,72526 | -0,508361 | 2,2634502 | 0,725194 | Q0PNE2 | ELP6 |
| 0,24948 | 0,175615 | -0,858045 | 0,175615 | 0,6192093 | 0,725517 | Q9BV73;E7 | CEP250 |
| -0,443697 | 0,639209 | 0,183434 | 0,183434 | 0,5437079 | 0,726327 | P39019;M0 | RPS19 |
| -0,690982 | 2,04265 | -3,16725 | -0,690982 | 2,6060092 | 0,726427 | Q06210 | GFPT1 |

| | | | | | | | |
|---|---|---|---|---|---|---|---|
| 4,00709 | -0,221975 | -1,71788 | -0,221975 | 2,9692312 | 0,726599 | Q1X8D7;J3( | LRRC36 |
| -0,162903 | -1,82642 | 1,00088 | -0,162903 | 1,4209917 | 0,726823 | P60953;Q5. | CDC42 |
| 0,22125 | 1,31238 | -3,14932 | 0,22125 | 2,3258717 | 0,727166 | Q92783;A6 | STAM |
| 1,9805 | 1,05673 | -1,70617 | 1,05673 | 1,9182676 | 0,727447 | Q96ST3;H3 | SIN3A |
| -0,961647 | 0,865289 | 0,818955 | 0,818955 | 1,0416642 | 0,727516 | H3BNV7;J3I | CIAO2B |
| 0,406388 | -0,0551318 | -0,757462 | -0,055132 | 0,5860624 | 0,727729 | P0CG39 | POTEJ |
| -0,513793 | 1,1351 | -0,0337987 | -0,033799 | 0,8480926 | 0,727863 | Q9NQ66;A( | PLCB1 |
| 1,09846 | -1,41365 | -0,570125 | -0,570125 | 1,278437 | 0,727952 | Q9HAV7 | GRPEL1 |
| 0,0770903 | -2,64985 | 1,20205 | 0,0770903 | 1,9806931 | 0,72812 | Q8N1W1 | ARHGEF28 |
| -1,3245 | 0,940149 | -0,403767 | -0,403767 | 1,1388953 | 0,728132 | D6R9L0 | RACK1 |
| 3,90371 | -3,04789 | -3,79182 | -3,04789 | 4,2445918 | 0,728241 | J3KPJ3;Q8N | CAMKK1 |
| -0,0362288 | -0,877004 | 1,8961 | -0,036229 | 1,4219062 | 0,728412 | Q5SQI0 | ATAT1 |
| -0,472761 | 2,45772 | -0,755525 | -0,472761 | 1,7791673 | 0,728491 | Q5TFQ8;P7 | SIRPB1 |
| -2,61241 | 3,73453 | 1,0801 | 1,0801 | 3,1875873 | 0,728542 | P68036;A0/ | UBE2L3 |
| -0,158101 | -1,49579 | 0,84416 | -0,158101 | 1,1739751 | 0,728958 | Q9P2F5;H0 | STOX2 |
| 2,55722 | -2,90005 | 2,49738 | 2,49738 | 3,1336248 | 0,72975 | Q9H0H5 | RACGAP1 |
| 1,82487 | -0,700935 | -0,206704 | -0,206704 | 1,3386092 | 0,730605 | O15381;E9I | NVL |
| -0,458168 | -0,00046897 | 0,951009 | -0,000469 | 0,7188624 | 0,730707 | Q96HE7;G3 | ERO1A |
| 1,84803 | -1,15213 | -2,10978 | -1,15213 | 2,0648775 | 0,730782 | A0A0G2JRX | PRAME |
| 7,58746 | -3,59376 | -0,0813454 | -0,081345 | 5,717915 | 0,730964 | Q9H313;A0 | TTYH1 |
| 0,57483 | -0,950097 | -0,144924 | -0,144924 | 0,7628621 | 0,731816 | P26599;A0/ | PTBP1 |
| 0,679953 | -0,856151 | -0,358117 | -0,358117 | 0,7837137 | 0,73186 | P51116;I3L | FXR2 |
| 0,890045 | 0,638918 | -3,01947 | 0,638918 | 2,1882708 | 0,732093 | Q96EI5;A2F | TCEAL4 |
| 0,109053 | -1,17394 | 2,23859 | 0,109053 | 1,7236763 | 0,732168 | Q14667;K7I | KIAA0100 |
| -0,579041 | -1,03757 | 0,919723 | -0,579041 | 1,0236811 | 0,732227 | Q8NI08 | NCOA7 |
| 1,88618 | -2,62371 | 2,68449 | 1,88618 | 2,8622076 | 0,732424 | O14511 | NRG2 |
| -0,189325 | 0,0472087 | 0,312697 | 0,0472087 | 0,2511501 | 0,7328 | P98196;E9F | ATP11A |
| -0,163812 | 1,06727 | -0,374599 | -0,163812 | 0,778779 | 0,73284 | O75914;A0. | PAK3 |
| -1,10072 | 1,75912 | -1,98638 | -1,10072 | 1,957543 | 0,733095 | A0A2R8YG4 | ARHGEF7 |
| -1,91938 | 1,69921 | -1,0597 | -1,0597 | 1,8905391 | 0,733609 | P62081;A0/ | RPS7 |
| 0,0137446 | 0,451704 | -0,231573 | 0,0137446 | 0,346135 | 0,734087 | A0A087WT | ALMS1 |
| 3,53338 | -2,65221 | -3,46581 | -2,65221 | 3,8277959 | 0,734251 | Q14145 | KEAP1 |
| 0,520402 | -3,13791 | 1,08177 | 0,520402 | 2,2914362 | 0,736089 | Q96M86 | DNHD1 |
| -0,194639 | 0,846361 | -0,240154 | -0,194639 | 0,6145822 | 0,736286 | Q9Y6V0;A0 | PCLO |
| -2,73834 | 1,27711 | 3,61137 | 1,27711 | 3,2117344 | 0,736361 | Q15274;C9. | QPRT |
| 2,90915 | -2,38317 | -2,61645 | -2,38317 | 3,125042 | 0,736553 | H3BUX2;O4 | CYB5B |
| 0,950466 | 1,4034 | -1,36242 | 0,950466 | 1,4834841 | 0,73678 | Q2M389;A( | WASHC4 |
| -1,92118 | -0,269531 | 4,37021 | -0,269531 | 3,2618177 | 0,73683 | O75131;A0. | CPNE3 |
| -0,588902 | 1,24304 | -1,6229 | -0,588902 | 1,4513658 | 0,737086 | O60884;A0. | DNAJA2 |
| -1,77377 | 1,05377 | 2,0373 | 1,05377 | 1,9784914 | 0,737702 | Q9UII2 | ATP5IF1 |
| 0,854948 | 0,0783115 | -0,485282 | 0,0783115 | 0,6729312 | 0,737738 | Q9H307 | PNN |
| 0,527017 | -1,55922 | 0,27429 | 0,27429 | 1,1385674 | 0,737752 | P05455;B5I | SSB |
| 0,400617 | 0,12199 | -0,291226 | 0,12199 | 0,3480965 | 0,738106 | P35222;A0/ | CTNNB1 |
| -1,08404 | 2,02977 | -2,47902 | -1,08404 | 2,3083533 | 0,738279 | Q16822;H0 | PCK2 |
| 0,54985 | 4,39187 | -2,61414 | 0,54985 | 3,508469 | 0,738579 | O94887;H7 | FARP2 |
| 0,113186 | -1,25823 | 2,35474 | 0,113186 | 1,8238649 | 0,738639 | A0A0C4DGI | UHRF1BP1L |
| -0,372706 | 0,277067 | 0,362267 | 0,277067 | 0,4020052 | 0,738642 | P28331;B4I | NDUFS1 |
| 0,719176 | 0,940621 | -3,20325 | 0,719176 | 2,3311703 | 0,739066 | O60518;A0. | RANBP6 |
| -0,252257 | -0,109903 | 0,703604 | -0,109903 | 0,515708 | 0,739067 | A0A0A0MR | GPSM1 |
| 0,0734324 | 0,626688 | -0,37043 | 0,0734324 | 0,4995581 | 0,739848 | Q8IWI9 | MGA |
| 0,220752 | -0,137406 | -0,244001 | -0,137406 | 0,2434594 | 0,739877 | P13489;H0Y | RNH1 |
| -2,5013 | 1,93782 | 2,33126 | 1,93782 | 2,6837231 | 0,740311 | Q5THN1;Q9 | HEBP2 |

| | | | | | | | |
|---|---|---|---|---|---|---|---|
| -1,21838 | -0,960016 | 1,2758 | -0,960016 | 1,3715295 | 0,740536 | Q9NP79;A0 | VTA1 |
| 0,699848 | -0,739114 | 0,561506 | 0,561506 | 0,7938684 | 0,740628 | Q8NFW8;F5 | CMAS |
| -1,51 | 0,436503 | 0,351336 | 0,351336 | 1,100053 | 0,741129 | A0A0C4DH4 | RPL30 |
| -2,17539 | 0,335889 | 0,790806 | 0,335889 | 1,5974871 | 0,741135 | P06730;D6I | EIF4E |
| -1,7489 | -3,02399 | 2,77838 | -1,7489 | 3,0493064 | 0,74201 | Q9NPH2;J3 | ISYNA1 |
| -0,414047 | 1,573 | -0,410978 | -0,410978 | 1,1463372 | 0,742597 | A8MU46;E9 | SMTNL1 |
| 0,339337 | 0,526396 | -1,65325 | 0,339337 | 1,208046 | 0,742817 | Q8IV33 | KIAA0825 |
| -2,95741 | 1,67339 | 3,43585 | 1,67339 | 3,3021293 | 0,742908 | O95372;Q5 | LYPLA2 |
| 0,348411 | -1,24371 | 1,92608 | 0,348411 | 1,5849005 | 0,743376 | Q8NEZ3;D6 | WDR19 |
| 0,739009 | -0,936429 | 0,846767 | 0,739009 | 0,9998743 | 0,743726 | P04350;M0 | TUBB4A |
| 1,45786 | 0,808106 | -1,32361 | 0,808106 | 1,4550469 | 0,744385 | Q7Z4S6;A0 | KIF21A |
| -0,650836 | 0,0193499 | 1,25704 | 0,0193499 | 0,967903 | 0,744882 | L8E898 | MUC12 |
| -0,107685 | 0,00288151 | 0,208455 | 0,0028815 | 0,1604317 | 0,744963 | Q5THK1;C9 | PRR14L |
| -3,29985 | 2,43833 | 3,14519 | 2,43833 | 3,5347065 | 0,744965 | A0A2R8Y8E | BRAF |
| 0,414891 | -1,86021 | 3,02326 | 0,414891 | 2,4436296 | 0,745088 | Q86UW6;H | N4BP2 |
| 0,501925 | -0,617624 | 0,540397 | 0,501925 | 0,6577592 | 0,745111 | Q07157;G3 | TJP1 |
| 3,19984 | -2,32739 | -3,08323 | -2,32739 | 3,4302219 | 0,745545 | Q8TAP6 | CEP76 |
| 2,92432 | -3,42716 | 2,84935 | 2,84935 | 3,6455794 | 0,745854 | Q02413 | DSG1 |
| 2,54618 | -1,05476 | -0,275406 | -0,275406 | 1,894532 | 0,746523 | P49589 | CARS |
| 2,54617 | -1,05477 | -0,275411 | -0,275411 | 1,8945311 | 0,746526 | B4DKY1 | CARS |
| 0,597055 | -0,144396 | -0,94864 | -0,144396 | 0,77306 | 0,746623 | Q6T4P5 | PLPPR3 |
| -0,32849 | 1,16797 | -0,292791 | -0,292791 | 0,8538628 | 0,747114 | O95140;Q5 | MFN2 |
| -0,84805 | 0,663084 | 0,760433 | 0,663084 | 0,9018703 | 0,747917 | H0Y684 | OBSL1 |
| -2,76505 | 3,87854 | -3,7611 | -2,76505 | 4,1531816 | 0,748135 | Q5TDH0 | DDI2 |
| -3,18832 | 4,27077 | -3,97766 | -3,18832 | 4,551514 | 0,74865 | E7ERS3;Q8 | ZC3H18 |
| -1,49542 | 1,41528 | 1,09398 | 1,09398 | 1,5958488 | 0,748948 | P08758;D6I | ANXA5 |
| 2,68656 | -2,46612 | 1,48804 | 1,48804 | 2,6963538 | 0,749567 | O00241;H3 | SIRPB1 |
| 5,89585 | -3,51292 | 0,605181 | 0,605181 | 4,7165469 | 0,749599 | Q8N6Q8 | METTL25 |
| -2,26456 | 2,86916 | -2,53107 | -2,26456 | 3,0438077 | 0,74983 | A0A0D9SGJ | SYNJ1 |
| -1,37779 | 0,906178 | -0,250847 | -0,250847 | 1,142017 | 0,74994 | P61586;C9J | RHOA |
| 0,65071 | 0,372698 | -1,91067 | 0,372698 | 1,4054494 | 0,750429 | Q9Y4L1;A0 | HYOU1 |
| 4,36429 | 1,38137 | -3,31338 | 1,38137 | 3,8705104 | 0,751499 | Q9UKI9 | POU2F3 |
| 0,659494 | -0,498668 | -0,600316 | -0,498668 | 0,6998563 | 0,751662 | Q96E17 | RAB3C |
| -3,08014 | -1,12241 | 2,44461 | -1,12241 | 2,8011665 | 0,75181 | Q6UVK1 | CSPG4 |
| 3,36143 | -2,56744 | -3,03001 | -2,56744 | 3,5640795 | 0,751883 | P08621;M0 | SNRNP70 |
| -3,22646 | 0,540557 | 1,19014 | 0,540557 | 2,3846289 | 0,751929 | Q96DN6;F8 | MBD6 |
| 2,15072 | -0,880853 | -0,266172 | -0,266172 | 1,602583 | 0,752284 | A0A0A0MR | CDC42BPA |
| -0,313728 | 1,24563 | -0,361159 | -0,313728 | 0,9142955 | 0,753047 | Q9NQ35;E9 | NRIP3 |
| -2,5774 | 3,7424 | 0,808405 | 0,808405 | 3,1625906 | 0,753144 | P63096;C9J | GNAI1 |
| 1,90753 | -0,69462 | -2,63191 | -0,69462 | 2,2778203 | 0,753524 | O94913;E9I | PCF11 |
| 0,481369 | -3,25494 | 1,26888 | 0,481369 | 2,4167859 | 0,753658 | Q05086;A0 | UBE3A |
| -0,368538 | -0,590145 | 1,76778 | -0,368538 | 1,3020992 | 0,754111 | K7ES47 | ZNF532 |
| 0,904808 | 0,391603 | -2,40125 | 0,391603 | 1,7792052 | 0,754262 | Q9HBL0;E9 | TNS1 |
| 0,727316 | 1,46732 | -1,30405 | 0,727316 | 1,4349533 | 0,754389 | Q92834 | RPGR |
| -0,422852 | 1,03729 | -1,36421 | -0,422852 | 1,2100532 | 0,754768 | Q96GM5;F8 | SMARCD1 |
| 0,780827 | 0,110768 | -0,496059 | 0,110768 | 0,6387039 | 0,754892 | E5RHG8;Q1 | ELOC |
| -1,77586 | 0,140312 | 3,18091 | 0,140312 | 2,4995506 | 0,755273 | A0A0B4J1W | MED24 |
| 1,65969 | 1,89439 | -2,15072 | 1,65969 | 2,2707276 | 0,755361 | Q9NQA5;A0 | TRPV5 |
| -0,174067 | 0,460218 | -0,620695 | -0,174067 | 0,5431646 | 0,756144 | P55287 | CDH11 |
| 3,88716 | -2,88792 | 1,09047 | 1,09047 | 3,4046725 | 0,75694 | O95391 | SLU7 |
| 1,21662 | -0,577506 | -1,48069 | -0,577506 | 1,3729597 | 0,757244 | Q96M83;A0 | CCDC7 |
| 1,45058 | 1,69831 | -1,91521 | 1,45058 | 2,0185572 | 0,757913 | Q8NB16;I3I | MLKL |

| | | | | | | | |
|---|---|---|---|---|---|---|---|
| 3,25754 | -0,380336 | -1,38539 | -0,380336 | 2,4427128 | 0,758081 | Q9P0K7 | RAI14 |
| 2,51107 | -2,8243 | 2,12761 | 2,12761 | 2,9758648 | 0,758462 | Q2WGJ9 | FER1L6 |
| 1,28966 | -1,41001 | 1,02751 | 1,02751 | 1,4887604 | 0,758595 | O60733 | PLA2G6 |
| 1,36692 | -3,24022 | 0,395876 | 0,395876 | 2,4286436 | 0,75897 | Q9BQI6 | SLF1 |
| 1,58437 | -0,30687 | -2,52848 | -0,30687 | 2,0586353 | 0,759216 | P53621 | COPA |
| 1,68108 | -1,32484 | 0,566283 | 0,566283 | 1,5195763 | 0,759433 | B1AJZ9;H0Y | FHAD1 |
| -0,0799999 | 0,876662 | -1,533 | -0,08 | 1,2133207 | 0,759514 | Q9UER7 | DAXX |
| -1,77544 | -2,60532 | 2,6658 | -1,77544 | 2,8342553 | 0,760184 | Q5HYB6;Q5 | TPM3 |
| 2,66219 | 0,382868 | -1,71964 | 0,382868 | 2,1915095 | 0,760292 | P30084 | ECHS1 |
| -0,620128 | 0,0679809 | 1,06402 | 0,0679809 | 0,8467528 | 0,760399 | O00186 | STXBP3 |
| 0,149717 | -0,63106 | 0,962513 | 0,149717 | 0,7968401 | 0,760646 | P11498;E9F | PC |
| -0,412796 | 1,74526 | -0,554587 | -0,412796 | 1,2888371 | 0,760758 | O60309 | LRRC37A3 |
| -3,07482 | 4,14269 | -3,69767 | -3,07482 | 4,3579743 | 0,760795 | P13569;E7E | CFTR |
| 2,93556 | -0,241938 | -1,35186 | -0,241938 | 2,2252487 | 0,760974 | A0A087WZ | LIPI |
| -0,662176 | -0,812881 | 2,65503 | -0,662176 | 1,9601435 | 0,761343 | Q8TER5;G3 | ARHGEF40 |
| -1,04599 | 0,129035 | 0,444303 | 0,129035 | 0,7853928 | 0,761412 | Q9UN37;I3 | VPS4A |
| 0,40239 | -0,704005 | 0,760797 | 0,40239 | 0,7635673 | 0,761574 | Q9H040;L8 | SPRTN |
| 0,871656 | 0,321489 | -2,1664 | 0,321489 | 1,6187474 | 0,761621 | Q9NV72;M | ZNF701 |
| 0,72574 | 0,133049 | -1,5749 | 0,133049 | 1,1945234 | 0,762273 | E7ET52;Q8 | ZFPM2 |
| 2,78658 | -2,26255 | -2,27219 | -2,26255 | 2,9179034 | 0,762415 | A5YKK6;H3 | CNOT1 |
| -2,96991 | 1,60155 | -0,0200794 | -0,020079 | 2,3176651 | 0,762434 | Q58FG0 | HSP90AA5P |
| -1,29466 | 0,1877 | 0,528334 | 0,1877 | 0,9692549 | 0,763215 | P63167;F8\ | DYNLL1 |
| -1,84375 | 1,73408 | -1,0072 | -1,0072 | 1,8715101 | 0,76329 | A0A2R8YFA | MICAL2 |
| 1,09473 | 2,18622 | -1,98885 | 1,09473 | 2,1652954 | 0,763307 | Q8IYU2;H0 | HACE1 |
| 1,4993 | -2,42509 | -0,246891 | -0,246891 | 1,9661541 | 0,763419 | R4GMW8 | BIVM-ERCC5 |
| 3,2547 | -3,26064 | 2,07137 | 2,07137 | 3,470837 | 0,763926 | P51571;A6I | SSR4 |
| 0,632021 | -1,63375 | 0,278095 | 0,278095 | 1,2188884 | 0,764449 | Q92736;H7 | RYR2 |
| -0,955363 | 0,233574 | 0,302253 | 0,233574 | 0,7070933 | 0,764584 | O75150;H3 | RNF40 |
| 2,76229 | -0,169479 | -1,34353 | -0,169479 | 2,1146876 | 0,765544 | Q5T282;Q9 | SNAPC3 |
| -2,92983 | 3,75273 | 1,1676 | 1,1676 | 3,3696795 | 0,765564 | M0R063;M | SLC1A6 |
| 1,34863 | -2,49307 | -0,00354467 | -0,003545 | 1,9487078 | 0,766168 | Q92545;F6 | TMEM131 |
| 2,22646 | 0,731153 | -1,76983 | 0,731153 | 2,0191249 | 0,766482 | Q9UBB9;F6 | TFIP11 |
| -1,01388 | 0,659146 | -0,137107 | -0,137107 | 0,8368359 | 0,76668 | P16298;Q5 | PPP3CB |
| 3,61345 | -2,81931 | -3,0097 | -2,81931 | 3,7701186 | 0,766707 | H3BPZ4 | SH2B1 |
| -1,27237 | 1,254 | 0,811023 | 0,811023 | 1,3490305 | 0,766741 | Q02978;I3L | SLC25A11 |
| -2,42929 | -1,50661 | 2,42437 | -1,50661 | 2,5775297 | 0,767172 | Q6UN15;H( | FIP1L1 |
| -1,15642 | -0,102294 | 2,2888 | -0,102294 | 1,7653164 | 0,768266 | A0A2R8YEI | CHN2 |
| 2,23452 | -0,64706 | -0,628669 | -0,628669 | 1,6583975 | 0,770286 | E7EPT4;P19 | NDUFV2 |
| 0,76025 | -0,238042 | -0,195978 | -0,195978 | 0,5646132 | 0,770418 | Q9HCG1;M | ZNF160 |
| 1,00224 | -0,610492 | -1,00621 | -0,610492 | 1,0639054 | 0,770507 | A8TX70;E9F | COL6A5 |
| -1,25284 | 3,27038 | -0,605811 | -0,605811 | 2,4461883 | 0,770673 | C9IZY1;C9J | OARD1 |
| 1,05994 | -1,49803 | 1,3406 | 1,05994 | 1,5641718 | 0,770719 | P16403;Q0 | HIST1H1C |
| -0,235445 | -0,6339 | 0,529403 | -0,235445 | 0,5911899 | 0,771465 | A0A0U1RR | DENND4A |
| -2,47358 | 0,987052 | 3,10482 | 0,987052 | 2,8160096 | 0,771591 | Q02779;E9 | MAP3K10 |
| 1,13731 | 1,45079 | -1,618 | 1,13731 | 1,6885632 | 0,771653 | Q08722;A0 | CD47 |
| 1,03457 | -1,40594 | 1,21072 | 1,03457 | 1,4625336 | 0,771884 | Q92830 | KAT2A |
| -0,518165 | 0,500342 | 0,330831 | 0,330831 | 0,5457235 | 0,77201 | Q13907 | IDI1 |
| -0,947428 | 4,85428 | -1,8265 | -0,947428 | 3,6300919 | 0,772191 | Q05DH4 | FAM160A1 |
| 1,80852 | 2,97181 | -2,9758 | 1,80852 | 3,1521693 | 0,772422 | Q6PDA5 | WDR3 |
| -2,65009 | -0,229018 | 1,64857 | -0,229018 | 2,1550485 | 0,772975 | A0A087WY | AP2M1 |
| -3,1312 | 0,84791 | 0,953724 | 0,84791 | 2,3284872 | 0,772976 | Q58EX2;H7 | SDK2 |
| 1,45914 | 0,833499 | -1,42684 | 0,833499 | 1,5181919 | 0,773246 | Q96K21;H3 | ZFYVE19 |

| | | | | | | | |
|---|---|---|---|---|---|---|---|
| 2,27945 | 0,323919 | -1,52044 | 0,323919 | 1,900216 | 0,773392 | A4FU69;H0 | EFCAB5 |
| -1,68945 | 1,26321 | 1,42369 | 1,26321 | 1,7528831 | 0,773719 | A0A1X7SC7 | FAM92B |
| -0,313359 | 1,55119 | -0,577954 | -0,313359 | 1,1604459 | 0,773868 | J3KNC0;P52 | GTF2A1 |
| 0,984423 | 0,441744 | -2,48346 | 0,441744 | 1,8653655 | 0,774561 | Q99729;D6 | HNRNPAB |
| 1,05766 | -0,3754 | -1,37397 | -0,3754 | 1,2222675 | 0,774891 | Q8N9W4;F8 | GOLGA6L2 |
| -3,30406 | 4,49488 | 1,01923 | 1,01923 | 3,9071397 | 0,774999 | Q92503;K7I | SEC14L1 |
| -0,631936 | 1,08648 | -1,10573 | -0,631936 | 1,1534888 | 0,775418 | Q8N3C0;E5 | ASCC3 |
| 0,0226534 | 0,87931 | -1,61777 | 0,0226534 | 1,2688747 | 0,775571 | Q9Y6F6;H0 | MRVI1 |
| -0,438998 | 0,128623 | 0,605148 | 0,128623 | 0,5227349 | 0,775654 | H0Y2S9;H0 | MPRIP |
| 4,24417 | -3,30405 | -3,40917 | -3,30405 | 4,3886271 | 0,776147 | E9PNC0 | POLD3 |
| -2,24465 | 0,874345 | 0,422701 | 0,422701 | 1,6855698 | 0,776304 | A0A0U1RQ | NRXN3 |
| -2,41202 | 1,02283 | 0,365125 | 0,365125 | 1,8231522 | 0,776489 | Q96LX7;F2 | CCDC17 |
| -2,29332 | 5,62993 | -0,970953 | -0,970953 | 4,2445691 | 0,778139 | Q9BY89 | KIAA1671 |
| 0,566492 | -0,332749 | 0,0185395 | 0,0185395 | 0,4531905 | 0,778387 | P02533 | KRT14 |
| -2,57713 | 1,14918 | 0,338374 | 0,338374 | 1,9597175 | 0,77865 | A0A1B0GVI | ARID1B |
| 0,490843 | -1,14234 | 0,171106 | 0,171106 | 0,8655113 | 0,779009 | Q96JM4;H0 | LRRIQ1 |
| -1,0214 | -2,84503 | 2,39125 | -1,0214 | 2,6580204 | 0,779026 | Q15436;F5I | SEC23A |
| 1,35806 | -1,92959 | 1,6773 | 1,35806 | 1,9966727 | 0,779477 | A0A087WT | PCDH15 |
| -0,192086 | -0,250016 | 0,754217 | -0,192086 | 0,5638157 | 0,779563 | Q13510;A0 | ASAH1 |
| -1,99526 | 2,40799 | -1,78654 | -1,78654 | 2,4841584 | 0,77977 | A0A2R8Y3T | COCH |
| -1,00067 | -0,437406 | 0,898628 | -0,437406 | 0,9754987 | 0,779783 | A0A0D9SFE | DNM1 |
| -1,85961 | -0,302177 | 1,29532 | -0,302177 | 1,5775074 | 0,781197 | A0A2R8Y5Z | EPB41 |
| 0,741751 | 0,173707 | -0,55801 | 0,173707 | 0,6515958 | 0,78146 | O43761;Q9 | SYNGR3 |
| -0,861428 | -3,52884 | 2,68285 | -0,861428 | 3,1161431 | 0,781705 | A0A075B74 | PDE4DIP |
| -1,96225 | 1,52369 | -0,519959 | -0,519959 | 1,7515937 | 0,781971 | A0A0G2JNE | KANSL1 |
| -2,05286 | 2,21207 | 1,04693 | 1,04693 | 2,2043849 | 0,781996 | Q04760 | GLO1 |
| 0,692109 | -0,508092 | -0,573628 | -0,508092 | 0,7126088 | 0,782155 | P37837 | TALDO1 |
| -1,32024 | 0,567557 | 1,54644 | 0,567557 | 1,4571573 | 0,782918 | P0C0S5;Q7 | H2AFZ |
| 1,48923 | -0,717133 | -1,64999 | -0,717133 | 1,6120878 | 0,782979 | Q9H2J7;F8 | SLC6A15 |
| 4,52853 | -3,38512 | -3,67721 | -3,38512 | 4,6555584 | 0,783099 | Q9NVR2;E5 | INTS10 |
| 0,462603 | -1,71582 | 2,35939 | 0,462603 | 2,0392263 | 0,783784 | O14646 | CHD1 |
| -1,45209 | -0,483008 | 1,20575 | -0,483008 | 1,3450611 | 0,783864 | Q8NCU4;C9 | CCDC191 |
| 1,58466 | 0,107074 | -2,94006 | 0,107074 | 2,3072847 | 0,78432 | P26232;A0 | CTNNA2 |
| -0,863927 | 1,00005 | -0,691797 | -0,691797 | 1,0300797 | 0,784924 | Q9H479 | FN3K |
| 1,25652 | -0,790522 | 0,0855995 | 0,0855995 | 1,0270528 | 0,78583 | P30050 | RPL12 |
| -1,86462 | -1,95458 | 2,46399 | -1,86462 | 2,525494 | 0,786005 | P52907 | CAPZA1 |
| -2,05054 | 7,15461 | -2,22588 | -2,05054 | 5,3659284 | 0,786092 | A0A1B0GW | MBD5 |
| -2,9385 | 2,89138 | 1,69731 | 1,69731 | 3,0796114 | 0,786296 | O14980;C9 | XPO1 |
| -2,16602 | 1,0034 | 0,275146 | 0,275146 | 1,6600637 | 0,78677 | Q16775;H3 | HAGH |
| 2,03106 | -0,433906 | -2,91985 | -0,433906 | 2,4754624 | 0,786876 | Q9NZR2 | LRP1B |
| -0,076449 | 3,79918 | -2,11744 | -0,076449 | 3,0053435 | 0,786943 | O75164 | KDM4A |
| -0,182267 | -0,0678893 | 0,157854 | -0,067889 | 0,1730725 | 0,787259 | O60934;A0 | NBN |
| 0,764237 | -2,26196 | 0,594762 | 0,594762 | 1,7003652 | 0,788126 | Q00839;A0 | HNRNPU |
| -3,05092 | 5,50126 | -0,142093 | -0,142093 | 4,3483424 | 0,788205 | P41440 | SLC19A1 |
| -0,824047 | 1,4893 | -1,49625 | -0,824047 | 1,5661492 | 0,788293 | Q9UKV8 | Aug-02 |
| -1,25904 | -0,12984 | 0,833984 | -0,12984 | 1,0476003 | 0,788644 | P31943;G8 | HNRNPH1 |
| -1,06106 | 0,807945 | 0,826107 | 0,807945 | 1,0843515 | 0,789124 | Q15814 | TBCC |
| -0,942553 | 0,996214 | 0,475431 | 0,475431 | 1,0033869 | 0,789549 | P40123;E9F | CAP2 |
| -0,496418 | -2,979 | 2,12997 | -0,496418 | 2,5548223 | 0,789808 | P0C7V7 | SEC11B |
| 2,72455 | 0,609126 | -2,0693 | 0,609126 | 2,4024287 | 0,789936 | D6R9D1;D6 | PACRGL |
| 0,677375 | -0,879272 | 0,67413 | 0,67413 | 0,8977953 | 0,79005 | O15254 | ACOX3 |
| -2,98667 | 2,32981 | 2,2605 | 2,2605 | 3,04966 | 0,790108 | Q9BXL6;I3L | CARD14 |

| | | | | | | | |
|---|---|---|---|---|---|---|---|
| -2,51008 | -2,37986 | 3,18153 | -2,37986 | 3,2491137 | 0,790121 | Q8IZJ4;B5M | RGL4 |
| 2,21778 | 1,58839 | -2,46989 | 1,58839 | 2,5442752 | 0,79035 | Q9Y6R4;F5I | MAP3K4 |
| 0,342348 | 0,304294 | -0,42098 | 0,304294 | 0,4301434 | 0,790574 | P29966 | MARCKS |
| 1,6376 | -1,34943 | 0,500743 | 0,500743 | 1,5076434 | 0,791087 | Q9UIA9;E7I | XPO7 |
| -0,898626 | 0,976432 | 0,425915 | 0,425915 | 0,9637877 | 0,791327 | Q92932;A0 | PTPRN2 |
| 0,59612 | -0,321272 | -0,602245 | -0,321272 | 0,6267153 | 0,791422 | Q9Y6D9;C9 | MAD1L1 |
| 1,11251 | 1,36818 | -1,61869 | 1,11251 | 1,6556073 | 0,792089 | Q3B7T1 | EDRF1 |
| 1,65446 | 2,00811 | -2,39027 | 1,65446 | 2,4437217 | 0,792095 | Q68DE3;C9 | USF3 |
| -0,363637 | 1,20633 | -1,56633 | -0,363637 | 1,3903783 | 0,792163 | P23396;H0\ | RPS3 |
| -0,402305 | 1,46661 | -1,95542 | -0,402305 | 1,7134419 | 0,79231 | P38398;Q5\ | BRCA1 |
| -3,03729 | -0,941534 | 2,52103 | -0,941534 | 2,8070288 | 0,792592 | E7EQA9;Q9 | TANK |
| -0,486208 | -1,18803 | 1,07472 | -0,486208 | 1,1582377 | 0,793253 | P50454;E9F | SERPINH1 |
| 3,20446 | -3,44315 | 2,0772 | 2,0772 | 3,5575202 | 0,793564 | C9J3G2;F2Z | CBWD2 |
| 0,842086 | -1,73259 | 1,84319 | 0,842086 | 1,8446937 | 0,793697 | P41229;F8\ | KDM5C |
| 2,12852 | 2,48224 | -3,0209 | 2,12852 | 3,0802109 | 0,793809 | E9PJ95;Q9F | COMMD9 |
| -1,134 | 1,03036 | -0,468404 | -0,468404 | 1,1085851 | 0,79386 | Q9H269 | VPS16 |
| 3,38713 | 1,4036 | -3,08395 | 1,4036 | 3,3153022 | 0,794321 | P05556 | ITGB1 |
| -2,0377 | -1,8579 | 2,55801 | -1,8579 | 2,6029835 | 0,794685 | Q70J99;K7E | UNC13D |
| -1,70128 | 0,836317 | 1,79227 | 0,836317 | 1,805461 | 0,794781 | P24752;H0\ | ACAT1 |
| -0,712973 | 0,323611 | 0,782885 | 0,323611 | 0,7662713 | 0,794803 | O94776 | MTA2 |
| 0,261677 | -0,439752 | 0,411443 | 0,261677 | 0,4544164 | 0,794804 | Q5VU43;A0 | PDE4DIP |
| -0,784239 | -2,5471 | 2,12185 | -0,784239 | 2,3576869 | 0,795017 | A0A1B0GVI | LCORL |
| 0,0374406 | 0,358678 | -0,663832 | 0,0374406 | 0,5228931 | 0,795404 | K7ERF1;Q9I | EIF3K |
| 1,41325 | 2,52453 | -2,56757 | 1,41325 | 2,6774165 | 0,795489 | H3BV82 | KLHDC4 |
| -1,92688 | 2,48797 | 0,570552 | 0,570552 | 2,213766 | 0,795711 | P31751;M0 | AKT2 |
| 2,06649 | -2,05487 | -1,11573 | -1,11573 | 2,1600199 | 0,795721 | P43003;A0/ | SLC1A3 |
| 0,36066 | -0,875189 | 1,00142 | 0,36066 | 0,9539005 | 0,796003 | Q7LBC6 | KDM3B |
| -0,166458 | -0,434729 | 0,986639 | -0,166458 | 0,7551919 | 0,796011 | P07814;V9( | EPRS |
| -0,904661 | 0,695555 | 0,675725 | 0,675725 | 0,9182142 | 0,796861 | O00159 | MYO1C |
| -0,812844 | 1,4257 | -0,0391959 | -0,039196 | 1,1369206 | 0,798244 | O95861;A6 | BPNT1 |
| 1,41665 | -2,35944 | 2,16444 | 1,41665 | 2,4249926 | 0,798552 | Q9UKN8 | GTF3C4 |
| 3,62919 | -3,52907 | -1,99343 | -1,99343 | 3,7685703 | 0,799081 | Q96F86;H3 | EDC3 |
| 0,462113 | -0,957243 | 1,00251 | 0,462113 | 1,012195 | 0,799513 | Q70CQ2;H7 | USP34 |
| -0,736805 | -2,29496 | 1,95442 | -0,736805 | 2,1497197 | 0,799555 | Q01813;Q5 | PFKP |
| -0,429934 | -0,909706 | 2,16967 | -0,429934 | 1,6568383 | 0,799626 | Q8WYP5;H | AHCTF1 |
| -1,89643 | -0,662962 | 1,65588 | -0,662962 | 1,8035788 | 0,799634 | P62805;B2F | HIST1H4A |
| -2,43415 | 1,9172 | 1,74583 | 1,74583 | 2,464273 | 0,800507 | Q9HBD1 | RC3H2 |
| -0,209063 | -0,876689 | 1,76516 | -0,209063 | 1,3737204 | 0,802085 | G3V559 | ATP6V1D |
| 0,170864 | -2,65834 | 1,44935 | 0,170864 | 2,102064 | 0,802359 | P16870;C9J | CPE |
| -2,97338 | 0,993621 | 0,866945 | 0,866945 | 2,2546707 | 0,802475 | O15015;C9. | ZNF646 |
| -1,05543 | 1,12992 | 0,478882 | 0,478882 | 1,1220308 | 0,802619 | P51513;F8\ | NOVA1 |
| 1,21106 | 0,0711894 | -2,11406 | 0,0711894 | 1,689726 | 0,80297 | Q8IZ02;G3\ | LRRC34 |
| -0,718753 | 0,497106 | 0,579175 | 0,497106 | 0,7268271 | 0,803112 | Q9H8V3;C9 | ECT2 |
| -1,10961 | 2,03751 | -0,136542 | -0,136542 | 1,6113001 | 0,803409 | P35789 | ZNF93 |
| 0,68117 | -0,529686 | -0,48864 | -0,48864 | 0,6875454 | 0,8037 | Q8IVL1;A0/ | NAV2 |
| -0,546921 | 1,16084 | -0,17417 | -0,17417 | 0,8979275 | 0,803944 | P50416 | CPT1A |
| 1,76524 | -1,06413 | -1,58263 | -1,06413 | 1,8019624 | 0,804152 | Q9Y2J2;A0/ | EPB41L3 |
| -1,09675 | 0,775789 | -0,13694 | -0,13694 | 0,9363681 | 0,804223 | B8ZZM6;Q9 | ITM2C |
| 0,0667259 | -1,49546 | 0,846317 | 0,0667259 | 1,1924839 | 0,804458 | Q96GE4 | CEP95 |
| -0,311553 | -0,244519 | 0,883683 | -0,244519 | 0,6715557 | 0,804677 | Q9UMY4;A( | SNX12 |
| 1,14774 | -1,99834 | 0,0712413 | 0,0712413 | 1,5989494 | 0,804839 | Q8IYY4;C9J | DZIP1L |
| 0,872208 | -1,51109 | 0,0489409 | 0,0489409 | 1,2104802 | 0,804862 | B5BUI8;P51 | DUSP3 |

| | | | | | | | |
|---|---|---|---|---|---|---|---|
| -2,86494 | 0,137826 | 1,61493 | 0,137826 | 2,2828226 | 0,804925 | Q5SNV9;H7 | C1orf167 |
| -2,65208 | 1,63676 | 2,32979 | 1,63676 | 2,6985634 | 0,804961 | P30049 | ATP5F1D |
| 2,02304 | 0,0956622 | -3,47431 | 0,0956622 | 2,7892754 | 0,805382 | P04626;J3C | ERBB2 |
| -0,472649 | -0,500106 | 1,54057 | -0,472649 | 1,1703392 | 0,805704 | P51784;G5I | USP11 |
| 1,21235 | 0,610825 | -2,89853 | 0,610825 | 2,2202379 | 0,806024 | Q6ZMV8 | ZNF730 |
| -0,653891 | 0,6522 | -0,327531 | -0,327531 | 0,6797374 | 0,806026 | Q9H4A3;F5 | WNK1 |
| 1,30195 | 0,468095 | -2,82616 | 0,468095 | 2,1828392 | 0,806223 | P0C671 | C6orf222 |
| -0,170013 | 1,07599 | -1,53639 | -0,170013 | 1,3066521 | 0,806746 | Q08945;E9I | SSRP1 |
| -0,892126 | 1,09725 | -0,73888 | -0,73888 | 1,1069835 | 0,806862 | Q9UL25 | RAB21 |
| -0,23289 | 0,0858743 | 0,0614727 | 0,0614727 | 0,1774145 | 0,806864 | Q96QK1;A0 | VPS35 |
| -2,43426 | 2,87593 | 0,850411 | 0,850411 | 2,6798603 | 0,806871 | Q96RU2;F5 | USP28 |
| 0,387327 | -0,501128 | -0,100189 | -0,100189 | 0,44493 | 0,807331 | P25788;G3Y | PSMA3 |
| 0,40025 | 0,518288 | -1,44857 | 0,40025 | 1,1030714 | 0,807504 | Q9NVU0;A0 | POLR3E |
| -1,22969 | 1,36857 | 0,495804 | 0,495804 | 1,3222459 | 0,807696 | P49841;B5E | GSK3B |
| 0,159527 | 1,01845 | -0,753155 | 0,159527 | 0,8859384 | 0,807885 | O60216 | RAD21 |
| -1,37209 | -0,697721 | 1,38163 | -0,697721 | 1,4353541 | 0,807912 | P37840;E7E | SNCA |
| 1,94255 | -1,97316 | 1,00954 | 1,00954 | 2,0453114 | 0,808229 | Q96FA7 | ZBED6CL |
| 3,14873 | -2,39729 | -2,2604 | -2,2604 | 3,16322 | 0,808843 | B7WPN9;Q | ATP13A4 |
| -0,405568 | 0,285494 | -0,0445888 | -0,044589 | 0,3456461 | 0,809091 | P26641 | EEF1G |
| 0,946906 | -1,14157 | -0,304808 | -0,304808 | 1,051086 | 0,809552 | P13797;A0 | PLS3 |
| 0,437158 | -1,71696 | 0,65654 | 0,437158 | 1,3116056 | 0,809555 | Q8IUD2;X6 | ERC1 |
| 0,859838 | 0,0854008 | -1,51875 | 0,0854008 | 1,2131731 | 0,810502 | P15880;E9F | RPS2 |
| -0,875588 | 0,928756 | 0,383671 | 0,383671 | 0,9254286 | 0,810772 | H0Y6H0;Q8 | KDM1B |
| -0,661043 | -0,328097 | 1,55179 | -0,328097 | 1,1931372 | 0,810952 | Q9P2D7 | DNAH1 |
| 1,79074 | -2,11731 | -0,602106 | -0,602106 | 1,9703811 | 0,811051 | A0A0A0MS | FSCN1 |
| -0,104704 | -1,53083 | 1,03165 | -0,104704 | 1,2839678 | 0,811436 | O00764;F2 | PDXK |
| 1,07007 | -0,106929 | -0,566279 | -0,106929 | 0,8439952 | 0,811476 | P13611;E9F | VCAN |
| 3,18653 | -1,56588 | -0,455133 | -0,455133 | 2,4859897 | 0,812011 | O60763 | USO1 |
| -0,549984 | 1,82311 | -0,622376 | -0,549984 | 1,3914751 | 0,812462 | Q9H2U1;E7 | DHX36 |
| -3,0912 | 3,03853 | 1,54709 | 1,54709 | 3,1966576 | 0,812529 | Q9H0Q3 | FXYD6 |
| 0,335969 | 0,667397 | -0,676623 | 0,335969 | 0,7001877 | 0,812856 | Q9C0D2 | CEP295 |
| -1,18827 | 3,31992 | -0,950898 | -0,950898 | 2,537059 | 0,813339 | P28072;A0 | PSMB6 |
| 0,541642 | -0,32334 | -0,47302 | -0,32334 | 0,5477434 | 0,813484 | P55884;C9J | EIF3B |
| -2,40617 | -0,458655 | 1,87071 | -0,458655 | 2,1412792 | 0,813781 | P14625;A0 | HSP90B1 |
| 0,304375 | 1,33916 | -2,58829 | 0,304375 | 2,0356471 | 0,813842 | Q6ZNF0 | ACP7 |
| -1,8968 | 3,14007 | -2,69825 | -1,8968 | 3,1648687 | 0,815537 | O60239 | SH3BP5 |
| -2,28943 | -3,28422 | 3,80868 | -2,28943 | 3,8402644 | 0,815588 | E9PGC0;P2 | RASA1 |
| -3,06144 | 2,92522 | 1,57843 | 1,57843 | 3,1406564 | 0,815739 | Q14774 | HLX |
| 2,55221 | 2,17307 | -3,23886 | 2,17307 | 3,2395788 | 0,815886 | Q8NFP7;Q9 | NUDT10 |
| 0,550421 | 0,299177 | -0,577831 | 0,299177 | 0,5923427 | 0,815897 | Q6ZTR5;A0 | CFAP47 |
| -3,01349 | 2,92047 | 1,51539 | 1,51539 | 3,1009914 | 0,815943 | Q8WVM8;C | SCFD1 |
| 1,26031 | -2,24801 | 2,03274 | 1,26031 | 2,2814388 | 0,816182 | Q92820 | GGH |
| -0,209795 | 0,989026 | -0,429813 | -0,209795 | 0,763619 | 0,81637 | Q16720 | ATP2B3 |
| 1,07479 | -1,47296 | 1,07003 | 1,07003 | 1,469572 | 0,816525 | Q3ZCQ8;M | TIMM50 |
| -1,29599 | -3,10632 | 2,98186 | -1,29599 | 3,1263192 | 0,817622 | A6NKZ9;E7 | TIAL1 |
| 0,905882 | -2,44733 | 2,73444 | 0,905882 | 2,6280029 | 0,817776 | Q96I24 | FUBP3 |
| 1,38547 | 1,1261 | -1,72775 | 1,1261 | 1,7274196 | 0,817854 | K7EM11;O1 | GIPC1 |
| -1,84954 | 0,0559713 | 1,11309 | 0,0559713 | 1,5014243 | 0,818062 | C9JFP8 | SHANK2 |
| -3,2321 | 4,07398 | -2,68394 | -2,68394 | 4,069168 | 0,818268 | Q92667;I3L | AKAP1 |
| 1,5139 | -1,30307 | 0,43236 | 0,43236 | 1,4210774 | 0,8183 | Q9BS26 | ERP44 |
| 1,69546 | 1,43312 | -2,15529 | 1,43312 | 2,1515027 | 0,818388 | Q49MG5;A | MAP9 |
| -1,49963 | -0,519119 | 1,36231 | -0,519119 | 1,4544115 | 0,818789 | P25054;E9F | APC |

| | | | | | | | |
|---|---|---|---|---|---|---|---|
| 3,07945 | -3,21218 | -1,32441 | -1,32441 | 3,2285773 | 0,818798 | Q9BVG8;F5 | KIFC3 |
| 0,752538 | 0,945787 | -2,59338 | 0,752538 | 1,9899001 | 0,819389 | Q9UIF8;C9J | BAZ2B |
| 3,07388 | -1,29505 | -3,22998 | -1,29505 | 3,229297 | 0,819557 | P16118 | PFKFB1 |
| 0,230768 | -1,47018 | 0,722589 | 0,230768 | 1,1506048 | 0,819631 | Q13492;B5 | PICALM |
| 0,701582 | 0,998792 | -2,59677 | 0,701582 | 1,9956422 | 0,819632 | J3KN36;P69 | NOMO3 |
| -2,72215 | -2,80637 | 3,82396 | -2,72215 | 3,8039437 | 0,820049 | Q71DI3 | HIST2H3A |
| 0,526695 | -1,79629 | 2,15737 | 0,526695 | 1,9869066 | 0,820551 | P54098;A0/ | POLG |
| 1,26409 | -2,04557 | 1,69519 | 1,26409 | 2,0466632 | 0,820695 | Q99615 | DNAJC7 |
| 2,2103 | 2,16628 | -3,03236 | 2,16628 | 3,014224 | 0,820883 | Q9P2R3 | ANKFY1 |
| -0,911864 | -0,898636 | 2,75126 | -0,898636 | 2,1110974 | 0,821013 | H0Y6T8 | RAB18 |
| 2,48116 | -2,31034 | -1,28888 | -1,28888 | 2,5237226 | 0,822026 | Q9BY76 | ANGPTL4 |
| -0,427639 | 0,410308 | -0,172281 | -0,172281 | 0,4294906 | 0,822624 | O95989 | NUDT3 |
| -3,03236 | 3,20218 | 1,23619 | 1,23619 | 3,187348 | 0,822764 | P62328 | TMSB4X |
| -0,441069 | 0,213866 | 0,0751157 | 0,0751157 | 0,3451179 | 0,822936 | P38919 | EIF4A3 |
| 0,911105 | -2,12097 | 2,18341 | 0,911105 | 2,2113317 | 0,823101 | M0R366;Q9 | FSD1 |
| -2,87645 | 0,406314 | 3,97908 | 0,406314 | 3,4287872 | 0,823169 | A0A075B6T | AMBRA1 |
| -0,847898 | -1,38874 | 1,5491 | -0,847898 | 1,563597 | 0,823313 | P19086 | GNAZ |
| 0,988623 | 1,27468 | -1,57407 | 0,988623 | 1,5686831 | 0,823445 | O15021;E7I | MAST4 |
| -0,23478 | -3,04524 | 2,14016 | -0,23478 | 2,5957465 | 0,823541 | Q52LJ0;H0Y | FAM98B |
| -3,15239 | 4,21243 | -2,89175 | -2,89175 | 4,1788731 | 0,823852 | Q5VZ89 | DENND4C |
| 1,14916 | -0,95481 | -0,697104 | -0,697104 | 1,1475912 | 0,823941 | Q8IWY7 | TTBK2 |
| 0,257101 | -1,44005 | 0,689808 | 0,257101 | 1,125748 | 0,823957 | A0A2R8YFH | SEC23B |
| -2,32306 | 2,5448 | 0,861198 | 0,861198 | 2,4721811 | 0,82396 | Q9P107;K7 | GMIP |
| -1,67376 | 0,113878 | 2,4702 | 0,113878 | 2,0784733 | 0,823988 | O15126;A0 | SCAMP1 |
| 0,451803 | -0,961444 | 0,182871 | 0,182871 | 0,7504497 | 0,824979 | O15031;A6 | PLXNB2 |
| -1,05225 | -0,0245792 | 0,694716 | -0,024579 | 0,8780075 | 0,825066 | Q9NPH3;C9 | IL1RAP |
| -0,184848 | 0,444878 | -0,462303 | -0,184848 | 0,4648501 | 0,825095 | A1L4K2;B5E | MAPK8 |
| 1,25767 | 0,445226 | -1,16852 | 0,445226 | 1,234952 | 0,826042 | A0A1W2PP | TCEA1 |
| 0,952325 | -0,664908 | 0,0629799 | 0,0629799 | 0,8099586 | 0,826079 | Q68DA7;H0 | FMN1 |
| -1,08961 | 0,946143 | -0,300535 | -0,300535 | 1,0264125 | 0,826093 | G3V1C3 | API5 |
| -0,557105 | 0,490859 | 0,308364 | 0,308364 | 0,5598466 | 0,826134 | Q16576;E9I | RBBP7 |
| -1,09931 | -0,0829966 | 1,8219 | -0,082997 | 1,4829583 | 0,826593 | P51153 | RAB13 |
| -0,123923 | -1,34922 | 0,97285 | -0,123923 | 1,1616277 | 0,826831 | F8WAD8;Q9 | ADAM22 |
| -0,445574 | 1,61341 | -0,63213 | -0,445574 | 1,2461051 | 0,827135 | O15347;E7I | HMGB3 |
| 1,74936 | -2,07857 | -0,495842 | -0,495842 | 1,9234954 | 0,827514 | Q9H5L6;D6 | THAP9 |
| 1,9588 | -1,1897 | -1,60361 | -1,1897 | 1,9482958 | 0,827748 | Q9UFE4 | CCDC39 |
| -1,38297 | 1,83369 | -1,22736 | -1,22736 | 1,8138882 | 0,827814 | Q9P265 | DIP2B |
| -1,92923 | 3,97164 | -3,77484 | -1,92923 | 4,0462842 | 0,827818 | P27482 | CALML3 |
| 1,71704 | -1,48125 | 0,453425 | 0,453425 | 1,6108357 | 0,827931 | O95433;G3 | AHSA1 |
| -2,50069 | 2,26755 | -0,799227 | -0,799227 | 2,4164785 | 0,828182 | Q8TBC5;M0 | ZSCAN18 |
| -1,74017 | 2,89797 | -0,151518 | -0,151518 | 2,3571004 | 0,8283 | Q6ZRK6;H0 | CCDC73 |
| 0,46585 | -0,88621 | 0,120525 | 0,120525 | 0,7024755 | 0,828336 | E5RHU1;Q9 | TMEM68 |
| -0,175512 | 0,342911 | -0,315131 | -0,175512 | 0,3467168 | 0,828624 | P61978;Q5 | HNRNPK |
| 3,94269 | -1,99246 | -3,64256 | -1,99246 | 3,9892533 | 0,829352 | P11387 | TOP1 |
| 0,734069 | -1,73132 | 0,42912 | 0,42912 | 1,3440386 | 0,829946 | Q69YQ0;C9 | SPECC1L |
| -1,71382 | -2,23613 | 2,78359 | -1,71382 | 2,7597431 | 0,829973 | A0A0A0MS | C10orf90 |
| -2,09811 | 1,48648 | 1,48475 | 1,48475 | 2,0690648 | 0,830225 | Q66K14 | TBC1D9B |
| 1,29673 | -0,724369 | -0,135772 | -0,135772 | 1,039499 | 0,831001 | Q15746;F8Y | MYLK |
| -0,0435993 | -1,78114 | 1,19709 | -0,043599 | 1,4960064 | 0,831179 | P63010;A0/ | AP2B1 |
| 2,66957 | -3,09382 | 1,72018 | 1,72018 | 3,0901079 | 0,831244 | Q9UBW7 | ZMYM2 |
| -1,80299 | -0,672413 | 1,72447 | -0,672413 | 1,8012138 | 0,832212 | P60981;F6F | DSTN |
| 2,94868 | -1,72041 | -0,239338 | -0,239338 | 2,3859811 | 0,833163 | Q99666 | RGPD5 |

| | | | | | | | |
|---|---|---|---|---|---|---|---|
| 0,0573773 | 0,801097 | -1,29994 | 0,0573773 | 1,065347 | 0,833198 | E9PM92;O( | C11orf58 |
| 3,24995 | -2,87671 | -1,72027 | -1,72027 | 3,2551601 | 0,833422 | A6NHJ4 | ZNF860 |
| -0,406877 | 0,233192 | 0,340921 | 0,233192 | 0,4042474 | 0,833467 | Q9NZK5;B4 | ADA2 |
| -0,056507 | -0,372714 | 0,644194 | -0,056507 | 0,5204278 | 0,833713 | P61081;M0 | UBE2M |
| 0,53406 | 0,34582 | -0,623999 | 0,34582 | 0,6214343 | 0,834226 | Q9NSE4 | IARS2 |
| 0,244808 | 0,321704 | -0,831994 | 0,244808 | 0,6450368 | 0,834297 | P25205;J3K | MCM3 |
| -1,5888 | -0,0486393 | 1,08678 | -0,048639 | 1,3428825 | 0,834893 | Q13535;H0 | ATR |
| 2,08237 | 0,287236 | -1,61219 | 0,287236 | 1,8475253 | 0,83493 | Q15751 | HERC1 |
| -0,236534 | 0,18313 | 0,148324 | 0,148324 | 0,2328966 | 0,83587 | P27695;G3 | APEX1 |
| -1,44849 | 0,682989 | 1,36331 | 0,682989 | 1,4669844 | 0,83589 | O43663 | PRC1 |
| -3,22247 | 1,54205 | 0,650889 | 0,650889 | 2,5330393 | 0,836309 | P10074;Q5 | ZBTB48 |
| 0,533306 | 0,142353 | -0,999105 | 0,142353 | 0,7962468 | 0,836398 | P54764;E9I | EPHA4 |
| 3,00643 | -3,42301 | -0,898808 | -0,898808 | 3,2393461 | 0,836456 | P04049 | RAF1 |
| -2,8532 | 0,976032 | 0,980205 | 0,976032 | 2,2120138 | 0,836679 | Q14126;J3I | DSG2 |
| 1,02262 | 1,37594 | -3,49802 | 1,02262 | 2,7177352 | 0,83705 | Q01432;E9I | AMPD3 |
| 2,12513 | 0,636988 | -1,93229 | 0,636988 | 2,0525762 | 0,837153 | K7ESP5 | HEATR6 |
| -0,609746 | 1,34897 | -1,29176 | -0,609746 | 1,3708371 | 0,837631 | Q32P51 | HNRNPA1L2 |
| -0,568403 | 0,708298 | -0,421368 | -0,421368 | 0,6985378 | 0,837679 | Q13043;F5I | STK4 |
| 0,779396 | -0,890895 | -0,225609 | -0,225609 | 0,8408837 | 0,838483 | O94759;E9I | TRPM2 |
| 0,622027 | -2,08048 | 0,811418 | 0,622027 | 1,6177395 | 0,83885 | P16615;H7( | ATP2A2 |
| 0,508867 | 1,32446 | -2,67702 | 0,508867 | 2,1145098 | 0,839228 | Q15477;H7 | SKIV2L |
| 0,529442 | -3,4903 | 4,56834 | 0,529442 | 4,0293238 | 0,83925 | F5H881;Q9 | MMP17 |
| -1,42053 | 0,0115781 | 2,11768 | 0,0115781 | 1,7797719 | 0,839537 | Q96AH8 | RAB7B |
| -0,475188 | -1,18146 | 2,41351 | -0,475188 | 1,9046962 | 0,839869 | Q8NEZ2;E5 | VPS37A |
| 1,32662 | -1,43997 | -0,443339 | -0,443339 | 1,4011929 | 0,839897 | Q9H4A5;Q5 | GOLPH3L |
| 1,48325 | 0,676739 | -1,53867 | 0,676739 | 1,5647416 | 0,839985 | A0A0A0MS | TOR1AIP1 |
| -1,04741 | 0,253663 | 0,468832 | 0,253663 | 0,8203737 | 0,840383 | P12270 | TPR |
| -0,219607 | 1,46599 | -1,91304 | -0,219607 | 1,6895165 | 0,840963 | Q6ZN16 | MAP3K15 |
| 0,613021 | 1,52503 | -1,52124 | 0,613021 | 1,5634679 | 0,840988 | O95671 | ASMTL |
| -1,40436 | -0,898459 | 1,65599 | -0,898459 | 1,6404719 | 0,841076 | Q9Y2Z0 | SUGT1 |
| -0,161004 | 0,712923 | -0,332755 | -0,161004 | 0,5607568 | 0,842435 | E7ENI6;B9Z | ICA1 |
| -1,72149 | -2,52427 | 3,06948 | -1,72149 | 3,024563 | 0,843193 | P33151;H3I | CDH5 |
| 2,55712 | -1,37036 | -0,394179 | -0,394179 | 2,0448388 | 0,843707 | Q9NPF5;Q5 | DMAP1 |
| -1,7591 | -0,577782 | 1,66358 | -0,577782 | 1,7384837 | 0,843827 | D6RAR6;D6 | FASTKD3 |
| -2,67739 | 3,94637 | -2,74469 | -2,67739 | 3,8438048 | 0,845156 | E7EVZ1;Q8( | ZFHX4 |
| -1,83819 | 1,63698 | -0,470899 | -0,470899 | 1,7506877 | 0,84516 | Q14498;G3 | RBM39 |
| 0,488973 | -0,937377 | 0,161935 | 0,161935 | 0,7472076 | 0,845366 | P63261;P6( | ACTG1 |
| -1,77075 | 0,700171 | 0,540444 | 0,540444 | 1,3827859 | 0,845369 | Q96FJ2 | DYNLL2 |
| 1,08216 | -1,42337 | -0,137167 | -0,137167 | 1,2529137 | 0,845986 | P60891;B1 | PRPS1 |
| -1,20977 | 0,107379 | 0,727485 | 0,107379 | 0,989307 | 0,847109 | Q9P227;A0 | ARHGAP23 |
| -0,550129 | 0,0558771 | 0,324556 | 0,0558771 | 0,4480524 | 0,847196 | P08581 | MET |
| -3,26777 | 1,5228 | 2,97733 | 1,5228 | 3,2676825 | 0,847829 | Q96FT9;G3 | IFT43 |
| -0,223259 | -3,08226 | 2,29188 | -0,223259 | 2,6889029 | 0,847893 | P49770 | EIF2B2 |
| -0,937846 | -1,60757 | 3,617 | -0,937846 | 2,8428649 | 0,847905 | P68431 | HIST1H3A |
| -3,36126 | 4,28021 | 0,518579 | 0,518579 | 3,8208874 | 0,848185 | Q9ULA0;A0 | DNPEP |
| 2,76274 | -2,05228 | -1,71991 | -1,71991 | 2,689146 | 0,848522 | O75436;S4I | VPS26A |
| 1,10961 | -1,25718 | -0,298768 | -0,298768 | 1,1905025 | 0,848704 | Q96C90 | PPP1R14B |
| -1,2711 | 1,70312 | 0,125274 | 0,125274 | 1,4880324 | 0,848861 | Q96G01;A8 | BICD1 |
| 2,32965 | -1,40797 | -0,208049 | -0,208049 | 1,9082946 | 0,849078 | Q96DT5;A0 | DNAH11 |
| 2,86476 | -3,00964 | 1,27995 | 1,27995 | 3,0392099 | 0,849272 | Q9NZC9;C9 | SMARCAL1 |
| -1,38296 | 0,63168 | 0,344338 | 0,344338 | 1,0897169 | 0,849286 | F5H212;A0 | HIVEP1 |
| -0,736729 | -0,129923 | 0,61411 | -0,129923 | 0,6765802 | 0,849355 | P51665;H3I | PSMD7 |

| | | | | | | | |
|---|---|---|---|---|---|---|---|
| -1,79636 | 1,44903 | 1,00369 | 1,00369 | 1,7593163 | 0,849429 | Q9NQ76;D6 | MEPE |
| -0,366698 | -0,160647 | 0,747964 | -0,160647 | 0,5930855 | 0,849859 | Q6R327 | RICTOR |
| 0,785215 | -0,983362 | 0,55555 | 0,55555 | 0,9616705 | 0,849992 | O76094;D6 | SRP72 |
| -0,489277 | -0,0690584 | 0,801913 | -0,069058 | 0,6585776 | 0,8507 | O75306 | NDUFS2 |
| -1,39901 | -1,77645 | 2,33653 | -1,39901 | 2,2735187 | 0,851036 | Q5W041 | ARMC3 |
| -3,26067 | 4,83288 | -3,29781 | -3,26067 | 4,6835715 | 0,85126 | B4DTR8;Q9 | LIPG |
| 1,85459 | -1,72517 | -0,813916 | -0,813916 | 1,8603758 | 0,851458 | E5RFK7 | UBR5 |
| 1,31644 | -2,5002 | 2,08681 | 1,31644 | 2,456314 | 0,851572 | Q5W5X9 | TTC23 |
| -0,295938 | -0,806324 | 1,55951 | -0,295938 | 1,2450135 | 0,851723 | Q14019 | COTL1 |
| -0,0953559 | 1,11966 | -0,686182 | -0,095356 | 0,9207248 | 0,851733 | Q8NBX0 | SCCPDH |
| -0,568167 | -0,634946 | 1,68697 | -0,568167 | 1,3217032 | 0,852189 | Q96HA7 | TONSL |
| -1,39848 | -0,232765 | 2,32835 | -0,232765 | 1,9064567 | 0,852358 | O75382;E9I | TRIM3 |
| -1,00073 | -1,00716 | 1,48398 | -1,00073 | 1,4364078 | 0,852722 | Q6ZRP7;H0 | QSOX2 |
| 4,30358 | -3,56707 | 0,691109 | 0,691109 | 3,939737 | 0,853658 | A8MT87;P1 | PNMT |
| 0,481015 | -1,52489 | 0,610188 | 0,481015 | 1,1971423 | 0,853697 | Q9NTJ4;B4I | MAN2C1 |
| -0,169435 | 0,429306 | -0,417223 | -0,169435 | 0,4352204 | 0,853981 | C9JRZ6;Q9N | CHCHD3 |
| 0,130797 | -1,23493 | 1,62047 | 0,130797 | 1,4281483 | 0,853983 | P17812 | CTPS1 |
| -0,0964063 | 1,48465 | -2,02275 | -0,096406 | 1,7565304 | 0,854109 | P18065;C9J | IGFBP2 |
| -1,88167 | 0,723057 | 0,626573 | 0,626573 | 1,4767755 | 0,854486 | P30085;Q5 | CMPK1 |
| -1,28437 | 3,33346 | -3,25483 | -1,28437 | 3,3816325 | 0,855955 | Q16853 | AOC3 |
| 0,148711 | -0,45578 | 0,179428 | 0,148711 | 0,3581997 | 0,856041 | Q14BN4;H7 | SLMAP |
| 3,23651 | -1,59953 | -2,76462 | -1,59953 | 3,1821973 | 0,856824 | Q6ZS81 | WDFY4 |
| 0,0472087 | 2,70245 | -3,93209 | 0,0472087 | 3,3392176 | 0,856925 | E9PFK9;Q9 | RABGEF1 |
| -0,0797528 | 1,16285 | -1,56549 | -0,079753 | 1,3659744 | 0,857303 | Q9NZM3;A | ITSN2 |
| -0,97957 | -1,54244 | 1,8761 | -0,97957 | 1,8329434 | 0,857605 | O60336 | MAPKBP1 |
| 2,58175 | -2,51188 | -0,989075 | -0,989075 | 2,6145361 | 0,857925 | P16152;A8I | CBR1 |
| -1,49684 | 1,44262 | -0,469841 | -0,469841 | 1,4917919 | 0,858035 | Q96K17;E9I | BTF3L4 |
| -1,01791 | -1,76148 | 3,84626 | -1,01791 | 3,0457564 | 0,858438 | Q96Q89;AC | KIF20B |
| 2,66892 | -0,429256 | -3,27958 | -0,429256 | 2,9751105 | 0,858732 | Q4J6C6;H7 | PREPL |
| -2,94407 | 2,35463 | 1,59044 | 1,59044 | 2,864204 | 0,858755 | P24394 | IL4R |
| 2,56381 | 0,0666111 | -3,73643 | 0,0666111 | 3,1725948 | 0,859099 | Q8IX12;A0 | CCAR1 |
| -1,4147 | 1,4802 | -0,584958 | -0,584958 | 1,4907378 | 0,859162 | Q96PC5;G3 | MIA2 |
| 1,643 | 0,167881 | -1,29853 | 0,167881 | 1,4707671 | 0,859202 | B1AK53;A0 | ESPN |
| 0,0899262 | -0,526611 | 0,639324 | 0,0899262 | 0,5832896 | 0,859577 | O75964;E9I | ATP5MG |
| -0,201659 | 1,68137 | -1,0015 | -0,201659 | 1,377397 | 0,859667 | Q12901;K7 | ZNF155 |
| 0,971518 | -1,06571 | -0,260407 | -0,260407 | 1,026032 | 0,860292 | O75592;H7 | MYCBP2 |
| 2,31152 | -0,699402 | -2,44344 | -0,699402 | 2,405444 | 0,860293 | J3KNE0;A6 | RGPD3 |
| -2,6864 | 2,3007 | -0,47224 | -0,47224 | 2,4987619 | 0,861188 | Q16836;A0 | HADH |
| -0,0909007 | -1,2577 | 1,89593 | -0,090901 | 1,5944852 | 0,86122 | Q9H2T7 | RANBP17 |
| -0,785313 | 0,564224 | 0,479951 | 0,479951 | 0,7560032 | 0,861558 | P99999;C9J | CYCS |
| -1,39368 | -0,038087 | 1,01802 | -0,038087 | 1,2089452 | 0,861626 | Q68DK2;A0 | ZFYVE26 |
| -3,0971 | 2,57986 | 1,55247 | 1,55247 | 3,0249492 | 0,861627 | Q5JSH3 | WDR44 |
| 0,66094 | -0,805297 | -0,106559 | -0,106559 | 0,7333872 | 0,861668 | K7EP59;Q1 | STK11 |
| 3,81121 | -2,79775 | 0,118928 | 0,118928 | 3,3120565 | 0,86176 | O94906 | PRPF6 |
| 0,67149 | 2,04424 | -3,74907 | 0,67149 | 3,0273265 | 0,861984 | Q9BWS9 | CHID1 |
| 1,08739 | -3,3882 | 1,39021 | 1,08739 | 2,6756871 | 0,862385 | Q96J02 | ITCH |
| 4,23965 | -2,56717 | -3,05523 | -2,56717 | 4,0781179 | 0,862885 | Q16787;A0 | LAMA3 |
| -0,446317 | 0,142575 | 0,459089 | 0,142575 | 0,4594807 | 0,863271 | Q99426;K7 | TBCB |
| 1,83488 | -1,16796 | -1,2614 | -1,16796 | 1,761284 | 0,863495 | Q96HH9;D6 | GRAMD2B |
| -2,11247 | 1,26312 | 1,53473 | 1,26312 | 2,0318484 | 0,863578 | Q06136;K7 | KDSR |
| 1,86136 | 1,88701 | -2,83314 | 1,86136 | 2,7178056 | 0,863803 | Q15572 | TAF1C |
| 1,10514 | -3,52605 | 3,63318 | 1,10514 | 3,6307364 | 0,864938 | X6RHV1 | SETDB1 |

| | | | | | | | |
|---|---|---|---|---|---|---|---|
| 1,08905 | 0,354614 | -1,97609 | 0,354614 | 1,6003469 | 0,865414 | O15085 | ARHGEF11 |
| -0,990086 | -0,160647 | 0,845045 | -0,160647 | 0,9189751 | 0,865436 | H3BP42 | PYCARD |
| 1,00759 | -1,62186 | 0,167872 | 0,167872 | 1,3430236 | 0,865539 | K7ES89 | DUSP3 |
| -1,28629 | -0,418113 | 1,27202 | -0,418113 | 1,300976 | 0,865549 | Q4LDE5;A0 | SVEP1 |
| -3,39617 | 4,11093 | 0,529605 | 0,529605 | 3,7548668 | 0,865928 | Q8IVL0;A0A | NAV3 |
| -0,195482 | -0,371953 | 0,771219 | -0,195482 | 0,615426 | 0,866037 | Q9UBN7;A( | HDAC6 |
| -1,13217 | 1,86003 | -0,221147 | -0,221147 | 1,5337602 | 0,866337 | Q08043;A0 | ACTN3 |
| -0,749059 | 2,14006 | -0,83263 | -0,749059 | 1,6926743 | 0,866534 | P07900 | HSP90AA1 |
| 0,99382 | -0,310371 | -1,01969 | -0,310371 | 1,0212958 | 0,866792 | Q8WYA0;H | IFT81 |
| -0,603946 | -0,507906 | 1,50315 | -0,507906 | 1,1897775 | 0,866929 | A0A0J9YXM | WDR81 |
| 0,07272 | 1,36163 | -1,0393 | 0,07272 | 1,2015505 | 0,866969 | P54886 | ALDH18A1 |
| -2,50192 | 4,34672 | -3,21014 | -2,50192 | 4,1735592 | 0,867621 | Q8TD23 | ZNF675 |
| 1,5164 | -2,15326 | 0,035792 | 0,035792 | 1,8461922 | 0,868243 | C9J6P4;Q7Z | ZC3HAV1 |
| 1,43817 | -1,51282 | 0,567227 | 0,567227 | 1,5162167 | 0,868523 | P61764;A0A | STXBP1 |
| -1,66368 | 1,65516 | -0,537447 | -0,537447 | 1,6877315 | 0,869074 | A0A0G2JQF | KANSL1 |
| 1,42046 | -2,00399 | 0,0279477 | 0,0279477 | 1,7221459 | 0,869423 | P68363;C9J | TUBA1B |
| -0,443039 | 0,627194 | -0,010511 | -0,010511 | 0,5383844 | 0,869455 | O15212 | PFDN6 |
| 3,15666 | -3,17794 | 1,05778 | 1,05778 | 3,226809 | 0,869978 | O60229;H7 | KALRN |
| 1,74331 | -1,54958 | 0,335435 | 0,335435 | 1,6521964 | 0,870349 | Q14586;I3L | ZNF267 |
| 1,58371 | -2,6072 | 0,334529 | 0,334529 | 2,1516641 | 0,870381 | A0A0C4DG | ZSCAN5A |
| 2,01725 | -1,04165 | -1,59646 | -1,04165 | 1,9460894 | 0,870848 | Q14CX7 | NAA25 |
| 1,46304 | -2,94126 | 2,37229 | 1,46304 | 2,841901 | 0,872609 | Q9Y5F7 | PCDHGC4 |
| -0,798366 | 1,26429 | -0,136225 | -0,136225 | 1,0531241 | 0,873222 | Q96A65 | EXOC4 |
| 0,916045 | -2,26352 | 2,04439 | 0,916045 | 2,2338638 | 0,873656 | Q9Y623 | MYH4 |
| 1,00387 | -2,85429 | 2,74385 | 1,00387 | 2,86508 | 0,873714 | Q9Y285;K7I | FARSA |
| 1,74933 | -2,09007 | 0,973781 | 0,973781 | 2,0301745 | 0,87372 | O43896 | KIF1C |
| -3,17762 | 1,46125 | 2,67739 | 1,46125 | 3,0897479 | 0,874032 | P27449 | ATP6V0C |
| -1,51303 | -0,387838 | 2,55272 | -0,387838 | 2,0993357 | 0,874244 | H3BPZ3 | PLCG2 |
| -1,91273 | 2,30258 | 0,264591 | 0,264591 | 2,1080388 | 0,874265 | Q92608;E7I | DOCK2 |
| 3,27086 | -2,98754 | 0,692822 | 0,692822 | 3,1453382 | 0,874307 | P98174 | FGD1 |
| 0,287193 | 2,39289 | -3,62818 | 0,287193 | 3,0555248 | 0,87433 | Q9NYF5;D6 | FAM13B |
| 0,445845 | 1,28766 | -1,32121 | 0,445845 | 1,3314991 | 0,874584 | Q7L014;A0, | DDX46 |
| -1,2981 | 1,05233 | 0,631527 | 0,631527 | 1,2533327 | 0,875328 | Q9BT09;A0 | CNPY3 |
| -1,04317 | 0,73495 | 0,612334 | 0,612334 | 0,9930961 | 0,875949 | E7EVA0;B5I | MAP4 |
| -0,714218 | 0,583412 | 0,34208 | 0,34208 | 0,690151 | 0,875989 | Q9HC77;F6 | CENPJ |
| 1,49854 | -0,726051 | -0,405633 | -0,405633 | 1,2025908 | 0,876416 | Q9UBU9;E9 | NXF1 |
| 0,25828 | -0,928389 | 0,443212 | 0,25828 | 0,7442751 | 0,876496 | Q99798;A2 | ACO2 |
| 0,235795 | -1,53071 | 0,913512 | 0,235795 | 1,261881 | 0,877536 | P09543;K7E | CNP |
| -0,448806 | 0,0818247 | 0,512407 | 0,0818247 | 0,4814735 | 0,877618 | P82909 | MRPS36 |
| -3,19385 | 2,66411 | 1,46153 | 1,46153 | 3,0939311 | 0,877968 | P05386 | RPLP1 |
| -0,752136 | 1,15355 | -0,109934 | -0,109934 | 0,969575 | 0,878182 | H7C3P7;P1 | RALA |
| -0,727885 | 1,3883 | -1,05838 | -0,727885 | 1,3275108 | 0,878523 | Q6NSJ2 | PHLDB3 |
| 0,649169 | -3,24299 | 1,80226 | 0,649169 | 2,6436423 | 0,878664 | Q6IN85;G3 | PPP4R3A |
| 3,08772 | -3,29829 | -0,750458 | -0,750458 | 3,2146587 | 0,878852 | Q14511 | NEDD9 |
| 1,68218 | -2,27634 | 0,0012629 | 0,0012629 | 1,986741 | 0,879062 | Q08J23 | NSUN2 |
| -1,88286 | 2,26527 | 0,236562 | 0,236562 | 2,0742303 | 0,879068 | O75843;H0 | AP1G2 |
| -1,93152 | 0,437293 | 2,0955 | 0,437293 | 2,0239325 | 0,879599 | Q14674 | ESPL1 |
| 1,02139 | -0,595275 | -0,713609 | -0,595275 | 0,9693495 | 0,879798 | P62277;J3K | RPS13 |
| 1,20718 | 2,81114 | -3,1108 | 1,20718 | 3,0628694 | 0,879912 | A0A2R8YH( | FAM208B |
| 0,666089 | 2,02665 | -3,55164 | 0,666089 | 2,9085415 | 0,880311 | O43719;Q5 | HTATSF1 |
| 2,99292 | -3,11234 | -0,787429 | -0,787429 | 3,0814079 | 0,880712 | Q16875;A0 | PFKFB3 |
| -0,985828 | 2,2583 | -0,741903 | -0,741903 | 1,8067044 | 0,880964 | Q9Y2F5 | ICE1 |

| | | | | | | | |
|---|---|---|---|---|---|---|---|
| 0,316128 | -0,11239 | -0,294742 | -0,11239 | 0,3135926 | 0,88235 | A0A1B0GTU | PXN |
| -0,0498048 | -3,05091 | 2,32339 | -0,049805 | 2,6932566 | 0,88298 | Q9P2D3 | HEATR5B |
| -0,779506 | 1,69269 | -0,522438 | -0,522438 | 1,359205 | 0,883437 | Q9Y281;F8V | CFL2 |
| -1,69876 | -0,498288 | 1,70279 | -0,498288 | 1,725129 | 0,883826 | A0A087WV | MTHFD1L |
| -0,945189 | -0,382282 | 1,73178 | -0,382282 | 1,4113992 | 0,883845 | P25787;A0A | PSMA2 |
| 1,06863 | -1,98498 | 1,45253 | 1,06863 | 1,8836307 | 0,884567 | Q6DKJ4;A0A | NXN |
| -0,344529 | -3,17219 | 2,68427 | -0,344529 | 2,9288056 | 0,884738 | Q9Y2Q0;H0 | ATP8A1 |
| 0,329838 | -0,762583 | 0,641127 | 0,329838 | 0,7371891 | 0,885361 | O43295;A0 | SRGAP3 |
| -0,534779 | -0,56973 | 0,872532 | -0,534779 | 0,8227865 | 0,885653 | Q99536;K7I | VAT1 |
| -0,426181 | -0,135117 | 0,437618 | -0,135117 | 0,4394869 | 0,885861 | P18859;A8I | ATP5PF |
| -0,800807 | 0,174763 | 0,442299 | 0,174763 | 0,6542967 | 0,886098 | O43390;B4 | HNRNPR |
| 0,247295 | 0,775589 | -0,79809 | 0,247295 | 0,8008735 | 0,886155 | Q12791;A0 | KCNMA1 |
| -1,70373 | 1,31021 | -0,0297545 | -0,029755 | 1,5100515 | 0,886306 | A6NNF4;A0 | ZNF726 |
| -1,34857 | 1,15726 | -0,159762 | -0,159762 | 1,2534616 | 0,886397 | Q15046;H3 | KARS |
| 2,01137 | 0,0893003 | -1,59659 | 0,0893003 | 1,8052679 | 0,886739 | M0R2L6;A6 | RPL37A |
| 1,46755 | 0,0156142 | -1,12234 | 0,0156142 | 1,2981132 | 0,887246 | Q6PKG0;A0 | LARP1 |
| -2,41376 | 2,05111 | 1,00545 | 1,00545 | 2,3352183 | 0,888327 | Q96S59 | RANBP9 |
| -1,15965 | 0,548163 | 0,915185 | 0,548163 | 1,1072692 | 0,888725 | Q9H4B7 | TUBB1 |
| 1,58071 | -3,17793 | 2,42171 | 1,58071 | 3,0196 | 0,889216 | B2R5W2;B4 | HNRNPC |
| -0,0314317 | -2,98353 | 3,96638 | -0,031432 | 3,4880422 | 0,889327 | Q8TBZ2;H0 | MYCBPAP |
| 0,0595213 | 1,92851 | -1,52287 | 0,0595213 | 1,7276721 | 0,89074 | Q8N4Y2;E9 | CRACR2B |
| -2,36753 | 4,43135 | -3,18542 | -2,36753 | 4,181485 | 0,891145 | Q86YT6 | MIB1 |
| -1,85681 | 1,10454 | 1,21968 | 1,10454 | 1,7439248 | 0,891231 | P36578;H3I | RPL4 |
| -2,93692 | 2,48631 | 1,20808 | 1,20808 | 2,8350873 | 0,89157 | Q76I76;F5H | SSH2 |
| 0,680268 | -1,90988 | 1,72968 | 0,680268 | 1,873345 | 0,891664 | Q9Y608;C9 | LRRFIP2 |
| -1,01801 | 0,675201 | 0,113018 | 0,113018 | 0,8623841 | 0,891855 | P11908;H7 | PRPS2 |
| -0,720145 | 0,98193 | -0,507417 | -0,507417 | 0,9274038 | 0,892498 | E9PIK4;E9P | TP53I11 |
| 2,1121 | -2,10751 | 0,557386 | 0,557386 | 2,134007 | 0,893107 | Q9UH77;D6 | KLHL3 |
| 3,04913 | -2,33868 | -0,00024514 | -0,000245 | 2,7017113 | 0,893295 | O00291;C9 | HIP1 |
| 5,15863 | -1,28813 | -2,76528 | -1,28813 | 4,2136887 | 0,893529 | Q86WZ0 | HEATR4 |
| -0,299701 | 1,34145 | -1,40392 | -0,299701 | 1,3814083 | 0,893575 | F8VVM2;F8 | SLC25A3 |
| -2,15286 | -2,30512 | 3,57997 | -2,15286 | 3,3546686 | 0,893755 | J3KT75 | MPDU1 |
| 2,35701 | -1,21804 | -0,638098 | -0,638098 | 1,9186795 | 0,894027 | A0FGR9 | ESYT3 |
| 1,59895 | 1,41914 | -2,42475 | 1,41914 | 2,2729564 | 0,89403 | Q9HC10;A0 | OTOF |
| 0,743321 | -0,0930617 | -0,487233 | -0,093062 | 0,6283802 | 0,894673 | Q9NVA2;D6 | SEPT11 |
| 0,850097 | 0,333752 | -1,5044 | 0,333752 | 1,2375427 | 0,894841 | P16885;A0A | PLCG2 |
| -2,61039 | -0,309378 | 2,28599 | -0,309378 | 2,4496642 | 0,894963 | O95477 | ABCA1 |
| -1,41697 | -0,365687 | 2,27474 | -0,365687 | 1,9020065 | 0,894963 | B7Z524;H0Y | HBS1L |
| 0,508188 | 2,12628 | -3,36434 | 0,508188 | 2,8213945 | 0,894974 | O14977 | AZIN1 |
| -1,55245 | 0,977465 | 0,239185 | 0,239185 | 1,3009921 | 0,895207 | H7C3T2;Q2 | ATG2A |
| 1,38329 | -1,21665 | -0,513308 | -0,513308 | 1,3448334 | 0,895338 | P37268;A0A | FDFT1 |
| -2,09081 | -0,697353 | 2,22109 | -0,697353 | 2,2004361 | 0,895367 | P68371;M0 | TUBB4B |
| 0,58535 | 0,404608 | -1,25056 | 0,404608 | 1,0118312 | 0,895429 | C9JAV2;H7 | DCUN1D2 |
| 1,5241 | 0,1349 | -2,13469 | 0,1349 | 1,8469642 | 0,895432 | O60285 | NUAK1 |
| -0,535521 | 1,05588 | -0,775942 | -0,535521 | 0,9954841 | 0,895756 | Q9NZL3 | ZNF224 |
| 0,311319 | 1,7387 | -1,61942 | 0,311319 | 1,6853357 | 0,896255 | Q7L099;D6 | RUFY3 |
| -1,96312 | 1,59121 | -0,0805567 | -0,080557 | 1,7782065 | 0,896677 | Q8IXB1;A0A | DNAJC10 |
| 0,616088 | 0,185106 | -1,0153 | 0,185106 | 0,8453941 | 0,897154 | Q92688 | ANP32B |
| -1,49873 | -0,306377 | 1,43285 | -0,306377 | 1,4742669 | 0,897459 | Q5JU85;A0 | IQSEC2 |
| 2,25796 | -0,300394 | -2,56612 | -0,300394 | 2,4135188 | 0,897604 | J3KR97;Q9I | TBCD |
| 0,0966954 | -0,641033 | 0,714415 | 0,0966954 | 0,6786089 | 0,898213 | P61158;B4I | ACTR3 |
| 0,0387589 | -3,45078 | 2,64591 | 0,0387589 | 3,058969 | 0,898285 | A5D8W1 | CFAP69 |

| | | | | | | | |
|---|---|---|---|---|---|---|---|
| 0,327052 | -0,934454 | 0,834726 | 0,327052 | 0,9109636 | 0,89865 | O60568;H7 | PLOD3 |
| 2,74107 | -1,29108 | -0,896103 | -0,896103 | 2,2227336 | 0,898791 | E7END4 | LOXL3 |
| -2,42394 | 0,37323 | 1,54348 | 0,37323 | 2,038548 | 0,89894 | F8VVA7;F8' | COPZ1 |
| 0,112939 | 0,498322 | -0,487909 | 0,112939 | 0,4970228 | 0,899196 | Q92747;A0 | ARPC1A |
| -0,064748 | 1,66286 | -2,05845 | -0,064748 | 1,8622399 | 0,899592 | Q96AX1;A0 | VPS33A |
| -1,24581 | 3,45671 | -3,03942 | -1,24581 | 3,3548581 | 0,899687 | M0QXN5;P | NUP62 |
| 2,61193 | -1,42972 | -1,78366 | -1,42972 | 2,4420422 | 0,899958 | Q9UBT7 | CTNNAL1 |
| 0,882228 | 0,0560731 | -1,19581 | 0,0560731 | 1,046262 | 0,900024 | A0A0D9SF3 | NCAM1 |
| 2,23608 | -2,12304 | 0,422584 | 0,422584 | 2,189783 | 0,900635 | P51587;H0' | BRCA2 |
| -1,2369 | 0,45825 | 1,06912 | 0,45825 | 1,1947401 | 0,90123 | Q8TAA3;A0 | PSMA8 |
| -0,245923 | 0,606847 | -0,502038 | -0,245923 | 0,5805798 | 0,901257 | P78527;H0' | PRKDC |
| 1,18176 | -0,70366 | -0,23947 | -0,23947 | 0,982359 | 0,901314 | E7EM64;Q | COPS6 |
| -2,6642 | 1,37602 | 0,760231 | 0,760231 | 2,1767446 | 0,901465 | P62136;E9I | PPP1CA |
| -0,0299626 | -0,582062 | 0,777731 | -0,029963 | 0,6838884 | 0,901562 | Q29RF7;H0 | PDS5A |
| 4,79885 | -0,143989 | -3,63128 | -0,143989 | 4,2359562 | 0,901827 | Q16760;H7 | DGKD |
| -2,67391 | 1,37982 | 0,7675 | 0,7675 | 2,1852142 | 0,902093 | P62140;E7I | PPP1CB |
| -2,67391 | 1,37982 | 0,767508 | 0,767508 | 2,1852159 | 0,902095 | P36873;F8\ | PPP1CC |
| 1,39903 | -1,16812 | 0,0754651 | 0,0754651 | 1,2837826 | 0,903031 | F8VSL2;F8V | PRKAG1 |
| -0,630498 | 2,69431 | -1,53378 | -0,630498 | 2,2266197 | 0,903275 | P49368;B4I | CCT3 |
| 0,237116 | 1,46212 | -2,13431 | 0,237116 | 1,8284148 | 0,903313 | Q6ZT07 | TBC1D9 |
| -0,608719 | -2,36976 | 2,40822 | -0,608719 | 2,4163429 | 0,904098 | H3BQ65;Q3 | MGAT5B |
| 0,37774 | -1,14278 | 0,546789 | 0,37774 | 0,9305198 | 0,904683 | B2RTY4;H3I | MYO9A |
| -1,74126 | 5,17365 | -2,44903 | -1,74126 | 4,2115351 | 0,905107 | O94986 | CEP152 |
| 1,37433 | -1,71445 | -0,0210231 | -0,021023 | 1,5467852 | 0,905112 | Q58FG1 | HSP90AA4P |
| 2,33916 | 0,344167 | -3,35556 | 0,344167 | 2,8895736 | 0,90545 | Q92556 | ELMO1 |
| 2,17128 | -2,22963 | 0,576391 | 0,576391 | 2,2280572 | 0,905504 | P11388 | TOP2A |
| -2,06448 | 2,89036 | -0,243921 | -0,243921 | 2,5062785 | 0,905628 | O15078;J3I | CEP290 |
| 0,797001 | -2,97953 | 2,87086 | 0,797001 | 2,9662024 | 0,905686 | Q6PIK3 | RAB4B |
| -0,230764 | 1,06794 | -1,08752 | -0,230764 | 1,085255 | 0,906244 | O43143 | DHX15 |
| -0,543457 | 1,46534 | -0,64833 | -0,543457 | 1,1912084 | 0,906657 | Q8WXX5 | DNAJC9 |
| -0,015133 | -0,554801 | 0,716229 | -0,015133 | 0,6379197 | 0,906784 | P28066 | PSMA5 |
| -1,91684 | 2,59893 | -0,161502 | -0,161502 | 2,276451 | 0,907045 | Q5BJF6;S4F | ODF2 |
| -1,22661 | 0,362448 | 0,637128 | 0,362448 | 1,0061539 | 0,908269 | P07339;A0 | CTSD |
| 0,454028 | -1,56504 | 0,821379 | 0,454028 | 1,2849499 | 0,908366 | Q16543;K7I | CDC37 |
| 1,84142 | -0,506427 | -0,994 | -0,506427 | 1,5160101 | 0,908559 | Q9GZU2;M | PEG3 |
| 0,564856 | -0,858582 | 0,130152 | 0,130152 | 0,7294676 | 0,908837 | Q460N5 | PARP14 |
| 4,42849 | -0,748353 | -2,84335 | -0,748353 | 3,7431799 | 0,909114 | Q5TAH2 | SLC9C2 |
| -0,695577 | -1,93743 | 2,16493 | -0,695577 | 2,1037291 | 0,909538 | Q8WVE0 | EEF1AKMT1 |
| 0,882453 | -0,719534 | 0,0138799 | 0,0138799 | 0,8019432 | 0,910359 | P62906 | RPL10A |
| -0,521754 | -1,02621 | 1,28106 | -0,521754 | 1,2129928 | 0,910533 | Q6ZMV9;H | KIF6 |
| -0,146835 | -0,177911 | 0,39556 | -0,146835 | 0,3224973 | 0,910715 | P63128;P6 | ERVK-9 |
| 0,652406 | 2,39895 | -2,5066 | 0,652406 | 2,4864351 | 0,910912 | A0A0A0MS | ERBB4 |
| -1,85923 | 1,19437 | 1,03911 | 1,03911 | 1,7199299 | 0,911513 | P20645;F5 | M6PR |
| 2,08987 | -1,78546 | 0,116315 | 0,116315 | 1,9377758 | 0,911708 | P13010;C9J | XRCC5 |
| -0,365634 | -0,669748 | 1,25967 | -0,365634 | 1,0373646 | 0,912073 | Q9NRF8 | CTPS2 |
| -0,227458 | 1,26936 | -1,32144 | -0,227458 | 1,3006092 | 0,912591 | Q13023;G3 | AKAP6 |
| -0,451872 | -0,11294 | 0,690903 | -0,11294 | 0,5869374 | 0,912632 | Q9UNZ2;F2 | NSFL1C |
| 2,22946 | 1,5861 | -3,1861 | 1,5861 | 2,9584933 | 0,913466 | O00472;D6 | ELL2 |
| -2,68633 | 0,883288 | 2,35289 | 0,883288 | 2,5915131 | 0,913705 | O15400 | STX7 |
| 0,081469 | 1,16019 | -1,52815 | 0,081469 | 1,3528787 | 0,913869 | Q9UBB4;A | ATXN10 |
| 0,0946959 | 0,174682 | -0,326128 | 0,0946959 | 0,2690419 | 0,914203 | Q02790;F5I | FKBP4 |
| -1,12297 | -0,611675 | 1,44784 | -0,611675 | 1,3608875 | 0,914279 | Q6BDS2;H7 | UHRF1BP1 |

| | | | | | | | |
|---|---|---|---|---|---|---|---|
| 0,86073 | -0,992181 | 0,332399 | 0,332399 | 0,954544 | 0,914372 | Q6Q0C0;H3 | TRAF7 |
| 0,213785 | -0,538222 | 0,232043 | 0,213785 | 0,4395369 | 0,914497 | Q13459;M( | MYO9B |
| -0,545794 | 0,232585 | 0,220693 | 0,220693 | 0,446004 | 0,915618 | P18621;A0/ | RPL17 |
| -0,451779 | 0,921917 | -0,313819 | -0,313819 | 0,7564299 | 0,915932 | Q10570;E9I | CPSF1 |
| 2,89208 | -2,35021 | -1,10752 | -1,10752 | 2,7393029 | 0,915996 | Q5T5C0 | STXBP5 |
| -3,06077 | 2,34047 | 1,31154 | 1,31154 | 2,8679024 | 0,916132 | P62829;J3K | RPL23 |
| -1,74065 | 0,769486 | 0,67785 | 0,67785 | 1,4235122 | 0,916177 | P31749;A0/ | AKT1 |
| 1,05351 | -1,94202 | 0,559194 | 0,559194 | 1,6059068 | 0,916574 | Q03164;E9I | KMT2A |
| 0,599685 | -1,57763 | 0,716826 | 0,599685 | 1,2922171 | 0,917784 | P23246;H0' | SFPQ |
| -2,23257 | 1,80354 | 0,0204214 | 0,0204214 | 2,0226083 | 0,917804 | Q15121 | PEA15 |
| 0,943649 | -1,22124 | 0,508773 | 0,508773 | 1,1451933 | 0,917865 | P21359;J3K | NF1 |
| -1,48505 | 0,281271 | 0,95018 | 0,281271 | 1,2581517 | 0,917988 | B7ZM68;Q6 | FGD5 |
| 0,0236407 | 0,856918 | -0,721572 | 0,0236407 | 0,7896543 | 0,918081 | Q86UE4;E5 | MTDH |
| 3,39336 | -1,51788 | -2,50935 | -1,51788 | 3,1608352 | 0,918404 | Q86YM6;Q8 | HOMER1 |
| 2,84504 | -2,6679 | -0,737563 | -0,737563 | 2,7974319 | 0,918486 | P78524;B4I | ST5 |
| 0,69374 | 2,52085 | -2,68872 | 0,69374 | 2,6431984 | 0,919045 | O75069;A0. | TMCC2 |
| -0,41467 | 3,85716 | -2,77714 | -0,41467 | 3,3626312 | 0,919484 | Q8TF32;Q9 | ZNF431 |
| 0,716483 | -0,648854 | -0,203981 | -0,203981 | 0,696337 | 0,920314 | Q9Y265;B5I | RUVBL1 |
| 0,443858 | -2,25747 | 2,25722 | 0,443858 | 2,2718524 | 0,920538 | Q13885 | TUBB2A |
| -0,472073 | 1,52751 | -0,808995 | -0,472073 | 1,263006 | 0,920594 | O94804 | STK10 |
| 1,67227 | 0,357733 | -2,43847 | 0,357733 | 2,0994024 | 0,920821 | Q969N2;A0 | PIGT |
| -2,51261 | 1,14512 | 1,81865 | 1,14512 | 2,330681 | 0,921221 | Q68CZ1;A0. | RPGRIP1L |
| -0,64543 | -2,83674 | 2,92102 | -0,64543 | 2,9061201 | 0,921413 | A0A0C4DG! | NDUFA6 |
| 3,52587 | -1,4604 | -2,70179 | -1,4604 | 3,2961515 | 0,921431 | C9JC11 | ACSL3 |
| -0,837668 | -1,30641 | 2,54878 | -0,837668 | 2,1035779 | 0,921699 | Q14CN2 | CLCA4 |
| 1,76837 | -2,68801 | 1,39237 | 1,39237 | 2,471511 | 0,922151 | Q6IBS0;D6F | TWF2 |
| -2,0243 | -0,754526 | 2,34898 | -0,754526 | 2,249802 | 0,922236 | Q8TET4;H3 | GANC |
| -1,13497 | -3,20182 | 3,6639 | -1,13497 | 3,5222893 | 0,922245 | Q9BX63;J3( | BRIP1 |
| -0,599437 | 0,923648 | -0,174102 | -0,174102 | 0,7858915 | 0,922259 | Q8IVF5;E9F | TIAM2 |
| 1,0106 | 0,745061 | -1,49452 | 0,745061 | 1,3760972 | 0,922758 | O60716;C9. | CTNND1 |
| -2,07499 | 1,44616 | 0,99141 | 0,99141 | 1,9152069 | 0,922943 | Q92626;H7 | PXDN |
| 3,02887 | -3,33876 | -0,291497 | -0,291497 | 3,184791 | 0,923139 | K7EM48;K7 | KCTD15 |
| 1,09904 | -1,77853 | 0,396805 | 0,396805 | 1,5003125 | 0,923304 | F5H039;Q9 | GPHN |
| 0,631899 | 0,745469 | -1,17552 | 0,631899 | 1,0777956 | 0,923765 | P46926;D6I | GNPDA1 |
| 2,2565 | -2,22918 | 0,394674 | 0,394674 | 2,253602 | 0,923775 | P22105;A0/ | TNXB |
| -0,133526 | -0,94099 | 0,901636 | -0,133526 | 0,9236548 | 0,92381 | P30101 | PDIA3 |
| -2,1866 | 1,02348 | 1,54075 | 1,02348 | 2,0192953 | 0,923876 | Q14974;J3K | KPNB1 |
| -0,902558 | 0,628194 | 0,429756 | 0,429756 | 0,8324302 | 0,924012 | P48681 | NES |
| 1,91023 | -2,37909 | 0,0697467 | 0,0697467 | 2,1518382 | 0,924497 | P81274 | GPSM2 |
| 1,40876 | 0,378328 | -2,12273 | 0,378328 | 1,8160627 | 0,924761 | A6NGW2;Q | STRCP1 |
| 0,250175 | -0,557302 | 0,4021 | 0,250175 | 0,5156796 | 0,925024 | H7C3G9;Q9 | NAGK |
| -2,68639 | 1,95634 | 1,18435 | 1,18435 | 2,4877546 | 0,925655 | Q9Y2E4;A0. | DIP2C |
| -1,05878 | -1,97729 | 2,5968 | -1,05878 | 2,419684 | 0,926089 | H0YDG5 | ATP2B4 |
| -0,687461 | -0,161872 | 1,00645 | -0,161872 | 0,8670404 | 0,926222 | Q9ULI3 | HEG1 |
| -0,189557 | 4,00113 | -3,16334 | -0,189557 | 3,5994183 | 0,926674 | E5RFR8;E5F | CHMP7 |
| -0,543368 | 0,554163 | 0,0883024 | 0,0883024 | 0,550849 | 0,926754 | P12236 | SLC25A6 |
| 1,15165 | -0,796871 | -0,545344 | -0,545344 | 1,0598576 | 0,926793 | Q96MU6;A( | ZNF778 |
| 0,890309 | -1,06719 | 0,00089061 | 0,0008906 | 0,9801074 | 0,926892 | A0A1B0GVI | BEGAIN |
| 3,41742 | -2,84327 | -1,15576 | -1,15576 | 3,2392876 | 0,926896 | E9PCE7;E9F | TTC3 |
| -1,80339 | 1,14716 | 0,383631 | 0,383631 | 1,5314363 | 0,927523 | Q9BXW7;A: | HDHD5 |
| 0,0646689 | 2,44645 | -2,98814 | 0,0646689 | 2,7241908 | 0,928694 | K7EPE7;Q9 | VMP1 |
| 0,696965 | -1,51005 | 0,595023 | 0,595023 | 1,2458357 | 0,928726 | P46108;I3L. | CRK |

| | | | | | | | |
|---|---|---|---|---|---|---|---|
| 0,210796 | -3,46574 | 3,89821 | 0,210796 | 3,6819763 | 0,928858 | Q96F07;A0. | CYFIP2 |
| -1,74804 | 1,37331 | 0,658987 | 0,658987 | 1,6353828 | 0,929217 | Q9Y678;H0 | COPG1 |
| -2,49653 | 1,09327 | 1,80381 | 1,09327 | 2,3052279 | 0,929243 | A6NI56;B7Z | CCDC154 |
| -0,752269 | 2,5032 | -2,16592 | -0,752269 | 2,3943395 | 0,929418 | O15344;C9. | MID1 |
| -0,946768 | 0,808433 | -0,0135072 | -0,013507 | 0,8781887 | 0,929588 | P07197;E7E | NEFM |
| -1,55898 | 1,34089 | -0,0321235 | -0,032124 | 1,450615 | 0,929757 | P31323 | PRKAR2B |
| -0,266581 | -0,269069 | 0,463688 | -0,266581 | 0,4223411 | 0,930607 | P23527;Q1( | HIST1H2BO |
| -2,54219 | 0,78338 | 1,39868 | 0,78338 | 2,1200812 | 0,93082 | O60293 | ZFC3H1 |
| -2,48396 | -3,44389 | 6,89669 | -2,48396 | 5,7132252 | 0,930935 | A0A0U1RQ. | CBL |
| -0,917954 | -0,570996 | 1,28823 | -0,570996 | 1,1863353 | 0,931093 | Q9Y266;A0. | NUDC |
| -0,56038 | -2,0347 | 3,03482 | -0,56038 | 2,6076527 | 0,931318 | P36543;C9J | ATP6V1E1 |
| -0,874743 | 0,936177 | 0,0905985 | 0,0905985 | 0,9061198 | 0,931663 | Q63ZY3 | KANK2 |
| 0,216165 | -1,36483 | 1,37662 | 0,216165 | 1,3760904 | 0,932528 | Q9NZI5 | GRHL1 |
| -0,870266 | -0,0824124 | 0,813492 | -0,082412 | 0,8424566 | 0,932704 | P05165;A0/ | PCCA |
| 0,393112 | -1,13874 | 0,922168 | 0,393112 | 1,0703436 | 0,932818 | Q9ULH0;A0 | KIDINS220 |
| -1,14642 | 1,71738 | -0,328106 | -0,328106 | 1,4750707 | 0,932936 | P19338;H7I | NCL |
| -0,365584 | -1,18599 | 1,80534 | -0,365584 | 1,5456408 | 0,933124 | O95155 | UBE4B |
| -0,0925953 | 0,977083 | -1,05072 | -0,092595 | 1,0144128 | 0,933249 | Q99460;A0. | PSMD1 |
| -2,85664 | 1,9926 | 1,2926 | 1,2926 | 2,6211105 | 0,933398 | Q8TEK3;A0. | DOT1L |
| -1,85503 | -1,78615 | 4,19932 | -1,78615 | 3,4757673 | 0,934583 | F5GZI8 | TCP1 |
| -1,39975 | 2,083 | -0,987798 | -0,987798 | 1,9030262 | 0,934805 | Q01130;J3K | SRSF2 |
| 0,373069 | -0,250966 | -0,0709135 | -0,070914 | 0,3211851 | 0,935072 | A0A0B4J20 | PRPS1L1 |
| -3,05962 | 0,879898 | 2,64445 | 0,879898 | 2,920327 | 0,935169 | Q9H2P0;A0 | ADNP |
| 0,374152 | -0,505155 | 0,0612561 | 0,0612561 | 0,4457028 | 0,936244 | Q8TD57;Q5 | DNAH3 |
| -2,46982 | 0,554911 | 2,29093 | 0,554911 | 2,4092703 | 0,936411 | Q9UBT2;K7 | UBA2 |
| -1,48435 | 0,408153 | 0,881979 | 0,408153 | 1,2520378 | 0,936799 | P61247;D6I | RPS3A |
| -1,25997 | 0,21154 | 0,880556 | 0,21154 | 1,0950476 | 0,937537 | Q13442;F8Y | PDAP1 |
| -1,40882 | 1,18486 | 0,0247862 | 0,0247862 | 1,2992417 | 0,937539 | Q15911 | ZFHX3 |
| -2,43832 | 2,33433 | -0,255812 | -0,255812 | 2,3892246 | 0,938636 | Q6IA86 | ELP2 |
| 4,8603 | -2,06538 | -3,4645 | -2,06538 | 4,457669 | 0,938793 | P14136;A0/ | GFAP |
| 1,72209 | 1,03491 | -3,14485 | 1,03491 | 2,6340626 | 0,939996 | Q96L73;D6I | NSD1 |
| -1,78767 | 2,14097 | -0,649971 | -0,649971 | 2,0214649 | 0,940192 | Q9P015 | MRPL15 |
| 0,106004 | 0,440178 | -0,626154 | 0,106004 | 0,5454038 | 0,940246 | P17661 | DES |
| 0,279873 | -0,8192 | 0,43901 | 0,279873 | 0,6851251 | 0,94033 | P02794;G3Y | FTH1 |
| -1,76126 | 0,371027 | 1,16857 | 0,371027 | 1,5147402 | 0,940364 | O75533;B4 | SF3B1 |
| 0,38099 | 2,04001 | -2,77844 | 0,38099 | 2,4478496 | 0,940492 | P0DP23;P0I | CALM1 |
| 0,919936 | -0,984709 | 0,204841 | 0,204841 | 0,9621211 | 0,940671 | Q8IYD8;H0Y | FANCM |
| -0,992595 | 0,226668 | 0,905692 | 0,226668 | 0,9618705 | 0,940784 | E9PNW5;A( | C4orf50 |
| -1,43786 | -0,284558 | 1,50771 | -0,284558 | 1,4842907 | 0,941049 | Q8NEV8;E9 | EXPH5 |
| 0,725262 | -0,346073 | -0,472134 | -0,346073 | 0,6579523 | 0,942425 | Q7L7V1 | DHX32 |
| 0,897147 | -2,21874 | 1,60898 | 0,897147 | 2,0358004 | 0,942466 | H8Y6P7;P0( | GCOM1 |
| 0,497913 | -1,73761 | 1,46811 | 0,497913 | 1,6439529 | 0,94337 | H3BQX2;H3 | PDP2 |
| 1,63545 | -2,20345 | 0,846159 | 0,846159 | 2,0273237 | 0,944073 | Q8IXI1 | RHOT2 |
| 0,632486 | 1,72657 | -2,09003 | 0,632486 | 1,9653477 | 0,944205 | Q01955;H7 | COL4A3 |
| 1,27084 | -2,95531 | 2,05222 | 1,27084 | 2,6940136 | 0,944357 | A0AVI2;A0/ | FER1L5 |
| -1,27065 | -2,59345 | 4,36689 | -1,27065 | 3,6963493 | 0,944554 | Q92769;E5I | HDAC2 |
| -0,344038 | -0,213898 | 0,496477 | -0,213898 | 0,4524074 | 0,944625 | P54296 | MYOM2 |
| 1,09285 | -2,57406 | 1,1928 | 1,09285 | 2,1465264 | 0,94523 | Q6P1X5;H0 | TAF2 |
| 5,03523 | -3,49223 | -0,954954 | -0,954954 | 4,3786917 | 0,945256 | P01023;F5H | A2M |
| 1,99039 | -0,304584 | -1,94928 | -0,304584 | 1,9787593 | 0,945722 | Q96RY5;J3( | CRAMP1 |
| -0,212599 | -0,668714 | 0,782524 | -0,212599 | 0,7421144 | 0,945735 | Q5T5U3;A0 | ARHGAP21 |
| -1,96389 | 1,43095 | 0,771589 | 0,771589 | 1,8001176 | 0,945958 | Q9Y243;F8Y | AKT3 |

| | | | | | | | |
|---|---|---|---|---|---|---|---|
| 1,11925 | -1,04137 | -0,221942 | -0,221942 | 1,0907594 | 0,946159 | P11234;C9J | RALB |
| -0,794885 | -0,0802814 | 0,773596 | -0,080281 | 0,7852704 | 0,947269 | A0A0A0MT | CUL2 |
| -0,220469 | 3,38729 | -2,77134 | -0,220469 | 3,0943925 | 0,947895 | Q6NZY4 | ZCCHC8 |
| 0,902453 | -0,977957 | 0,19532 | 0,19532 | 0,9497857 | 0,948568 | Q8IVG5 | SAMD9L |
| 2,09117 | -2,13961 | -0,218046 | -0,218046 | 2,1183479 | 0,94871 | Q8NFC6;Q1 | BOD1L1 |
| -2,5525 | 0,784548 | 1,49686 | 0,784548 | 2,1618123 | 0,948874 | Q96LB3 | IFT74 |
| -0,984457 | 0,95367 | -0,0901032 | -0,090103 | 0,970023 | 0,949187 | Q8NEZ4;H0 | KMT2C |
| -0,855527 | 2,06742 | -1,44362 | -0,855527 | 1,8804642 | 0,949756 | A4D1E1;A0 | ZNF804B |
| -2,57052 | 1,03192 | 1,27411 | 1,03192 | 2,1531918 | 0,949916 | O76003 | GLRX3 |
| -0,922037 | 0,756919 | 0,270222 | 0,270222 | 0,8638334 | 0,950389 | P00451 | F8 |
| -2,77773 | 1,70357 | 0,788515 | 0,788515 | 2,3677514 | 0,950809 | O60343 | TBC1D4 |
| -3,35423 | 2,43771 | 1,28635 | 1,28635 | 3,0661371 | 0,950818 | A8CG34 | POM121C |
| 1,79592 | 0,182768 | -2,22006 | 0,182768 | 2,0208883 | 0,951299 | Q9BX82 | ZNF471 |
| -0,841192 | 0,352808 | 0,579026 | 0,352808 | 0,7630891 | 0,951563 | H0Y326 | SYNE1 |
| -1,65735 | 0,170252 | 1,30959 | 0,170252 | 1,496716 | 0,951638 | Q9NVE7;A0 | PANK4 |
| -0,561568 | 1,13625 | -0,694944 | -0,561568 | 1,0209184 | 0,951965 | Q8WXA9 | SREK1 |
| 0,428873 | 0,215369 | -0,715696 | 0,215369 | 0,6086192 | 0,952125 | Q5TF21;E9I | SOGA3 |
| -2,92478 | 1,46921 | 1,17021 | 1,17021 | 2,4551134 | 0,952602 | F8WJN3;Q1 | CPSF6 |
| 0,943344 | 0,708061 | -1,83001 | 0,708061 | 1,5377828 | 0,952637 | Q9NZL4 | HSPBP1 |
| -1,15855 | -0,793414 | 2,16283 | -0,793414 | 1,8213671 | 0,95279 | P29373 | CRABP2 |
| 1,20383 | 1,5442 | -2,49341 | 1,20383 | 2,2393351 | 0,953632 | D6REX3;O9 | SEC31A |
| 2,44615 | -2,407 | 0,235873 | 0,235873 | 2,4297862 | 0,953839 | Q9BVV6;A0 | KIAA0586 |
| -1,79401 | -0,520465 | 2,56805 | -0,520465 | 2,2430786 | 0,953898 | Q16695;B4 | HIST3H3 |
| 0,323733 | -2,66346 | 2,07069 | 0,323733 | 2,3939979 | 0,954169 | E7EV03 | MAP2 |
| -0,821914 | 0,963012 | -0,241774 | -0,241774 | 0,9104974 | 0,954905 | Q8IWJ2;B8 | GCC2 |
| 2,20833 | -2,06466 | -0,381123 | -0,381123 | 2,1524408 | 0,955009 | Q14683;G8 | SMC1A |
| 5,94416 | -1,68376 | -3,70118 | -1,68376 | 5,0873656 | 0,955169 | Q92766 | RREB1 |
| 1,08206 | 0,144988 | -1,36259 | 0,144988 | 1,23337 | 0,955184 | A0A1B0GTS | OBSCN |
| -2,44408 | 0,721842 | 1,97132 | 0,721842 | 2,2759619 | 0,955365 | A0A0A0MS | ARHGAP5 |
| -1,31862 | -1,58012 | 3,18797 | -1,31862 | 2,6805602 | 0,955993 | Q99941 | ATF6B |
| 1,46378 | -2,14485 | 0,890315 | 0,890315 | 1,9392137 | 0,955993 | A2RUB6;C9 | CCDC66 |
| 3,44762 | -0,4604 | -2,65388 | -0,4604 | 3,0906385 | 0,956012 | Q3KQU3 | MAP7D1 |
| -0,649152 | -1,0236 | 1,52445 | -0,649152 | 1,3758219 | 0,956035 | Q08257;A6 | CRYZ |
| 0,0220708 | 1,87113 | -1,70362 | 0,0220708 | 1,7877298 | 0,956748 | A0A0A0MR | ASAP1 |
| -2,06429 | 0,528179 | 1,34821 | 0,528179 | 1,7813152 | 0,956974 | Q8WVY7 | UBLCP1 |
| -0,608916 | -0,00937188 | 0,55705 | -0,009372 | 0,5830614 | 0,957162 | Q9ULE4 | FAM184B |
| 0,338727 | 0,732213 | -0,977056 | 0,338727 | 0,8951459 | 0,957222 | P62899;B7Z | RPL31 |
| 1,8279 | -0,829497 | -0,839043 | -0,829497 | 1,537012 | 0,95771 | Q9ULV0 | MYO5B |
| 0,136194 | 0,347645 | -0,441875 | 0,136194 | 0,4087006 | 0,958119 | Q9NYB0 | TERF2IP |
| 1,26923 | -3,08376 | 1,55153 | 1,26923 | 2,5985293 | 0,958716 | Q9UKA9 | PTBP2 |
| -2,49999 | 6,79333 | -3,71409 | -2,49999 | 5,7481259 | 0,958895 | L8E6V6 | LAMA3 |
| 1,44879 | 1,03579 | -2,71278 | 1,03579 | 2,2927789 | 0,9594 | Q9BUQ8;H0 | DDX23 |
| -1,52647 | 0,952477 | 0,444495 | 0,444495 | 1,3094482 | 0,95966 | P32926 | DSG3 |
| 0,49265 | -2,50755 | 2,24961 | 0,49265 | 2,4055034 | 0,960198 | Q14669;C9 | TRIP12 |
| -2,46242 | 3,11422 | -0,377249 | -0,377249 | 2,817718 | 0,960253 | O95819;A0 | MAP4K4 |
| -0,190359 | 0,378323 | -0,15711 | -0,15711 | 0,3191638 | 0,960565 | Q13153;B3 | PAK1 |
| 0,92516 | -0,634839 | -0,213915 | -0,213915 | 0,8070797 | 0,96138 | P49189 | ALDH9A1 |
| -2,17692 | 0,928909 | 1,0745 | 0,928909 | 1,8366229 | 0,961461 | P07919 | UQCRH |
| 1,24457 | -2,89526 | 1,89548 | 1,24457 | 2,5984953 | 0,961568 | P53355;F8V | DAPK1 |
| -2,19579 | 0,0449126 | 1,95708 | 0,0449126 | 2,0785998 | 0,961965 | Q9Y224;G3 | RTRAF |
| 2,1089 | -2,69426 | 0,816647 | 0,816647 | 2,4855158 | 0,962038 | P22735;H0Y | TGM1 |
| -0,895137 | 2,28632 | -1,58284 | -0,895137 | 2,0641787 | 0,962121 | B7ZAQ6;P0 | GPR89A |

| | | | | | | | |
|---|---|---|---|---|---|---|---|
| -0,187659 | 1,41055 | -1,10481 | -0,187659 | 1,2729542 | 0,96216 | P26373;J3C | RPL13 |
| -3,30372 | 2,12195 | 1,45283 | 1,45283 | 2,9583324 | 0,962619 | P42785;E9F | PRCP |
| 0,83268 | -1,07919 | 0,336305 | 0,336305 | 0,9920737 | 0,963072 | E9PCX7;D6I | NNT |
| -0,747654 | -0,516877 | 1,36889 | -0,516877 | 1,1611154 | 0,963332 | P11277 | SPTB |
| 0,243992 | -0,466066 | 0,186895 | 0,186895 | 0,394504 | 0,963619 | Q9UNH7;A( | SNX6 |
| -1,24373 | 0,281667 | 1,06672 | 0,281667 | 1,1748279 | 0,963657 | Q9NZJ4 | SACS |
| -1,27775 | 2,23166 | -0,784767 | -0,784767 | 1,8999045 | 0,963678 | A6H900;B1. | AXDND1 |
| -0,204469 | 0,751274 | -0,60823 | -0,204469 | 0,6981784 | 0,964105 | O94813;X6I | SLIT2 |
| 2,81529 | -1,43686 | -1,59853 | -1,43686 | 2,5029557 | 0,964123 | Q9NR28;AC | DIABLO |
| -2,16777 | 1,07215 | 1,26237 | 1,07215 | 1,927828 | 0,964712 | Q5XKE5 | KRT79 |
| -3,33169 | 3,0421 | 0,561585 | 0,561585 | 3,2128841 | 0,96546 | Q8NHU2;C9 | CFAP61 |
| -0,672634 | 3,09657 | -2,19356 | -0,672634 | 2,7235268 | 0,965489 | Q9Y239;G3 | NOD1 |
| -1,27498 | 1,77815 | -0,638591 | -0,638591 | 1,6107571 | 0,965696 | P14550;Q5 | AKR1A1 |
| 5,98626 | -1,76905 | -3,78357 | -1,76905 | 5,1583703 | 0,965701 | P32019 | INPP5B |
| -0,822321 | 0,969571 | -0,222702 | -0,222702 | 0,9121344 | 0,966249 | P35813;B5E | PPM1A |
| -2,18142 | 1,60596 | 0,415638 | 0,415638 | 1,9367423 | 0,96633 | C9JEZ4;Q9L | CDC42EP3 |
| -2,40382 | 2,25021 | -0,0384237 | -0,038424 | 2,3271205 | 0,96633 | Q8IY21;D6F | DDX60 |
| 0,097485 | 0,602517 | -0,754982 | 0,097485 | 0,6861196 | 0,967304 | O43776;K7I | NARS |
| 1,91672 | -1,07274 | -0,713303 | -0,713303 | 1,6321298 | 0,967332 | Q15233;C9 | NONO |
| -0,922815 | 2,33444 | -1,57843 | -0,922815 | 2,0956341 | 0,967523 | A0AUZ9 | KANSL1L |
| 0,0821003 | 0,661072 | -0,691507 | 0,0821003 | 0,6786195 | 0,968934 | O43290 | SART1 |
| 0,87386 | -2,75899 | 2,07679 | 0,87386 | 2,5175851 | 0,968936 | P00390;A0/ | GSR |
| -2,98892 | 1,31134 | 1,87779 | 1,31134 | 2,6613896 | 0,969304 | H3BQZ9;P0 | APRT |
| 0,367624 | -1,45033 | 0,990137 | 0,367624 | 1,2680929 | 0,970211 | Q8N4C8 | MINK1 |
| -0,8973 | -3,31032 | 3,93853 | -0,8973 | 3,6912903 | 0,970252 | Q96I99;E9P | SUCLG2 |
| -2,37785 | -2,29048 | 4,39258 | -2,29048 | 3,8839337 | 0,971027 | O43252 | PAPSS1 |
| 2,10503 | -1,36435 | -0,873199 | -0,873199 | 1,8773951 | 0,971195 | P07864;F5H | LDHC |
| 0,885209 | 0,546113 | -1,52262 | 0,546113 | 1,3033472 | 0,971415 | Q5T9S5;J3K | CCDC18 |
| -2,67802 | 1,13954 | 1,38036 | 1,13954 | 2,2767743 | 0,97166 | Q92823;F8V | NRCAM |
| 1,10618 | -0,7865 | -0,252683 | -0,252683 | 0,9758506 | 0,971985 | A5A3E0 | POTEF |
| 0,455164 | 1,29723 | -1,86284 | 0,455164 | 1,6364728 | 0,972458 | K7EK35;P42 | STAT5A |
| 0,950502 | -0,545785 | -0,349969 | -0,349969 | 0,8132695 | 0,972528 | Q15393;I3L | SF3B3 |
| 1,99584 | -3,05859 | 1,24604 | 1,24604 | 2,727615 | 0,972578 | P22748;K7E | CA4 |
| 1,35967 | -0,134631 | -1,14135 | -0,134631 | 1,2584064 | 0,972863 | Q5T655 | CFAP58 |
| 0,556683 | -1,87132 | 1,20852 | 0,556683 | 1,6230375 | 0,973317 | A0A087WU | LDHAL6A |
| 2,44073 | 1,1383 | -3,38078 | 1,1383 | 3,0552792 | 0,97352 | M0QXL7 | ZNF208 |
| -2,07598 | 1,02067 | 1,17329 | 1,02067 | 1,833498 | 0,973739 | P16435;H0Y | POR |
| 0,251374 | 0,0141495 | -0,250077 | 0,0141495 | 0,2508466 | 0,97487 | Q8TAQ5;K7 | ZNF420 |
| -0,00054229 | -2,94481 | 2,77206 | -0,000542 | 2,8588645 | 0,975261 | Q9ULK5 | VANGL2 |
| 2,95822 | 0,565937 | -3,33195 | 0,565937 | 3,1749745 | 0,975292 | Q9NSV4 | DIAPH3 |
| 1,79835 | 1,84029 | -3,83195 | 1,79835 | 3,2628296 | 0,975819 | P49815;A0/ | TSC2 |
| -0,373014 | 0,210679 | 0,181635 | 0,181635 | 0,3289318 | 0,976053 | Q5VTR2;C9 | RNF20 |
| -0,488004 | -0,958653 | 1,52036 | -0,488004 | 1,3165955 | 0,977154 | O94855 | SEC24D |
| -0,0284116 | 1,75936 | -1,83109 | -0,028412 | 1,7952302 | 0,977234 | Q00796;H0 | SORD |
| 1,81601 | -2,94068 | 1,26844 | 1,26844 | 2,6026469 | 0,977453 | Q13332;A0. | PTPRS |
| 3,36002 | -2,7528 | -0,437947 | -0,437947 | 3,0862509 | 0,977615 | Q6AWC2;D | WWC2 |
| 1,91672 | 0,0622647 | -2,08824 | 0,0622647 | 2,0043029 | 0,977753 | P52742;M0 | ZNF135 |
| 0,0126399 | -1,93756 | 1,82446 | 0,0126399 | 1,8814341 | 0,978208 | Q8N7M2 | ZNF283 |
| 0,851627 | -1,86741 | 0,930916 | 0,851627 | 1,5932188 | 0,978259 | H3BM40;H3 | NDRG4 |
| 0,112971 | 0,543123 | -0,689049 | 0,112971 | 0,6253685 | 0,978492 | Q8N5A5;V9 | ZGPAT |
| -0,32292 | 0,114539 | 0,223602 | 0,114539 | 0,2892379 | 0,978522 | Q15006;E5I | EMC2 |
| 1,74326 | -0,229714 | -1,42927 | -0,229714 | 1,6019003 | 0,978527 | Q16799;Q2 | RTN1 |

| | | | | | | | |
|---|---|---|---|---|---|---|---|
| -3,05817 | 1,66097 | 1,2607 | 1,2607 | 2,6167135 | 0,978708 | P52943;H0Y | CRIP2 |
| 0,421519 | -3,1099 | 2,84289 | 0,421519 | 2,993595 | 0,978933 | P29375 | KDM5A |
| 2,25895 | -3,09767 | 0,701503 | 0,701503 | 2,7553806 | 0,979672 | A0A1B0GV9 | HSD17B12 |
| -1,61296 | -0,985689 | 2,7133 | -0,985689 | 2,3378236 | 0,979983 | Q5T0R9 | CAP1 |
| 0,222924 | -0,6504 | 0,400273 | 0,222924 | 0,5624443 | 0,980259 | Q6YP21 | KYAT3 |
| -2,35471 | 2,0678 | 0,392041 | 0,392041 | 2,2327638 | 0,980781 | H0YDU8;P5 | PPP5C |
| 0,845956 | -0,0810788 | -0,802812 | -0,081079 | 0,8265116 | 0,981266 | P23497 | SP100 |
| 2,41741 | -1,96215 | -0,557316 | -0,557316 | 2,2361835 | 0,981372 | K7EMM8;Q | GLYR1 |
| 1,94921 | -0,908154 | -1,11748 | -0,908154 | 1,7133269 | 0,981794 | Q9C0C2;A0 | TNKS1BP1 |
| -2,57009 | 2,66474 | 0,0198948 | 0,0198948 | 2,6174629 | 0,982139 | P42892;B4I | ECE1 |
| -0,00697179 | 2,55482 | -2,66054 | -0,006972 | 2,6078146 | 0,98236 | P39687;H0Y | ANP32A |
| -0,201161 | 1,28558 | -1,13649 | -0,201161 | 1,2214515 | 0,982598 | Q9C0B1;A0 | FTO |
| -2,48382 | 0,814422 | 1,76155 | 0,814422 | 2,2285503 | 0,983121 | P39023;H7( | RPL3 |
| 5,47493 | -2,8904 | -2,77875 | -2,77875 | 4,7978197 | 0,983475 | A0A0J9YY0 | MYO15B |
| -1,29584 | 0,389966 | 0,860155 | 0,389966 | 1,1336765 | 0,983538 | P62280;M0 | RPS11 |
| -2,60231 | 2,73507 | -0,026062 | -0,026062 | 2,6692236 | 0,983683 | P53618;E9F | COPB1 |
| 0,350594 | -1,06144 | 0,747828 | 0,350594 | 0,9508843 | 0,984122 | Q9P2Q2;Q5 | FRMD4A |
| 1,29304 | -0,811999 | -0,52539 | -0,52539 | 1,1416378 | 0,984145 | A0A2R8Y7E | CCDC88A |
| 1,12354 | -0,39668 | -0,765745 | -0,39668 | 1,0013887 | 0,98415 | Q99747;J3K | NAPG |
| 0,550922 | -0,0785332 | -0,491563 | -0,078533 | 0,5249734 | 0,985091 | Q4FZB7 | KMT5B |
| -1,63432 | 2,67895 | -0,962302 | -0,962302 | 2,3207263 | 0,98552 | P10253 | GAA |
| 2,48971 | -3,43507 | 1,04937 | 1,04937 | 3,0899784 | 0,986259 | Q9BUY5 | ZNF426 |
| -1,77829 | -2,67954 | 4,5882 | -1,77829 | 3,9615757 | 0,986566 | Q9UJY4;H3 | GGA2 |
| -0,720505 | -1,62752 | 2,2808 | -0,720505 | 2,0455428 | 0,986585 | Q9NQX4;H( | MYO5C |
| 0,92002 | -1,94134 | 1,07712 | 0,92002 | 1,6991744 | 0,986593 | Q12904;D6 | AIMP1 |
| 7,33605 | -3,50755 | -3,62281 | -3,50755 | 6,2940919 | 0,98666 | Q7Z553;G3 | MDGA2 |
| 2,12307 | -2,80275 | 0,760915 | 0,760915 | 2,5435827 | 0,986962 | Q00526 | CDK3 |
| -0,360918 | 0,696381 | -0,316469 | -0,316469 | 0,5980137 | 0,987034 | Q15435;H7 | PPP1R7 |
| 6,03767 | -3,30041 | -2,57458 | -2,57458 | 5,194507 | 0,987216 | O75419 | CDC45 |
| 3,07071 | -0,574354 | -2,58337 | -0,574354 | 2,8662187 | 0,987606 | Q5JNZ5 | RPS26P11 |
| 1,35531 | -1,31362 | -0,0023374 | -0,002337 | 1,3345321 | 0,987962 | P17252;J3K | PRKCA |
| 3,76527 | -0,189746 | -3,4697 | -0,189746 | 3,6227301 | 0,988075 | Q6UXK2;Q2 | ISLR2 |
| 1,09499 | -1,44956 | 0,317746 | 0,317746 | 1,3039819 | 0,988472 | O00139;D6 | KIF2A |
| -0,645342 | -0,653058 | 1,268 | -0,645342 | 1,1069027 | 0,988787 | Q7L7X3;J3C | TAOK1 |
| -1,89152 | 1,30247 | 0,635267 | 0,635267 | 1,6848039 | 0,988801 | P78318 | IGBP1 |
| -1,27366 | 0,146289 | 1,15944 | 0,146289 | 1,2222047 | 0,989287 | Q9Y6M1;F8 | IGF2BP2 |
| -0,949979 | 1,0701 | -0,145952 | -0,145952 | 1,0170186 | 0,989633 | Q6JQN1;F8 | ACAD10 |
| 1,8365 | 1,42106 | -3,18784 | 1,42106 | 2,788624 | 0,989794 | K7ELQ6 | MISP3 |
| 2,92493 | 0,32267 | -3,17137 | 0,32267 | 3,0590016 | 0,989827 | Q08AN1 | ZNF616 |
| -0,675059 | 0,0148967 | 0,676748 | 0,0148967 | 0,6759522 | 0,989984 | Q8WVV9;B | HNRNPLL |
| 2,55091 | -2,3367 | -0,274183 | -0,274183 | 2,4536999 | 0,990022 | Q6GPH4;C9 | XAF1 |
| 1,16691 | 0,977708 | -2,18923 | 0,977708 | 1,8854252 | 0,990341 | Q96M96;B7 | FGD4 |
| 0,103685 | -1,29828 | 1,22099 | 0,103685 | 1,2623125 | 0,991466 | P35749 | MYH11 |
| 0,154827 | -0,349677 | 0,2011 | 0,154827 | 0,3055107 | 0,991649 | Q9Y6I3 | EPN1 |
| -2,77337 | -0,0678594 | 2,89592 | -0,067859 | 2,8356253 | 0,992127 | K7EK07;P84 | H3F3B |
| -0,375693 | -0,212465 | 0,57943 | -0,212465 | 0,5108818 | 0,993025 | C9J813 | CALD1 |
| 0,324616 | -2,381 | 2,09441 | 0,324616 | 2,253953 | 0,993113 | Q7L8J4 | SH3BP5L |
| 0,328937 | -0,510008 | 0,173649 | 0,173649 | 0,4463425 | 0,993212 | Q4G0X9;I3I | CCDC40 |
| -0,889647 | 1,41027 | -0,540285 | -0,540285 | 1,2393773 | 0,993524 | Q9NQ29;A8 | LUC7L |
| -2,3729 | 0,638968 | 1,70046 | 0,638968 | 2,1130695 | 0,993534 | V9GYY6 | MCAT |
| -1,16103 | -3,4969 | 4,59397 | -1,16103 | 4,1641024 | 0,99373 | G3V2A3;G3 | TUBB3 |
| -0,284833 | 0,0129495 | 0,275956 | 0,0129495 | 0,2805742 | 0,994075 | P00367 | GLUD1 |

| | | | | | | | |
|---|---|---|---|---|---|---|---|
| 0,691656 | 0,530386 | -1,23635 | 0,530386 | 1,0696239 | 0,99454 | Q9NWH9;H | SLTM |
| 1,34477 | -2,7611 | 1,44717 | 1,34477 | 2,4006315 | 0,994754 | Q96SN8;A0 | CDK5RAP2 |
| -0,0705674 | 0,901614 | -0,820984 | -0,070567 | 0,8636749 | 0,995244 | Q5TZA2;B1. | CROCC |
| 0,476512 | 0,566863 | -1,03378 | 0,476512 | 0,8991851 | 0,995645 | P30613 | PKLR |
| 2,54307 | -1,61808 | -0,944966 | -0,944966 | 2,2336311 | 0,996349 | Q9BWM7;A | SFXN3 |
| 0,392519 | -0,511192 | 0,115076 | 0,115076 | 0,4629398 | 0,996828 | Q15819;G3 | UBE2V2 |
| -1,28463 | 0,979933 | 0,296465 | 0,296465 | 1,1615532 | 0,997108 | A0A0A0MS | ZNF300 |
| 1,75063 | 0,525423 | -2,26205 | 0,525423 | 2,0564021 | 0,99722 | Q14117 | DPYS |
| -0,603661 | 1,10664 | -0,496786 | -0,496786 | 0,958082 | 0,997359 | O43242;H0 | PSMD3 |
| 1,08863 | -0,149716 | -0,945436 | -0,149716 | 1,0250281 | 0,997402 | Q7L2R6;D6 | ZNF765 |
| -1,53066 | 2,37295 | -0,854667 | -0,854667 | 2,0861707 | 0,997577 | Q15811;F8' | ITSN1 |
| -2,25395 | -0,0681248 | 2,33298 | -0,068125 | 2,2943068 | 0,998059 | Q86VS3;H3 | IQCH |
| -0,881606 | 0,539842 | 0,338788 | 0,338788 | 0,7692311 | 0,998421 | Q9UQ80;F8 | PA2G4 |
| -1,60925 | 0,559272 | 1,04551 | 0,559272 | 1,4134276 | 0,99871 | Q9NX40;D6 | OCIAD1 |
| -2,39337 | 2,5776 | -0,177204 | -0,177204 | 2,490344 | 0,998848 | Q6ZU65 | UBN2 |
| -1,6943 | -0,547617 | 2,24749 | -0,547617 | 2,0275278 | 0,998877 | Q12792;F8' | TWF1 |
| -1,35236 | 0,00106841 | 1,35319 | 0,0010684 | 1,3527751 | 0,999426 | Q08379;A0. | GOLGA2 |
| 0,449148 | 1,04853 | -1,49932 | 0,449148 | 1,332124 | 0,999498 | P16989 | YBX3 |
| -0,861936 | 1,57386 | -0,711403 | -0,711403 | 1,3649291 | 0,999844 | Q6ZU52;E9 | KIAA0408 |
